# Structural features of the amino acid sequences of Chromista nucleolin-like proteins

**DOI:** 10.1101/2025.03.02.641111

**Authors:** Darya P. Petrova, Nadezhda A. Potapova, Yekaterina D. Bedoshvili

**Author notes:** **Correspondence** Ye. Bedoshvili, Department of Ultrastructure of cell, Limnological Institute, Siberian Branch, Russian Academy of Sciences, 3 Ulan-Batorskaya st., Irkutsk 664033, Russia.

## Abstract

Nucleolins are multifunctional proteins localized in the centrosome and nucleus and involved in the processes of microtubule nucleation. Their structure is highly conserved in organisms that are evolutionarily distant from each other. The phylogeny of the kingdom Chromista was supported by marker genes, cell morphology and cytoskeletal features. In this study we identified general patterns of the nucleolin protein structure characteristic of this group - the structure of the N-terminal and central domains, including bipartite sequences that ensure nuclear localization, as well as RNA recognition motifs. We also noted features in the primary structure of these proteins, which may affect the conformation of the protein molecule. Each of the six clades identified during phylogenetic reconstruction has structural features of the N-terminal domains, bipartite nuclear localization sequences and sequences of RNA recognition motifs. The features we described allow us to classify the studied nucleolin sequences of Chromista as nucleolin-like proteins. In this study, we described for the first time the structure of nucleolin-like proteins in Chromista.

## Introduction

The composition and functioning of the cytoskeleton and the microtubule organizing center (MTOC) are the essential factors reflecting the process of eukaryote evolution and, in particular, unicellular organisms from the kingdom Chromista (Cavalier-Smith, 2018). For eukaryotes the organization of the cytoskeleton, along with the compartmentalization, determines the symmetry of the cell at different stages of the cell cycle and the spatial distribution of the processes occurring in it. In the kingdom Chromista, diatoms (Bacillariophyta) are the most morphologically diverse. As has been shown, in this group of microalgae, the work of the cytoskeleton determines the symmetry and micro- and nanostructure of species-specific silica frustules. Recent searches and analyses (De Martino et al. 2009; Khabudaev et al. 2022; Petrova et al. 2023) have shown that the set of structural proteins that make up the microtubule center in diatoms is characterized by specificity, their features were identified, and an analysis of potential functional activity was carried out. It is known that the composition of the microtubule center in different organisms determines its morphology and functional features (Teixidó-Travesa et al. 2012; Sulimenko et al. 2017; Chumová et al. 2021).

Sequences of potential MTOC components in diatoms were identified in genomes and transcriptomes of large databases (Petrova et al. 2023). As expected, not all known MTOC components were found in diatoms. Of the identified sequences, nucleolins in general stand out for their functional diversity, and preliminary analysis showed that the structure of nucleolins in diatoms differs from those in other model eukaryotes.

Nucleolins are multifunctional phosphoproteins localized in nucleolus, nucleoplasm and cytoplasm and detected in a variety organism, from yeast to plants and mammals. This protein is localized in the nucleolus and binds with ribosome biogenesis (Bouvet et al., 1998); however, the nucleolin functions are much wider. Nucleolins were identified in a mature centrosome (Gaume et al., 2015). The localization of phosphorylated nucleolins in the nucleus and cytoplasm in *Xenopus laevis* oocytes at different stages of centrosome maturation suggests that their functioning is associated with post-translational modifications regulation is associated with phosphorylation and other modifications (Schwab & Dreyer, 1997; Salvetti et al., 2016). It is supposed that nucleolins take part in the interactions of microtubules and kinetochore and stabilization of such interactions (Ma et al., 2007). It is known that nucleolin is encoded by one gene in the genomes of humans (*Homo sapiens*) and other mammals, and also yeast (*Saccharomyces cerevisiae, Schizosaccharomyces pombe*) (Srivastava et al., 1989; Kondo & Inouye, 1992; Gulli et al., 1995). Also, it was found that tetraploid organisms such as *Xenopus laevis* have up to three genes encoding this protein (Messmer & Dreyer, 1993), and in plant genomes two genes were found (Tajrishi et al., 2011). There are nucleolin-like proteins that have structural similarities and low sequence homology with known nucleolins (Gulli et al., 1995; Tong et al., 1997).

Nucleolin protein structure consists of three structural domains, N- and C-terminal and central. It is suggested that terminal domains participate in binding with other proteins while central domain is necessary for specific interactions with nucleic acids (Ginisty et al., 1999). Nucleolin structure is highly conserved in evolutionarily distant organisms and the domain arrangement is similar in yeasts, plants and animals (Figure 1). N-terminal domain has various number of acidic regions in different organisms and is important for protein-protein interactions (Bharti et al., 1996). Central domain includes two RNA Recognition Motifs (RRM) in yeasts and plants and four in animals; it was proposed that the presence of only two RRMs is characteristic of nucleolin-like proteins (Ginisty et al. 1999). RRMs are less conserved within the same protein compared to RRMs of different proteins (Ginisty et al., 1999). C-terminal domain contains GAR (Glycine- and Arginine-Rich) or RGG (Arg-Gly-Gly repeats) domain and it has been suggested that this GAR/RGG sequence is poorly evolutionarily conserved (Ginisty et al., 1999; Tajrishi et al., 2011).

**Figure 1.**
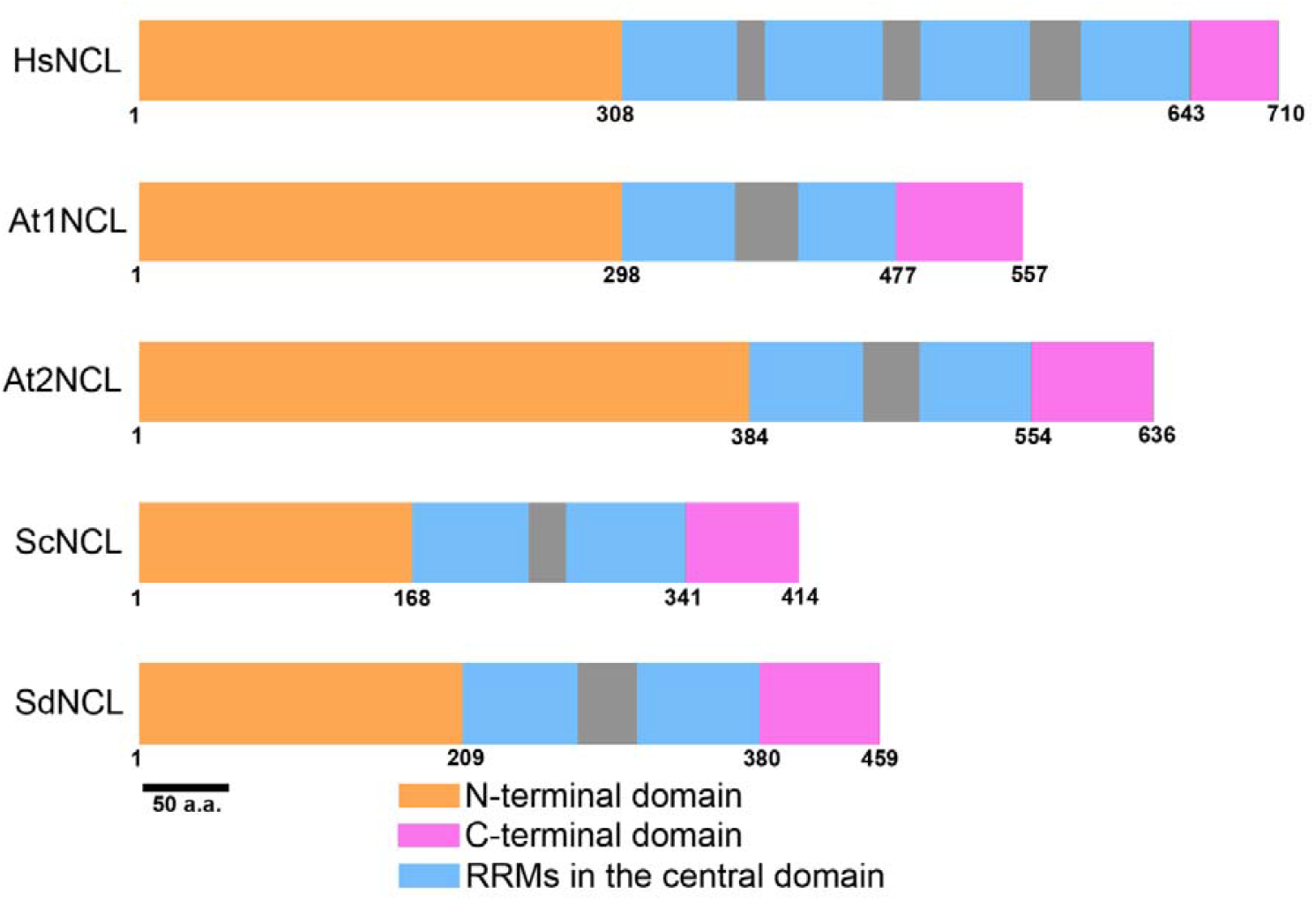
Domain organization of nucleolin amino acid sequences of *Homo sapiens* (HsNCL), *Arabidopsis thaliana* (At1NCL and At2NCL), *Saccharomyces cerevisiae* (ScNCL) and *Skeletonema dohrnii* (SdNCL), as one of the representatives of Chromista.

Analysis of the phylogeny and conserved domains within nucleolins of 15 organisms (González-Camacho & Medina, 2004) showed that nucleolin is a highly conserved protein and that ancestors of the eukaryotes had a gene encoding nucleolin. Also, it was shown that the sequence of that gene was evolved very slowly. Even with a small sample of 15 sequences, the resulting tree was very similar to the taxonomic organization of the represented groups. Currently, single-celled organisms of the kingdom Chromista, which are very diverse in the composition of microorganisms, are being actively studied. Chromista includes all chromophyte algae containing chlorophyll a and c and resulting from the symbiogenic enslavement of another eukaryote (red algae), as well as all heterotrophic protozoa that evolved from them as a result of the loss of photosynthesis or entire plastids (Cavalier-Smith, 1981). Chromista kingdom and their place in the tree of life of Earth’s organisms are mostly studied using phylogenetic approach, revealing global events in the evolution of single-celled organisms.

Since previously the kingdom formally included many protozoan species; to revise Chromista was used both marker genes phylogeny and morphology data, including information on the organization of centrioles in their cells (Cavalier-Smith, 2018). Previously, the morphology of the proximal flagellar microtubules was used for phylogenetic analysis of the Excavata group (Simpson, 2003). Later Cavalier-Smith (2018) examined such details as the symmetry and relative position of the centrioles and the structure of the cell apex, often in chromists having flagella. It is noted that centriole structure has remained conserved for hundreds of millions of years; however, the stability of the structure is alternated by large shifts that give rise to modifications in the microtubular apparatus structure and associated cellular structures. The structures formed by microtubules in chromists can be diverse (Beech & Wetherbee, 1990), but have a common eukaryotic features. The most important structure organizing microtubules in the cell and provide the cell morphology is the microtubule organizing center and its modifications (Zheng et al., 1995).

Among the representatives of the kingdom Chromista, diatoms (Bacillariophyta) are distinguished by morphological diversity and have a special acentriolar microtubular organizing center (De Martino et al. 2009). In this group of microalgae, the functioning of the cytoskeleton has been shown to determine the symmetry and micro- and nanostructure of species-specific silica shells (Pickett-Heaps et al., 1990). Recent studies (De Martino et al. 2009; Khabudaev et al. 2022; Petrova et al. 2023) have shown that the set of structural proteins that make up the microtubule organizing center in diatoms is specific; their features have been identified and potential functional activity has been analyzed. As expected, not all known microtubule organizing center components were found in diatoms, and preliminary analysis of nucleolins showed the presence of distinctive structural features, in particular serine and lysine repeats in the N-terminal domain. The present study was made to test whether these features are truly unique only to diatoms or to the entire Chromista.

The presence of nucleolins in the microtubule organizing center, as well as their belonging to the group of conserved proteins necessary in the processes of microtubule nucleation (Teixidó-Travesa et al., 2012; Sulimenko et al., 2017; Chumová et al., 2021), suggests their important role in the cell cycle and accompanying processes, such as valve morphogenesis of diatoms, a large and morphologically diverse class of the kingdom Chromista. Chromista are a very large and well-studied group of organisms. However, data on their evolution, systematics and morphology remain in demand. In this regard, we believe that the study of the components of the chromist microtubule organizing center can expand the understanding of evolution and cytology both in these organisms and in general. In this work, we undertook a search for nucleolins in diatoms and other species of the kingdom Chromista. Comparative analysis of the predicted amino acid sequences in silico allowed us to identify the characteristic features of nucleolins in these single-celled organisms.

## Materials and methods

The search of the nucleolin sequences was performed in the set of amino acid sequences predicted according to published genomic data in Nucleotide NCBI and transcriptomic data in MMETSP (Marine Microbial Eukaryote Transcriptome Sequencing Project – Keeling et al., 2014) using BLAST+ (Camacho et al. 2009). As a query, the sequences of *Homo sapiens* (NP_005372.2), *Arabidopsis thaliana* (NP_001185316.1) and *Saccharomyces cerevisiae* (NP_011675.1) were used. All obtained sequences were searched against the NCBI nr database (NCBI Resource Coordinators, 2018) with default parameters to remove contaminants and sequences not identified as nucleolins or nucleolin-like proteins. The data set was manually curated to remove sequences without nucleolin specific domains.

Alignment and analysis of the predicted amino acid sequences were performed using ClustalW BioEdit v. 7.2.5 (Hall, 1999). Only 48 completely identified sequences of Chromista were selected for phylogenetic analysis. We used the maximum likelihood method with WAG+F evolutionary model as best model according to BIC/AICc with 1000 botstrap recplications in the MEGA 7.0 program (Kumar et al., 2016a). Evolutionary model is a model which in the best possible way predicts the rules on how the sequence evolve and accumulate mutations. Here WAG+F is a Whelan and Goldman model and F is F-distributed rates. BIC and AICc characteristics are measured in a process of model selection and give an opportunity to select the best one (Aho et al., 2014)

We used SMART service (Letunic & Bork, 2017) for identification of RRM and coil-coil domains and NetPhos-3.1 (Blom et al., 2004) for analysis of predicted phosphorylation sites. α-helices were calculated using an online server PROTEUS2 (Montgomerie et al., 2008). Homologous 3D-modeling of protein molecules was carried out using the SWISS-MODEL server (Waterhouse et al., 2018). We carried out a search of conserved elements according to data published earlier (Ginisty et al., 1999; Tajrishi et al., 2011).

## Results and discussion

As a result of the database search, 172 sequences were found, however, after a detailed analysis of their structure, we excluded 66 sequences from further analysis, since structures described for nucleolins were not identified in the structure of their N- and C-terminal domains (Tuteja & Tuteja, 1998). The presence of RRMs, also known as RBD (RNA binding domain), in the structure of a protein is not the only necessary condition for classifying it as a member of the nucleolin family. These motifs may be included of various proteins involved in RNA metabolism (Cléry & Frédéric, 2013). Thus, the final sample included 106 sequences, of which 83 belonged to 71 species of Bacillariophyta and 23 sequences to 22 species of other chromists (Supplementary table S1; Supplementary data_1). The length of predicted amino acid sequences of Chromista nucleolins varies from 104 to 789 amino acids (a.a.) and is 460-484 a.a. in most sequences. This length is comparable to the nucleolin-like protein of *S. cerevisiae* (414 a.a.) (Supplementary table S1).

That there is only one gene of nucleolin in the genome of *Homo sapiens* whereas *Arabidopsis thaliana* has two genes encoding nucleolin-like proteins (Srivastava et al., 1990; Ginisty et al. 1999; Pontvianne et al., 2007). In our study, we find eight species with two sequences and two diatoms that have three genes of nucleolin-like proteins: *Chaetoceros tenuissimus* and *Fistulifera solaris* (Supplementary table S1). Nucleolin sequences of N- and C-terminal domains of Chromista have low identity (no more than 9 %) with model organisms (*H. sapiens, A. thaliana* and *S. cerevisiae*); the identity of central domain increase to 10-15 %. The comparison of sequences within one species showed that identity did not exceed 20 %. Only the diatoms *Nitzschia inconspicua* and *Perkinsus olseni* (class Perkinsea) had identity above 80 %; whereas identity between two isoforms of *Arabidopsis thaliana* was 47,4%. Among some genera identity increased to 90 %. Within Bacillariophyta identity varied from 5 to 80 % and identity increase was obvious in all three domains and generally throughout the entire length of the sequence. It is noted that nucleolins have high homology that is conserved during evolution possibly because of important nucleolin functions (Tuteja & Tuteja, 1998).

Only fully identified sequences of nucleolins and nucleolin-like proteins were used for phylogenetic analysis as our main goal was to reveal features that were specific both for fragments (short sequences without start and stop codons, as well as some specific nucleolin domains) and all protein molecules (Supplementary data_2). It was shown that six clades are formed on the tree (Figure 2). Clades 1-3 include different Bacillariophyta classes; clade 4 consists of nucleolins of reference group (one sequence of *H. sapiens*, two sequences of *A. thaliana* and one sequence of *S. cerevisiae*) and species of Ciliophora and Haptophyta. Clades 5 and 6 include sequences of Ochrophyta and Cryptophyta species and Bacillariophyta and Bolidopyceae species, respectively (Figure 2).

**Figure 2.**
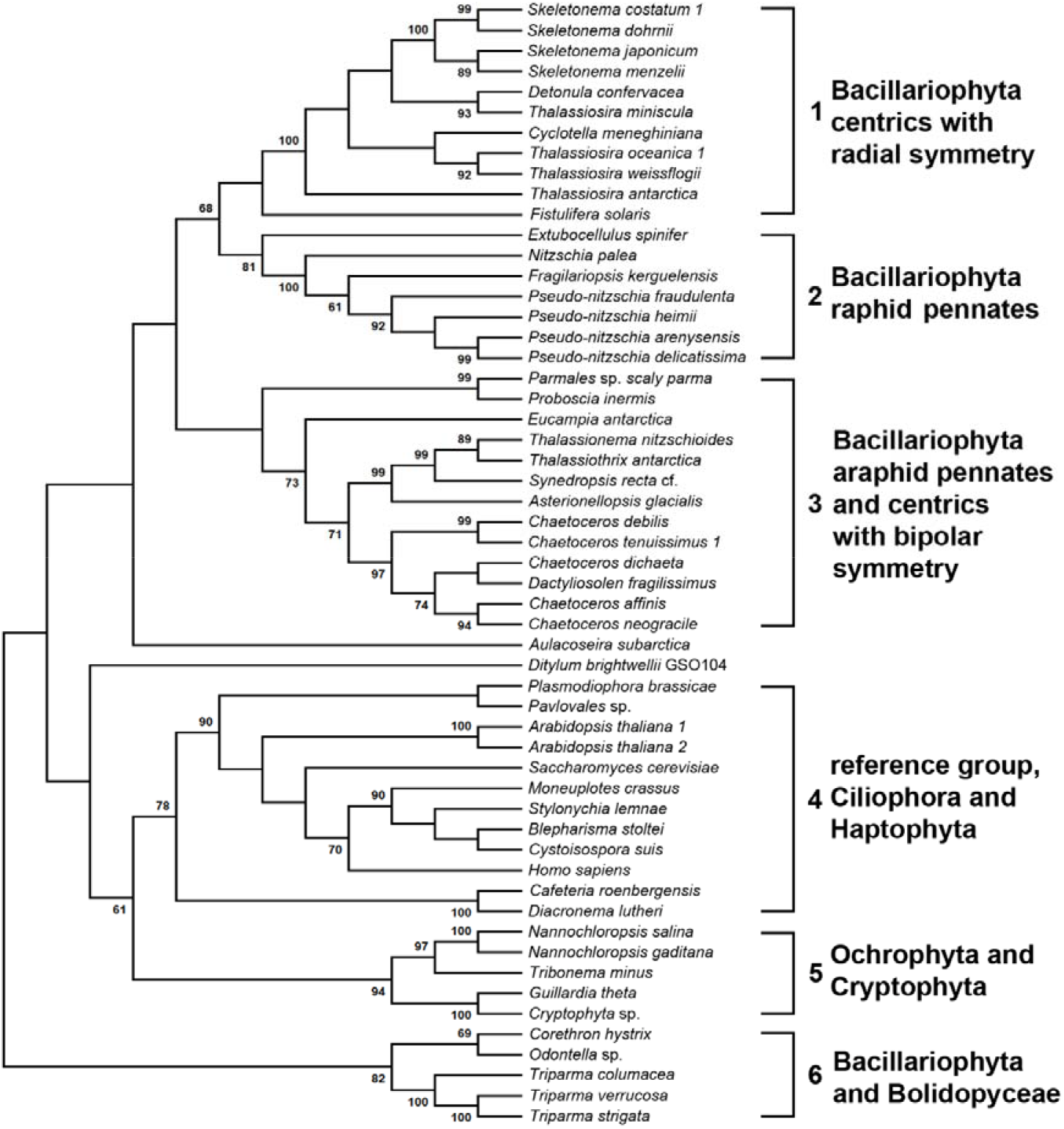
Phylogenetic reconstruction of Chromista nucleolins.

The N-terminal domain has a different length in nucleolins of different organisms. For instance, the *H. sapiens* nucleolin has an N-terminal domain with a length of 307 a.a., *Saccharomyces cerevisiae* – 168 a.a., and in different Chromista species the length of this domain varies from 202-336 a.a. (Figure 1; Supplementary Table S1). The N-terminal domain has the greatest length of 473 a.a. in *Cystoisospora suis* (Ciliophora). However, the length of N-terminal domain did not depend on the clade according to phylogenetic reconstruction (Figure 2). It is known that the N-terminal domain of nucleolins include several highly charged fragments formed by acidic a.a. repeats that alternate with the basic residues. The number of acidic regions varied in different organisms; for example, there are four such regions in the *H. sapiens* sequence, seven – in nucleolin-like proteins of *A. thaliana* and four – in *S. cerevisiae*. In nucleolins of mammals the number of glutamate (E), aspartate (D) and serin (S) residues in highly charged acidic repeats can reach several dozen; in plant nucleolin-like proteins, the number of such repeats is greater than in mammal nucleolins, but they are shorter in length and are represented by several E and D residues (González-Camacho & Medina, 2004). Analysis of the sequences within each clade showed that there are 5 short highly charged acidic repeats in the sequences of clades 1-3 (Figure 3, Supplementary data_2). The maximum number of such repeats was 10 and was noted for sequences of clade 5. The longest repeats are found in clade 4, which includes reference group and the *Diacronema lutheri* (NCLDlu) sequence, in the N-terminal part of which a fragment of twenty-five residues D is localized (Supplementary data_2). The length of repeats in other clades are 3-7 a.a. (Figure 3).

**Figure 3.**
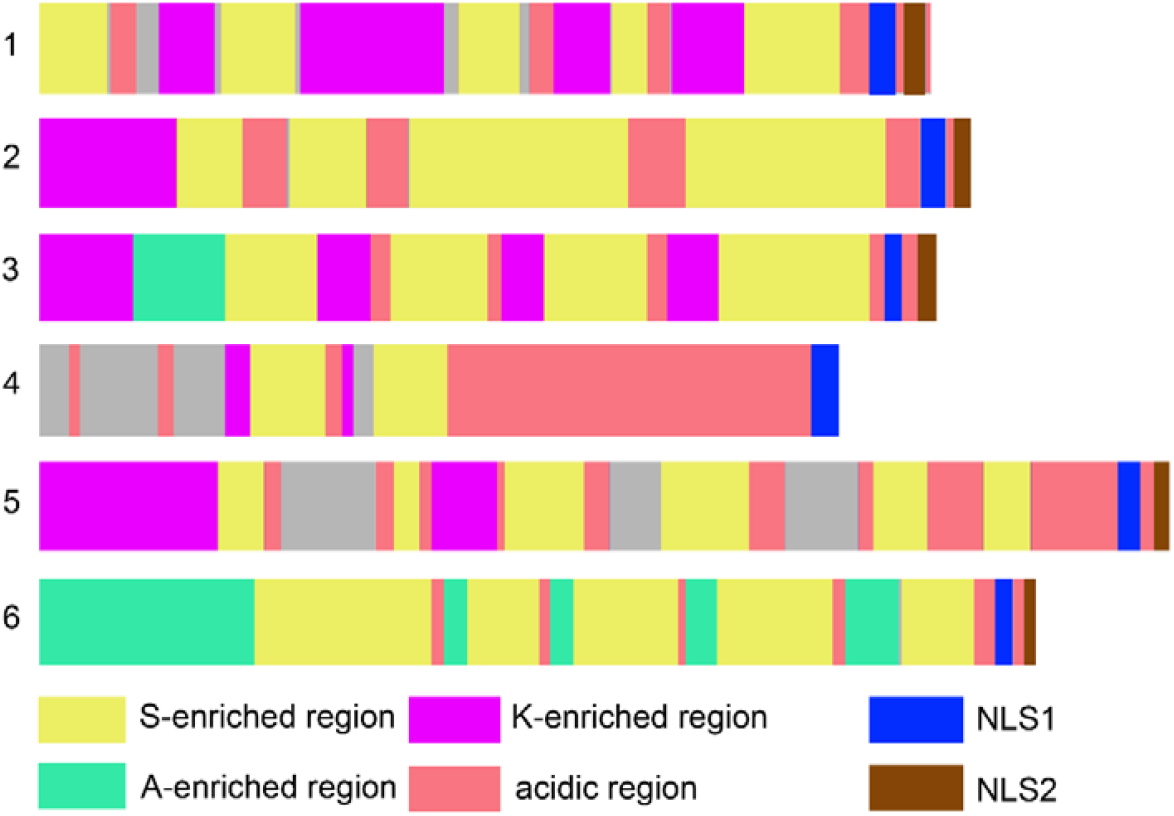
Organization of the N-terminal domains of nucleolins and nucleolin-like proteins of Chromista. The numbers indicate clades according to phylogenetic reconstruction on the Figure 2. The colored boxes correspond to regions in sequences presented in the Supplementary_data 2.

It was shown (González-Camacho & Medina, 2004) that all studied nucleolins excluding those from *Tetrahymena thermophila* had bipartite consensus nuclear localization sequences (NLSs), that are signals for protein import into the nucleus via nuclear pore complexes (Kalderon et al., 1984). We identified this sequence for all clades except the species of Ciliophora and Haptophyta belonging to the clade 4 (Figure 3, Table 1).

**Table 1.** Consensus of nuclear localization sequences belonging to different clades.

| Clades, according to<br>Figure 2 | Consensus NLS1 | Consensus NLS2 |
| --- | --- | --- |
| 1 | PSKKRKA | RKQX |
| 2 | XSKKRKX | KKQX |
| 3 | XSKKRKA | KKXA |
| 4 | XKR/KKKX | - |
| 5 | SKKKKX | KKKX |
| 6 | XXKRKX | KKAX |

The main segments of nucleolins located between the acidic repeats are long and enriched in proline (P) and lysine (K) (Bogre et al., 1996). There are fewer P residues in the N-terminal domain sequences of Chromista than in the sequences of *H. sapiens* NCL (22) and *A. thaliana* nucleolin-like proteins (16-21), while in diatoms their number varies from 8 to 12. K residues are in comparable abundance between Chromista and model organisms; moreover, in some cases they are distributed in small repeats of 3-5 a.a., which are located in homologous positions within phylogenetic groups. However, the position of P and K, as well as acidic boxes (short repeats), were noted only for 1-3 clades (Bacillariophyta) and does not have a general pattern for all Chromista.

A coil-coil region was found in the structure of the N-terminal domain in some predicted sequences (Supplementary table S1), including in the *S. cerevisiae* sequence.

It is known that a region of a polypeptide carrying several consecutive E residues at pH 7.0 does not form an α-helical configuration, since the negatively charged carboxyl groups repel each other (Boyle, 2004). Under these conditions, the α-helical configuration will not be formed in areas with a large number of closely spaced K or arginine (R) residues, which will repel each other due to the positive charges of these residues. Since Chromista sequences are characterized by the presence of both E-, K- and R-rich regions, the N-terminal domain should not have a stable α-helical configuration. In silico analysis provided results consistent with this assumption, since in the sequences of model organisms and various species of Chromista, extended regions with such a conformation are not calculated. Presumably, absence of α-helices in the N-terminal domain contributes not only to the stabilization of the tertiary structure, but also allows the correct folding of the protein chain to perform biological functions. However, in polypeptide chains, a loop or bend appears after the P residue, which is not capable of forming an intrachain hydrogen bond (Szent-Gyorgyi & Cohen, 1957). Thus, we can assume that the saturation of P residues in the N-terminal domain affects the conformation of nucleolin, which is indirectly confirmed by the results of homology modeling of the 3D structure of the protein molecule (Figure 4). According to these 3D-modeling results N-terminal domains of the most studied organism proteins have a more compact conformation than that of a representative of Chromista (*Skeletonema dohrnii*).

**Figure 4.**
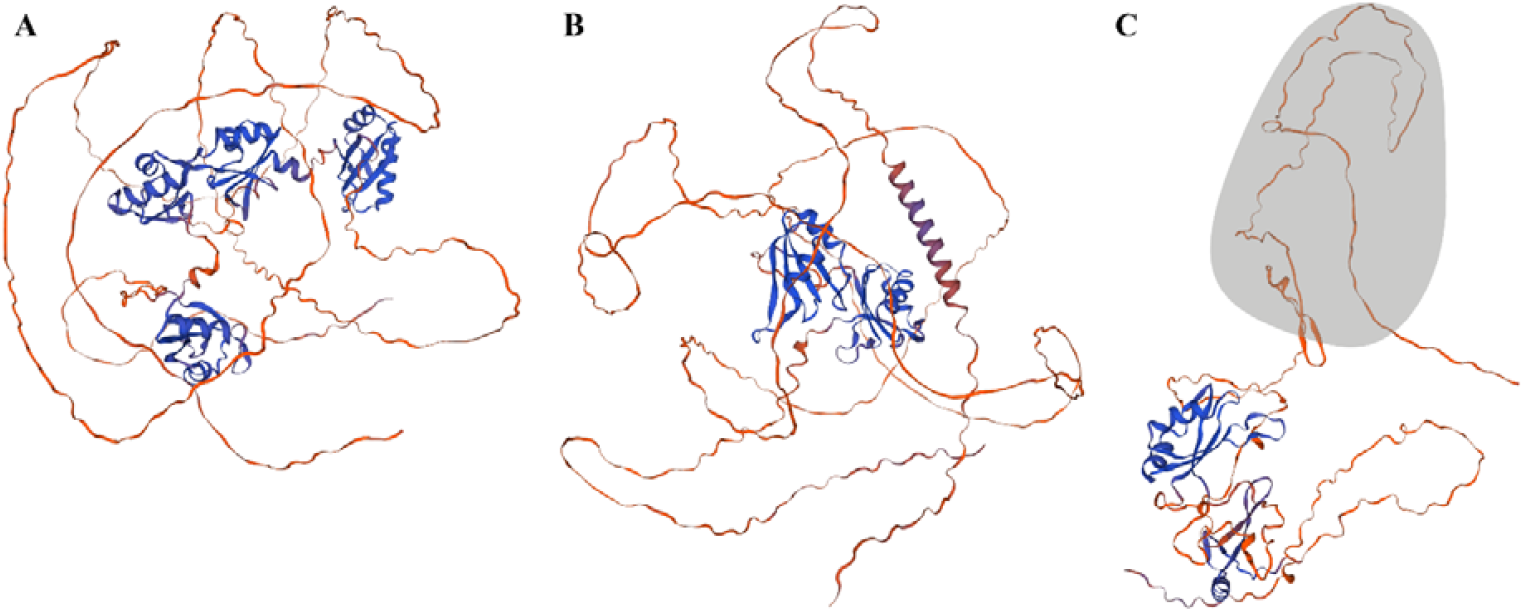
3D-models of nucleolins and nucleolin-like proteins. A – nucleolin of *H. sapiens*, B – nucleolin-like protein of *A. thaliana* (At2NCL), C – nucleolin-like protein of *Skeletonema dohrnii*, the N-terminal domain of *S. dohrnii* is highlighted in gray.

At the beginning of the N-terminal domain of clades 3 and 6, there is an enrichment in alanine (A), which is not present in model organisms (Figure 3). It was previously noted that polyalanine regions are characteristic of DNA and RNA-binding proteins (Veitia, 2004) and are well conserved during evolution (Kumar et al., 2016b). Single diatom species from the classes Coscinodiscophyceae and Mediophyceae, belonging to clades 1 and 2, have A repeats of up to 6 a.a.

Chromista N-terminal domain is also more enriched in serines (S) compared to *A. thaliana* and *H. sapiens*, but similar serine repeats are present in the structure of the *S. cerevisiae* nucleolin-like protein. Serines are located in several repeating sequences of 2-10 a.a. Interestingly, in different species such repeats have different lengths and numbers; the three longest boxes (two of 10 and one of 7 a.a.) are observed in species of Bacillariophyta, included in clades 2, 3 and 6. Post-translational modifications, such as glycosylation (Carpentier et al., 2005), ADP-ribosylation (Paul et al., 2006), and acetylation (Das et al., 2013) play an important role in the localization and functioning of nucleolins. Nucleolins are substrates for a variety of kinases, and it was shown by many studies that phosphorylation has been implicated in its diverse physiological functions (Jordan et al., 1994; Schwab & Dreyer, 1997). Analysis of the post-translational modification sites in Chromista showed the presence of multiple predicted phosphorylation sites, which are also located in serine repeats.

Octapeptide motifs (XTPXKKXX, where X is a nonpolar residue), which are likely responsible for the ability of nucleolin to modulate DNA condensation in chromatin (Erard et al., 1990), are not found in Chromista sequences. However, at homologous sequence positions within each clade there are single motifs with the structure (A/V)P(A/V)KK.

The structure of the central domain contains two RRMs with a length of 71-78 aa. It was previously discussed that, according to RRM sequence identity, nucleolins and nucleolin-like proteins can be divided into three groups within which these motifs are very similar (Ginisty et al., 1999). We noted that the first RRM of Chromista has the maximum similarity to the RRM of *S. cerevisiae*, but the second RRM differs from the RRM of all model organisms and therefore cannot be unambiguously assigned to any of the three identified groups (Ginisty et al., 1999).

For chromist sequences, the presence of conserved a.a. in both RRM domains, which are absent in model objects, is noted. A later study showed the presence of two types of RRM with conserved sequences: RRM1 (R/K)G(F/Y)(G/A)(F/Y)VX(F/Y), and RRM2 (L/I)(F/Y)(V/I)(G/K)(G/N)L (Tajrishi et al., 2011). Analysis of the predicted amino acid sequences showed that these sequences in the Chromista RRM vary among clades and have the structure shown in Table 2, but can be generalized for this group of organisms to the following structure: SGTAXXXF for RRM1 and VFXGNL for RRM2.

**Table 2.**
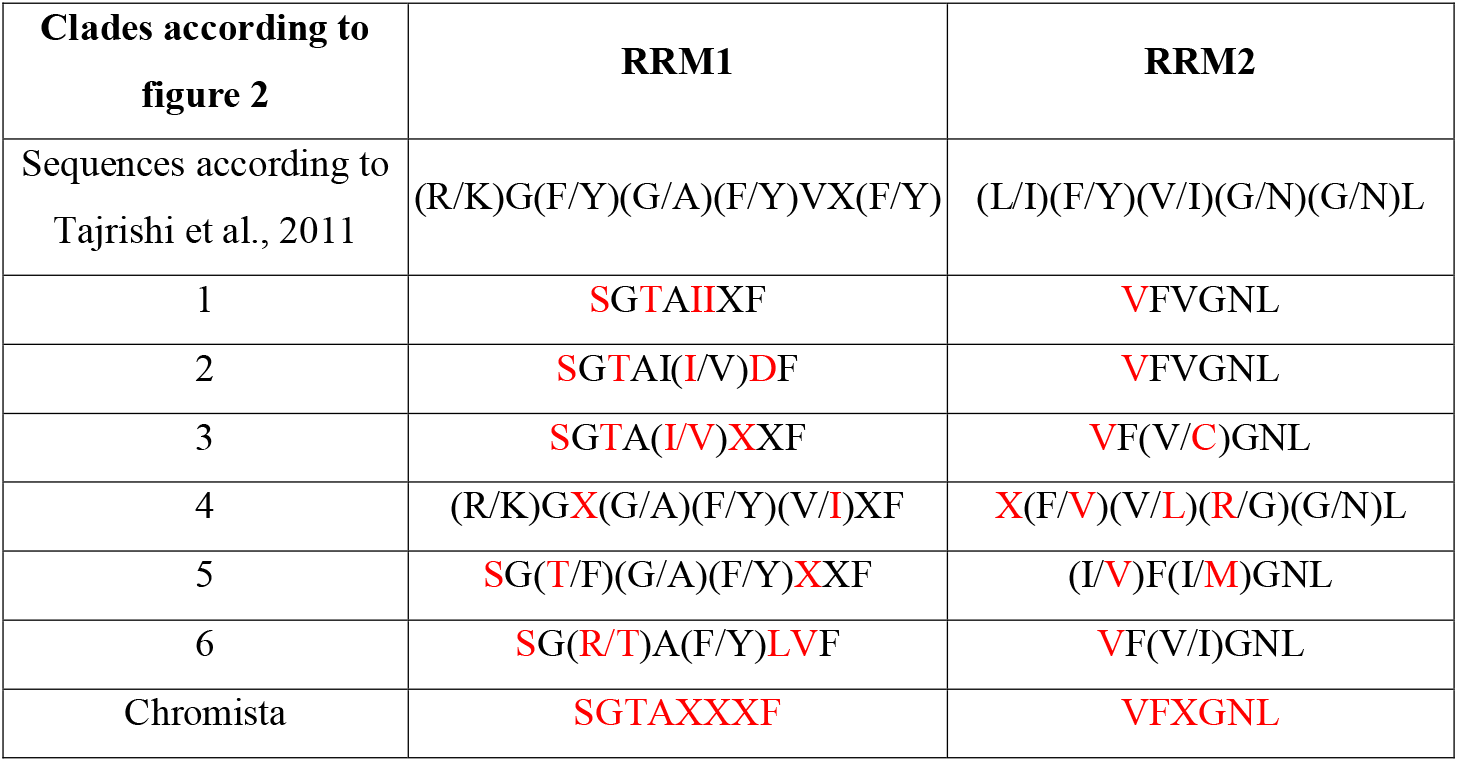
Conserved sequences of RRM1 and RRM2 of Chromista; a.a. specific for Chromista are highlighted by red.

The C-terminal domain of Chromista nucleolin-like proteins has characteristic RGG repeats, but its length varies among both different phyla and among the same phylogenetic clade. We note that the number of RGG repeats and aromatic a.a. in this domain also has no common features.

Thus, the amino acid sequences of various Chromista phyla identified during the analysis were classified by us as nucleolin-like proteins, since their structure clearly defines three domains characteristic of this family of proteins. These proteins have low homology with the model organism proteins, also some structural features of each of the domains. In the structure of the central domain, only two RRM domains are identified, which have their own specific conserved structures. All Chromista are characterized by the presence of S and A boxes and a smaller amount of P in the N-terminal domain, which can significantly affect the conformation of the protein molecule. The proteins under study have characteristics of DNA and RNA-binding proteins and conserved motifs responsible for protein localization in the nucleus. Due to the similar features of not only nucleolins, but also other chromist cytoskeletal proteins (Morozov et al., 2018), their use expands the repertoire of tools for studying the evolution of this kingdom.

## Supporting information

Features of nucleolins and nucleolin-like proteins of the reference group and representatives of the Chromista kingdom

analyzed amino acid nucleolin sequences

alignment of sequences used for phylogenetic reconstruction

## Acknowledgments

This research was funded by Russian Science Foundation, grant number 22-24-00080. We are grateful to A.A. Morozov and E.M. Bayramova for assistance in collecting primary information.

## SUPPORTING INFORMATION

Supplementary materials include:

Supplementary Table S1. Features of nucleolins and nucleolin-like proteins of the reference group and representatives of the Chromista kingdom.

Supplementary data_1 – analyzed amino acid nucleolin sequences.

Supplementary data_2.pdf – alignment of sequences used for phylogenetic reconstruction.

## Notes

### Competing Interest Statement

The authors have declared no competing interest.

