## Supplementary material for "Structural features of the amino acid sequences of Chromista nucleolin-like proteins": analyzed amino acid nucleolin sequences

```
                ....|....| ....|....| ....|....| ....|....| ....|....| ....|....|
                    5         15         25         35         45         55

NCLHs          ---------- ---------- ---------- ---------- ---------- ----------
NCLAt1         ---------- ---------- --MGKS--KS ATKVVAEIKA TKPLKKGKRE PEDDIDTKVS
NCLAt2         ---------- ---------- --MGKSSKKS VTEVETPASM TKPLKKGKRD AEEDLDMQVT
Nsr1pSc        ---------- ---------- ---------- -------MAK TTKVKG---- NKKEV-----
NCLAs          ---------- ---------- ------XKVD KVDKKAAKKA KKEAT----- KAAA------
NCLAsu         ---------- ---------- ---------- -------MGS KVKPD----- KKAAK-----
NCLCd          ---------- ---------- --MG--SKQD KK---EKRAK KKAVA----- EKLA------
NCLCp          ---------- ---MG----- -KSEKGAKSK D-----KLVK KKSSKSKTKV DKKA------
NCLCh          ---------- ---MG----- --KDKKSKGK D------KSA KKASKSKIKA ELKAK-----
NCLCw          ---------- ---------- ---------- ---------- ---------- ----------
NCLCm          ---------- ---------- ---------- -------MSS SSSSS----- SSS-------
NCLCcr         ---------- ---------- ---------- ---------- ---------- ----------
NCLEa          ---------- ---------- --MGSKVRKE KKEKKSKKSP KKEDK----- ERVE------
NCLEs          ---------- ---------- ---------- -------MAK KVLEN----- EKAVE-----
NCLO           ---------- ---MG----- ----SKSKSK D-------SK KDAKRAEKKV MKKAA-----
NCLOs          ---------- ---MGSKSKD KKADKKAKKK DEKSKAKLSK KEDKKAEKEA AKKAKKEAAK
NCLMp          ---------- ---------X RQANMGS-KD KKA--AKKEA KKKAA----- EEAK------
NCLPi          ---------- ---------- ---------- -------MGS DKDKK----- --LKK-----
NCLDf          ---------- ---------- --MG--SKPD KK---SKKSK ESKKA----- AKLAA-----
NCLDc          ---------- ---------- ---------- -------MAS SSSSS----- SSS-------
NCLPj          ---------- ---------- ---------- ---------- ---------- ----------
NCLLda         ---------- ----X----- -RVFSRLFKM G------KAK TGSSKAKKAK SKDSK-----
NCLCa          ---------- ---------- --MG--SKSD KK---AKKAQ KKALA----- EKLA------
NCLCc          ---------- ---------- ---X--FPQD KK---AKKAA KKALA----- EKLL------
NCLChd         ---------- ---------- --MG--SKTD KK---AKKAA KKALV----- EKLA------
NCLCmu         ---------- ---------- ---------- ---------- ---------- ----------
NCLCn          ---------- ---------- --MG--SKTD KK---LKKAA KKALA----- EKLA------
NCLC           ---------L YPKLEDISKS TIMG--SKDD KKA--AKKAA KKALA----- AKLA------
NCLCt1         ---------- ---------- --MGSKSKAD KK---AKKAA KKAEA----- ERLA------
NCLCt2         ---------- ---------- ---------- ---------- ---------- ----------
NCLDbGSO10     ---------- ---MG----- -SKTPKSEKK S-----KKAK KAESKESKLK EKKTD-----
NCLDbGSO10     ---------- ---------- --------XK TKIFMQGLPY TAIKEDVAHF FRNCG-----
NCLSco1        ---------- ---------- ---------- -------MAS SSSSS----- SSS-------
NCLSco2        ---------- ---------- ---------- ---------- ---------- ----------
NCLSd          ---------- ---------- ---------- -------MAS SSSSS----- SSS-------
NCLSj          ---------- ---------- ---------- -------MAS SSSSS----- SSS-D-----
NCLSma         ---------- ---------- ---------- ---------- ---------- ----------
NCLSme         ---------- ---------- ---------- -------MAS SSSSS----- SSSSD-----
NCLTa          ---------- ---------- ---------- -------MAS SSSSS----- SDS-------
NCLTg          ---------- ---------- ---------- ---------- ---------- ----------
NCLTm          ---------- ---------- ---------- -------MAS DSSSS----- SSS-------
NCLTn          ---------- ---------- --MGSKV-KN DKK--AKKQA KKEAA----- EKARK-----
NCLTp          ---------- ---------- ---------- ----VPTKKR KAESV----- SES-------
NCLTo1         ---------- ---------- ---------- -------MSS SSSSS----- SSS-------
NCLTo2         ---------- ---------- ---------- ---------- ---------- ----------
NCLTw          ---------- ---------- ---------- -------MTS SSSSS----- SSS-------
NCLAg          ---------- ---------- --MGSKS-KE DKK--AKKAA KKAAA----- EKAA------
NCLAf          ---------- ---------- ---------- ---------- ---------- ----------
```

```
NCLGo       ---------- ---------- ---------- --------SS SSSSD----- SSS-------
NCLScom     ---------- --------LL FTMGSLN-KE EKK--ARKAA KKAAA----- EKAA------
NCLSr       ---------- ---------- --MGTKTDKS EKK--AKKQA KKELL----- EKAK------
NCLTan      ---------- ---------- --MGTKD-KS DKKKAAKKQA KKEAA----- EKAR------
NCLA1       ---------- ---------- ---------- -----PFVLF DNGNS----- SNTNN-----
NCLA2       ---------- ---------- ---------- ---------- ---------- ----------
NCLAp       ---------- ---------- ---------- ---QPPDPLV NPQKE----- ETVIM-----
NCLAr1      ---------- ---------- ---------- --------XS SSSDD----- DDDDD-----
NCLAr2      ---------- ---------- ---------- --------QT EGLSSTAKAF AKEAS-----
NCLE        ---------- ---------- ---------- ---------X NTKQE----- VKTMV-----
NCLFc1      ---------- ---------- ---------- ---------- ---------- ----------
NCLFc2      ---------- ---------- ---------- ---------- ---------- ----------
NCLFk       MAKSSKKEKK VKDSKPDKKA AKKAKAKEEK AAKKASKKAA ELEAQ----- QKAAE-----
NCLFs1      ---------- ---------M NLVKQVTSTM GADKKEILRQ KTEAA----- EAAAK-----
NCLFs2      ---------- ---------- ---------- ---------- ---------- ----------
NCLFs3      ---------- ---------- ---------- ---------- ---------- ----------
NCLPa       ---------- ---------- ---------- MASSKE---K DLKAK----- QKAAK-----
NCLPau      ---------- ---------- ---------- ---------- ---------- ----------
NCLPd       ---------- ---------- ---------- MAGNKE---K ELKAK----- LKAAK-----
NCLPf       ---------- ---------- ---------M AKASKD---K DLKAK----- QKAAK-----
NCLPh       ---------- ---------- ---------M AKANKE---K VLKAK----- QKAAK-----
NCLPp       --XSKRLQKK PKESEQQRDK PDHKGIEQIK IMANKE---K ELKAK----- QKAAK-----
NCLNpa      ---------- ---------- ---------- ---------- ---------- ----------
NCLNp       ---------- ---------- ---------M AKISKEKLAK ELKAK----- QKAVE-----
NCLNi1      ---------- ---------- ---------- ---------- ---------- ----------
NCLNi2      ---------- ---------- ---------- ---------- ---------- ----------
NCLPsp      ---------- ---------- ---------- -------MGK TKTDKKAKLA QELAE-----
NCLTv       ---------- ---------- ---------- -------MGK SATKD----- --AKK-----
NCLTs       ---------- ---------- ---------- -------MGK SATKD----- --AKK-----
NCLTc       ---------- ---------- ---------- -----MASAT VDKKA----- --AKK-----
NCLNsa      ---------- --------MG KSSKSKADKT LGADVVVQSK GGSAEGTKAV GKKGKKAA--
NCLNg       ---------- --------MG KSSKSKADKT LGADVVVQSK GGSAEGTKAV GKKGKKAA--
NCLTmi      ---------- ---------- ---------- -----MVRSP PPSKAVAAAN GKKAQ-----
NCLCr       ---------- ---------- --MASTDVEL HQLVHQFLKA EGLSKSAKAF AKEAA-----
NCLBs       ---------- ---------- ---------- -------MG- ---------- ----------
NCLMc       ---------- ---------- ---------- -------MPK TKKSKKEAKL SKKEL-----
NCLSl       ---------- ---------- ---------- ---------- ---------- ----------
NCLCsu      ---------- ---------- MKQKSAPVAV SSSASSSDSD SSEEEPQAKV QKATLKRVAE
NCLEv       ---------- ---------- ---------- ---------- ---------- ----------
NCLPg       ---------- ---------- ---------- ---------- ---------- ----------
NCLEt       ---------- MHGGLNAKEN VEDLKAKVAQ -VVANDDS-L SSSEE---EV QPKKA--IAK
NCLEn       ---------- MHGGLNAEEN VEDLKAKVAQ -PVANDDS-L SSSEE---EV QPKK---VAK
NCLEm       ---------- ---------- --MAKAKVAQ -PVASDDSSL SSSEE---EV QPKKA-VSAK
NCLEb       ---------- ---------- --MAKAKVAQ -PVASDDSSL SSSEE---EV QPKKT-VASK
NCLEmi      ---------- ---------- --MAKAKVAQ -PVASDDSSL SSSEE---EV QPKKA-VPAK
NCLEa       ---------- ---------- --MAKAKVVQ QPVASDDSSL SSSEE---EV QPKKAAIAAK
NCLEp       ---------- ---------- --MAKAKVAQ -PVASDDSSL SSSEE---EV QPKKP-VAAK
NCLPo1      ---------- ---------- ---------- ---------- ---------- ----------
NCLPo2      ---------- ---------- ---------- ---------- ---------- ----------
NCLPm       ---------- ---------- ---------- ---------- ---------- ----------
NCLCc1      ---------- ---------- ---------- ---------- ---------- ----------
NCLCc2      ---------- ---------- ---------- ---------- ---------- ----------
NCLSmi      ---------- ---------- ---------- ---------- ---------- ----------
NCLPbr      ---------- ---------- ---------- -------MGS TKAATVVKKP TRSKD-----
```

```
NCLGt         ---------- ---------- ---------- ---------- ---------- ----------
NCLG          ---------- ---------- --MTKASAGI KRKAGDEPTK KKASKESESD SSSDSESDEE
NCLDlu        ---------- ------MAAT ADDHTILSLV HEFLTSKGLT RTANALLSEV KTPVQPLKPG
NCLP          ---------- ---------- ---------- ---------- ---------- ----------

              ....|....| ....|....| ....|....| ....|....| ....|....| ....|....|
                    65         75         85         95        105        115

NCLHs         ---------- ---------- ---------- ---------- ---------- ----------
NCLAt1        LKKQKKDVIA AVQKEK---- ---------A VKKVPKKVES SDDSDSESEE EEKAKKVPAK
NCLAt2        -KKQKKELID VVQKEK---- ---------A EKTVPKKVES SSSDASDSDE EEKTKETPSK
Nsr1pSc       ------KASK QAKEEK---- ---------- -AKAVSSSSS ---------- ---------E
NCLAs         ---------K EAAA-AAA-- ---------- KAAAEAAAAA AKAAAEAAAD SDSDDSSSS-
NCLAsu        ------KAEA AR-------- ---------- --LKADAEEL ARKAEELKKK ATKKDETSSD
NCLCd         ---------K EAKELA---- ---------- ----AAAAAA AAAAAANDSD SD---SDSDS
NCLCp         -------AKA AAEAAAAA-- ---------- ---AKAAAEE AARLAAAA-- --------AA
NCLCh         ------ADRE AAEALAAA-- ---------- ----KAAAEE AERLAKAAEE AAKKEKPSSD
NCLCw         ---------- ---------- ---------- ---------- ---------- ----------
NCLCm         ---------D SSSEDE---- ---------- -KKVVH-KPI ATKKKDSSSS SSS-----SS
NCLCcr        ---------- ---------- ---------- ---------- ---------- ----------
NCLEa         ---------K LAAEKAAA-- ---------- EKAAAEKAAA EKAAAEKAAE SDSDSSSSSD
NCLEs         ------EESS SSDSDSSS-- ---------- SSDSSDSEDE APPKKVEDKV VAKKKEKESN
NCLO          ----AEAAKK AAEAAAEA-- ---------- ---AKKAAEE DARKAAEDSS DSGSDSDSSS
NCLOs         KAKKEAEAKA KAEAEAKAKK EAEEKAKKEA EEKAKKEAEE KARKEAEAKK KKEEEESSSS
NCLMp         ---------K AAEEAA---- ---------- -RKAKELAEA AAKAADDSDD DDSSVSSSSS
NCLPi         ------AAKK EKAAKA---- ---------- KAEAEAAAKK AADLAAAAAK AAA-----AA
NCLDf         --------KK EAEEAARK-- ---------- AKEAAEAAKR AAEEAAAKAK EE---ASSDS
NCLDc         ------SDSS S-EEDK---- ---------- -KKVIE-KKP IATKKS---- ----------
NCLPj         ---------- ---------- ---------- ---------- ---------- ----------
NCLLda        -------KYE LAKAKAEE-- ---------- ---LKRLAEE AKRLAEEAAK AELEASKMDD
NCLCa         ---------R EAEEAA---- ---------- -KKAAEAAKK AAEEAAAAAE AE---GSDSD
NCLCc         ---------K EAEAAK---- ---------- -KAAEEAAK- ---AAAAAAD AE---DSGSE
NCLChd        ---------K EAAAAA---- ---------- -AAAAEAAKK AAEAS-DSDS SD---SDSSD
NCLCmu        ---------- ---------- ---------- ---------- ---------- ----------
NCLCn         ---------K EAEEAA---- ---------- -KAAAEAAKK AAIEAAAESD SD---SSSSS
NCLC          ---------K EAEEAA---- ---------- -KAAAEAAKK AAEEALEDES ST---SSSSS
NCLCt1        ---------K EAEEAA---- ---------- -KAAAEAAKK AAEEAADDDS SS---SSSSS
NCLCt2        ---------- ---------- ---------- ---------- ---------- ----------
NCLDbGSO10    -----KEAKK AAKKLKKE-- ---------- EEKAKKAAAE AAAKKAAEEA AAKKAAEEAA
NCLDbGSO10    ---------- ---------- ---------- ---TIKHIDL PLDYDGRSSG SALVSFETTD
NCLSco1       ------DSSS SSSEDE---- ---------- -KKVVD-TPK IAKKKGSSSS SSS-----SS
NCLSco2       ---------- ---------- ---------- ---------- ---------- ----------
NCLSd         ------DSSS SSSEDE---- ---------- -KKVVD-TPK IAKKKGSSSS SSS-----SS
NCLSj         ------SSSS SSSEEE---- ---------- -KKVVD-TPK IAHKKGSSSS SSS-----SS
NCLSma        ---------- ---------- ---------- ---------- ---------- ----------
NCLSme        ------SSSS SSSEDE---- ---------- -KKVVD-TPK IAHKKGSSSS SSS-----SS
NCLTa         --------SS SEEETK---- ---------- -KVVTTPVAK KTKAKASSSS KKSKKEVVAS
NCLTg         ---------- ---------- ---------- ---------- ---------- ----------
NCLTm         ------SDSS SSEEET---- ---------- -KKVVETKKP IAKKKGSSDN SSS-----SG
NCLTn         ------AAEQ AAEEAA---- ---------- -RKAAELKKI AEAEE----- -----SSDSD
NCLTp         ------NGAD APAEKL---- ---------- -QKTGEEGDE NTKIYVRGLP WRATEDEVRE
NCLTo1        ------SSSS SSSEDE---- ---------- -KPKVEEKKV IARKKSSSSS SSSS-SSSSE
NCLTo2        ---------- ---------- ---------- ---------- ---------- ----------
NCLTw         ------SSES SSSEDE---- ---------- -KKIVNEKPI AVKKGSSSSA SSSSGSDSSD
```

```
NCLAg       ---------K EAEEAA---- ---------- -RKAAELAKK AAEAE----- -----SSDSS
NCLAf       ---------- ---------- ---------- ---------- ---------- ----------
NCLGo       -------SSE DEKPTT---- ---------- -KKTMTVTKK KTKKEESSDS SSS-----SD
NCLScom     ---------K EAAEAL---- ---------- -ERAKKLEAA AAAAEA---- -----SSSSE
NCLSr       ---------K AAEEAA---- ---------- -AKAAELAKA AESSD----- --------SS
NCLTan      ---------K EAEEAA---- ---------- -KKAAELAKK AAAEEKSS-- -----SDDSD
NCLA1       ------TTMV KLATLQ---- ---------- -KHVEEAQKA AAEAQANADK AAKLAQECLA
NCLA2       ---------- ---------- ---------- ---------- ---------- ----------
NCLAp       ------GSSS EYETLQ---- ---------- -KRAAAAQKE AEVAQQAAEK AAQVAQQALA
NCLAr1      ------D--- --DESS---- ---------- -DNSDDSDDE KVKPVVKTDP SKATAKSKSK
NCLAr2      ------KAGI KNVTEV---- ---------- -DSIDTPLKQ VYSAWADSRP AKR--ERASS
NCLE        ------D--- --AALK---- ---------- -KQAKAAKKA AAEAHKAAER AATLAKEALE
NCLFc1      ---------- ---------- ---------- ---------- ---------- ----------
NCLFc2      ---------- ---------- ---------- ---------- ---------- ----------
NCLFk       ------EAKK AADDAI---- ---------- -KAAEKLAEE LLKKE-DSGS DSD-------
NCLFs1      ------EAQE ATARAE---- ---------- -RLAKEVKEL AEQLKSSGKK TKATKSVVSD
NCLFs2      ---------- ---------- ---------- ---------- ---------- ----------
NCLFs3      ---------- ---------- ---------- ---------- ---------- ----------
NCLPa       ------EAKK AADDAI---- ---------- -KKAEKLAEE VAKMESEIAA AAK------K
NCLPau      ---------- ---------- ---------- ---------- ---------- ----------
NCLPd       ------EAKK AADEAV---- ---------- -KKAEKLAEE VAKLESDINA EAK------K
NCLPf       ------EAKK AADDAI---- ---------- -KKAEKLAEE VAKLESEMEA EAK------K
NCLPh       ------EAKK AADDAI---- ---------- -KKAEKLAEE VAKLESEIKK EAK------K
NCLPp       ------EAKK AADEAI---- ---------- -KKAEELAEE VAKLESEVEA AAK------K
NCLNpa      ---------- ---------- ---------- ---------- ---------- ----------
NCLNp       ------EAKQ IAQEAL---- ---------- -KKAERLAEE VAKMESEIEG DGKPSKPEAK
NCLNi1      ---------- ---------- ---------- ---------- ---------- ----------
NCLNi2      ---------- ---------- ---------- ---------- ---------- ----------
NCLPsp      ------QKKK MADLEA---- ---------- KLAAEAVVES SSDSDSDSDS SSS-----ED
NCLTv       ------IAKK AKAAAK---- ---------- QKELELAMKK AQEEIKRLQE AND-----SS
NCLTs       ------IAKK AKAAAK---- ---------- QKELELAMKK AQEEIKRLQE AND-----SS
NCLTc       ------LAKK QKAEAL---- ---------- KLKAKEAAEA AAKAAKLAAE EAAKAAKMAE
NCLNsa      ----KPVEPS SSSSSD---- ---------- -SSSDEEVTP TTNGTAGKKG GKVVAKPVAA
NCLNg       ----KPVEPS SSSSSD---- ---------- -SSSDEETAP TTNGTAGKKG GKVVAKPVAA
NCLTmi      ---------E SSEESE---- ---------- -SDSDE---- ---------- ----------
NCLCr       ------KVG- -AVTEE---- ---------- -PGQSVTLSE VFKAFSAQRA SKRPREEPKA
NCLBs       ---------K YAK------- ---------- ---------- ---------- ----------
NCLMc       ------KAKA KAQESE---- ---------- -ESSDSDSDS SVEEPVKTKK TKKAPVVEKK
NCLSl       ---------- ---------- ---------- ---------- ---------- ----------
NCLCsu      GG-AKQKAQR KNRGKAPPTD SSEDTS---S DDESTGTVRK SLVESSAAAV QRAKQAVRAT
NCLEv       ---------- ---------- ---------- ---------- ---------- ----------
NCLPg       ---------- ---------- ---------- ---------- ---------- ----------
NCLEt       P---VSTASK KKTAPVPPPS SSDDSS--DE DQPSATVVKK AAAAKAKTSG KGVAAKAQSR
NCLEn       P---VSTASK KKTAPVPPPS SSDDSS--DE DQPSATVVKK AAAAKAKAAG KGIAAKAQSQ
NCLEm       P--VAAALKK RAAAAAAPSS SSDDSS--DE DQPSAAVVKK AAAAKANNAA KGITAKAQST
NCLEb       A--SAASIKK KAVAPPS--- SSDDSS--DE DQPSATVVKK AAAAKNNNAA KGLAAKAQST
NCLEmi      A--AGAALKK KAPAPAAPSS SSDDSS--DE DQPSATVVKK AAAAKKNNVG KGPAAKAQST
NCLEa       AGGAAAAVKK KAAAAVAPAS SSSDDSSDDD DQPSAAVVKK AVAAKTN-AT KGITAKAQST
NCLEp       A--GAAALKK KA-APPPPSS SSDDSS--DE DQPSAVVVKK AAAAKANNAG KAVAAKAQST
NCLPo1      --------MA KPAPKKVVES SSDESS---S DEEVQATTKP IVHKKHAAAA KKKAEKKPVV
NCLPo2      --------MA KPAPKKVVES SSDESS---S DEEVQATTKP IVHKKHAAAA KKKAEKKPVV
NCLPm       --------MA KPAPKKVVES SSDESS---S DEEVQAP-KP VVHKKKVAAT KKKAEKEPVV
NCLCc1      ---------- ---------- ---------- ---MHKQKSQ KIEGGISTAT LGNISKKQHK
NCLCc2      ---------- ---------- ---------- -----KQKSQ KIEGGVSTAI QGNIAKKQRK
```

```
NCLSmi         ---------- ---------- ---------- ---------- ---------- ----------
NCLPbr         ---------K YNKNKD---- ---------- -KNKGAVKDA PVKNAAKDVE AKKAAAAAKK
NCLGt          ---------- ---------- ---------- ---------- ---------- ----------
NCLG           EVKAKKPVKA AAKAAPAK-- --------AD KKKAAKKEES SSEEESSDDE DAKKPAAKKA
NCLDlu         TPPLAQLVRL RASAPAAPPA ETSEESSDDS DESSKAAAKE SSDDSDDSDE SSKAAAEESS
NCLP           ---------- ---------- ---------- ---------- ---------- ----------

               ....|....| ....|....| ....|....| ....|....| ....|....| ....|....|
                   125        135        145        155        165        175

NCLHs          ---------- ---------- ---------- ---------- ---------- ----------
NCLAt1         KAASSSDE-- ---------- ---------- -------SSD DSSSDD---- ----------
NCLAt2         LKDESSSE-- ---------- ---------- -------EED DSSSDE---- ----------
Nsr1pSc        SSSSSS---- ---------- ---------- -------SSS ESESE----- ----------
NCLAs          SSSSSSS--- ---------- ---------- -------SSS SSEDEAPKKV VV--------
NCLAsu         EDSDASS--- ---------- ---------- -------SSD SESE------ ----------
NCLCd          SSSSSSS--- ---------- ---------- -------SSS SEAPKE---- ----------
NCLCp          SSSSSSD--- ---------- ---------- -------SSD SSSSDD---- ----------
NCLCh          SSSSSDS--- ---------- ---------- -------SSD SSSDSD---- ----------
NCLCw          ---SSSS--- ---------- ---------- -------SSD SSSD------ ----------
NCLCm          SDSSDSD--- ---------- ---------- -------AKA KTK------- ----------
NCLCcr         ---------- ---------- ---------- ---------- ---------- ----------
NCLEa          SSSSSSS--- ---------- ---------- -------SSS SSEDEKP--- ----------
NCLEs          SSSESSS--- ---------- ---------- -------SSD SSDDDS---- ----------
NCLO           SSSSSSSXDE APAPPKKKEG KKDKAKAKEA AKKKDSTSSS SSSSSS---- ----------
NCLOs          SSSSSSD--- ---------- ---------- -------SDS SSDSDS---- ----------
NCLMp          SSSSSSS--- ---------- ---------- -------DSD SEDEKP---- ----------
NCLPi          AAKTADS--- ---------- ---------- -------SDS SDSDSD---- ----------
NCLDf          SDSDSDS--- ---------- ---------- -------SDD SSSSED---- ----------
NCLDc          ---------- ---------- ---------- ---------- ---------- ----------
NCLPj          ---------- ---------- ---------- ---------- ---------- ----------
NCLLda         ESTSDSS--- ---------- ---------- -------SSS DSSDSE---- ----------
NCLCa          SSSSSSS--- ---------- ---------- -------SSD SSDSEN---- ----------
NCLCc          SDSSSSS--- ---------- ---------- -------SSS SSDSES---- ----------
NCLChd         SDSDSSS--- ---------- ---------- -------CSD SSVEAP---- ----------
NCLCmu         ---------- ---------- ---------- ---------- ---------- ----------
NCLCn          SSDSSDS--- ---------- ---------- -------EAE APKSKK---- ----------
NCLC           SSSSSSS--- ---------- ---------- -------SSS SSSESEKPKE ----------
NCLCt1         SSSSSSS--- ---------- ---------- -------SSS SDSDAE---- ----------
NCLCt2         ---------- ---------- ---------- ---------- ---------- ----------
NCLDbGSO10     AKKAAEE--- ---------- ---------- -------AAK KAAEEA---- ----------
NCLDbGSO10     AAAAAVALHG ---------- ---------- -------EMF DEENGR---- ----------
NCLSco1        S--DSSD--- ---------- ---------- -------SEA EKKA------ ----------
NCLSco2        ---------- ---------- ---------- ---------- ---------- ----------
NCLSd          SSSDSSD--- ---------- ---------- -------SEA EKKA------ ----------
NCLSj          ---DSSD--- ---------- ---------- -------SEA QKK------- ----------
NCLSma         ---------- ---------- ---------- ---------- ---------- ----------
NCLSme         SSSDSSD--- ---------- ---------- -------SEA EKK------- ----------
NCLTa          SSSDSSD--- ---------- ---------- -------SES DSKK------ ----------
NCLTg          ---------- ---------- ---------- ---------- ---------- ----------
NCLTm          SDSSDSD--- ---------- ---------- -------SDA KAKAKAKASS KKG-------
NCLTn          SSDSDSS--- ---------- ---------- -------DEE TPAPKA---- ----------
NCLTp          FFAQCGE--- ---------- ---------- -------MES VELPL----- ----------
NCLTo1         SEAEAEA--- ---------- ---------- -------KPA PPPKK----- ----------
```

```
NCLTo2      ---------- ---------- ---------- ---------- ---------- ----------
NCLTw       SEAKKKK--- ---------- ---------- -------KKT KAVTK----- ----------
NCLAg       SSDSDSS--- ---------- ---------- -------DSE SEDEKP---- ----------
NCLAf       ---------- ---------- ---------- ---------- ---------- ----------
NCLGo       SDSSSDE--- ---------- ---------- -------EEV KAK------- ----------
NCLScom     SESDSSS--- ---------- ---------- -------ESE SEDEAP---- ----------
NCLSr       SSDSDSS--- ---------- ---------- -------SDE -EEKAA---- ----------
NCLTan      SSDDDSS--- ---------- ---------- -------SDD GSSSSS---- ----------
NCLA1       ELQKHQA--- ---------- ---------- -------EAT N--------- ----------
NCLA2       ---------- ---------- ---------- ---------- ---------- ----------
NCLAp       TAQKLQK--- ---------- ---------- -------KAD QAQKP----- ----------
NCLAr1      KAQDSED--- ---------- ---------- -------DDD S--------- ----------
NCLAr2      TSSSDSDS-- ---------- ---------- -------SSD SSSSES---- ----------
NCLE        LSKELEE--- ---------- ---------- -------AEE K--------- ----------
NCLFc1      ---------- ---------- ---------- ---------- ---------- ----------
NCLFc2      ---------- ---------- ---------- ---------- ---------- ----------
NCLFk       SSSSSSS--- ---------- ---------- -------SSS SSSS------ ----------
NCLFs1      DEQKKAE--- ---------- ---------- -------ESD SDSS------ ----------
NCLFs2      ---------- ---------- ---------- ---------- ---------- ----------
NCLFs3      ---------- ---------- ---------- ---------- ---------- ----------
NCLPa       AAESS----- ---------- ---------- -------DSD SDSS------ ----------
NCLPau      ---------- ---------- ---------- ---------- ---------- ----------
NCLPd       AEESS----- ---------- ---------- -------SSD SDSS------ ----------
NCLPf       AAKKAES--- ---------- ---------- -------SSD SDSS------ ----------
NCLPh       SDSSSS---- ---------- ---------- -------DSD SDSS------ ----------
NCLPp       AKDDSG---- ---------- ---------- -------SDS DSSS------ ----------
NCLNpa      ---------- ---------- ---------- ---------- ---------- ----------
NCLNp       AVENDD---- ---------- ---------- -------SSD SDSS------ ----------
NCLNi1      ---------- ---------- ---------- ---------- ---------- ----------
NCLNi2      ---------- ---------- ---------- ---------- ---------- ----------
NCLPsp      EKKEVTK--- ---------- ---------- -------KVT DESDSD---- ----------
NCLTv       SSSDSDS--- ---------- ---------- -------DSD SDSDDE---- ----------
NCLTs       SSSDSDS--- ---------- ---------- -------DSD SDSDDE---- ----------
NCLTc       SDSDSDS--- ---------- ---------- -------DSD SDSDSD---- ----------
NCLNsa      SSDSDDD--- ---------- ---------- -----SSSSS -EDEAP---- ----------
NCLNg       SSDSDDD--- ---------- ---------- -----SSSSS SEDEAP---- ----------
NCLTmi      ---------- ---------- ---------- -------SSS EEDERP---- ----------
NCLCr       ETSSSSSS-- ---------- ---------- -------SSS SSSSDS---- ----------
NCLBs       ---------- ---------- ---------- ---------- ---------- ----------
NCLMc       NDSSDSDEAP AAGKKRKPET VKATTKKPKV EE---NLSSD DSDSDSVDVI AKP-------
NCLSl       ---------- ---------- ---------- ---------- ---------- ----------
NCLCsu      ALSKQ----- ---------- ------KAPA KPSAGSDGSS SDSTEDETPQ TK---HASKA
NCLEv       ---------- ---------- ---------- ---------- ---------- ----------
NCLPg       ---------- ---------- ---------- ---------- ---------- ----------
NCLEt       AQDSSESSSS EDDAPAAKKG KGAVTKPALP QKAVTHSSDS EDSSENEPPA KK---PAQKV
NCLEn       AQDSSESSSS EDDTPAAKKG KGAVTKPALP QKAVAQSSDS EDSSENEPPA KK---PAQKV
NCLEm       AQDSS-DSSS DDERPAPKKG Q-FAAKAAAS QRAAAQDSDS DDSSEEEPPA KK---PAQKA
NCLEb       AQDSS-DSSS EDEAPAPKKG H-TAGKAAAL QKAAAQDSDS DDSSENEPPA KK---PVQKA
NCLEmi      AQDSS-DSSS DDEMPAPKKG Q-AAAKAAAL QKAAAQDSDS DDSSEDEPPA TK---PAQKA
NCLEa       IQDSSSDSSS EDEMPPPKKG Q-VPAKAAP- QKAAVQDSDS DDSSEDEAPA KK---PVQKA
NCLEp       AQDSS-DSSS EDEMPAPKKG Q-VPAKAAAL QKAAAQDSDS DDSSEEEPQP KK---PVQKA
NCLPo1      EDSSSSDSDS SVDEAPAPKA KKAVAKKAPA PKKAAKKAAE SSSSSDESSS DEEEVPAKKA
NCLPo2      EDSSSSDSDS SVDEAPAPKA KKAVAKKAPA PKKAAKKAAE SSSSSDESSS DEEEVPAKKA
NCLPm       EDSSS-DSDS SVDEAPAPKA KKAVAKKAAP ---AAKKAAP AE-------- ----------
```

```
NCLCc1        VKSSN----- ---------- --FKIQPFVP ATTKNQEPDL TDTISDKDTS AV---AVQKA
NCLCc2        GESSK----- ---------- --FKLQPFVA ATKEKLQADL SDTSSDEDTS TV---AVQKA
NCLSmi        ---------- ---------- ---------- ---------- ---------- ----------
NCLPbr        VAAAAPA--- ---------- ---------- -------QKP ANDAVP---- ----------
NCLGt         ---------- ---------- ---------- ---------- ---------- ----------
NCLG          DAKKAAP--- ---------- ---------- ----AKKAAK KEESSS---- ----------
NCLDlu        DDSDESS--- ---------- ---------- -----KAAAE ESSDDSDEGS KAA-------
NCLP          ---------- ---------- ---------- ---------- ---------- ----------

              ....|....| ....|....| ....|....| ....|....| ....|....| ....|....|
                 185        195        205        215        225        235

NCLHs         ---------- ---------- ---------- ---------- ------MVKL AKAGKNQGDP
NCLAt1        ---------- -----EPAP- -------KKA VAATNG---- ---------- ----TVAKKS
NCLAt2        ---------- -----EIAPA KKRPEPIKKA KVESSSSDDD STSDEETAPV KKQPAVLEKA
Nsr1pSc       ---------- ------SESE SESSSSS--- ---------- ---------- ----SSSDSE
NCLAs         ---------- -----KKDKK KAVKLSKKEE KKAA------ ---------- --KVKEEKKA
NCLAsu        ---------- ---------A EVKVETKTKG ---------- ---------- ------KKND
NCLCd         ---------- -------TAK IATKKTKA-- ---------- ---------- ----PVKKSD
NCLCp         ---------- ------SDEE -------MKP AAP------- ---------- ---PAAKKKA
NCLCh         ---------- ------SDSS ----VEAKKP AAK------- ---------- ---AAPAKKP
NCLCw         ---------- ---------E EEKAKPAP-- ---------- ---------- ----------
NCLCm         ---------- ---KAVAKKK EPKKEVKKEE ---------- ---------- ----------
NCLCcr        ---------- ---------- ---------- ---------- ---------- ----------
NCLEa         ---------- -------EKK KSEKVVKKIE T--------- ---------- ----KEVKKS
NCLEs         ---------- -----SDDEE DEKAEAKPAE ---------- ---------- ------KPVA
NCLO          ---------- ------SSSE EEKKKADKSK AKASKFK--- ---------- ---AEAKKKE
NCLOs         ---------- ------SDSE DEKAPAKKKD AAA------- ---------- ---AAAKKSS
NCLMp         ---------- -------AKK EDKAKAVPA- ---------- ---------- ----KKDDSS
NCLPi         ---------- -------SDS DSDSSSSSSS SED------- ---------- ----KKKKTK
NCLDf         ---------- -------ESI VTKNKTKK-- ---------- ---------- ----SEEKTN
NCLDc         ---------- --------KK ESKKEASKKK ---------- ---------- -----AAKKA
NCLPj         ---------- ---------- ---------- ---------- ---------- ----------
NCLLda        ---------- ------SEPE ----PETKKP VEK------- ---------- ---KEESESS
NCLCa         ---------- -------EAA APEKKASD-- ---------- ---------- ----AKKKDD
NCLCc         ---------- -------EAE APPAKKEA-- ---------- ---------- ----PKAASS
NCLChd        ---------- -------AKK VAVKKTK--- ---------- ---------- ----KAKKVE
NCLCmu        ---------- ---------- ---------- ---------- ---------- ----------
NCLCn         ---------- -------ETK VVKEKTKV-- ---------- ---------- ----VKK-DD
NCLC          ---------- -----KGTKE EASKQSKDST SDSSDDDDD- ---------- --EGKEKQKE
NCLCt1        ---------- -------MKD ADKKEEK--- ---------- ---------- ----KEEKAE
NCLCt2        ---------- ---------- ---------- ---------- ---------- ----------
NCLDbGSO10    ---------- ------AKKA AEEAAAKKDS SDS------- ---------- ---SSSDSSS
NCLDbGSO10    ---------- -----WVKIK YDTPRVPPNK EYQ------- ---------- ----KQQQHQ
NCLSco1       ---------- --AVVVAKKK ESKKEAKKKA ---------- ---------- ----EAKKKA
NCLSco2       ---------- ---------- ---------- ---------- ---------- ----------
NCLSd         ---------- --AVVVAKKK ESKKEAKKKA ---------- ---------- ----EAKKKA
NCLSj         ---------- ---VVAKKKK ESKKEAKKKA ---------- ---------- ----EAEAKK
NCLSma        ---------- ------AKKK ESKKEAKKKA ---------- ---------- ----EAKKKA
NCLSme        ---------- ---VVSKKK- ESKKEAKKKA ---------- ---------- ----EAEAKK
NCLTa         ---------- --KAKAKKAK KSKKESKKEV AVK------- ---------- ----KKKSKK
NCLTg         ---------- ---------- ---------- ---------- ---------- ----------
NCLTm         -EKGKKGKKK EEAPVAKKKK DSKKEAAVKK KKK------- ---------- ----EVKKKE
NCLTn         ---------- -------SNK KKKAGAEVK- ---------- ---------- ----KEESDS
```

```
NCLTp       ---------Q DDGRSSGTAI IDFKEASAAA AAL------- ---------- ----EQNGAD
NCLTo1      ---------A ES-KKEKKVK AKKEAEKKEE SSS------- ---------- ----------
NCLTo2      ---------- ---------- ---------- ---------- ---------- ----------
NCLTw       ---------K ETPKKETPKK VEKKADKKKD TPA------- ---------- ----KKTKKK
NCLAg       ---------- -------TKT KTKATK---- ---------- ---------- ----KEESSD
NCLAf       ---------- ---------- ---------- ---------- ---------- ----------
NCLGo       ---------- ---VVVSKKK KEKKVAVATK KVE------- ---------- ----SKKKSK
NCLScom     ---------- -------TKK VETKKE---- ---------- ---------- ----KGSSSS
NCLSr       ---------- -------PVK KGKDSKKKK- ---------- ---------- ----KEESSS
NCLTan      ---------- -------EEK PKKKAPPAK- ---------- ---------- ----KKESQK
NCLA1       ---------- -----VSVSS EEDKEAKKAN KSK------- ---------- ----DSDDSS
NCLA2       ---------- ---------- ---------- ---------- ---------- ----------
NCLAp       ---------- -PTETVSLSS DDGAQKKKKK QDD------- ---------- ----SSDDSS
NCLAr1      ---------- -----SDDDS SDDDEKAKAT PSK------- ---------- ----PQNIKA
NCLAr2      ---------- ----SAAPPA AKRTKAAD-- ---------- ---------- ----PESSSS
NCLE        ---------- ----KVSESS KKTKKVDKKK KKK------- ---------- ----EESSSS
NCLFc1      ---------- ---------- ---------- ---------- ---------- ----------
NCLFc2      ---------- ---------- ---------- ---------- ---------- ----------
NCLFk       ---------- -----SDSDS DSDDDTKKKP TIA------- ---------- ----TIAKKD
NCLFs1      ---------- ---SSSRDSD SDSDDPKKTA VKP------- ---------- ----KVAAKK
NCLFs2      ---------- ---------- ---------- ---------- ---------- ----------
NCLFs3      ---------- ---------- ---------- ---------- ---------- ----------
NCLPa       ---------- -----SSSSS DSDSDD--EK PAA------- ---------- ----ASKKED
NCLPau      ---------- -----XSSDS DSDSDD---- ---------- ---------- ----------
NCLPd       ---------- -----SSSSS DSDSDDDAKK PTT------- ---------- ----AVKKED
NCLPf       ---------- -----SSSSS DSDSDDDEKK PTP------- ---------- ----AATKKE
NCLPh       ---------- -----SSSSS SSDSDSGNAT KKA------- ---------- ----ATKTKD
NCLPp       ---------- -----SSSDS DSDSDDEDAK KKP------- ---------- ----TEAKND
NCLNpa      ---------- ---------- ---------- ---------- ---------- ----------
NCLNp       ---------- -----SSSSS DSDSDDDDDE KKK------- ---------- ----PDTSSG
NCLNi1      ---------- ---------- ---------- ---------- ---------- ----------
NCLNi2      ---------- ---------- ---------- ---------- ---------- ----------
NCLPsp      ---------- -------SDS DSSSSDSDSD SDA------- ---------- ----APPAKK
NCLTv       ---------- -------AAT PVAAKAPVVA KAK------- ---------- ----VSDSSD
NCLTs       ---------- -------AAA PVAAKAPVVA KAK------- ---------- ----VSDSSD
NCLTc       ---------- -------SDS DNEKKVVKAE PEK------- ---------- ----VSEDSD
NCLNsa      ---------- -------APK KATPAAAKKE KLT------- ---------- ----KKKAAP
NCLNg       ---------- -------APK KATPAAGKKE KLT------- ---------- ----KKKAAP
NCLTmi      ---------- -------ATK VLKAAAKK-- ---------- ---------- ----------
NCLCr       ---------- ----GSSDSD SDSDSDSD-- ---------- ---------- ----SEKKVE
NCLBs       ---------- ---------- ---------- ---------- ---------- ----------
NCLMc       -AKKSKKKSA PAKKESSDSD SDSSSEEEKK VAKKTTKKAK KTTKKAKKPA PVKESSSDSD
NCLSl       ---------- ---------- ---------- ---------- ---------- ----------
NCLCsu      KSVSKSAVAD PSSSDDSSGE DASPENFKSK PVKDTQRKTG AAPAKGTQKA VTPTAESSSD
NCLEv       ---------- ---------- ---------- ---------- ---------- ----------
NCLPg       ---------- ---------- ---------- ---------- ---------- ----------
NCLEt       KVGVP--LKK TAPPNDSGSD DSSSEDEKPA PKKAQPKAK- --AAPAQKKA SVLQDSDSDE
NCLEn       KAGVP--QKK IAPPNDSGSD DTSSEDEKPA PKKAQPKAK- --AAPAQKKA AVLQDSDSDE
NCLEm       KVTAAPNQKK PAPPQSSDSD SS--EEDGQP TRKTAAPVKG KPTAAPQKKA PAPQDSDSDE
NCLEb       KVTAAATQKK AAPPASSDSD SS--EEERHP AKKAPPPAKG KSAAPAQKKA PVSKDSDSDD
NCLEmi      KVTAAATQKK PVPPQSSDSD SS--EEERPP PKKAPAPVKG KPAAAPQKKA PVSNDSDSDD
NCLEa       KLTAVAAQKK GAPPQDSDSD SSS-EEERPP PKKAAAPAKG KPAAAPQKKA PLPEDSDSDD
NCLEp       KMTAAPAPKK AAPPQSSDSD SSSEEEEKPQ AKKAPAPVKG KPAAAPQKKP AVPQDSDSDD
NCLPo1      PAKKAPVKKA PVEESSSDDD SSSSEDEKPA PKKAKK-AAA PAASKKVVES EDSSSSDSSD
```

```
NCLPo2        PAKKAPVKKA PVEESSSDDD SSSSEDEKPA PKKAKKAAAA PAASKKVVES EDSSSSDSSD
NCLPm         ---------- --ESSSSDDD SSSSEDEKPA AKKAKK---- ---------- ----------
NCLCc1        KQSLAQNQAS NVVSK----- ------KELQ KKIAAEKLSK QDIVFKKKTV LRPRKENS--
NCLCc2        KQNLAKNQVS KAAFKTKPLV KAP-PVQQPI KKKANAKAKA ANLKAPVAPV AEPSSDDSDE
NCLSmi        ---------- ---------- ---------- ---------- ---------- ----PVSHGA
NCLPbr        ---------- -----VKQAK KEKKKGKKTT AAP------- ---------- ----EPAPAP
NCLGt         ---------- ---------- ---------- ---------- ---------- ----------
NCLG          ---------- -----EESSS DEDEKPAAKK ATPAKADKK- ---------- --ATPAKKAA
NCLDlu        -AKESSDDSD GDSDDSSDDE SSDGPAKAPV VKPPVASGS- ---------- --LIDGAAQT
NCLP          ---------- ---------- ---------- ---------- ---------- ----------

              ....|....| ....|....| ....|....| ....|....| ....|....| ....|....|
                  245        255        265        275        285        295

NCLHs         KKMAPPPKEV EEDSEDEEMS EDEEDDSSGE EVVIPQKKGK KAAATSAKKV VVSPTKKVAV
NCLAt1        KD-------- -----DSSSS DDDSSDEE-- VAVTKK---- ---------- ----------
NCLAt2        KV-------- -----ESSSS DDDSSSDEET VPVKKQ---- ---------- ----------
Nsr1pSc       SS-------- -----SSSSS DSESEAETK- ---------- ---------- ----------
NCLAs         DS-------- -----SSSSS SSDSDDEKK- ---------- ---------- ----------
NCLAsu        EG-------- -----SSSES SSSSESEKVV AKKSV----- ---------- ----------
NCLCd         SS-------- -----SDSSS DSDEEEKPK- ---------- ---------- ----------
NCLCp         KE-------- -----DSSSS DSDSSSS--- -EEEK----- ---------- ----------
NCLCh         DS-------- -----DDS-S DSDSSSD--- -DEAAP---- ---------- ----------
NCLCw         ---------- -----TKKET SSSSSS---- ---------- ---------- ----------
NCLCm         ---------- --------TA VKSKKKE--- ---------- ---------- ----------
NCLCcr        ---------- ---------- ---------- ---------- ---------- ----------
NCLEa         DS-------- -----SDDSS SDDSDD---- ---------- ---------- ----------
NCLEs         VA-------- -----AAKKD DSSSDS---- ---------- ---------- ----------
NCLO          SD-------- -----SDSSS DSDSSSDSDS DDEKKE---- ---------- ----------
NCLOs         SS-------- -----SSSDS DSDSDSD--- -EEMKD---- ---------- ----------
NCLMp         SD-------- -----SSDSS DSDSDDEDE- ---------- ---------- ----------
NCLPi         S--------- ---------- KSKAKKEEEK PAAPKK---- ---------- ----------
NCLDf         DS-------- -----SDSDS DSESDSDK-- ---------- ---------- ----------
NCLDc         ES-------- -----SDSSS SSSSSSSEEE APKKK----- ---------- ----------
NCLPj         ---------- ---------- ---------- ---------- ---------- ----------
NCLLda        SD-------- -----DDSSS DDDSSSDDDS SSDEEK---- ---------- ----------
NCLCa         DS-------- -----SSSDS DSDSDEEKE- ---------- ---------- ----------
NCLCc         SS-------- -----SSSDS SSSSDSDS-- ---------- ---------- ----------
NCLChd        SS-------- -----SDS-S SSSDSSDDE- ---------- ---------- ----------
NCLCmu        ---------- ---------- ---------- ---------- ---------- ----------
NCLCn         SS-------- -----SDSDS ESDSDSD--- ---------- ---------- ----------
NCLC          KE-------- -----SSSDS ESSDSDEEM- ---------- ---------- ----------
NCLCt1        ST-------- -----DSS-S DSSDSESED- ---------- ---------- ----------
NCLCt2        ---------- ---------- ---------- ---------- ---------- ----------
NCLDbGSO10    SS-------- -----DSSDS DSDSSSD--- --DEEE---- ---------- ----------
NCLDbGSO10    TP-------- ------VEKP DGCLEVYIG- ---------- ---------- ----------
NCLSco1       ES-------- -----SDSSS SSSSSSS-ED EAPAKK---- ---------- ----------
NCLSco2       ---------- ---------- ---------- ---------- ---------- ----------
NCLSd         ES-------- -----SDSSS SSSSSSSSED EAPAKK---- ---------- ----------
NCLSj         KA-------- -----EESSS SSSSSSS-ED EAPAK----- ---------- ----------
NCLSma        ES-------- -----SDSSS SSSSSSS-ED EAPAKK---- ---------- ----------
NCLSme        KA-------- -----EESSS SSSSSSSDEE EAPAK----- ---------- ----------
NCLTa         EV-------- -----VKAES SDSSDSDSED EAPPP----- ---------- ----------
NCLTg         ---------- ---------- ---------- ---------- ---------- ----------
```

```
NCLTm       ES-------- -----SDSSS SSSSSSEEEE TKKKAP---- ---------- ----------
NCLTn       SS---EEEVK ------AKKT KTTVKKKE-- ---------- ---------- ----------
NCLTp       FG-------- -----GRWLS IKYSNSRPVT EARAPS---- ---------- ----------
NCLTo1      ---------- -----SSSSS SSSSSSSEEE APPPPK---- ---------- ----------
NCLTo2      ---------- ---------- ---------- ---------- ---------- ----------
NCLTw       EE-------- -----SSDSS SSSSDSESED EKPAKK---- ---------- ----------
NCLAg       SD-------- -----SSDS- DSDSDSDSD- ---------- ---------- ----------
NCLAf       ---------- ---------- ---------- ---------- ---------- ----------
NCLGo       KK-------- -----QESSS SSSSSSSSSD SSDSEE---- ---------- ----------
NCLScom     DS-------- -----SSDS- -DNSDSEDE- ---------- ---------- ----------
NCLSr       DS-------- ------SDSD DSSDEEEE-- ---------- ---------- ----------
NCLTan      NNKKKEEPVK KKKEEPAKKK KSEGKKKEEP VKKKKP---- ---------- ----------
NCLA1       SD-------- -----SDSSS SSDDEDEKKK TVAAKK---- ---------- ----------
NCLA2       ---------- ---------- ---------- ---------- ---------- ----------
NCLAp       SD-------- -----SDDSS SSEDEAPTKK ETPAVV---- ---------- ----------
NCLAr1      ND-------- -----DDNRD DDDDESSDED SSX------- ---------- ----------
NCLAr2      ES-------- -----SSDSE SEEEVKKPA- ---------- ---------- ----------
NCLE        SD-------- ------DSST SSSEKSPEKK KPKVVA---- ---------- ----------
NCLFc1      ---------- ---------- ---------- ---------- ---------- ----------
NCLFc2      ---------- ---------- ---------- ---------- ---------- ----------
NCLFk       SS-------- -----GSDSS SSDSDSDDDN KKDTKM---- ---------- ----------
NCLFs1      EE-------- -----SDDSS SDDSTSDDED DKAADK---- ---------- ----------
NCLFs2      ---------- ---------- ---------- ---------- ---------- ----------
NCLFs3      ---------- ---------- ---------- ---------- ---------- ----------
NCLPa       ---------- -------SDS SSSSDSD--- ---------- ---------- ----------
NCLPau      ---------- ---------- DSDSDDDE-- ---------- ---------- ----------
NCLPd       ---------- -------SDS DSDSDSDDEE EK-------- ---------- ----------
NCLPf       DS-------- -----DSSDS DSDSDSDSDD ---------- ---------- ----------
NCLPh       ---------- -------EES DSDSDGEEEK ---------- ---------- ----------
NCLPp       VS-------- -----DSSDS DSDSDDDEA- ---------- ---------- ----------
NCLNpa      ---------- ---------- ---------- ---------- ---------- ----------
NCLNp       ---------- -------KEK DTAVKEKSDS ---------- ---------- ----------
NCLNi1      ---------- ---------- ---------- ---------- ---------- ----------
NCLNi2      ---------- ---------- ---------- ---------- ---------- ----------
NCLPsp      AAV------- ----VVVATK KEAPKKEAPK KEAPKS---- ---------- ----------
NCLTv       SS-------- ----DSDDSS DSEDEAPAAK PEVKKA---- ---------- ----------
NCLTs       SS-------- ----DSDDSS DSEDEAPAAK PEVKKA---- ---------- ----------
NCLTc       SSS------- ----DSDSDS DSDSDDEKAA PAAKAA---- ---------- ----------
NCLNsa      ESS------- -----DSDDS GSESDDSSEE EEAAVK---- ---------- ----------
NCLNg       ESS------- -----DSDDS GSESDDSSDE EEAAVK---- ---------- ----------
NCLTmi      ---------- ---------- ---------- ---------- ---------- ----------
NCLCr       KK-------- -----ASPSA KRAKPAESS- ---------- ---------- ----------
NCLBs       ---------- ---------- ---------- ---------- ---------- ----------
NCLMc       SS-------- -----SSEEE KPKKAAKTKK AAKAKKA--- ---------- ----------
NCLSl       ---------- ---------- ---------- ---------- ---------- ----------
NCLCsu      DSDEEEPP-- RKTVSAPPRT AAPPQPKTGS SHKQTP---- KSQSTKRLVE PEAES----E
NCLEv       ---------- ---------- ---------- ---------- ---------- ----------
NCLPg       ---------- ---------- ---------- ---------- ---------- ----------
NCLEt       SSSDDEVQ-- AKKPVPKAKA APPQKKPAAP KDSES----- -DESSEDE-P PQAKKLAAKG
NCLEn       SSSDDGFQ-- AKKPAPKAKE APPQKKPAAP KDSDS----- -DESSEDE-P PQAKKPAAKG
NCLEm       SSSDEDAAPP PKKATAPSRA NAAAQKKAAP PSSDS----- DDSSSDEEPA PPARKPAPKA
NCLEb       SSSDEDAAPP PKKAAPPAKT NAA-QKKAAV PSSDS----- DDSSSEEEAA PPAKKPAPKA
NCLEmi      SSSEDDAPPP PKKAAVSAKA NAAAQKKAVA -SSDS----- DDSSSEEEAA PPAKKPAPKA
NCLEa       SS-DDDDAPP PKKAAAPAKP NAAPQKKTAP ASSDS----- EDSSSEEEVA PPPKKPAPKA
```

```
NCLEp       SSSDEDVAPP PKKAVAPAKP NAAPQKKAPA PASDSE---- DDSSSEEEAA PPAKKPAPKA
NCLPo1      SDSDASDMPS AKAVANAKAR LAGS------ ---------- ---------- ----------
NCLPo2      SDSDASDMPS AKAVANAKAR LAGASKKAAQ SSDSES---- ---------- --SDDDEEEK
NCLPm       ---------- ---------- ---------- ---------- ---------- ----------
NCLCc1      ---------- ------IAKT KAAK------ ---------- ---------- ----------
NCLCc2      DSSEEESAPV PAKAATITAA KATKGKAPVA PVAEPSSDDS DEDSSEEESA PMPAKAATIT
NCLSmi      DS-------- ------EESS DGDSS----- ---------- ---------- ----------
NCLPbr      VPE------- -----YSSSS ASDSDSAEDE PMPPAKE--- ---------- ----------
NCLGt       ---------- ---------- ---------- ---------- ---------- ----------
NCLG        KK-------- ------EESS SDESSSDEED AKPAAK---- ---------- ----------
NCLDlu      PGGAAAESSD ESDGDSDDSS DDSSDDDSSA DENEQN---- ---------- ----------
NCLP        ---------- ---------- ---------- ---------- ---------- ----------

            ....|....| ....|....| ....|....| ....|....| ....|....| ....|....|
               305        315        325        335        345        355

NCLHs       ATPAKKAAVT PGKKAAATPA KKTVTPAKAV TTPGKKGATP GKALVATPGK KGAAIPAKGA
NCLAt1      ---------- -------PAA AAKNGS---- ---------- ---------- -VKAKKESSS
NCLAt2      ---------- -------PAV LEKAKIESSS SDDDSSSDEE TVPMKKQTAV LEKAKAESSS
Nsr1pSc     ---------- --------KE ESKDS----- ---------- ---------- ----------
NCLAs       ---------- -------VET KAAKVEE--- ---------- ---------- ------SSSD
NCLAsu      ---------- -------AKT VAKKDE---- ---------- ---------- ------SSSS
NCLCd       ---------- --------PK KAKVESS--- ---------- ---------- ------SSSS
NCLCp       ---------- ----KPVAPP AAKKTATK-- ---------- ---------- ---------D
NCLCh       ---------- ----AKTSKP AAKKSAKKX- ---------- ---------- -----AKKPE
NCLCw       ---------- ---------- ---------- ---------- ---------- -------SSS
NCLCm       ---------- -------VKK EVKKEE---- ---------- ---------- -------SSS
NCLCcr      ---------- ---------- ---------- ---------- ---------- ----------
NCLEa       ---------- ---------- ---------- ---------- ---------- --------SD
NCLEs       ---------- ---------- ---------- ---------- ---------- ------GSSS
NCLO        ---------- ----KPKPKA KAKAVVKKD- ---------- ---------- -----DSASD
NCLOs       ---------- ----APAADE KEKKKDSS-- ---------- ---------- ---------D
NCLMp       ---------- ------KPAK KEDKPAA--- ---------- ---------- ------AAKK
NCLPi       ---------- ----KVEKKK AESSSSS--- ---------- ---------- -------SDS
NCLDf       ---------- ---------N ENKKTAA--- ---------- ---------- ------VKKK
NCLDc       ---------- ----AKSKK- EAKKEP---- ---------- -EKKKKE--- AKKVAKKEES
NCLPj       ---------- ---------- ---------- ---------- ---------- ----------
NCLLda      ---------- ----APVAAA KKKSSSSS-- ---------- ---------- ----------
NCLCa       ---------- --------KS DVKKEET--- ---------- ---------- ------AKKV
NCLCc       ---------- ---------S DSDDEKP--- ---------- ---------- ------AVTK
NCLChd      ---------- --------KE TKEKPKA--- ---------- ---------- ------KAKK
NCLCmu      ---------- ---------- ---------- ---------- ---------- ----------
NCLCn       ---------- ---------S DSDEEEK--- ---------- ---------- ------KSSK
NCLC        ---------- --------KE EPKKEEN--- ---------- ---------- ------NKKS
NCLCt1      ---------- --------EK PAEKKAA--- ---------- ---------- ------PAKK
NCLCt2      ---------- ---------- ---------- ---------- ---------- ----------
NCLDbGSO10  ---------- ----KAKAKP KESSSSSD-- ---------- ---------- ----------
NCLDbGSO10  ---------- ---------- ---------- ---------- ---------- -------NLS
NCLSco1     ---------- ----ETKKVA ETKKED---- ---------- ---------- -------SSS
NCLSco2     ---------- ---------- ---------- ---------- ---------- ----------
NCLSd       ---------- ----ETKKVA ETKKED---- ---------- ---------- -------SSS
NCLSj       ---------- -------KVV ETKKKD---- ---------- ---------- -------SSS
NCLSma      ---------- ----ETKKVA ETKKED---- ---------- ---------- -------SSS
NCLSme      ---------- -------KVA ETKKED---- ---------- ---------- -------SSS
```

```
NCLTa       ---------- -------KKK ESKKSP---- ---------- ---------A KKVVAKKAES
NCLTg       ---------- ---------- ---------- ---------- ---------- ----------
NCLTm       ---------- ----AKTKET KAKKESSPSS SDSDSSSDSD DEKKKKAPPA KKEVAKKDES
NCLTn       ---------- --------KT ESSSSDD--- ---------- ---------- ------SSDS
NCLTp       ---------- ----QKDEGC TTVFVGNL-- ---------- -SFHIDEDXK PKAVTQPSSD
NCLTo1      ---------- ----KKESKQ KAEKV----- ---------- ------VAAK KKEESSASSS
NCLTo2      ---------- ---------- ---------- ---------- ---------- ----------
NCLTw       ---------- ----SKKEEP KKEKK----- ---------- ------KAAK EK-------S
NCLAg       ---------- ------EPKK KEE-PKK--- ---------- ---------- ------VAKK
NCLAf       ---------- ---------- ---------- ---------- ---------- ----------
NCLGo       ---------- ----EGEKAP PKKKTAK--- ---------- ---------- -----KKEDD
NCLScom     ---------- ------KEVK KVEVKVT--- ---------- ---------- ------TVKK
NCLSr       ---------- --------KP KKKAEKK--- ---------- ---------- ------TEKK
NCLTan      ---------- ----AEKKKA PPAKEDS--- ---------- ---------- ------SSSS
NCLA1       ---------- ----KEAVKD DNSDSS---- ---------- ---------- ------SDDS
NCLA2       ---------- ---------- ---------- ---------- ---------- -------MTA
NCLAp       ---------- ----KKEESD DSSDDS---- ---------- ---------- ------DSDD
NCLAr1      ---------- ----DDDDDD DNDDDD---- ---------- ---------- ------DDES
NCLAr2      ---------- --------RK DSTSSE---- ---------- ---------- ---------S
NCLE        ---------- ----KKEDSS SSSDSS---- ---------- ---------- ------SSSD
NCLFc1      ---------- ---------- ---------- ---------- ---------- ----------
NCLFc2      ---------- ---------- ---------- ---------- ---------- ----------
NCLFk       ---------- ----KEVKKE VKKEES---- ---------- ---------- -------SSD
NCLFs1      ---------- ----MEVEKT EKSAKK---- ---------- ---------- EESSDSSSSD
NCLFs2      ---------- ---------- ---------- ---------- ---------- ----------
NCLFs3      ---------- ---------- ---------- ---------- ---------- ----------
NCLPa       ---------- ---------- --SSD----- ---------- ---------- ----------
NCLPau      ---------- ---------- -KKVEK---- ---------- ---------- -------KDD
NCLPd       ---------- -----VAVKK EESSDS---- ---------- ---------- -------SSV
NCLPf       ---------- --------DE KKQTPA---- ---------- ---------- -------AVK
NCLPh       ---------- -------VES KKKSDS---- ---------- ---------- -------SSS
NCLPp       ---------- ---------- -KKVET---- ---------- ---------- -------KKE
NCLNpa      ---------- ---------- ---------- ---------- ---------- ----------
NCLNp       ---------- ---------- -SDSDS---- ---------- ---------- -------SDS
NCLNi1      ---------- ---------- ---------- ---------- ---------- ----------
NCLNi2      ---------- ---------- ---------- ---------- ---------- ----------
NCLPsp      ---------- ----KAKAKK EESLSSS--- ---------- ---------- -------DSS
NCLTv       ---------- -----AAVAK KEETDSD--- ---------- ---------- -------SSS
NCLTs       ---------- -----AAVAK KEETDSD--- ---------- ---------- -------SSS
NCLTc       ---------- ----PAAAAK KEESDSD--- ---------- ---------- -------SSS
NCLNsa      ---------- ----PATPAR GKTTAPS--- ---------- ---------- ------KKAE
NCLNg       ---------- ----PATPAR GKTTAPS--- ---------- ---------- ------KKAE
NCLTmi      ---------- ---------- ---------- ---------- ---------- ----------
NCLCr       ---------- --------SS ESSSSD---- ---------- ---------- ---------S
NCLBs       ---------- --------GS DESSGS---- ---------- ---------- ------EQEY
NCLMc       ---------V KTKKAAPVKE ESSDSDSSDE EMEA------ -----PKKTE KKAKKPAPVK
NCLSl       ---------- ---------- ---------- ---------- ---------- ----------
NCLCsu      SASEHSSEEE NVPSKS--VH VTVTKAKAAE APSKQ----- -PQGK----- -SQHRYAASS
NCLEv       ---------- ---------- ---------- ---------- ---------- ----------
NCLPg       ---------- ---------- ---------- ---------- ---------- ----------
NCLEt       KTAVAAPQKK AAPSQD--PD SDETSS-DDE PPAKT----A APKAK---AS APQKKIAATA
NCLEn       KTAVAAPQKK AAPPQD--SD SDETSS-DDE PPAKP----A APKAK---AS APPKKTAATT
NCLEm       KAAPALAQKK AAAAVQ-DSD SDDSSSEEDN PPPKK--AAP APKGKSAVTA TPQKK-AAAA
NCLEb       KAAPPATQKK AAAAAQ-DSD SDDSSSEEDV PPPKKPAVAA ATKGKPAVTA AAQKK-AAPA
```

```
NCLEmi       KAASAA-QKK AAAAAQ-DSD SDDSSSEDDA PPPKK-APAA APKGKAAVT- AAAKK-APAA
NCLEa        KAAPAGTQKK AAAAAANDSE SNDSSSEDDM PPPKK-AAPA APKGKPAVTA AAQKKAAAAA
NCLEp        KAAPAAPQKK APAAAQ-DSD SDDSSSEDDT PPPKK-PAAA APKGKAAMTP AAQKK-AAAA
NCLPo1       -------KKA APAKKAEESD SSSDSSSDEE EEEKP----A AKKAKASPKK AAPKKAAAEE
NCLPo2       KPAVQAPKKA APAKKAEESD SSSDSSSDEE EE-KP----A AKKAKASPKK AAPKKAAAEE
NCLPm        ---------A APAKKA---- ---------- ---------- -------PKK AA--AAAAEE
NCLCc1       ----MKGKTP VAPVVDPSSD ESEEDSSDEE ---------E ITPAPT---K AVIAPVADPS
NCLCc2       VAKATKGKAP VAPVAEPSSD DSDEDSSEEE SAPVPAKAAT ITAAQATKGK APVAPVAEPS
NCLSmi       ---------- ---------- ---------- ---------- ---------- ---------D
NCLPbr       ---------E PKKVAAESSS DESSSSSDD- ---------- ---------- ------EKEA
NCLGt        ---------- ---------- ---------- ---------- ---------- ----------
NCLG         ---------- ----KATPAK AAPAKADKK- ---------- ---------- ---KAAKKEE
NCLDlu       ---------- ----AANGAQ PAKAAVTHDA S--------- ---------- ------SSSS
NCLP         ---------- ---------- ---------- ---------- ---------- ----------

             ....|....| ....|....| ....|....| ....|....| ....|....| ....|....|
                 365        375        385        395        405        415

NCLHs        KNGKNAKKED SDEEEDD--- ---------- -DSEEDEEDD EDEDEDEDEI EPAAMKAAAA
NCLAt1       EDDSSSEDE- --PAKKP--- ---------- ---------- -AAKIAKPAA KDSSSSDDDS
NCLAt2       SDDGSSSDEE PTPAKKEPIV VKKDSSDESS SDEETPVVKK KPTTVVKDAK AESSSSEEES
Nsr1pSc      -SSSSSDSSS DEEEE----- ---------- ------EEKE ETKKEES--- KESSSSDSSS
NCLAs        SDSSDDEKEE KKKPV----- ---------- ------AVKK EAA------- -KSSSSSSSS
NCLAsu       SESESDSDDD DAEPV----- ---------- ------PKTL ESKSEATSDS DEDDSEKGRK
NCLCd        SSSSDSDSDD EPKKG----- ---------- ------DKAA DKTTAKKTVK KESSDSSSDS
NCLCp        SSSS-DSSSD D--------- ---------- ---------- ---------- --SSDSDT--
NCLCh        SDSS-DSDSS SDEE------ ---------- -----APPAK AAVAKKKADS DSDSDSDS--
NCLCw        SDSSDSSDDD DEPPT----- ---------- ------KKQK QGQSEPMTPS N------GKR
NCLCm        DSSSSSDSSD SEDEA----- ---------- ------PVKK ESKKETKKEE VTKEEVKKEV
NCLCcr       ---------- ---------- ---------- ---------- ---------- ----------
NCLEa        SDEEPTKKEV AKKAV----- ---------- ------AKKE SSD------- -SSSSSSSDS
NCLEs        SDSDSDSSDD DSDDE----- ---------- ------DKAE TAKNAKVADA KKTEEAAGDS
NCLO         SDSSSDSDSD DDDEKME--- ---------- -DTEEKKKEE DKPKKKAAAK KESSDSDSSS
NCLOs        SDSS-DSDSS DEEEEEA--- ---------- -----PKKKK EEAPKKKEES SSGSDSDS--
NCLMp        DDSSDSSDSS DSSDS----- ---------- ------EDEK -KADVKAKK- -EKADDGSSS
NCLPi        DSSSSDSDSD ---------- ---------- ---DEKAKTK AAPAKTAAKT EKKDDSDSDS
NCLDf        VESSESSSDS DSSSD----- ---------- ------ESED EKEVSKPEK- -AAESTDSSS
NCLDc        SSSSSSDSSD SEAEA----- ---------- ------PAKK AVAKKEAAKK EESSSSSSDS
NCLPj        ---------- ---------- ---------- ---------- ---------- ----------
NCLLda       SSSSEDEESS SDEDEP---- ---------- -----KKAND DKKTGSSGST SSDSSSDS--
NCLCa        MEEAKKEVKK ESDGS----- ---------- ------SSSS ---------- --SSDSEMED
NCLCc        KEEAAPASKA DAS-S----- ---------- ------DSSS ---------- --SSDSDSSS
NCLChd       EESSSSSS-D SSD------- ---------- --------DD EPK------- --AKTTKAET
NCLCmu       ---------- ---------- ---------- ---------- ---------- ----------
NCLCn        TEEKKVDVKK DSD-S----- ---------- ------SSDS ---------- --SSDSSSDS
NCLC         EDSSSSSSDS SSSDS----- ---------- ------ESDD DDK------- -PSEKDKAVE
NCLCt1       EDSSSSSSSD SSDSE----- ---------- ------SEDE KPAEKK---- -PAEKAAAEV
NCLCt2       ---------- ---------- ---------- ---------- ---------- ----------
NCLDbGSO10   SDSS-DSDSS DDEEEE---- ---------- ------KKSK AVAAKKESSD DSSSSSDS--
NCLDbGSO10   WHVDEQTIRD TFQECG---- ---------- ----AIESIR FAEDRTTGEF KGFGHIKFQE
NCLSco1      SSSS--DSSD SEDET----- ---------- ------XKEK KTXDATPAET KSASKSVSS-
NCLSco2      ---------- ---------- ---------- ---------- ---------- ----------
NCLSd        SSSSSSDSSD SEDET----- ---------- ------PKEK KTVDATPAET KSASKSVSS-
NCLSj        SSSDS-SDSD SDDEK----- ---------- ------PKEK KTVDVTPKET KTASKSVSS-
```

```
NCLSma     SSSS--DSSD SEDET----- ---------- ------PKEK KTVDATPAET KSASKSVSS-
NCLSme     SSSDS-SDSE SEDEK----- ---------- ------PKEK -TVDATPAET KTTSKSVSSS
NCLTa      SSSDSSDSSD SDDEK----- ---------- ------APPA SAKKEAAKKS DSSSDSDSSS
NCLTg      ---------- ---------- ---------- ---------- ---------- ----------
NCLTm      SSSSSDSSSD EEEEK----- ---------- ------KATT SAKK----KK KDSSSSSSDS
NCLTn      SDD--DSSED EDQKN----- ---------- ------SSSK EKNKAKVEKV SKDKDSDDSS
NCLTp      SSSDSDSSSD EEEEK----- ---------- ------AAPA AKKEKTPEKK EEAEKEGSSS
NCLTo1     SSSSDSDSSD DEPTE----- ---------- ------KKEE PKEQVAAKKD ESSG-SSSSS
NCLTo2     ---------- ---------- ---------- ---------- ---------- ----------
NCLTw      SSSSDSSGSE DEAPK----- ---------- ------KKEE TKK---AKED SSSGEDSSGS
NCLAg      EESSDSDSSD SDSDS----- ---------- ------DDDK PKSKVEPKKV AKKEESSDSS
NCLAf      ---------- ---------- ---------- ---------- ---------- ----------
NCLGo      SSSDSDSSSD SDTST----- ---------- ------KSSD PPPKKKEDDS SSDSDSSSDS
NCLScom    SDDSDSSSES SSDSD----- ---------- ------SDEK EPAKTTAVKV DEKKVSSSSS
NCLSr      TEK----KTE KKTEK----- ---------- ------KTEK KADSKKKKEV AKKESSSDSS
NCLTan     SDDSSDSSSD EEETK----- ---------- ------KKAP EKTTKKTPEK TKKADSSSDS
NCLA1      SDDDDSDDED DKETE----- ---------- ------KKAA PVKKEAPAKM EVDDDDDDSS
NCLA2      STDPIHDDEE VLVGK----- ---------- ------QKKK KKEGAADPKD QDDEPVESNN
NCLAp      SDSDDSDSDD DDDDD----- ---------- ------KKTA -VKKED--KM PVDKKEDDDS
NCLAr1     SDEEEEEPKT KSGHE----- ---------- ------KKAG PSKKDSSTKM EVDDDGDD--
NCLAr2     SS----DSSS SEEEV----- ---------- ------KKPA RKDSTSSESS SXSSSSEEEV
NCLE       SDSDDSDDDE KKVVV----- ---------- ------EKEE PKKAK---KM DVEKASSSSS
NCLFc1     ---------- ---------- ---------- ---------- ---------- ----------
NCLFc2     ---------- ---------- ---------- ---------- ---------- ----------
NCLFk      SSSDDDSSSD EENEK----- ---------- ------KEVV PATKVTAKKD ESSDDSSGSD
NCLFs1     SDSSSSESED EEEEA----- ---------- ------PKVA SKKAEIKKKP DDSSSSSSSD
NCLFs2     ---------- ---------- ---------- ---------- ---------- ----------
NCLFs3     ---------- ---------- ---------- ---------- ---------- ----------
NCLPa      ---DDSSDDE DEKAK----- ---------- ------AAP- AKAKAAVKKD DSSD-SDSSD
NCLPau     KK-SSDSDS- ---------- ---------- ---------- ---------- ----------
NCLPd      SDSDSDSDDE EEKAK----- ---------- ------AAP- AKAAAAVKKE DSS--SDSSD
NCLPf      KEESDSSDSD SDSDS----- ---------- ------DDDE KKQTPAAVKK EES-------
NCLPh      SDDDSSSDDE AEDTK----- ---------- ------AAEP TKSKAVVKKD DSSSSSDSSD
NCLPp      KSDSSDSDSD SSSDD----- ---------- ------DSSS AEEEKNAPKK SKAAAKDYSS
NCLNpa     ---------- ---------- ---------- ---------- ---------- ----------
NCLNp      DSDDDEDEKE DSKAK----- ---------- ------AKKV SEDKAEKEDK SDSDSSSSSS
NCLNi1     ---------- ---------- ---------- ---------- ---------- ----------
NCLNi2     ---------- ---------- ---------- ---------- ---------- ----------
NCLPsp     SDSSSDSSSD SSSDSSS--- ---------- -DSEATAPKK AAPAPVSTKN -KKDDSDSSS
NCLTv      -DSDSDSDSE DEKPA----- ---------- ---AKPEVKK APAAAVAKKD -DSDSSSSDS
NCLTs      -DSDSDSDSE DEKPA----- ---------- ---AKPEVKK APAAAVAKKD -DSDSSSSDS
NCLTc      SDSDSDSDSD DEKAAPA--- ---------- -AKAAPAAKA APAAAAKKEE SDSDSSSSDS
NCLNsa     PESSSSESSD GSDEE----- ---------- -------EEE EEAAAKPKAA EESSDESSSD
NCLNg      PESSSSESSD SSDEE----- ---------- -------EEE -VVAAKPKAA EESSDESSSD
NCLTmi     -ESASSE--- ---------- ---------- ---------- ---------- -ESSDE----
NCLCr      SDSSDGDSSD DEKAA----- ---------- ------PAPA AAAAASSDSS S-SSDSDSSD
NCLBs      KSSSKKYEHD SDSDDP---- ---------- -------RYK GSSSKKNKGY GSPGSSGS--
NCLMc      ESSSDSDSSS EEEEK----- ---------- ------PKKT AAKKKAAPAK EESSESDSSS
NCLSl      ---------- ---------- ---------- ---------- ---------- ---MPKNQKA
NCLCsu     DSSDGD--SS DDE------- ---------- -TCVKSKKNV TSAKVVKSAI AVKGQVSDDS
NCLEv      ---------- ---------- ---------- ---------- ---------- ----------
NCLPg      ---------- ---------- ---------- ---------- ---------- ----------
NCLEt      EDSDSDESDS EESSSDY--- ---------- -EAPPAKKPA AKAKAAPTKK VAAAPD---S
NCLEn      EDSGSDESDS EESSSDY--- ---------- -EAPTGKKPA AKAKAAPTKK VAAAPD---S
```

```
NCLEm          EDSDSDDSSS EDE------- ---------- -AVPATKKPA SKSAAAQRKA TAAAAMSEDS
NCLEb          QDSDSDESSS EEE------- ---------- -ATPAPKKPA PKAAPAAQKK AAAAAMSEDS
NCLEmi         QDSDSDYSSS EEE------- ---------- -AVPATKKPA PKAAAAAQKK AAAAAMSEDS
NCLEa          QDSSSDDSSS EEE------- ---------- -VAAPAKKPA PKAAAAANKN --AAAMPEES
NCLEp          QDSDSDDSSS EEE------- ---------- -AAPAPKKPA PKAAAATQKK TAAAAMSEDS
NCLPo1         SSSSDSDSSE DEAP------ ---------- -----AKKPA AKAKAAPKKA AAEESSSSDS
NCLPo2         SSSSDSDSSE DEAP------ ---------- -----AKKPA AKAKAAPKKA AAEESSSSDS
NCLPm          SDSS-SDSSD DEE------- ---------- ------EKPA TKKAAAKKAA VAEESSSSDS
NCLCc1         SDESEEDSSD EEEA------ ---------- -IPAPTKAVT APTAKVAKGK T---------
NCLCc2         SDESDEDSS- EEES------ ---------- -APVPAKAAT ITAAKATKGK APVAPVAEPS
NCLSmi         STSSDSESSD DEVQQVS--- ---------- -KAVAATPAK IPVDSSSDDD SDSSSSESE-
NCLPbr         DVSSSDDESS SSDDEPE--- ---------- -PPKKVAKAA PAPAKEVKAS SSEDSSSSEE
NCLGt          ---------- ---------- ---------- ---------- ---------- ----------
NCLG           SSSEEESSDE DEKPAAK--- ---------- -KATPAAPAP AKADKKKAAK KEESSSEEES
NCLDlu         SSSDDDDDDD DDDDDDD--- ---------- -DDDDDDDDD DDSESSPSSD KKAKEEEKEG
NCLP           ---------- ---------- ---------- ---------- ---------- ----------

               ....|....| ....|....| ....|....| ....|....| ....|....| ....|....|
                  425        435        445        455        465        475

NCLHs          APASEDEDDE DDEDDEDDDD DEEDDSEEEA METTPAKGKK AAKVVPVKAK NVAEDEDEEE
NCLAt1         DEDSEDEKPA TK-------- ---------- ---------- ---------- ----------
NCLAt2         SSDDEPTPAK KP-------- ---------- ---------- ---------- ----------
Nsr1pSc        ---------- ---------- ---------- ---------- ---------- ----------
NCLAs          S----SSDSD ---------- ---------- ---------- ---------- ----------
NCLAsu         EDD------- ---------- ---------- ---------- ---------- ----------
NCLCd          SSDSESEEEK ---------- ---------- ---------- ---------- -----STPKA
NCLCp          -ASS--EEPK AK-------- ---------- ---------- ---------- --------PK
NCLCh          -SSSDTKSSE EP-------- ---------- ---------- ---------- --------PK
NCLCw          KRE------- ---------- ---------- ---------- ---------- ----------
NCLCm          S--------- ---------- ---------- ---------- ---------- ----------
NCLCcr         ---------- ---------- ---------- ---------- ---------- ----------
NCLEa          SDDSDSSDSD ---------- ---------- ---------- ---------- ----------
NCLEs          DSD------- ---------- ---------- ---------- ---------- ----------
NCLO           KSDSSSDSDD DE-------- ---------- ---------- ---------- --------EK
NCLOs          -SSSSDEEDA KA-------- ---------- ---------- ---------- --------PA
NCLMp          S---SSSDSD ---------- ---------- ---------- ---------- ----SSSD--
NCLPi          DSSSSS---- ---------- ---------- ---------- ---------- ----------
NCLDf          DSDSDSDDKE ---------- ---------- ---------- ---------- -----GEKKM
NCLDc          SDSEAEAP-- ---------- ---------- ---------- ---------- ----------
NCLPj          ---------- ---------- ---------- ---------- ---------- ----------
NCLLda         -SDSEDEPDP VA-------- ---------- ---------- ---------- --------KA
NCLCa          AEXKKTEAAK ---------- ---------- ---------- ---------- -----PTEAK
NCLCc          DS-------- ---------- ---------- ---------- ---------- ----------
NCLChd         KK----ASDS ---------- ---------- ---------- ---------- -----SSDSS
NCLCmu         ---------- ---------- ---------- ---------- ---------- ----------
NCLCn          DSDSDDDDAK ---------- ---------- ---------- ---------- -----SNDKK
NCLC           KSKSG-SSSS ---------- ---------- ---------- ---------- -----SSSSS
NCLCt1         KKDDGSDSDS ---------- ---------- ---------- ---------- -----SSSSS
NCLCt2         ---------- ---------- ---------- ---------- ---------- ----------
NCLDbGSO10     ---SDSDSDD EV-------- ---------- ---------- ---------- --------KE
NCLDbGSO10     TESSDKAVAL SG-------- ---------- ---------- ---------- ----------
NCLSco1        ---------- ---------- ---------- ---------- ---------- ----------
NCLSco2        ---------- ---------- ---------- ---------- ---------- ----------
```

```
NCLSd      ---------- ---------- ---------- ---------- ---------- ----------
NCLSj      ---------- ---------- ---------- ---------- ---------- ----------
NCLSma     ---------- ---------- ---------- ---------- ---------- ----------
NCLSme     ---------- ---------- ---------- ---------- ---------- ----------
NCLTa      DEEEKP---- ---------- ---------- ---------- ---------- ----------
NCLTg      ---------- ---------- ---------- ---------- ---------- ----------
NCLTm      SDSEEEAP-- -A-------- ---------- ---------- ---------- ----------
NCLTn      DDDDSTKSSE ---------- ------APEE ---------- ---------- ----AAEEKP
NCLTp      DSDSSSDSKD EA-------- ---------- ---------- ---------- ----------
NCLTo1     DSSESEDEKP EA-------- ---------- ---------- ---------- ----------
NCLTo2     ---------- ---------- ---------- ---------- ---------- ----------
NCLTw      DSSDSEDEKE DD-------- ---------- ---------- ---------- ----------
NCLAg      S---DSSDSD ---------- ---------- ---------- ---------- ----SSDDEE
NCLAf      ---------- ---------- ---------- ---------- ---------- ----------
NCLGo      ---------- ---------- ---------- ---------- ---------- ----------
NCLScom    SSSSDSSDSD ---------- ---------- ---------- ---------- ----SSDEEE
NCLSr      DSDDSSSDSD ---------- ---------- ---------- ---------- -----SDEKP
NCLTan     DSDSSSSSSE EEKAPVSKKE KGSKRKAMDS ---------- ---------- ----DSEEKP
NCLA1      SSSSGSSSSE SEPAKKEE-- ---------- ---------- ---------- -----TKKEE
NCLA2      NDDTNTKRKR KR-------- ---------- ---------- ---------- ----------
NCLAp      DSDS-SSSSE SE-------- ---------- ---------- ---------- --------EE
NCLAr1     -----SDSTD DD-------- ---------- ---------- ---------- ----------
NCLAr2     ---------- ---------- ---------- ---------- ---------- ----------
NCLE       DDSSSSSDSD DD-------- ---------- ---------- ---------- ------DKKD
NCLFc1     ---------- ---------- ---------- ---------- ---------- ----------
NCLFc2     ---------- ---------- ---------- ---------- ---------- ----------
NCLFk      SSDSDD--DD DD-------- ---------- ---------- ---------- ----------
NCLFs1     SDSDEEEEK- -V-------- ---------- ---------- ---------- ----------
NCLFs2     ---------- ---------- ---------- ---------- ---------- ----------
NCLFs3     ---------- ---------- ---------- ---------- ---------- ----------
NCLPa      SSDSDS-SDD ED-------- ---------- ---------- ---------- ----------
NCLPau     ---------- ---------- ---------- ---------- ---------- ----------
NCLPd      SSDSDS-SDD ED-------- ---------- ---------- ---------- ----------
NCLPf      ---------- -D-------- ---------- ---------- ---------- ----------
NCLPh      SSDSDS-SDD ED-------- ---------- ---------- ---------- ----------
NCLPp      SSDSDSNSDD ED-------- ---------- ---------- ---------- ----------
NCLNpa     ---------- ---------- ---------- ---------- ---------- ----------
NCLNp      SSDSDSDDDD DD-------- ---------- ---------- ---------- ----------
NCLNi1     ---------- ---------- ---------- ---------- ---------- ----------
NCLNi2     ---------- ---------- ---------- ---------- ---------- ----------
NCLPsp     DSSSDS---- ---------- ---------- ---------- ---------- ----------
NCLTv      DSDSDSEDEK PA-------- ---------- ---------- ---------- ----------
NCLTs      DSDSDSEDEK PA-------- ---------- ---------- ---------- ----------
NCLTc      DSDSDSDDEK AA-------- ---------- ---------- ---------- ----------
NCLNsa     SEEEEEEEEK ---------- ---------- ---------- ---------- ----------
NCLNg      SEEEEEEEEE EE-------- ---------- ---------- ---------- ----------
NCLTmi     --EEEEEEEE ---------- ---------- ---------- ---------- ----------
NCLCr      ---------- ---------- ---------- ---------- ---------- ----------
NCLBs      --DSHPQKR- ---------- ---------- ---------- ---------- ----------
NCLMc      EEESEPKKAK KT-------- ---------- ---------- ---------- ----------
NCLSl      VVATKKEETK DNSRRQSMDE QKNRRQQKEQ KNVKEPVTKK NVPPPKVVDD SDDESGNESE
NCLCsu     DDSDDSSDEE SPPRVAHKKS EGTA------ ---------- ---------- QQKGAILKKA
NCLEv      ---------- ---------- ---------- ---------- ---------- ----------
NCLPg      ---------- ---------- ---------- ---------- ---------- ----------
```

```
NCLEt        SDSSSSDEEP KTKKAAPKAK VGPP------ -QKKPTAAKN SDSDESSS-- EDDVPPATKP
NCLEn        SDSSSSDEEP QAKKAAPKAK AGPP------ -QKKPTVAKN SDSDESSS-- EDDGPPAKKP
NCLEm        DDSSDEDEAP PTKKPVAGLA KGKPGAAAH- --KKQPEHED SDSDDDSS-E EEAVPPPKKP
NCLEb        DDSSSEEEAP PAKKPTAAVA KGKP-AAAP- --KKQPVQED SDSDDDSSSE EEAAPPAKKP
NCLEmi       DDSSSEDEAP PAKKPAVAAT KGKAAAPAP- --KKQPVQED SDSDDDSS-E EEAAPPAKKP
NCLEa        DESSDEDEAP PAKKPAAPAN KGKATAPAAA APKKQPVQQD SDSEDDSS-E EDEEPPAKKP
NCLEp        DDSSDEDEAP PAKKPAAAVA KGKATTTAPA --KKQPAQED SDSDEDSS-E EEAAPPAKKA
NCLPo1       DSSEDEAPAK KAAPKTKAAA KKAE------ ---------- ---------E ESSSSDSDSD
NCLPo2       DSSEDEAPAK KAAPKAKAAA KKAE------ ---------- ---------E ESSSSDSDSD
NCLPm        DDEE----ME EAKPVAKKAA AKKA------ ---------- ---------E ESDSSDSEDE
NCLCc1       ---------- LEITPAPTKA AAT------- ---------- -------TIA DEETTDD---
NCLCc2       SDDSDEDSSE EESAPVPAKA AAAS------ ---------- -------AVA DEETTDDEDS
NCLSmi       ---DESAAHS AQ-------- ---------- ---------- ---------- --------NG
NCLPbr       DDDEEPTKNG AT-------- ---------- ---------- ---------- ----------
NCLGt        ---------- ---------- ---------- ---------- ---------- ----------
NCLG         SDDEKPAAKK QA-------- ---------- ---------- ---------- --------VG
NCLDlu       GGNHASDSDS DD-------- ---------- ---------- ---------- ----------
NCLP         ---------- ---------- ---------- ---------- --------MG LADDLLSGVA

             ....|....| ....|....| ....|....| ....|....| ....|....| ....|....|
                485        495        505        515        525        535

NCLHs        DDEDEDDDDD EDDEDDDDED DEEEEEEEEE EPVKEAPGKR KKEMAKQKAA PEAKKQKVEG
NCLAt1       KAAPAAAKAA SSSDSSDEDS DEE-SEDEKP AQKKADT--K ASKK-SSSDE SS-ESEEDES
NCLAt2       TVVKNAKPAA KDSSSSEEDS DEEESDDEKP PTKKAKVSSK TSKQESSSDE SSDESDKEES
Nsr1pSc      ---------- ---------S SSSDSESE-- ---------- ---------- ---KEESNDK
NCLAs        ---------- ----SSDDDD S-STKASDTP A--------- ---------A MEVDES-SSK
NCLAsu       ---------- ---ASSSSEG SSSDEEEDVE P--------- -AKNNKRKN- --EDVSATAV
NCLCd        KEIKKEESDS ---SSSSDSS SDSDSDSDSE D--------- ---------V ID-------K
NCLCp        AVAAKRAKEE KAGSSSSSSS DSSDSSS-SD SD-------- --------SD SDAEAAPAAK
NCLCh        AAPAAATKPP KKAASSSDSS SSSDSDSXSD SS-------- -------SDE EDDKEKPVAK
NCLCw        ---------- ---ETSTPNG HTTPKRPAVT P--------- -DED------ ----------
NCLCm        ---------- ----DSSSDS SDSDSSDEDE K--------- -SNDTKESEI P----ATNSK
NCLCcr       ---------- ---------- ---------- ---------- ---------- ---NGGG---
NCLEa        ---------- ----SDDSDS SDSTKKSEPP AN-------- --------DD NEAENSGSSK
NCLEs        ---------- ---SSSSSSS SSSDSDSDSD D--------- -DDDSTKSSN PPDDMDEDEK
NCLO         EKPKAKKEEV KKGSSSSDSS SSSDSDSDSD DD-------- --------DS TKSSEPPAAD
NCLOs        KKAAPQSKKD ESDSSSSSDS DSSDSDS-DD DDEKKDATAA KKKAKSTSDD TSSSSAPSSD
NCLMp        ---------- --------SD SDSDSDSDND T--------- -KDSDLPDVT EED-----SK
NCLPi        ---------- -------SSS SSSDSDDDDT ---------- ---------- -KDSDPPASK
NCLDf        EVEKEDSSSS ---SSDSDSS SDSDDDDDTK S--------- ---------S DPPIEKKESK
NCLDc        --VKKAKAEE KKKD--SDSG SSSSSDSSSD S--------- -DDEKE-AEV P---DGVPSK
NCLPj        ---------- ---------- ---------- ---------- ---------- ----------
NCLLda       KTAAAVTKEK SDDSSSDSSS SSSDDESEKE KTKASPSASK KE---SESSS SDSSSSSSDE
NCLCa        AVEKEDSSS- ---DSSSDSS SDSDSDSDSD S--------- ---------T KS-SDAPNSK
NCLCc        ---------- ---DSDSDSD EDT-KESDTP M--------- ---------T EE-KDAG-SN
NCLChd       S--------- ---SSDSDSS SDSDSDEDTK ---------- ---------E SDTPAASKKR
NCLCmu       ---------- ---------- ---------- ---------- ---------- ---NGGG---
NCLCn        EIVKEAKATA AKEDSSSDSS SDS-SDSDSD S--------- ---------T QS-SEAP-VK
NCLC         S--------- ---SSDSDSS -DSDSDSTAE S--------- ---------E VPEAPVSKKR
NCLCt1       S--------- ---SSDSDSS SDSDSDSDSE ---------- ---------D TKDSAVPESK
NCLCt2       ---------- ---------- ---------- ---------- ---------- ----------
NCLDbGSO10   EKMPGAKKSS NDTTESSSVA SSSDSSSSDE DD-------- --------DE KEEEKANPKK
NCLDbGSO10   ------IDVL GKTMKVQYAL GKPDVPKQEE Q--------- --------TY KSSEVSNGGK
```

| | | | | | | |
|---|---|---|---|---|---|---|
| NCLSco1 | ---------- | ----GSSSSS | S----DSDSD | ---------- | -DDDTKESDV | P-ADDGVPSK |
| NCLSco2 | ---------- | ---------- | --------AP | PSDN-GAPAD | DTNNAP---- | ---SGGG--- |
| NCLSd | ---------- | ----GSSSSS | SSSSSDSDSD | ---------- | -DDDTKESDV | P-ADDGVPSK |
| NCLSj | ---------- | ----GSSSSS | SSSDSDSD-- | ---------- | -DEDTKESEV | PAADEGVPSK |
| NCLSma | ---------- | ----GSSSSS | S----DSDSD | ---------- | -DDDTKESDV | P-ADDGVPSK |
| NCLSme | ---------- | ----GSSSSS | SSSSSDSDSD | S--------- | -DDDTEESAV | P-ADDGVPSK |
| NCLTa | --VAKKEEKA | VAKKEDSSSS | SSSSSGSNSD | S--------- | -SSDSD-ADV | P---SGVPSK |
| NCLTg | ---------- | ---------- | ---------- | ---------- | ---------- | -----XEEHV |
| NCLTm | --AKKTKVES | KKDDDGSSSS | SSSDSDSSSD | S--------- | -EDETEKSDV | P---VGVPTK |
| NCLTn | SKEGKTKAVS | KDEDSSSDSD | SSSDDSSSDD | S--------- | TASSEAPEDV | DMKDTESKKR |
| NCLTp | --PAKEEAKK | EERKDESGSS | SSSDSDSDSD | SDSST----- | -EESEVPAGV | PTKKRKAESV |
| NCLTo1 | ----KKAKVE | DVKKDESGSS | SDSDSDSDSD | SSSGSDSSD- | SEDEKEDSEE | AEKTSNGDSK |
| NCLTo2 | ---------- | ---SRSRSRS | RSPDRGADAP | PADTSGAPPA | DDAGAPPPQA | DNGGGGG--- |
| NCLTw | ----KKAKEA | TKKKSESSSS | SDS-SDSE-- | ---------- | -EDETKESEA | PATTAATPNK |
| NCLAg | PA----KKKV | EAKKDDSSSS | SDSDSDSDDS | T--------- | -KSSEAPGDT | EMENVDTTSK |
| NCLAf | ---------- | ----SERDRS | RSRSPEPSSN | QPHGEDRISG | DDHGAPSSDG | ANGGGHGGAD |
| NCLGo | ---------- | ----DTSTKS | SDPPATMDVD | ---------- | -DDKDFKRKT | SSATEEPPVK |
| NCLScom | AAPVKKKEVV | NKKDDDASPS | SDNSSTSDDS | T--------- | -QSSKLPEEV | VMEETPVESN |
| NCLSr | KKAEAKKADT | KKKDDS-DSD | SDSDDSSDDD | T--------- | -KSSDVPEDD | KMEEATKKRK |
| NCLTan | TKKVDKKKQA | AKEDSSDDDS | SSSDDSSSSD | SDS------- | TKSSEAPVKS | KESATPSKKR |
| NCLA1 | TKKEETKKET | KPKEEDSDSD | SS-SDSSDSS | D--------- | -DDDDDEKEN | DKVKEDVNQK |
| NCLA2 | ---------- | KRKRKDKADN | GDDDDKKDGE | D--------- | ----AATTTT | TAADDGNDLE |
| NCLAp | APKKKTVEAK | --KEEDSDSD | SSDSDSSDSD | D--------- | -SDDDDDDKE | EEAEKET--K |
| NCLAr1 | --NPAKEKPK | TAAANEEDES | SDDSDDDDED | E--------- | -DEDDSDDVP | ETPKESNGKK |
| NCLAr2 | ---------- | ---------K | KPTRKDST-- | ---------- | ---------- | --SSSSSDSS |
| NCLE | TKKAVPAKKE | AAAVKKSDSS | SSSSSSSDSD | S--------- | -SDSDDDDKK | ETMKEEPTSK |
| NCLFc1 | ---------- | ---------- | ---------- | --NG----GG | ADH------- | HNGDQNG--D |
| NCLFc2 | ---------- | ---------- | ---------- | ---------- | ---------- | ---KPTTGRR |
| NCLFk | --DNKVEKMD | VDKKDDDDSS | SDSDSDSDSD | E--------- | ---------- | --DEKASKKR |
| NCLFs1 | --APKAKAAA | KKEESDDSSS | SSSDSDSDND | D--------- | -DDDDEAEKA | P----EKSSG |
| NCLFs2 | ---------- | ---------- | ---------- | ---------- | ---------- | ---------- |
| NCLFs3 | ---------- | ---------- | ---------- | ---------- | ---------- | ---------- |
| NCLPa | --DKKEDKMD | VEKKAADSDS | SSSDSDSDSD | SDSDSDS--- | -DSDSTPASV | PPVD-DSSSK |
| NCLPau | ---------- | --------SS | SDSDSDSDSD | D--------- | -----TKASE | PPAE-KEKSK |
| NCLPd | --DKKTEKMD | VEKKASDSDS | SGSDSDSDSD | SDSDS----- | ---DSTPASV | PPVD-DSASK |
| NCLPf | --EEEADKME | VDKKSSDSDG | SSSDSDSDSD | S--------- | -----TPASV | PPVD-DSSSK |
| NCLPh | --DEKEADKK | ADKMDVDKKS | SDSDSSSDSD | SDSDS----- | ---DATPASV | PPVE-DSASK |
| NCLPp | --EKNKAKMD | VEKKSSDDSS | SSSDSDSDSE | D--------- | -----TPSSE | PPQD-SSK-K |
| NCLNpa | ---------- | ---------- | ---------- | ---------- | ---------- | ------G--A |
| NCLNp | --KDAGEKDT | EMKVADEKKE | EDSDSSSDSD | SSSDSDS--- | -DSEQTPASI | PPKETESTSK |
| NCLNi1 | ---------- | ---------- | ---------- | -------ANG | GSQGG---GG | GASDQNG-AA |
| NCLNi2 | ---------S | TVSTTMSERE | RSRSPDRGGP | PADDEDYANG | GAQGG---GG | GASDQNG-AA |
| NCLPsp | ---------- | -------SSD | SSSDSDADSD | ---------- | ---------- | -AEKAKPPSK |
| NCLTv | SKPEEKKAAV | AKKDDSDSDS | SSSDSDSDSD | ---------- | ---------- | -DEKE-TKAP |
| NCLTs | SKPEEKKAAV | AKKDDSDSDS | SSSDSDSDSD | ---------- | ---------- | -DEKEETKAP |
| NCLTc | PAAKAAPAAA | AKKEESDSDS | SSSDSDSDSD | S--------- | ------DSDS | DDEKAATSNK |
| NCLNsa | -------KAA | TKAESSESGG | ESSSSEEDED | E--------- | ---------E | EEETAVAASK |
| NCLNg | -------KAA | TKAESSESGS | ESSSSEEDED | ED-------- | ------EDEE | EEETAVAASK |
| NCLTmi | ---------A | PKADTAADG- | ---------- | ---------- | ---------- | ------ASAK |
| NCLCr | ---------- | ---------S | DSSDSEDE-- | ---------- | ---------- | --KEEEEEKP |
| NCLBs | ---------- | ---------- | ---------- | ---------- | ---------- | ---QREPASY |
| NCLMc | ---------- | -KKAAPVKES | SSSESDSSDE | APKPSKA--- | -KKSAEIESS | SESEEESEEK |
| NCLSl | DSVDLEQLIQ | AKQKKTTAAL | GKRSRGASDV | SATKKVAKKP | VQAAESDDEE | SDGSDDEKKE |
| NCLCsu | CPATKPKAP- | ------VGES | TSSEESDDSD | ENEAAQMQNK | VVNGKT---- | IRSARKEEQD |

```
NCLEv      ---------- ---------- ---------- ---------- ---------M AGPPQGIQQI
NCLPg      ---------- ---------- ---------- ---------- ---------M AN----VAQV
NCLEt      AAKSRAAAG- -PKKAATAQD PDSDDSSSEE DATERKPTAK AKAAAP---L KKAMAGDSDS
NCLEn      AAKGKATAG- -PKKAAAAQD SDSDDSSSEE EATEKKPAAK AKVAAP---L KKAVAGDSDS
NCLEm      APKAKAAALS QKKAAPVCQD SDSDDRSS-D EGNTP-PPKK AAVPK---GK AAAAAQESSS
NCLEb      APKAKAAPAV QKKAAAAAQD SDSDDSS--E EDSAP-PSKK AAAAP--KGK AAAAAQDSSS
NCLEmi     APKAKAAPA- QKKAPAPAQD SDSDDSSSDE EDSAP-PPKK AAPAA--KGK AAAVAQDSSS
NCLEa      APKAKAAAP- QKKAAAAAQD SDSDDSSSEE EDSAP-PTKK AAAAQ----- KKAAAQASSS
NCLEp      APKAKAAAP- QKKA-APAKD SDSDDSSSEE EDDAP-PPKK AAAAAPPKGK AAAAADDSSS
NCLPo1     EEMEDAKPAA KPAAKAAAAE SSDSDSDSDD EEEEAPKKVK AAAAAE---- ----EEE--S
NCLPo2     EEMEDAKPAA KAAAKAAAAE SSDSDSDSDD EEEEAPKKVK AAAVAE---- ----EEEEES
NCLPm      EEAPAAAAKK PAAAKAAAEE SSDSDSD-DE EEEEAPKKVK AAAVAE---- ----EEDDES
NCLCc1     --DVEMASGL EGVAVAIKHD SDSDSVDDED EESEDEITAN DANTSG---- --SAAKRPLV
NCLCc2     DEDVEMASGQ EVVADAANNG SDSDSIDD-D EESEDEETAD DANTNG---- --NTTKHSLA
NCLSmi     DKKRKAESS- -------ESS SSSDSSSESD SE-------- -----DSDKE DAAKPVAKKS
NCLPbr     --------EA KAASSDEDDS SDESDEEESD DE-------- --------ES GDDEEESESE
NCLGt      ---------- ---------- ---------- ---------- --------MA EASKKRKEPE
NCLG       AKKPEAAVVT KKASTKKDES SSDESSSDEE DA-------- -----KPVKK AAAKKATPAK
NCLDlu     DAPSHKAAAK AVTADESDGS SSSDDSSDDD SEDEPDSSSD DDDDDASDAP AAQPPAAGSK
NCLP       PQDRMLSLFA PPPAEPSAPV EEEDEDEDDE SEDGDADADG DGAPDTKPKA DAVLRKPRVD

           ....|....| ....|....| ....|....| ....|....| ....|....| ....|....|
               545        555        565        575        585        595

NCLHs      TEPTTAFN-- ---------- ---------- ---------- ---------- ----------
NCLAt1     EDEEETPKKK SSDVEMVDAE -KSSAKQPKT PSTPAA---- ---------- ----------
NCLAt2     KDEKVTPKKK DSDVEMVDAE QKSNAKQPKT PTNQTQ---- ---------- ----------
Nsr1pSc    KRKSE--DAE EE-------E DEESSNKKQK NEETEE---- ---------- ----------
NCLAs      KRKADDSLDN EP-------I KKAAVSG--- SEG--D---- ---------- ----------
NCLAsu     TKVAKVANIA ATNG------ ---------E VSDDAN---- ---------- ----------
NCLCd      KRKSGAVEDE PA-------T KKAAVVS--- DEG--K---- ---------- ----------
NCLCp      KRKTVEPPVA AP-------A AAPAAAADAD AEPEAT---- ---------- ----------
NCLCh      KRKVEKNDEP TP-------I KKPAETPAEE HK-------- ---------- ----------
NCLCw      ---------T ETN------- ---------- ---------- ---------- ----------
NCLCm      KRKSEAEDTT TDAKR--QAV SDEYVV---- LDDSEN---- ---------- ----------
NCLCcr     DD-ADGVK-- ---------- ---------- ---------- ---------- ----------
NCLEa      KRKADDADES EE-------N KRPAVSD--- DGA--K---- ---------- ----------
NCLEs      ESKKKRKAEA DTEDGFKDHS KKAAITPAED GSDGIN---- ---------- ----------
NCLO       DAPKSKKRKA ED-------T SPVEATPVKK AAASDD---- ---------- ----------
NCLOs      DEKKEPTPAA AG-------K KRKASNDAGD DEEDESGGAP TKKA------ ----------
NCLMp      KRKADADAGA TE-------T KKQAVED--- PNA--T---- ---------- ----------
NCLPi      KRKADDTQDE VV------HK TPKKEPE--- AAGGN----- ---------- ----------
NCLDf      KRKKEDEAET EP-------A KKPAVED--- DAV--N---- ---------- ----------
NCLDc      KRKAEAVLES TE-------D KKVAVG---- DD--EN---- ---------- ----------
NCLPj      ---------- ---------- ---------- ---------- ---------- ----------
NCLLda     DEDSVTETPV AN-------K RKAEATPQLN GSNKSS---- ---------- ----------
NCLCa      KRKADNESGN Q--------S KKVAVED--- -GV--N---- ---------- ----------
NCLCc      KRKLDDDGES AP-------T KKAAVED--- -GE--N---- ---------- ----------
NCLChd     KFDGVGEEGE AV-------S KKAAAVA--- DDDGAG---- ---------- ----------
NCLCmu     DSGVEEVK-- ---------- ---------- ---------- ---------- ----------
NCLCn      KRKATDDGGN E--------S KKAAVED--- -GE--N---- ---------- ----------
NCLC       KAEEKVQSDK PS-------A KKAAVED--- ESV--N---- ---------- ----------
NCLCt1     KRKAEDAESE PS-------A KKTAVAD--- ESE--N---- ---------- ----------
NCLCt2     ---------- ---------- ---------- ---------- ---------- ----------
```

```
NCLDbGSO10   RQRTDDAAAE SS-------F STPAKKKSAV SNGKTK---- ---------- ----------
NCLDbGSO10   KGKKEVYSPK PN-------- KENMTEEQQK LLDGYT---- ---------- ----------
NCLSco1      KRKAETVLSN DN------GD KKQHTG---- EEGGEN---- ---------- ----------
NCLSco2      ED-SEGVK-- ---------- ---------- ---------- ---------- ----------
NCLSd        KRKAETVLSN DN------GD KKQHTG---- EEGGEN---- ---------- ----------
NCLSj        KRKAETVLSN DN------GD KKQHTG---- EEGGEN---- ---------- ----------
NCLSma       KRKAETVLSN DN------GD KKQHTG---- EEGGEN---- ---------- ----------
NCLSme       KRKAETVLSN DN------GD KKQHTG---- EEGGEN---- ---------- ----------
NCLTa        KRKAEAVVAD ATEE---QPE KIAKVD---- EE--DN---- ---------- ----------
NCLTg        EKKQAVASE- ---------- ---------- ---DEN---- ---------- ----------
NCLTm        KRKAEAVTPD AADE---HAD KKMHVA---- ED--EQ---- ---------- ----------
NCLTn        KGSD---AME PP-------T KKVAIDG--- DAN------- ---------- ----------
NCLTp        SESSDVANAP AE-------- KLQKTG---- EEGDEN---- ---------- ----------
NCLTo1       KRKADSVLES -G-------- EDKKQS---- LDPEAN---- ---------- ----------
NCLTo2       DD-GEGVK-- ---------- ---------- ---------- ---------- ----------
NCLTw        KRKADTQEET NG-------- DYKRQN---- VDSDDN---- ---------- ----------
NCLAg        KRKAE-DSGE EP-------S KKVAVEG--- EN-------- ---------- ----------
NCLAf        SGNGEEVK-- ---------- ---------- ---------- ---------- ----------
NCLGo        KIKTTTATDG DD-------- ----DA---- DNSNDA---- ---------- ----------
NCLScom      KRKAEDTPVE PD-------S KRAAGEA--- PK-------- ---------- ----------
NCLSr        AGDD---DDA PP-------A KKAAGAE--- EET--E---- ---------- ----------
NCLTan       KNDDNEDSEE PP-------T KKVAVEE--- SGK--S---- ---------- ----------
NCLA1        KRKEPDTPAQ HD------TP KRAAVD---- GEG------- ---------- ----------
NCLA2        EHRTKTAQVH R--------- ---------- ---------- ---------- ----------
NCLAp        KRKEPDTAPE TP------LA KKAAVA---- DESGGE---- ---------- ----------
NCLAr1       RKADDDAPKE TP-------A KKPAVE---- SKSDGN---- ---------- ----------
NCLAr2       SSESEEEETK VT-------T PKAAASSSDE SG-------- ---------- ----------
NCLE         KRKEPDTPTT TQEDKSKSAA KRVAVDTS-G PEDDNN---- ---------- ----------
NCLFc1       TGTGEEVK-- ---------- ---------- ---------- ---------- ----------
NCLFc2       PRGPDGPK-- ---------- ---------- ---------- ---------- ----------
NCLFk        KEANDDDEQT SE-----PPM KKQKDS---T ESEGPP---- ---------- ----------
NCLFs1       KRKAEDIEAA AP------TP KKAAVS---- DEATEG---- ---------- ----------
NCLFs2       ---------- ---------- ---------- ---------- ---------- ----------
NCLFs3       ---------- ---------- ---------- ---------- ---------- ----------
NCLPa        KRKAD--DEP VE-----PAV KKQAVG---- GDEEGT---- ---------- ----------
NCLPau       KRKAD--EET SE-----PPV KKQAVN---- DGEESM---- ---------- ----------
NCLPd        KRKAD--EET VE-----PAF KKQAVG---- GDDEG----- ---------- ----------
NCLPf        KRKAD--DEE SE-----PPV KKQAVT---- DSDENT---- ---------- ----------
NCLPh        KRKAD--DEP SE-----PAT KKQAVA---- DGDEAT---- ---------- ----------
NCLPp        RKADD--EET FE-----PPV KKQAVA---- DSDGTS---- ---------- ----------
NCLNpa       GPTGDEVK-- ---------- ---------- ---------- ---------- ----------
NCLNp        KRKEESSNES SE-----PPM KKQAFSPKES ESGDSN---- ---------- ----------
NCLNi1       DGEAEGIK-- ---------- ---------- ---------- ---------- ----------
NCLNi2       DGEAEGIK-- ---------- ---------- ---------- ---------- ----------
NCLPsp       KRKAEDAPAE EP------KA KVEASSS--- SVNG------ ---------- ----------
NCLTv        KRKASDAALE EA------PA KKAAVSD--- DAGNK----- ---------- ----------
NCLTs        KRKASDAALE EA------PA KKAAVSD--- DAGNK----- ---------- ----------
NCLTc        RKAEEEAAPV AA------KA SKAEADS--- SPADAK---- ---------- ----------
NCLNsa       KRKGGEEGGQ EK------KK SKGAASP--- DED------- ---------- ----------
NCLNg        KRKGGEEGGQ EK------KK SKGAASP--- DED------- ---------- ----------
NCLTmi       KAPKAEEG-- ---------V STGAGT---- ---------- ---------- ----------
NCLCr        AAKEEEAEEA VE-------T PKPAAATPSK GGNGRG---- ---------- ----------
NCLBs        EDKGDE---- ---------- ---------- ---------- ---------- ----------
NCLMc        PQKEEHQDAE ME-------E EKQEAPAQEE AEETEG---- ---------- ----------
```

```
NCLSl       NDTENSPE-- ---------- ---------- ---------- ---------- ----------
NCLCsu      EEAESGADET G---KGKTAR KRKTPSPDDG EEEPAA-KKQ KESFSQ---- ----PKPRVQ
NCLEv       FERNQEAT-- ---------- ---------- ---------- ---------- ----------
NCLPg       YERNQEAT-- ---------- ---------- ---------- ---------- ----------
NCLEt       GSSEVEAEKQ ---PHKKTAN KKGKAADVDM EVDGSS-KKR KQGQD---HQ --QAPKKKLQ
NCLEn       GSSEEEAEKQ ---PPKKAAN KKGKAADMEM EIDGSSSKKR KQGQD---HQ --QAPKKKLQ
NCLEm       DESESEAEEP ---PKNAAPH KKGKATDANM EVDGAS-KKR KQAQAQNGNQ --QGPKKKLQ
NCLEb       DESEEEEEEQ EEPPKKAAAQ KKGKATGGSM EVDGAS-KKR KQSETQNGQQ --QGPKKKLQ
NCLEmi      DESDSDEEEE --PPKKPVAQ KKGKAADAGM EVDGGS-KKR KQTQAQNGQQ --QGPKKKLQ
NCLEa       EESSEEEEED EEPPKVAAAQ KKGKGTDGSM EVDGAS-KKR KQAQAQNGNQ PQQGPKKKLQ
NCLEp       DESEEEEEEE E-QPKKAVTQ KKGKGSDENM EVDGGS-KKR KQQQTQNGQQ --QGPKKKLQ
NCLPo1      EDSDDEMEDA P-------AA GKRTASEATT EEDEAP---- ---------- --AQKKTKMN
NCLPo2      EDSDDEMEDA P-------AA GKRTASEATT EEDEAP---- ---------- --AQKKTKMN
NCLPm       EDSDDDMEEA --------AT GKRAASEATT EEDEAP---- ---------- --AQKKAKLN
NCLCc1      SSTAEFPEKR ----QKVKSE DFAGKRERYF NADGAP---- ---------- ----------
NCLCc2      ASADGLSEKR ----QKVKNG DFGGERERYF NSDGGP---- ---------- ----------
NCLSmi      RVEPSTPAAG D--------- ---------- ---------- ---------- ----------
NCLPbr      SEKEDEPMPD ATAKRKKPDT KASSGGEPAK KSKVDE---- ---------- ----------
NCLGt       KAKKEDGAAG D--------- ---------- ---------- ---------- ----------
NCLG        KAKKEESSSD ESSSSDEEDE KPAKKKAAKD SDDEEESPKK KVKVEEKEEN A---------
NCLDlu      RVRDEYDASA GTP-----AP KRAAPASSGN GGGSDK---- ---------- ----------
NCLP        DAEKLSRT-- ---------- ---------- ---------- ---------- ----------

            ....|....| ....|....| ....|....| ....|....| ....|....| ....|....|
                 605        615        625        635        645        655

NCLHs       ---------- ---LFVGNLN FNKSAPELKT GISDV-FAKN D--------- ----------
NCLAt1      --------GG SKTLFAANLS FNIERADVEN FFKEAG---- ---------- ----------
NCLAt2      --------GG SKTLFAGNLS YQIARSDIEN FFKEAG---- ---------- ----------
Nsr1pSc     ---------- PATIFVGRLS WSIDDEWLKK EFEHIG---- ---------- ----------
NCLAs       ---------- -TKLYVRGLP WRGTEEDVQE FFKGCGAG-- ---------- ----------
NCLAsu      ---------- -CKIFVRGLP WAATEDEVED FFKGCG---- ---------- ----------
NCLCd       ---------- -LKMFVRGLP WRASEDDVRN YFTECGE--- ---------- ----------
NCLCp       ---------- --MAFIRGLP WTTGEDEVRD FFKDCG---- ---------- ----------
NCLCh       ---------- ---IIAQGIP WACTEEDVQQ FFADCG---- ---------- ----------
NCLCw       ---------- --KVFVMGLP WKATEDEVRD WFKDCG---- ---------- ----------
NCLCm       ---------- -PKVYVRGLP WKATYDEIKE FFGGCG---- ---------- ----------
NCLCcr      ---------- ---LYVGNLD YATDEPRLRE VFGEFGTVTD V--------- ----------
NCLEa       ---------- -SKIFIRGLP WRASEDEIRE HFGGCG-E-- ---------- ----------
NCLEs       ---------- -TKIFVRGLP WKTTEEEVTD FFKDCGN--- ---------- ----------
NCLO        ---------- -NVVIAKGLP WKATPEEVTD FFKDCGS--- ---------- ----------
NCLOs       ----AVESSS SHAIFVKGLP WKATEDEVRD FFASCGS--- ---------- ----------
NCLMp       ---------- -TKLYIRGLP WRATQEEVEE FFAQCGKG-- ---------- ----------
NCLPi       ---------- -TKIFIRGLP WRATEEEVMD FFKDCGDGPT SIEMPLQDDG RSSGTAIMDF
NCLDf       ---------- -LKMFVRGLP WRATEEEVRE YFATCGE--- ---------- ----------
NCLDc       ---------- -SKIYVRGLP WRATEDEVRE FFAACG---- ---------- ----------
NCLPj       ---------- ---------- ---------- ---------- ---------- ----------
NCLLda      ----KLNDDR EDSVYVRGLP WKATEEEVRE FFSECG---- ---------- ----------
NCLCa       ---------- -RKMYVRGLP WKATEDEVRD YFASCGE--- ---------- ----------
NCLCc       ---------- -KKMYIRGLP WRATEDEVRE FFSSCGE--- ---------- ----------
NCLChd      ---------- -KKMYIRGLP WRATDDELRE YFASCGE--- ---------- ----------
NCLCmu      ---------- ---LYVGNLD YDTNESKLRE VFGAFGTVTD V--------- ----------
NCLCn       ---------- -LTLFIRGLP WRATEDDLRE YFKSCGE--- ---------- ----------
NCLC        ---------- -TTMYIRGLP WKTTEDEVRD FFSSCGE--- ---------- ----------
```

```
NCLCt1       ---------- -TTMYIRGLP WRATEEEVRA FFDSCGE--- ---------- ----------
NCLCt2       ---------- ---------- ---------- ---------- ---------- ----------
NCLDbGSO10   ---------- ---VGVWGLP YSVTEEEIKD FFKDCG---- ---------- ----------
NCLDbGSO10   ---------- FTKVYMHGLP FNATEDDVRT FLQHCG---- ---------- ----------
NCLSco1      ---------- -TKIYIRGLP WRAGEDEVGT FFTSCG---- ---------- ----------
NCLSco2      ---------- ---LYVGNLD YSTDEPKLRE SFGEFGNVKD V--------- ----------
NCLSd        ---------- -TKIYIRGLP WRAGEDEVRD FFTSCG---- ---------- ----------
NCLSj        ---------- -TKIYIRGLP WRATEEEVRD FFTSCG---- ---------- ----------
NCLSma       ---------- -TKIYIRGLP WRAGEDEVRD FFTSCG---- ---------- ----------
NCLSme       ---------- -TKVYVRGLP WRATEEEVRD FFTSCG---- ---------- ----------
NCLTa        ---------- -TKIYVRGLP WRTSDEEVRE YFAACG---- ---------- ----------
NCLTg        ---------- -TKVYVRGLP WSTTEDEVKE FFATCG---- ---------- ----------
NCLTm        ---------- -TKIYIRGLP WRATEEEVRD FFASCG---- ---------- ----------
NCLTn        ---------- -TKIYIRGLP WRATEDELRD FFQTCGT--- ---------- ----------
NCLTp        ---------- -TKIYVRGLP WRATEDEVRE FFAQCG---- ---------- ----------
NCLTo1       ---------- -CKVYIRGLP WRATEDEVRE FFAECG---- ---------- ----------
NCLTo2       ---------- ---LYIGNLD YSTDEPQLRS VFGAFGAVTD V--------- ----------
NCLTw        ---------- -CKIYVRGLP WKAGEDDVKE FFKACG---- ---------- ----------
NCLAg        ---------- -TKIYIRGLP WRATEDEVRE FFQGCGKG-- ---------- ----------
NCLAf        ---------- ---LYIGNLD YQTDETKLRE EFGQFGPVTD V--------- ----------
NCLGo        ---------- -CKLYVKGLP WKATYDDVHG YFSQSSTKA- ---------- ----------
NCLScom      ---------- -TKIYIRGLP WRATEDEVRD FFSQCGQG-- ---------- ----------
NCLSr        ---------- -TKVYVRGLP WRASEEEIRE WFVSCGT--- ---------- ----------
NCLTan       ---------- -VKVFVRGLP WRASEDEVWE FFGECGK--- ---------- ----------
NCLA1        ---------- -TKIYIRGLP WRATVEEVQE FFATCG---- ---------- ----------
NCLA2        ---------- --TLYLEGLP FSYQRQQVLD FFAEHEIT-- ---------- ----------
NCLAp        ---------- -TKIYIRGLP WRASQDEVME FFTTCG---- ---------- ----------
NCLAr1       ---------- -TKIYIRGLP WRATQEEVQE YFESCG---- ---------- ----------
NCLAr2       ---------- EFPATIKGLP YEMDVDEISQ HFADCG---- ---------- ----------
NCLE         ---------- -TKVYIRGLP WRASEEEVQD FFKSCG---- ---------- ----------
NCLFc1       ---------- ---LYVGNLD YATDENRLRD EFGQFGTVTD V--------- ----------
NCLFc2       ---------- ---IYVGNLP YGVAEDDLKE LFAEHGEVVS V--------- ----------
NCLFk        ---------- -PKLYIRGLP WRADEEQVRD FFKSCGS--- ---------- ----------
NCLFs1       ---------- -NKVYVRGLP WKAEEHEVQE FFAACG---- ---------- ----------
NCLFs2       ---------- ---LYVGNVA FSTRTAHIRS HFSQLGPVKQ I--------- ----------
NCLFs3       ---------- ---LYVGNVA FSTRTAHIRS HFSQLGPVKQ I--------- ----------
NCLPa        ---------- -TKLYIRGLP WRATEDAVRD YFKACGS--- ---------- ----------
NCLPau       ---------- -TKVYIRGLP WRATEEGVQE FFAKCGS--- ---------- ----------
NCLPd        ---------- -TKLYIRGLP WRATEDGVRD FFKACGS--- ---------- ----------
NCLPf        ---------- -TKLYIRGLP WRTTEEAVKE YFQTCGS--- ---------- ----------
NCLPh        ---------- -TKLYIRGLP WRATEEAVKE FFQSCGS--- ---------- ----------
NCLPp        ---------- -TKLYIRGLP WRTTEEAVQE YFSKCGS--- ---------- ----------
NCLNpa       ---------- ---LYVGNLD YATDETRLRT EFAQFGAVTE V--------- ----------
NCLNp        ---------- -TKIYVRGLP WRATKDQVEV FFQSCGS--- ---------- ----------
NCLNi1       ---------- ---LYVGNLD YQTDEQRLRD EFSQFGTVTE V--------- ----------
NCLNi2       ---------- ---LYVGNLD YQTDEQRLRD EFSQFGTVTE V--------- ----------
NCLPsp       ---------- -HKVFVRGFP WATTENEIND FFQSCG---- ---------- ----------
NCLTv        ---------- --KLYVRGLP WKATEDEVRD FFKECG---- ---------- ----------
NCLTs        ---------- --KLYVRGLP WKATEDEVRD FFKECG---- ---------- ----------
NCLTc        ---------- -LKLYVRGLT WTAGEADVRD FFKDCG---- ---------- ----------
NCLNsa       ---------- -RKVFVQGLP WSATEEEVRA FFEKCG---- ---------- ----------
NCLNg        ---------- -RKVFVQGLP WSATEEEVRA FFQKCG---- ---------- ----------
NCLTmi       ---------- --KIYVRGLP WSATEEEVRD FFKECG---- ---------- ----------
NCLCr        ---------- EFAVTVRGLP YEMDEDSILE LFATSG---- ---------- ----------
```

```
NCLBs         ---------- QSEVFVGGLS WEADDSDLQK FFKNCG---- ---------- ----------
NCLMc         ---------- PTELFVGNLS WNVDDAYLKE TFEYYG---- ---------- ----------
NCLSl         ---------- ---IFVGNIP FTMENDALKE LFEEFGTIEN V--------- ----------
NCLCsu        VKGGGVVEKM RTEIFCGGLP YATTESELKE LFEADCG--- ---------- ----------
NCLEv         ---------- ---IYVGNLD PKIDEEILWE LFCQCGPVAN L--------- ----------
NCLPg         ---------- ---IYVGNLD PKIDEEVLWE LMVQCGPVAN V--------- ----------
NCLEt         LTDQSAAARM RSEIFCGGLP YSVTEEQLKE LFEADCG--- ---------- ----------
NCLEn         LTDQSAAARM RSEIFCGGLP YSVTEEQLKE LFEADCG--- ---------- ----------
NCLEm         LTDQSAAARM RSEIFCGGLP YSVTEEQLKD LFEADCG--- ---------- ----------
NCLEb         LTDQSAAARM RSEIFCGGLP YSVTEEQLKD LFESDCG--- ---------- ----------
NCLEmi        LTDQSAAARM RSEIFCGGLP YSVTEEQLKD LFEADCG--- ---------- ----------
NCLEa         LTDASAAARM RSEIFCGGLP YSVTEEQLKD LFEADCG--- ---------- ----------
NCLEp         LTDQSAAARM RSEIFCGGLP YSVTEEQLKD LFEADCG--- ---------- ----------
NCLPo1        DGSAQDSVER SKSVFVGNLP FSMTKEWLEQ IFS-WCG--- ---------- ----------
NCLPo2        DGSAQDSVER SKSVFVGNLP FSMTKEWLEQ IFS-WCG--- ---------- ----------
NCLPm         DGSAQESVER SKSVFVGNLP FSMTKEWLEQ IFS-WCG--- ---------- ----------
NCLCc1        --SKKVQLKP RVEIFCGGLP FQVTEDQIRE LFENDCG--- ---------- ----------
NCLCc2        --SKKVQLKA RVEIFCGGLP FQVTEDQIRD LFENDCG--- ---------- ----------
NCLSmi        ------AEAE QFVAVVRGLP WSAGEDDVKA CFT--G---- ---------- ----------
NCLPbr        --------SA KSSLFIGGLS YEATEDDLRD LFAECG---- ---------- ----------
NCLGt         ---------- KRRVFLGGLP FKATEKDIKK MFESCG---- ---------- ----------
NCLG          --EEKAVGVE NRRVFLGGLP FRATEDEIKD MFKSCG---- ---------- ----------
NCLDlu        ---------- -PTAMLTGLP HEVTDAAIRA FLAPAGCG-- ---------- ----------
NCLP          ---------- ---VFIGNVP VSTTQKVLKR LFTECGQIDS IRIRSAAFAN KKISRKAA--

              ....|....| ....|....| ....|....| ....|....| ....|....| ....|....|
                   665        675        685        695        705        715

NCLHs         ---------- ---------- ---------- ---------- ---------- ----------
NCLAt1        ---------- ---------- ---------- ---------- ---------- ----------
NCLAt2        ---------- ---------- ---------- ---------- ---------- ----------
Nsr1pSc       ---------- ---------- ---------- ---------- ---------- ----------
NCLAs         ---------- ---------- ---------- ---------- ---------- ----------
NCLAsu        ---------- ---------- ---------- ---------- ---------- ----------
NCLCd         ---------- ---------- ---------- ---------- ---------- ----------
NCLCp         ---------- ---------- ---------- ---------- ---------- ----------
NCLCh         ---------- ---------- ---------- ---------- ---------- ----------
NCLCw         ---------- ---------- ---------- ---------- ---------- ----------
NCLCm         ---------- ---------- ---------- ---------- ---------- ----------
NCLCcr        ---------- ---------- ---------- ---------- ---------- ----------
NCLEa         ---------- ---------- ---------- ---------- ---------- ----------
NCLEs         ---------- ---------- ---------- ---------- ---------- ----------
NCLO          ---------- ---------- ---------- ---------- ---------- ----------
NCLOs         ---------- ---------- ---------- ---------- ---------- ----------
NCLMp         ---------- ---------- ---------- ---------- ---------- ----------
NCLPi         ANATGFDAGL ALNGETFGER WLSIKAERVS FLVILNLSCC FDFYAFRETE VLPSFFXANF
NCLDf         ---------- ---------- ---------- ---------- ---------- ----------
NCLDc         ---------- ---------- ---------- ---------- ---------- ----------
NCLPj         ---------- ---------- ---------- ---------- ---------- ----------
NCLLda        ---------- ---------- ---------- ---------- ---------- ----------
NCLCa         ---------- ---------- ---------- ---------- ---------- ----------
NCLCc         ---------- ---------- ---------- ---------- ---------- ----------
NCLChd        ---------- ---------- ---------- ---------- ---------- ----------
NCLCmu        ---------- ---------- ---------- ---------- ---------- ----------
```

```
NCLCn        ---------- ---------- ---------- ---------- ---------- ----------
NCLC         ---------- ---------- ---------- ---------- ---------- ----------
NCLCt1       ---------- ---------- ---------- ---------- ---------- ----------
NCLCt2       ---------- ---------- ---------- ---------- ---------- ----------
NCLDbGSO10   ---------- ---------- ---------- ---------- ---------- ----------
NCLDbGSO10   ---------- ---------- ---------- ---------- ---------- ----------
NCLSco1      ---------- ---------- ---------- ---------- ---------- ----------
NCLSco2      ---------- ---------- ---------- ---------- ---------- ----------
NCLSd        ---------- ---------- ---------- ---------- ---------- ----------
NCLSj        ---------- ---------- ---------- ---------- ---------- ----------
NCLSma       ---------- ---------- ---------- ---------- ---------- ----------
NCLSme       ---------- ---------- ---------- ---------- ---------- ----------
NCLTa        ---------- ---------- ---------- ---------- ---------- ----------
NCLTg        ---------- ---------- ---------- ---------- ---------- ----------
NCLTm        ---------- ---------- ---------- ---------- ---------- ----------
NCLTn        ---------- ---------- ---------- ---------- ---------- ----------
NCLTp        ---------- ---------- ---------- ---------- ---------- ----------
NCLTo1       ---------- ---------- ---------- ---------- ---------- ----------
NCLTo2       ---------- ---------- ---------- ---------- ---------- ----------
NCLTw        ---------- ---------- ---------- ---------- ---------- ----------
NCLAg        ---------- ---------- ---------- ---------- ---------- ----------
NCLAf        ---------- ---------- ---------- ---------- ---------- ----------
NCLGo        ---------- ---------- ---------- ---------- ---------- ----------
NCLScom      ---------- ---------- ---------- ---------- ---------- ----------
NCLSr        ---------- ---------- ---------- ---------- ---------- ----------
NCLTan       ---------- ---------- ---------- ---------- ---------- ----------
NCLA1        ---------- ---------- ---------- ---------- ---------- ----------
NCLA2        ---------- ---------- ---------- ---------- ---------- ----------
NCLAp        ---------- ---------- ---------- ---------- ---------- ----------
NCLAr1       ---------- ---------- ---------- ---------- ---------- ----------
NCLAr2       ---------- ---------- ---------- ---------- ---------- ----------
NCLE         ---------- ---------- ---------- ---------- ---------- ----------
NCLFc1       ---------- ---------- ---------- ---------- ---------- ----------
NCLFc2       ---------- ---------- ---------- ---------- ---------- ----------
NCLFk        ---------- ---------- ---------- ---------- ---------- ----------
NCLFs1       ---------- ---------- ---------- ---------- ---------- ----------
NCLFs2       ---------- ---------- ---------- ---------- ---------- ----------
NCLFs3       ---------- ---------- ---------- ---------- ---------- ----------
NCLPa        ---------- ---------- ---------- ---------- ---------- ----------
NCLPau       ---------- ---------- ---------- ---------- ---------- ----------
NCLPd        ---------- ---------- ---------- ---------- ---------- ----------
NCLPf        ---------- ---------- ---------- ---------- ---------- ----------
NCLPh        ---------- ---------- ---------- ---------- ---------- ----------
NCLPp        ---------- ---------- ---------- ---------- ---------- ----------
NCLNpa       ---------- ---------- ---------- ---------- ---------- ----------
NCLNp        ---------- ---------- ---------- ---------- ---------- ----------
NCLNi1       ---------- ---------- ---------- ---------- ---------- ----------
NCLNi2       ---------- ---------- ---------- ---------- ---------- ----------
NCLPsp       ---------- ---------- ---------- ---------- ---------- ----------
NCLTv        ---------- ---------- ---------- ---------- ---------- ----------
NCLTs        ---------- ---------- ---------- ---------- ---------- ----------
NCLTc        ---------- ---------- ---------- ---------- ---------- ----------
NCLNsa       ---------- ---------- ---------- ---------- ---------- ----------
NCLNg        ---------- ---------- ---------- ---------- ---------- ----------
```

```
NCLTmi       ---------- ---------- ---------- ---------- ---------- ----------
NCLCr        ---------- ---------- ---------- ---------- ---------- ----------
NCLBs        ---------- ---------- ---------- ---------- ---------- ----------
NCLMc        ---------- ---------- ---------- ---------- ---------- ----------
NCLSl        ---------- ---------- ---------- ---------- ---------- ----------
NCLCsu       ---------- ---------- ---------- ---------- ---------- ----------
NCLEv        ---------- ---------- ---------- ---------- ---------- ----------
NCLPg        ---------- ---------- ---------- ---------- ---------- ----------
NCLEt        ---------- ---------- ---------- ---------- ---------- ----------
NCLEn        ---------- ---------- ---------- ---------- ---------- ----------
NCLEm        ---------- ---------- ---------- ---------- ---------- ----------
NCLEb        ---------- ---------- ---------- ---------- ---------- ----------
NCLEmi       ---------- ---------- ---------- ---------- ---------- ----------
NCLEa        ---------- ---------- ---------- ---------- ---------- ----------
NCLEp        ---------- ---------- ---------- ---------- ---------- ----------
NCLPo1       ---------- ---------- ---------- ---------- ---------- ----------
NCLPo2       ---------- ---------- ---------- ---------- ---------- ----------
NCLPm        ---------- ---------- ---------- ---------- ---------- ----------
NCLCc1       ---------- ---------- ---------- ---------- ---------- ----------
NCLCc2       ---------- ---------- ---------- ---------- ---------- ----------
NCLSmi       ---------- ---------- ---------- ---------- ---------- ----------
NCLPbr       ---------- ---------- ---------- ---------- ---------- ----------
NCLGt        ---------- ---------- ---------- ---------- ---------- ----------
NCLG         ---------- ---------- ---------- ---------- ---------- ----------
NCLDlu       ---------- ---------- ---------- ---------- ---------- ----------
NCLP         ---------- ---------- ---------- ---------- ---------- ----------

             ....|....| ....|....| ....|....| ....|....| ....|....| ....|....|
                  725        735        745        755        765        775

NCLHs        ---------- ---------- ---------- ---------- ---------- ----------
NCLAt1       ---------- ---------- --EVVD---- ---------- ---------- ----------
NCLAt2       ---------- ---------- --EVVD---- ---------- ---------- ----------
Nsr1pSc      ---------- ---------- --GVIG---- ---------- ---------- ----------
NCLAs        ---------- ---------- ---PTQ---- ---------- ---------- ----------
NCLAsu       ---------- ---------- --EIVK---- ---------- ---------- ----------
NCLCd        ---------- ---------- ---MKT---- ---------- ---------- ----------
NCLCp        ---------- ---------- --EMTT---- ---------- ---------- ----------
NCLCh        ---------- ---------- --EILS---- ---------- ---------- ----------
NCLCw        ---------- ---------- --EIAT---- ---------- ---------- ----------
NCLCm        ---------- ---------- --KIKS---- ---------- ---------- ----------
NCLCcr       ---------- ---------- ---------- ---------- ---------- ----------
NCLEa        ---------- ---------- ---IKE---- ---------- ---------- ----------
NCLEs        ---------- ---------- --GPKV---- ---------- ---------- ----------
NCLO         ---------- ---------- --GPSD---- ---------- ---------- ----------
NCLOs        ---------- ---------- --GPTS---- ---------- ---------- ----------
NCLMp        ---------- ---------- ---PKS---- ---------- ---------- ----------
NCLPi        KHFCLTYMII YVAHNSQNNF TPALPNXTFS LKPFLFHKTL TMDKTINIPI SAXNHFFNCF
NCLDf        ---------- ---------- ---IES---- ---------- ---------- ----------
NCLDc        ---------- ---------- --EITT---- ---------- ---------- ----------
NCLPj        ---------- ---------- ---------- ---------- ---------- ----------
NCLLda       ---------- ---------- --TITK---- ---------- ---------- ----------
NCLCa        ---------- ---------- ---ITL---- ---------- ---------- ----------
NCLCc        ---------- ---------- ---MDS---- ---------- ---------- ----------
```

```
NCLChd       ---------- ---------- ---IVS---- ---------- ---------- ----------
NCLCmu       ---------- ---------- ---------- ---------- ---------- ----------
NCLCn        ---------- ---------- ---ITS---- ---------- ---------- ----------
NCLC         ---------- ---------- ---IEH---- ---------- ---------- ----------
NCLCt1       ---------- ---------- ---MVS---- ---------- ---------- ----------
NCLCt2       ---------- ---------- ---------- ---------- ---------- ----------
NCLDbGSO10   ---------- ---------- --TIQS---- ---------- ---------- ----------
NCLDbGSO10   ---------- ---------- --DIKN---- ---------- ---------- ----------
NCLSco1      ---------- ---------- --EMTS---- ---------- ---------- ----------
NCLSco2      ---------- ---------- ---------- ---------- ---------- ----------
NCLSd        ---------- ---------- --EMTS---- ---------- ---------- ----------
NCLSj        ---------- ---------- --EMTS---- ---------- ---------- ----------
NCLSma       ---------- ---------- --EMTS---- ---------- ---------- ----------
NCLSme       ---------- ---------- --EMAS---- ---------- ---------- ----------
NCLTa        ---------- ---------- --EVTS---- ---------- ---------- ----------
NCLTg        ---------- ---------- --EMTS---- ---------- ---------- ----------
NCLTm        ---------- ---------- --EIES---- ---------- ---------- ----------
NCLTn        ---------- ---------- ---IVS---- ---------- ---------- ----------
NCLTp        ---------- ---------- --EMES---- ---------- ---------- ----------
NCLTo1       ---------- ---------- --EIKS---- ---------- ---------- ----------
NCLTo2       ---------- ---------- ---------- ---------- ---------- ----------
NCLTw        ---------- ---------- --DIVN---- ---------- ---------- ----------
NCLAg        ---------- ---------- ---PTN---- ---------- ---------- ----------
NCLAf        ---------- ---------- ---------- ---------- ---------- ----------
NCLGo        ---------- ---------- --KVVS---- ---------- ---------- ----------
NCLScom      ---------- ---------- ---PKL---- ---------- ---------- ----------
NCLSr        ---------- ---------- ---ITN---- ---------- ---------- ----------
NCLTan       ---------- ---------- ---IVT---- ---------- ---------- ----------
NCLA1        ---------- ---------- --DIAS---- ---------- ---------- ----------
NCLA2        ---------- ---------- --DVVD---- ---------- ---------- ----------
NCLAp        ---------- ---------- --PIES---- ---------- ---------- ----------
NCLAr1       ---------- ---------- --TVTA---- ---------- ---------- ----------
NCLAr2       ---------- ---------- --EVVR---- ---------- ---------- ----------
NCLE         ---------- ---------- --EITL---- ---------- ---------- ----------
NCLFc1       ---------- ---------- ---------- ---------- ---------- ----------
NCLFc2       ---------- ---------- ---------- ---------- ---------- ----------
NCLFk        ---------- ---------- --GPTT---- ---------- ---------- ----------
NCLFs1       ---------- ---------- --PIKS---- ---------- ---------- ----------
NCLFs2       ---------- ---------- ---------- ---------- ---------- ----------
NCLFs3       ---------- ---------- ---------- ---------- ---------- ----------
NCLPa        ---------- ---------- --GPKS---- ---------- ---------- ----------
NCLPau       ---------- ---------- --GPKT---- ---------- ---------- ----------
NCLPd        ---------- ---------- --GPKS---- ---------- ---------- ----------
NCLPf        ---------- ---------- --GPTF---- ---------- ---------- ----------
NCLPh        ---------- ---------- --GPKS---- ---------- ---------- ----------
NCLPp        ---------- ---------- --GPKS---- ---------- ---------- ----------
NCLNpa       ---------- ---------- ---------- ---------- ---------- ----------
NCLNp        ---------- ---------- --GPRS---- ---------- ---------- ----------
NCLNi1       ---------- ---------- ---------- ---------- ---------- ----------
NCLNi2       ---------- ---------- ---------- ---------- ---------- ----------
NCLPsp       ---------- ---------- --EIES---- ---------- ---------- ----------
NCLTv        ---------- ---------- --EIES---- ---------- ---------- ----------
NCLTs        ---------- ---------- --EIES---- ---------- ---------- ----------
NCLTc        ---------- ---------- --EMVA---- ---------- ---------- ----------
```

```
NCLNsa      ---------- ---------- --PMTS---- ---------- ---------- ----------
NCLNg       ---------- ---------- --PMTS---- ---------- ---------- ----------
NCLTmi      ---------- ---------- --EIES---- ---------- ---------- ----------
NCLCr       ---------- ---------- --EVVR---- ---------- ---------- ----------
NCLBs       ---------- ---------- --SITN---- ---------- ---------- ----------
NCLMc       ---------- ---------- --EIKR---- ---------- ---------- ----------
NCLSl       ---------- ---------- ---------- ---------- ---------- ----------
NCLCsu      ---------- ---------- --PTTR---- ---------- ---------- ----------
NCLEv       ---------- ---------- ---------- ---------- ---------- ----------
NCLPg       ---------- ---------- ---------- ---------- ---------- ----------
NCLEt       ---------- ---------- --PTTR---- ---------- ---------- ----------
NCLEn       ---------- ---------- --PTTR---- ---------- ---------- ----------
NCLEm       ---------- ---------- --PTTR---- ---------- ---------- ----------
NCLEb       ---------- ---------- --PTTR---- ---------- ---------- ----------
NCLEmi      ---------- ---------- --PTTR---- ---------- ---------- ----------
NCLEa       ---------- ---------- --PTTR---- ---------- ---------- ----------
NCLEp       ---------- ---------- --PTTR---- ---------- ---------- ----------
NCLPo1      ---------- ---------- --DIER---- ---------- ---------- ----------
NCLPo2      ---------- ---------- --DIER---- ---------- ---------- ----------
NCLPm       ---------- ---------- --DIER---- ---------- ---------- ----------
NCLCc1      ---------- ---------- --LVTR---- ---------- ---------- ----------
NCLCc2      ---------- ---------- --SVTR---- ---------- ---------- ----------
NCLSmi      ---------- ---------- --SIEE---- ---------- ---------- ----------
NCLPbr      ---------- ---------- --TISS---- ---------- ---------- ----------
NCLGt       ---------- ---------- --AIEN---- ---------- ---------- ----------
NCLG        ---------- ---------- --QVET---- ---------- ---------- ----------
NCLDlu      ---------- ---------- ---------- ---------- ---------- ----------
NCLP        ---------- ---------- ---------- ---------- ---------- ----------

            ....|....| ....|....| ....|....| ....|....| ....|....| ....|....|
                 785        795        805        815        825        835

NCLHs       ---------- ---------- ---------- ----LAVVDV RIGMTRKFGY VDFESAEDLE
NCLAt1      ---------- ---------- ---------- ---VRFSTNR DDGSFRGFGH VEFASSEEAQ
NCLAt2      ---------- ---------- ---------- ---VRLSS-F DDGSFKGYGH IEFASPEEAQ
Nsr1pSc     ---------- ---------- ---------- ---ARVIYER GTDRSRGYGY VDFENKSYAE
NCLAs       ---------- ---------- ---------- ----IELPLQ DDGRSSGTAI VDFGSAEDAA
NCLAsu      ---------- ---------- ---------- ----IDLPLS EDGRASGTAF VEFKTTSGSA
NCLCd       ---------- ---------- ---------- ----CELPLN HDGRSSGTAI IVFATKDACE
NCLCp       ---------- ---------- ---------- ----VDLPLQ SDGRSSGTAI ITFTTQAGFD
NCLCh       ---------- ---------- ---------- ----CKIPMQ DDGRSSGRAF VVFSTDEALQ
NCLCw       ---------- ---------- ---------- ----FEMPLT DEGRASGKAF ITFGTSEGYD
NCLCm       ---------- ---------- ---------- ----VDLPLL ADGRSSGTAV IEFESPAGSA
NCLCcr      ---------- ---------- ---------- ----FLPMER GTSRPRGFGF VTLATRQAAE
NCLEa       ---------- ---------- ---------- ----VEQPMN SDGRSSGTAL IVFGSASSAE
NCLEs       ---------- ---------- ---------- ----VELPLQ DDGRSSGTAI IDFGSTEGAE
NCLO        ---------- ---------- ---------- ----VDLPLD YSGRSSGTAI LKFDSSDGVN
NCLOs       ---------- ---------- ---------- ----VEMPLD HQGRASGNAI LTFSSEEDRN
NCLMp       ---------- ---------- ---------- ----VELPLQ DDGRSSGTAI LEFHSTEDAE
NCLPi       QPLWILLMPL DLMRRATEEE VMDFFKDCGD GPTSIEMPLQ DDGRSSGTAI MDFANATGFD
NCLDf       ---------- ---------- ---------- ----LELPLQ DDGRSSGTAI IVFKTKEACE
NCLDc       ---------- ---------- ---------- ----LELPMQ DDGRSSGTAI IDFKESAGAA
NCLPj       ---------- ---------- ---------- ---------- ---------- ----------
NCLLda      ---------- ---------- ---------- ----CEMPLD ATGRSSGTAI LYFEDNSATI
```

```
NCLCa        ---------- ---------- ---------- ----LELPLM DDGRSSGTAV IEYATKDACE
NCLCc        ---------- ---------- ---------- ----CELPLQ DDGRSSGTAV INFSTKEGAE
NCLChd       ---------- ---------- ---------- ----CELPLQ SDGRSSGTAE IEFSTKDACE
NCLCmu       ---------- ---------- ---------- ----FLPTER GTQRPRGFGF VTLAGRAAAE
NCLCn        ---------- ---------- ---------- ----LELPLQ DDGRSSGTAV MKFSSKKECE
NCLC         ---------- ---------- ---------- ----LELPLQ DDGRSSGTAI IKFSTVDACE
NCLCt1       ---------- ---------- ---------- ----CELPLQ DDGRSSGTAI IKFSDTAGAE
NCLCt2       ---------- ---------- ---------- ---------- ---------- ----------
NCLDbGSO10   ---------- ---------- ---------- ----IEF-QE KLGRFSGRAI VEFDSSDACE
NCLDbGSO10   ---------- ---------- ---------- ----IKFVLQ EDGRSTGRMI VEFQKIGGAK
NCLSco1      ---------- ---------- ---------- ----VELPLQ DDGRXSGTAI IQFSDAAGAA
NCLSco2      ---------- ---------- ---------- ----FLPMDR MTSRPRGFGF VTFDDRAAAE
NCLSd        ---------- ---------- ---------- ----VELPLQ DDGRSSGTAI IQFSDAAGAA
NCLSj        ---------- ---------- ---------- ----VELPLQ DDGRSSGTAI IAFSNAASAA
NCLSma       ---------- ---------- ---------- ----VELPLQ DDGRSSGTAI IQFSDAAGAA
NCLSme       ---------- ---------- ---------- ----VEMPLQ DDGRSSGTAI IAFSDAAGAA
NCLTa        ---------- ---------- ---------- ----VELPLQ DDGRSSGTAI IDFKDNAGSA
NCLTg        ---------- ---------- ---------- ----VELPLQ DDGRSSGTAI IQFNDTAGAA
NCLTm        ---------- ---------- ---------- ----CELPLM DDGRSSGTAI IAFKEATAAA
NCLTn        ---------- ---------- ---------- ----CELPLQ DDGRSSGTAV LEFESAESAA
NCLTp        ---------- ---------- ---------- ----VELPLQ DDGRSSGTAI IDFKEASAAA
NCLTo1       ---------- ---------- ---------- ----VDMPLQ DDGRSSGTAI IEFSDPSGSA
NCLTo2       ---------- ---------- ---------- ----FLPMER GTSRPRGFGF VTLSTRQAAE
NCLTw        ---------- ---------- ---------- ----IELPLM NDGRSSGTAI IEFKDSTGAA
NCLAg        ---------- ---------- ---------- ----LELPLQ DDGRSSGTAI IDFGSAEDAA
NCLAf        ---------- ---------- ---------- ----FLPMER GTQRPRGFGF VTLASREAAE
NCLGo        ---------- ---------- ---------- ----CELPVQ DDGRSSGTAI LEFESPKEAS
NCLScom      ---------- ---------- ---------- ----VELPLQ DDGRSSGTAV LDFATPECAA
NCLSr        ---------- ---------- ---------- ----CELPLQ DDGRSSGTAI LDFDSADAAV
NCLTan       ---------- ---------- ---------- ----CELPLQ DDGRSSGTAL VEFGSDAEAA
NCLA1        ---------- ---------- ---------- ----CELPLQ DDGRSSGTAI VDFKEADSAA
NCLA2        ---------- ---------- ---------- ---LRLPVWQ DSGRLRGYGH LVVQSPEDYD
NCLAp        ---------- ---------- ---------- ----CELPLQ DDGRSSGTAI LNFETAEGAA
NCLAr1       ---------- ---------- ---------- ----CELPLQ DDGRSSGTAV VEFDSPDAAS
NCLAr2       ---------- ---------- ---------- ---VDLARFE DTGKPKGACN IVFATREALD
NCLE         ---------- ---------- ---------- ----CEMPLQ DDGRSSGTAV IDFKDSDGAA
NCLFc1       ---------- ---------- ---------- ----FLPMER GTSRPRGFGF VTLSTRDAAE
NCLFc2       ---------- ---------- ---------- ----YLVKD- DDGKDRGFGF ITMANDEAVD
NCLFk        ---------- ---------- ---------- ----IELPLQ DDGRSSGTAV VEFKDTASAE
NCLFs1       ---------- ---------- ---------- ----CEMPLM DDGRSSGTAI IEFETNEGAA
NCLFs2       ---------- ---------- ---------- ----HMGLDR NKKMPCGFCF VEYYSREDAL
NCLFs3       ---------- ---------- ---------- ----HMGLDR NKKMPCGFCF VEYYSREDAL
NCLPa        ---------- ---------- ---------- ----VELPLQ EDGRSSGTAI IDFHDAASAA
NCLPau       ---------- ---------- ---------- ----VELPLQ DDGRSSGTAI VDFHDADSAA
NCLPd        ---------- ---------- ---------- ----VELPLQ EDGRSSGTAI LDFHDGASAA
NCLPf        ---------- ---------- ---------- ----VELPLQ PDGRSSGTAI IDFADAASAA
NCLPh        ---------- ---------- ---------- ----VELPLQ DDGRSSGTAI VDFHDSSSAA
NCLPp        ---------- ---------- ---------- ----VELPLQ DDGRSSGTAI LDFHDAASAA
NCLNpa       ---------- ---------- ---------- ----FLPVER GTQRPRGFGF VTLANRDAAE
NCLNp        ---------- ---------- ---------- ----VDLPLQ DDGRSSGTAI IEFEDAESAA
NCLNi1       ---------- ---------- ---------- ----FLPTER NTNRPRGFGF VTLATREAAE
NCLNi2       ---------- ---------- ---------- ----FLPTER NTNRPRGFGF VTLATREAAE
NCLPsp       ---------- ---------- ---------- ----IEQPLG DDGRASGTAI IAFKTQEGAN
NCLTv        ---------- ---------- ---------- ----CELPLD DTGRSSGTCF LVFKDTAGAE
```

```
NCLTs          ---------- ---------- ---------- ----CELPLD DTGRSSGTCF LVFKDTAGAE
NCLTc          ---------- ---------- ---------- ----CELPLS DDGRSSGTAF ITMKDQAGVD
NCLNsa         ---------- ---------- ---------- ----VELPLK D-GRSSGTAY IVFGEEDGVG
NCLNg          ---------- ---------- ---------- ----VELPLK D-GRSSGTAY IVFGEEDGVG
NCLTmi         ---------- ---------- ---------- ----CDLPTD HNGRASGTAF VVFADSDAAN
NCLCr          ---------- ---------- ---------- ---VEVSRFE DTGRPKGIAN VVFATEEGKD
NCLBs          ---------- ---------- ---------- ---IKILKN- DQGKSKGSAF IKFSSPEEAQ
NCLMc          ---------- ---------- ---------- ---ANVLIRD G--RSQGIGF VEFGNHAQAK
NCLSl          ---------- ---------- ---------- ----SVPMSG RKKK--GYAF IRFSSHEEAA
NCLCsu         ---------- ---------- ---------- ---IKMLEGK ------GIAF ITFQTEEGAQ
NCLEv          ---------- ---------- ---------- ----HLPRDK ITTTHQGYGF VEFKNEEDAD
NCLPg          ---------- ---------- ---------- ----HLPRDK ITNTHQGYGF VEFKNEEDAD
NCLEt          ---------- ---------- ---------- ---IKILEGK ------GIAF ITFESEEGAA
NCLEn          ---------- ---------- ---------- ---IKILEGK ------GIAF ITFESEEGAA
NCLEm          ---------- ---------- ---------- ---IKILEGK ------GIAF ITFASEEAAA
NCLEb          ---------- ---------- ---------- ---IKILEGK ------GIAF ITFATEEAAA
NCLEmi         ---------- ---------- ---------- ---IKILEGK ------GIAF ITFASEEAAA
NCLEa          ---------- ---------- ---------- ---IKILEGK ------GIAF ITFASEEAAA
NCLEp          ---------- ---------- ---------- ---IKILEGK ------GIAF ITFATEEAAA
NCLPo1         ---------- ---------- ---------- ---VSLPTDW ESGKIKGFAF LDFADEDSAE
NCLPo2         ---------- ---------- ---------- ---VSLPTDW ESGKIKGFAF LDFADEDSAE
NCLPm          ---------- ---------- ---------- ---VSLPTDW ESGKIKGFAF LDFADEDSAE
NCLCc1         ---------- ---------- ---------- ---VSILENR ------GVSF VTFETEEAAS
NCLCc2         ---------- ---------- ---------- ---VSILENR ------GVAF VTFETEEAAS
NCLSmi         ---------- ---------- ---------- ----VELPLD DTGRSSGTAV LTFTSRSDLD
NCLPbr         ---------- ---------- ---------- ---IRIPVFE DSGKPRGIAF IEFEEGDAAK
NCLGt          ---------- ---------- ---------- ----IELPMN ADSRPAGFGF LTFKDADSVA
NCLG           ---------- ---------- ---------- ----LELPMN NEGRPSGFGF LTFKSAAAVA
NCLDlu         ---------- ---------- ---------- -PVEIFMPGE STGQTGRGVA FVSVSSTDAL
NCLP           ---------- ---------- ---------- ----VISKAF DKDRRDTYNA YIVFVAPEGA

               ....|....| ....|....| ....|....| ....|....| ....|....| ....|....|
                   845        855        865        875        885        895

NCLHs          KALELTG-LK VF--GNEIKL EKPKGKDSKK ERDARTLLAK NLPYKVTQDE LKEVFEDAAE
NCLAt1         KALEFHGRPL LG---REIRL DIAQERGERG ERPAF----- ---------- ----------
NCLAt2         KALEMNGKLL LG---RDVRL DLANERG--- ---------- ---------- ----------
Nsr1pSc        KAIQEMQGKE ID--GRPINC DMSTSKPAGN ----N----- ---------- ----------
NCLAs          AGLAL-NGNT IGDEGRWLSI KLS-TPKAMA ---------- ---------- ----------
NCLAsu         AAIEL-NGNT FG--ERWLSI KYDTPKQQGG NF-------- ---------- ----------
NCLCd          ACIAL-NGAD FE--GRWLNI IYS-NDKPIT ---------- ---------- ----------
NCLCp          AALAL-DQAT FG--ERWLSI KQWFPKTNMG NNH------- ---------- ----------
NCLCh          AAIAM-DGQT MQ--ERWIGV RKWEPPRAS- ---------- ---------- ----------
NCLCw          KAISL-DGEY FG--ERWLGV K--PAKSSSA FK-------- ---------- ----------
NCLCm          AAMEQ-NGAD FG--GRWLNI KYS-SNKPVT ---------- ---------- ----------
NCLCcr         TAISKMDQSQ LD--GRTIRV NESKPRGEGP AD-S------ ---------- ----------
NCLEa          SAVEM-NGGD FG--GRWLDI KLA-DDKPIQ ---------- ---------- ----------
NCLEs          AAIAL-NGAT YGDSGRWLNI KYSTSKP--- IT-------- ---------- ----------
NCLO           AALAL-NGET FGDSGRWLKI EKYDCKPQAQ AVS------- ---------- ----------
NCLOs          VAIEM-NGAT FGDSGRWLNV AEYTAPAAST YNK------- ---------- ----------
NCLMp          AGIAL-NGAD FN--GRWLSI KYS-TPKPIN ---------- ---------- ----------
NCLPi          AGLAL-NGET FG--ERWLSI KAN-GDKPVH ---------- ---------- ----------
NCLDf          ACLAC-NGED FN--GRWLNI KYS-TPKPIF ---------- ---------- ----------
NCLDc          AAMEQ-NGAD FG--GRWLSI KYS-SSKPIG ---------- ---------- ----------
```

```
NCLPj        ---------- ---------- ----GGGGGG GG-------- ---------- ----------
NCLLda       KALEL-DQAT FG--ERWLSV KPNTKK---- ---------- ---------- ----------
NCLCa        ACLKL-NGED FN--GRWLSI KYS-SPKPIN ---------- ---------- ----------
NCLCc        ACLAL-NGED FG--GRWLNI KYS-TPKPVT ---------- ---------- ----------
NCLChd       KCMAL-NGED FG--GRWLSI KYA-SDKPAF ---------- ---------- ----------
NCLCmu       EAIAKMDQAQ LD--SRTIRV NESRPRGEGP ED-------- ---------- ----------
NCLCn        ACLQL-DGED FD--GRWLSI KYS-TPKPIN ---------- ---------- ----------
NCLC         ACVQM-NGQD FN--GRWVHV KYS-SPKPII ---------- ---------- ----------
NCLCt1       ACLAL-NGED FN--GRWLNI KYS-NSKPIH ---------- ---------- ----------
NCLCt2       ---------- ---------- ------GIPT PC-------- ---------- ----------
NCLDbGSO10   AAIKL-NKMT FFDSGRWVGV EDADAPAQTR TPT------- ---------- ----------
NCLDbGSO10   SALLL-NNEV FDSTERYVVV RQWTDNPVHR GKNGQ----- ---------- ----------
NCLSco1      AAMEH-NGAD FN--GRWLXI KYS-XXKPVT ---------- ---------- ----------
NCLSco2      SAISKMDQSQ LD--GRTIRV NESKPRGEGP ADRG------ ---------- ----------
NCLSd        AAMEH-NGAD FN--GRWLNI KYS-TSKPVT ---------- ---------- ----------
NCLSj        AAMEH-NGAD FN--GRWLNI KYS-TSKPVT ---------- ---------- ----------
NCLSma       AAMEH-NGAD FN--GRWLNI KYS-SAKPVT ---------- ---------- ----------
NCLSme       AAMEH-NGAD FG--GRWLNI KYS-SSKPIT ---------- ---------- ----------
NCLTa        AAFEL-NGAD YG--GRWLSI KYS-SNKPIN ---------- ---------- ----------
NCLTg        ASIEQ-NGAD FG--GRWLSI KYS-TNKPVN ---------- ---------- ----------
NCLTm        AALEH-NGAD FG--GRWLNI KYS-SSKPIS ---------- ---------- ----------
NCLTn        KGIEL-NGED FQ--GRWLSI KYS-SAKPIT ---------- ---------- ----------
NCLTp        AALEQ-NGAD FG--GRWLSI KYS-NSRPVT ---------- ---------- ----------
NCLTo1       SALEH-NGAD FG--GRWLNI KYS-TSRPIT ---------- ---------- ----------
NCLTo2       DAIAKMDQSQ LD--GRTIRV NESRPRGEGP G--A------ ---------- ----------
NCLTw        SALEH-NGAD FQ--GRWLNI KYS-TSKSIT ---------- ---------- ----------
NCLAg        AAMEL-NGAD FN--GRWLSI KYS-TPKPIL ---------- ---------- ----------
NCLAf        KAISKMDQAQ LD--GRTIRV NESRPKGERS S--------- ---------- ----------
NCLGo        AVMEECQGAD FD--GRWLNI SLY-TQRKPS RM-------- ---------- ----------
NCLScom      AAMEL-NGAD FH--GRWLSI QFS-SPKPVL ---------- ---------- ----------
NCLSr        KAIEL-NGED FQ--GRWLSI KYS-SPKPIT ---------- ---------- ----------
NCLTan       KAIEM-NGAD FQ--GRWLDI KLDGMQKPAA ---------- ---------- ----------
NCLA1        AAIE-LNGQD FQ--GRWLSI KYS-TPKPIL ---------- ---------- ----------
NCLA2        KALSLSGRAL PG-SHRYVTI QPAQAPKSPT SSG------- ---------- ----------
NCLAp        AALE-LNGQD FQ--GRWLSI KYS-TPKPIL ---------- ---------- ----------
NCLAr1       KAIEDLNGQD FQ--GRWLSI KYS-TPKAIL ---------- ---------- ----------
NCLAr2       NCLARD-GET IG--KRWLSI SEPRAFT--- ---------- ---------- ----------
NCLE         AAIA-LNGED FQ--GRWLSI QLS-TPKPIM ---------- ---------- ----------
NCLFc1       KAISKMDQAQ LD--GRTIRV NESRPKGSAA PD-------- ---------- ----------
NCLFc2       VAVAKIDGSF FQ--GRRLAA RRPLAEGEKS EQ-------- ---------- ----------
NCLFk        AAIEL-NGAD FE--GRWLSI KYS-TPKSII ---------- ---------- ----------
NCLFs1       AAIEL-NGGD FQ--GRWLSI KYS-TPKPIL ---------- ---------- ----------
NCLFs2       AAVSLLSSTK LD--GQVIRV EKQKQNSIMS GR-------- ---------- ----------
NCLFs3       AAVSLLSSTK LD--GQVIRA TKQKQNSIMS GR-------- ---------- ----------
NCLPa        AALEL-NGAD FE--GRWLSI KYS-TPKPIL ---------- ---------- ----------
NCLPau       AALEL-NGAD FE--GRWLSI KYS-SPKPIL ---------- ---------- ----------
NCLPd        AAMEL-NGAD FE--GRWLSI KYS-TPKPIL ---------- ---------- ----------
NCLPf        AAMEL-NGAD FE--GRWLSI KYS-TPKPIL ---------- ---------- ----------
NCLPh        AALEL-NGAD FE--GRWLSI KYS-SPKPIL ---------- ---------- ----------
NCLPp        AALEL-NGAD FE--GRWLSI KYS-TPKPIL ---------- ---------- ----------
NCLNpa       RAITQMDQSQ LD--GRTIRV NESRPKGESR TR-------- ---------- ----------
NCLNp        AAIEL-NGAD FD--GRWLSI KYS-TPKPIL ---------- ---------- ----------
NCLNi1       KAISKMDQAQ LD--GRTIRV NESRPKGERG P--------- ---------- ----------
NCLNi2       KAISKMDQAQ LD--GRTIRV NESRPKGERG P--------- ---------- ----------
```

```
NCLPsp      KSYEL-DRAT FG--ERWLSV KEN-SEKPK- ---------- ---------- ----------
NCLTv       QGLAL-DQAT FG--ERWLSV KYA-TDKPQG ---------- ---------- ----------
NCLTs       QGLAL-DQAT FG--ERWLSV KYA-TDKPQG ---------- ---------- ----------
NCLTc       AALAL-DHAD FQ--GRWLSI KLS-TEKPKT ---------- ---------- ----------
NCLNsa      KAVAM-NEEI FGDSGRWVRV RKYEPVAAGK DGGKG----- ---------- ----------
NCLNg       KAVAM-NEEI FGDSGRWVRV RKYEPTAAGK DGGKG----- ---------- ----------
NCLTmi      KS-------- ---RERWLKI VLATERAARP SFG------- ---------- ----------
NCLCr       NAIARD-GET CG--ARWLSI GEKRLR---- ---------- ---------- ----------
NCLBs       EAIRLNGSEH MG---RTLRI NLSGDKPNKH ---------- ---------- ----------
NCLMc       AALEQANEYE LD--GRPMRV SFSADKP--- ----N----- ---------- ----------
NCLSl       AAVKGMNNKE IE--GRELKC NFSSGKVVEK KP-------- ---------- ----------
NCLCsu      KAVEY-NNTQ YN--GRTLRI NLTADKQQNQ QTRAGDRGER GGG------- ----------
NCLEv       YAIKIMNMIK FF--GKPIRC NKSSQ----- ---------- ---------- ----------
NCLPg       YTVKIMNMIR LF--GKPIRC NKSSQ----- ---------- ---------- ----------
NCLEt       KAVEF-NQTE YE--GRTLRI NLSAERQQGD R--------- ---------- ----------
NCLEn       KAVEF-NQTE YE--GRTLRI NLSADRQQGD R--------- ---------- ----------
NCLEm       KAVEF-NQTE YE--GRTLRI NLSADRQQQQ Q--------- ---------- ----------
NCLEb       KAVEF-NQTE YE--GRTLRI NLSADRQQQQ QQ-------- ---------- ----------
NCLEmi      KAVEF-NQTE YE--GRTLRI NLSADRQQQQ Q--------- ---------- ----------
NCLEa       KAVEF-NQTE YE--GRVLRI NLSADRQQQQ H--------- ---------- ----------
NCLEp       KAVEF-NQTE YE--GRTLRI NLSADRQQQQ Q--------- ---------- ----------
NCLPo1      KAVGK-NGED CE--GRDLRI NYSFPKND-- HGGKG----- ---------- ----------
NCLPo2      KAVGK-NGED CE--GRDLRI NYSFPKND-- HGGKG----- ---------- ----------
NCLPm       KAVGK-NGED CE--GRDLRV NYSFPKNDNA HSGKG----- ---------- ----------
NCLCc1      KACEF-NKTT YE--GRSLRI NYADEKP--- ---------- ---------- ----------
NCLCc2      KACEF-NKTT YD--GRSLRI NYADEKP--- ---------- ---------- ----------
NCLSmi      AALAM-DGAT FPGSERWMKV TEGRPSRKSV GAA-P----- ---------- ----------
NCLPbr      KGLAKDGAEI RG---RWIKV QEASAKREPQ ---------- ---------- ----------
NCLGt       KAVAM-DGQE LMG--RWVKV KEADGTEGSA GKK-P----- ---------- ----------
NCLG        KAVAM-DGQE LQG--RWIKV KEADGEEKNK APG-R----- ---------- ----------
NCLDlu      AKLIAHSGKQ LG--GRAVGI REAVSADGGA ---------- ---------- ----------
NCLP        QAALGKNGTV VD--EKTLRV DKATAPRVGQ GP-------- ---------- ----------

            ....|....| ....|....| ....|....| ....|....| ....|....| ....|....|
                905        915        925        935        945        955

NCLHs       IRLVSKDGKS KGIAYIEFKT EADAEKTFEE KQGTEIDGRS ISLYYTGEKG QNQDYRGGKN
NCLAt1      ---------- ---------- ---------- ---------- ---------- -TPQSGNFRS
NCLAt2      ---------- ---------- ---------- ---------- ---------- -TPRNSNPGR
Nsr1pSc     ---------- ---------- ---------- ---------- ---------- -DRAKKFGDT
NCLAs       ---------- ---------- ---------- ---------- ---------- ---NVREASQ
NCLAsu      ---------- ---------- ---------- ---------- ---------- ---SPRPVSK
NCLCd       ---------- ---------- ---------- ---------- ---------- ---APRPTSE
NCLCp       ---------- ---------- ---------- ---------- ---------- ---NVS---E
NCLCh       ---------- ---------- ---------- ---------- ---------- ---------E
NCLCw       ---------- ---------- ---------- ---------- ---------- ---EHRQLTP
NCLCm       ---------- ---------- ---------- ---------- ---------- ---APREPSH
NCLCcr      ---------- ---------- ---------- ---------- ---GRGGAGG IGPGGRGGFN
NCLEa       ---------- ---------- ---------- ---------- ---------- ---TERAPSE
NCLEs       ---------- ---------- ---------- ---------- ---------- ---APRGPSE
NCLO        ---------- ---------- ---------- ---------- ---------- ---------P
NCLOs       ---------- ---------- ---------- ---------- ---------- ---SESPFGG
NCLMp       ---------- ---------- ---------- ---------- ---------- ---APRDPTQ
NCLPi       ---------- ---------- ---------- ---------- ---------- ---TPREASE
```

```
NCLDf        ---------- ---------- ---------- ---------- ---------- ---AKREPSE
NCLDc        ---------- ---------- ---------- ---------- ---------- ---EVRAPSQ
NCLPj        ---------- ---------- ---------- ---------- ------GGGG GGEGGGGGVG
NCLLda       ---------- ---------- ---------- ---------- ---------- ---------E
NCLCa        ---------- ---------- ---------- ---------- ---------- ---TPRQPSE
NCLCc        ---------- ---------- ---------- ---------- ---------- ---APRGPSE
NCLChd       ---------- ---------- ---------- ---------- ---------- ---AARETSE
NCLCmu       ---------- ---------- ---------- ---------- ---------G RGPGGRGGFN
NCLCn        ---------- ---------- ---------- ---------- ---------- ---APRQPSE
NCLC         ---------- ---------- ---------- ---------- ---------- ---APREKTE
NCLCt1       ---------- ---------- ---------- ---------- ---------- ---SPRGPSE
NCLCt2       ---------- ---------- ---------- ---------- ----GGGGKG TGPGGRGGFN
NCLDbGSO10   ---------- ---------- ---------- ---------- ---------- ---------A
NCLDbGSO10   ---------- ---------- ---------- ---------- ---------- ENQQQRASIK
NCLSco1      ---------- ---------- ---------- ---------- ---------- ---AXREPSQ
NCLSco2      ---------- ---------- ---------- ---------- ---GGRGGAG LGPGGVGGFN
NCLSd        ---------- ---------- ---------- ---------- ---------- ---AAREPSQ
NCLSj        ---------- ---------- ---------- ---------- ---------- ---AAREPSQ
NCLSma       ---------- ---------- ---------- ---------- ---------- ---AGREPSQ
NCLSme       ---------- ---------- ---------- ---------- ---------- ---AAREASQ
NCLTa        ---------- ---------- ---------- ---------- ---------- ---EARPVSQ
NCLTg        ---------- ---------- ---------- ---------- ---------- ---GPRPPSE
NCLTm        ---------- ---------- ---------- ---------- ---------- ---APRAPSQ
NCLTn        ---------- ---------- ---------- ---------- ---------- ---APRQPSE
NCLTp        ---------- ---------- ---------- ---------- ---------- ---EARAPSQ
NCLTo1       ---------- ---------- ---------- ---------- ---------- ---EARAPSQ
NCLTo2       ---------- ---------- ---------- ---------- ---RRSNEPG TGPGGYGAFN
NCLTw        ---------- ---------- ---------- ---------- ---------- ---APREPSK
NCLAg        ---------- ---------- ---------- ---------- ---------- ---AAREPSQ
NCLAf        ---------- ---------- ---------- ---------- ------TPSG GG--GRNGFN
NCLGo        ---------- ---------- ---------- ---------- ---------- ---GAYVPSE
NCLScom      ---------- ---------- ---------- ---------- ---------- ---SQREPSQ
NCLSr        ---------- ---------- ---------- ---------- ---------- ---SGREPTP
NCLTan       ---------- ---------- ---------- ---------- ---------- ---GAREPTE
NCLA1        ---------- ---------- ---------- ---------- ---------- ---TPREASV
NCLA2        ---------- ---------- ---------- ---------- ---------- -GHGTMDNQN
NCLAp        ---------- ---------- ---------- ---------- ---------- ---TPREASV
NCLAr1       ---------- ---------- ---------- ---------- ---------- ---SPREPTQ
NCLAr2       ---------- ---------- ---------- ---------- ---------- --TTPGGGSS
NCLE         ---------- ---------- ---------- ---------- ---------- ---TPRESSV
NCLFc1       ---------- ---------- ---------- ---------- ------GPPG RGPPGGSGFN
NCLFc2       ---------- ---------- ---------- ---------- ----AAPRKC LWEAKPKGAA
NCLFk        ---------- ---------- ---------- ---------- ---------- ---APRAASE
NCLFs1       ---------- ---------- ---------- ---------- ---------- ---SAREPTQ
NCLFs2       ---------- ---------- ---------- ---------- ----GKGGKG LESGGGSGFN
NCLFs3       ---------- ---------- ---------- ---------- ----GKGGKG LGNGGGSGFN
NCLPa        ---------- ---------- ---------- ---------- ---------- ---AAREVTQ
NCLPau       ---------- ---------- ---------- ---------- ---------- ---APRETTQ
NCLPd        ---------- ---------- ---------- ---------- ---------- ---AAREVSQ
NCLPf        ---------- ---------- ---------- ---------- ---------- ---AAREASQ
NCLPh        ---------- ---------- ---------- ---------- ---------- ---AARQVSE
NCLPp        ---------- ---------- ---------- ---------- ---------- ---APRDTAQ
NCLNpa       ---------- ---------- ---------- ---------- ------GGAG GSGGGGGGFN
NCLNp        ---------- ---------- ---------- ---------- ---------- ---GAREPTM
```

```
NCLNi1      ---------- ---------- ---------- ---------- ---------G LGPGGAGRFN
NCLNi2      ---------- ---------- ---------- ---------- ---------G LGPGGAGRFN
NCLPsp      ---------- ---------- ---------- ---------- ---------- ---FEREVSP
NCLTv       ---------- ---------- ---------- ---------- ---------- ---FNKGPSE
NCLTs       ---------- ---------- ---------- ---------- ---------- ---FNKGPSE
NCLTc       ---------- ---------- ---------- ---------- ---------- ---FS-GPSE
NCLNsa      ---------- ---------- ---------- ---------- ---------- -MGFARPPSE
NCLNg       ---------- ---------- ---------- ---------- ---------- -MGFARPPSE
NCLTmi      ---------- ---------- ---------- ---------- ---------- ----APDKKD
NCLCr       ---------- ---------- ---------- ---------- ---------- ---------E
NCLBs       ---------- ---------- ---------- ---------- ---------- ---RESGGSG
NCLMc       ---------- ---------- ---------- ---------- ---------- -KHEKRPVNP
NCLSl       ---------- ---------- ---------- ---------- ---------- RGEGQEGGDK
NCLCsu      ---------- ---------- ---------- ---------- ----RGGRGG GKNTGERDGS
NCLEv       ---------- ---------- ---------- ---------- ---------- ---------D
NCLPg       ---------- ---------- ---------- ---------- ---------- ---------D
NCLEt       ---------- ---------- ---------- ---------- ------AKGD GKGTSPRE-D
NCLEn       ---------- ---------- ---------- ---------- ------GKGD GKGTSPRE-D
NCLEm       ---------- ---------- ---------- ---------- ----GERRHE GKNGAPRE-D
NCLEb       ---------- ---------- ---------- ---------- ----GDRKHE GKNGAPRD-D
NCLEmi      ---------- ---------- ---------- ---------- ----GDRKHE GKNGAPRD-D
NCLEa       ---------- ---------- ---------- ---------- ----GDRRHE GKNGGPRE-D
NCLEp       ---------- ---------- ---------- ---------- ----GDRRQE GKNGPPRD-D
NCLPo1      ---------- ---------- ---------- ---------- --------KG KGGKGKGKGH
NCLPo2      ---------- ---------- ---------- ---------- --------KG KGGKGKGKGH
NCLPm       ---------- ---------- ---------- ---------- --------KG GKGKGKGKGH
NCLCc1      ---------- ---------- ---------- ---------- ---------- GAVSGRND--
NCLCc2      ---------- ---------- ---------- ---------- ---------- GSAPGRND--
NCLSmi      ---------- ---------- ---------- ---------- ---------- -SS-----AT
NCLPbr      ---------- ---------- ---------- ---------- ---------- ---KQRGQQP
NCLGt       ---------- ---------- ---------- ---------- ---------- -FTPNREPKP
NCLG        ---------- ---------- ---------- ---------- ---------- -FG---EQRP
NCLDlu      ---------- ---------- ---------- ---------- ---------- ----VTYSMT
NCLP        ---------- ---------- ---------- ---------- -----GKAVP GGPEGEQAAA

            ....|....| ....|....| ....|....| ....|....| ....|....| ....|....|
                 965        975        985        995       1005       1015

NCLHs       STWSGESKTL VLSNLS---- ---YSATEET LQEVFEKA-- --TFIKVPQN QN--GKSKGY
NCLAt1      GGDGGDEKKI FVKGFDASLS ---EDDIKNT LREHFSS-CG EIKNVSVPID RD-TGNSKGI
NCLAt2      KGEGSQSRTI YVRGFSSSLG ---EDEIKKE LRSHFSK-CG EVTRVHVPTD RE-TGASRGF
Nsr1pSc     PSEPSDT--L FLGNLS---- ---FNADRDA IFELFAK-HG EVVSVRIPTH PE-TEQPKGF
NCLAs       KEDGCIT--V FVGNLP---- ---WDIDEAS MKEFFSP-CG EISMVRFAED RE-TGDFKGF
NCLAsu      KEPGCTT--L FMGNLS---- ---WNVDEDS VREFFKG-CG EIAGVRFSED RE-TGEFKGF
NCLCd       KPPGCTT--V FVGNLS---- ---FSIDEPT LREAFAE-CG EISEVRFAED RE-TGEFKGF
NCLCp       KPEGCTT--V FMGNLS---- ---YDIDEPS VRALFGE-CG TITQIRFAED RE-TGEFRGF
NCLCh       KPEGCDT--V FVGNLS---- ---YDVDEPT VRDFFKD-CG AIQTIRFAED RE-TGEFRGF
NCLCw       KAEGCVS--I FMGNLS---- ---WNIDEDS VRNFFAD-CG EIKRINFSED RE-TGEFKGF
NCLCm       KEEGCTT--V FVGNLS---- ---FAIDEDT LREAFAS-CG EIASVRFAED RE-TGAFKGF
NCLCcr      PTNR-EEVKL YVGNLS---- ---FETTEEQ VRSLFEQYGP VTDCFLPTDR DS--GRVRGF
NCLEa       KPEGCLT--I FIGNLS---- ---WQIDEDS VREAFQE-CG EITTVRFAED KE-TGDFRGF
NCLEs       KQEGCVT--V FIGNLD---- ---WNIDEET IRATFGD-CG EITQVRFAED RE-SGQFKGF
NCLO        KPEGCTR--I FVGNLS---- ---FDIDEDT IKNFFAQ-CG TVTDVKWAED RD-TGRFRGF
NCLOs       KGEQMETNCI FIGNLN---- ---FDIDEQT VKDFFKE-CG AIREVRWGEE RE-TGKFRGF
```

```
NCLMp        KQEGCLT--V FIGNLA---- ---WDVDEDS IRQIFGD-CG TISQVRFATD RE-TGDFKGF
NCLPi        KQEGCTT--V FVGNLS---- ---WDIDEET LRQTFGE-CG TICSVRFATD RE-TGEFRGY
NCLDf        KQEGTLT--V FVGNLS---- ---FNIDEET LKEAFAD-CG EITQVRFAED KE-TGQFKGF
NCLDc        KDEGCMT--V FVGNLS---- ---FHIDEDT VRETFKD-CG EIASIRFAED RE-TGQFKGF
NCLPj        GGGG-DRTKL YVGNLS---- ---FYTTLDT IQGLFEEYGE VFDCYMPTDR ET--GASRGF
NCLLda       KPEGCTT--V FCGNLS---- ---FDIDEDS LREFFGE-CG NIREVRFATD RE-TGDFKGF
NCLCa        KPEGCTT--V FVGNLS---- ---WQIDEET LRAAFAD-CG EIAQVRFAED RE-TGDFKGF
NCLCc        KPEGCCT--V FCGNLS---- ---WNIDEDS LREAFQE-CG EISQVRFATD RE-TGDFKGF
NCLChd       KPAGCKT--V FCGNLS---- ---WNVDEDT LRETFAE-CG EISSIRFATD RE-TGEFKGF
NCLCmu       PTGA-ENVKL YVGNLS---- ---FDTTEEA VRGYFEKFGE VSDCFLPTDR NN--NNNN--
NCLCn        KPEGCTT--V FVGNLS---- ---FNIDEDT LREAFAE-CG EISQIRFATD RE-TGDFRGY
NCLC         KPEGCNT--V FVGNLS---- ---WNIDEET LRSSFXE-CG EISEIRFAED KE-TGDFKGY
NCLCt1       KPEGCNT--V FCGNLS---- ---WNVDEDS LRAVFQD-CG EITQIRFAED KE-TGEFKGF
NCLCt2       SSGA-ENVKL YVGNLS---- ---FDTTDEV VRGQFEKYGE VVDCFLPTDR ES--GKVRGF
NCLDbGSO10   KTPGSVT--V FVANLS---- ---FSIDEET VRDTFKD-CG TITAIRFGED RD-TGAFKGY
NCLDbGSO10   KPEGCRE--V HIRNLS---- ---SDVDEKI IRATFEKKCG TIDHVSFIKD KE-TKKFAGY
NCLSco1      KXEGCMT--V FVGXLS---- ---FHXDEET VRETFKD-XG EITAVRFAED KE-TGQFKGF
NCLSco2      PQGR-DEVKL YVGNLS---- ---FDTDEQQ VRDIFEKYGQ VSDCFLPQDR DS--GRPRGF
NCLSd        KEEGCMT--V FVGNLS---- ---FHIDEET VRETFKD-CG EITAVRFAED KE-TGQFKGF
NCLSj        KEEGCMT--V FVGNLS---- ---FHIDEET VRDTFKD-CG EITAVRFAED KE-TGQFKGF
NCLSma       KEEGCMT--V FVGNLS---- ---FHVDEET VRETFKD-CG EITAVRFAED KE-TGQFKGF
NCLSme       KEEGCMT--V FVGNLS---- ---FHIDEDT IRDTFKD-CG EITAVRFAED KE-TGQFKGF
NCLTa        KEEGCMT--V FVGNLS---- ---FNIEEDA LREAFQE-CG EITNIRFATD RE-TGDFKGF
NCLTg        KDPGCLT--V FVGNLS---- ---FNIDEDT VREAFKD-CG EISSIRFAED RE-TGQFKGF
NCLTm        KEDGCVT--V FVGNLS---- ---FNIDEET LRDAFKD-CG EIASVRFAED RE-TGQFKGF
NCLTn        KPPGCLT--V FVGNLS---- ---WEIDEES LRAAFAE-CG EITQVRFATD RE-TQEFKGF
NCLTp        KDEGCTT--V FVGNLS---- ---FHIDEDT VRETFKD-CG EIANIRFAED RE-TGQFKGF
NCLTo1       KEEGCVT--V FVGNMS---- ---FHIDEDT VRETFKD-CG EIASIRFAED RE-TGAFKGY
NCLTo2       PQGR-EDVKL YVGNLS---- ---FDTNEEA VRSMFEQYGT VSDCFLPSDR DT--GRPRGF
NCLTw        KDPGCVT--V FVGNLS---- ---FDIDEDA LRDAFKD-CG EICSIRFAED RE-TGAFKGF
NCLAg        KEEGCIT--V FIGNLS---- ---WDVDEDA IRQAFGD-CG EINQVRFATD RE-TGEFKGF
NCLAf        AAGA-AEVKL YVGNVA---- ---FDTDEET IRKLFEQYGK VSDCFMPTDR DS--GKVRGF
NCLGo        KTEGTTT--I FVGNMS---- ---FDIDEET LRGALEETCG EITSVRFAMD RD-TGNFKGF
NCLScom      KEEGCTT--V FIGNLP---- ---WSADEET IREVFGE-CG EIKMVRFSTD RE-TGDFKGY
NCLSr        KPPGCTT--V FVGNLS---- ---WSIDEDS LREAFAH-CG EIASVRFAMD RE-TQEFKGF
NCLTan       KPEGCLS--V FVGNLS---- ---WSIDEES LRAAFAE-CG EISQVRFATD RE-TQEFKGY
NCLA1        KEEGCCT--V FVGNLS---- ---FDIDEDT LRATFGE-CG TITQVRFATD KE-TEQFKGF
NCLA2        AATANPSRTV MLHNLS---- ---YQATEED IQAVLERVFS QQLHDNNNNN KKNKSLELSR
NCLAp        KEEGCCT--V FVGNLS---- ---FDIDEDT LRQVFGE-CG EITQVRFAMD KE-TEQFKGF
NCLAr1       KENGCVT--V FVGNLS---- ---YDIDEDT LKEAFQQ-CG EIKSVRFALD KE-TEAFKGY
NCLAr2       STAPSAT--L FMGGLS---- ---YQATEDD VWNAFSG-YA EPVSVRIATD RE-TGNPKGF
NCLE         KEEGCCT--V FVGNLS---- ---FDIDEET LKEAFGE-CG EITQVRFATD KE-TEQFKGF
NCLFc1       ASGS-TDVKL YVGNLA---- ---FDTTEDS VRQTFEQYGK VVDCFLPTDR ES--GKVRGF
NCLFc2       GTAS-STTKL YVGNLD---- ---FDTTAET LRDAFAEYGD VKDCTLPEDR DF--GGSRGF
NCLFk        KQEGCMT--V FVGNLS---- ---WDVDEDT LRDAFKD-CG TITQVRFSTD RE-TGDFKGY
NCLFs1       KQDGCVT--V FVGNLA---- ---WDVDEDT LRQAFGE-CG EITSVRFATD RE-TGEFKGF
NCLFs2       AAGK-PDVKL YVGNLA---- ---FETTQES IQSLFEQYGT VSDCFMPTDR DT--GKTRGF
NCLFs3       AAGK-PDVKL YVGNLA---- ---FETTQES IQSLFEQYGT VSDCFMPTDR DT--GKLRGF
NCLPa        KQEGCCT--V FVGNLS---- ---WDIDEET LRAAFAD-CG TITQVRFSTD RE-TGDFKGY
NCLPau       KQEGCCT--V FVGNLS---- ---WDIDEDS LRAAFAE-CG TISQIRFSTD RE-TGDFKGY
NCLPd        KQEGCCT--V FVGNLS---- ---WDIDEET LRAAFAD-CG TITQIRFSTD RE-TGDFKGY
NCLPf        KQEGCCT--V FVGNLS---- ---WDIDEDS LRAAFAD-CG TINQIRFSTD RE-TGDFKGY
NCLPh        KQEGCCT--V FVGNLS---- ---WDIDEDS LRAAFAD-CG TITQVRFSTD RE-TGDFKGY
NCLPp        KPEGCCT--V FVGNLS---- ---WDIDEES LRAAFAD-CG TIAQVRFSTD RE-TGDFKGY
```

```
NCLNpa      SAGK-EDVKL YVGNLS---- ---FDTQEQE IRRMFEEHGT VSDCFMPVDR DS--GKKRGF
NCLNp       KEEGCVT--V FVGNLS---- ---WDIDEDS LRDAFKD-CG TITQVRFSTD RE-TGDFKGY
NCLNi1      SAGN-EETRF PYIFFYVFIN ---SDTPETE IRRIFEEYGT VNDCFMPTDR DS--GKVRGF
NCLNi2      SAGN-EEVKL YVGNLSLYVN GLNSDTPEAE IRRIFEEYGT VNDCFMPTDR DS--GKVRGF
NCLPsp      KMDGCMT--V FMGNLS---- ---WDVDEDS IRETFGA-CG EITEVRFAMD RE-SGEFKGF
NCLTv       KMAGCMT--V FIGNLS---- ---WDVDEET IRSIFEG-CG TISQIRFAED RE-TGRFKGF
NCLTs       KMAGCMT--V FIGNLS---- ---WDVDEET IRSIFEG-CG TISQIRFAED RE-TGRFKGF
NCLTc       KQEGCMT--V FIGNLS---- ---WDVDEDA IRQTFES-CG EITQIRFAED RE-TGRFKGF
NCLNsa      KPEGCMS--V FIGNLS---- ---WNIDEET IRSAFAS-CG EVMSVRFATD RE-TGEFRGF
NCLNg       KPEGCMS--V FIGNLS---- ---WNIDEET IRSAFAS-CG EVTSVRFATD RE-TGEFRGF
NCLTmi      KPEGCTT--V FIGNLS---- ---WDVDEDA VRAAFGE-CG EIASVRFATD RE-TGDFKGF
NCLCr       EAEASAT--L FMGGLS---- ---YTATEDD VYNAFAG-VA EPVRVRIATD RD-SGEPKGF
NCLBs       R-GSTST--V FVGNLP---- ---YNADENN LTDFFSD-CG SIKQVRIAHD QD--GNKRGF
NCLMc       DAEGSCC--I FIGNVG---- ---YNTTKEA LWDFFAN-YG NITDLRV-AH DM-DGNPRGF
NCLSl       PEQK--STTV FVGNLS---- ---YTTTEKS LEKFFSGCGA IKAVRIAKME D---GKLKGF
NCLCsu      RPKAAPNKQI VVRNLS---- ---FNATEES IRGLFEE-CG DIQEVRMPVF ED-SGKFKGQ
NCLEv       KKTNEVGANL FIGNLEP--- ----EVDEKM LYDTFSAFGV LLFAKVMRDP DN--GLSRGF
NCLPg       KKTNEVGANL FIGNLEP--- ----EVDEKM LYDTFSAFGV LLFAKVMRDP DT--GASRGF
NCLEt       KIKTEPCNQI VLRNLS---- ---FDTTEET IRETFGD-CG EIRAVRIPVF ED-SGKPRGQ
NCLEn       KIKAEPCNQI VLRNLS---- ---FDTTEET IRETFGD-CG EIRAVRIPVF ED-SGKPRGQ
NCLEm       RPKADPSNQI VLRNLS---- ---FDTTEDT IRETFGE-CG EIRAVRIPVF ED-SGKPRGQ
NCLEb       RPKADPSNQI VLRNLS---- ---FDTTEDT LRETFGE-CG EIRAVRIPVF ED-SGKPRGQ
NCLEmi      RPKADPSNQI VLRNLS---- ---FDTTEDT IRETFGE-CG EIRAVRIPVF ED-SGKPRGQ
NCLEa       RPEADPSNQI VLRNLS---- ---FDTTEDT IRETFGD-CG EIRAVRIPVF ED-SGKPRGQ
NCLEp       RPKADPSNQI VLRNLS---- ---FDTTEET IRETFGD-CG EIRAVRIPVF ED-SGKPRGQ
NCLPo1      HELGEKSASV FVGNLP---- ---WSMTQDW LYEVFGD-CG SITRCFMPTD RE-TGNPRGF
NCLPo2      HELGEKSASV FVGNLP---- ---WSMTQDW LYEVFGD-CG SITRCFMPTD RE-TGNPRGF
NCLPm       HELGEKSASV FVGNLP---- ---WSMTQEW LSEVFGD-CG SITRCFMPTD RE-TGNPRGF
NCLCc1      --KAPTGRSV IIGNLS---- ---FGSTQES LRALFEE-CG NIDDIRIPVF ED-TGKPRGR
NCLCc2      --KAPTGRSV IIGNLS---- ---FGSTQES LRALFEE-CG PIDDVRIPVF ED-TGKPRGR
NCLSmi      KPEGCDT--V FVGNLP---- ---WDVTEDQ IYELFAP-CG EVARVRFATS QE-DGSFRGF
NCLPbr      KPDGCTT--V FIGNLS---- ---FQADEYA ISQVFED-CG EIVSVRICTD RDT-GKSRGF
NCLGt       KPDGCTT--I FMGNLS---- ---WDVDEDT IRSFFAD-CG EVVNVRFATD RE-TGDFKGF
NCLG        KPAGCVS--L FMGNLS---- ---WEVTEEE VRGLFAD-CG EVTQVRFATD RE-TGEFKGF
NCLDlu      VPPGVTK--L FVGNLS---- ---FKVTEAA LRDAFAP-CG HVTAVRLGMD KEDPTKFAGW
NCLP        PEYD-RKRSV FVGNMP---- ---FDAQEQS LREHFASCGN VESVRIVRDP YE--NMGKGF

            ....|....| ....|....| ....|....| ....|....| ....|....| ....|....|
                1025       1035       1045       1055       1065       1075

NCLHs       AFIEFASFED AKEALNSCNK REIEGRA-IR LELQGPRGSP NARSQPSKTL FVKG--LSED
NCLAt1      AYLEFSEGKE KALELN---G SDMGGGFYLV VDEPRPRGDS SGGGGFGRGN G---------
NCLAt2      AYIDLTSGFD EALQLS---G SEIGGGN-IH VEESRPR-DS DEGRSSNRAP A---------
Nsr1pSc     GYVQFSNMED AKKALDALQG EYIDNRP-VR LDFSSPRPNN DGGRGGSRG- FGG-RGGGR-
NCLAs       GHVEFVDTDS TDKAVA-LAG QDLMGRA-VR VDFANDRRNA PGGGG--RGG G---------
NCLAsu      GHVDFVNTES CDKGVL-LSG QELMGRA-IR LDFANPRPSG GGGAGGRGGG RG--------
NCLCd       GHIEFVDSAS TDEALK-LVG TEIMGRG-VR IDYAADKRGE GGG----GGR G---------
NCLCp       GHVEFAESEA TDAAVA-LAG SDLGGR---- ---------- ---------- ----------
NCLCh       GHVQFCDTMS TDEAVK-LNG QAMCGRN-AR IDYA-PDKRG GG-------- ----------
NCLCw       GHVDFYDTES TDKAVA-LAG QDLCGRP-IR VDYSNPREFG GG-----GGG RG--------
NCLCm       GHIEFVDTES TDLAVK-MAG EDVMGRA-IR VDYANSRNHG EGRG---GGG ----------
NCLCcr      AFVTMP-AKE AEVACTKANG ADLDGRS-LR VNEAQPKGFG GGG---GGGA GG--------
NCLEa       GHVEFANTDS TTQAVA-MAG TDIMGRA-VR VDYAQDRYSG GGGGGGGRGG G---------
NCLEs       GHIEFAETEA TDKAVA-LAG TDMLGRA-VR VDYANDRRGG GGGGGGFGGG RGGG------
```

```
NCLO        GHLEFEESDA TDKAVA-MAG SDLLGRP-IK IDYAKAQARK SF-------- ----------
NCLOs       GFVEFEESSS TDKALE-LNS VDVMGRP-IK IDKT-GPSNG GG-------- ----------
NCLMp       GHIEFEETEA TDAAVK-LAG TDICGRG-VR VDFANDKRQG GGF----GGG G---------
NCLPi       GHVEFEETEA TDKAVG-MAG QDVMGRA-IR VDFAAVRQN- SFGGGGGRGG G---------
NCLDf       GHIEFAATES TDAAVK-LAG TDIMGRA-VR VDYANDRRSN GGGR-GFGGG G---------
NCLDc       GHVEFVETES TDKAVA-MAG TYVMDRA-LR VDFANERKSF GGGG---GGG ----------
NCLPj       AFVTMS-PEA ANAAMDECDG YDLEGRI-IR VNEAQPNGGR GGGRGGGGQG GGR-------
NCLLda      GHVDFENTEA VDAAVK-LNG EYVANRR-IR IDYAADRKNG GG-------- ----------
NCLCa       GHVEFVNSES TEAAVK-MAG TDIMGRP-CR VDYAADKRKQ NGG---FGGG ----------
NCLCc       GHIEFVETES ADNAIA-MAG TDILGRA-VR VDYAADRRGG GGG---FGGG G---------
NCLChd      GHIEFFSSDS TDAAIA-MAG TDIMGRA-VR VDYAADKR-- AGGG--FGGG R---------
NCLCmu      -NNNNN-NNN NNNNNNNNNN NNNNNNN-NN NNNNQPKGMG GVG-----GP RG--------
NCLCn       GHMEFVNSES TDLAVQ-MAG TDIMGRA-CR VDFAADKR-A GGG---FGGG ----------
NCLC        CHIEFVNSES TDKAIE-MAG TEVLGRP-LR VDFAADKKK- -----QFGGG G---------
NCLCt1      GHIEFAETEA TDAAVA-LAG TDIMGRA-VR VDYAGDKRKQ AGGGRGFGGR G---------
NCLCt2      AFVTMK-AAD AEKACMAMNG CEVDGRT-LK VNEAQPK--- ----GMGGGR GG--------
NCLDbGSO10  AFMDFEETEA TDKAVA-MAG TEIMGRA-VR VDHTTGASNT PGRT------ ----------
NCLDbGSO10  GYIRFKKTES TDKAVK-MTG VQILGKS-AR INYALASSEK SDNDAKQKPL SKKPEGCKDV
NCLSco1     GHVEFAETES TDLAVA-LAG TYVMDRP-LR VDFANERKDR GFGG---GGG ----------
NCLSco2     AFVTMP-AKE AEEACDKVNG LEVDGRT-LR VNEAQPKGSG GGGR--GGGF GG--------
NCLSd       GHVEFADTES TDAAVA-LAG TYVMDRP-LR VDFANERKDR GFGG---GG- ----------
NCLSj       GHVEFAETES TDAAVA-LAG TYVMDRP-LR VDFANERRDR GFGG---GG- ----------
NCLSma      GHVEFAETES TDLAVA-LAG TYVMDRP-LR VDFANERKDR GFGG---GGG ----------
NCLSme      GHVEFADTES TDKAVA-LAG TYVMDRP-LR VDFANERRDR GFGG---GG- ----------
NCLTa       GHVEFAETES TDKAVL-MAG TYVMDRA-LR VDYANDRRGG AGGG---GGG ----------
NCLTg       GHVEFVETES TDKAVA-MAG EYVMDRP-IR VDYANERKRT FGGE---GGG ----------
NCLTm       GHVEFVESES TDKAVE-LAG TYVMDRP-LR VDYANDRKKF GGGG---GGG GFGG------
NCLTn       GHIEFAETEA TDKAIA-MAG TDIVGRQ-VR VDFAADRRQS FG-----GGG G---------
NCLTp       GHVEFTETEA TDKAVA-LAG TYVMDRA-IR VDFANERK-S FGG----GGG ----------
NCLTo1      GHVEFVETEA TDKAVA-LAG TYVMDRP-IR VDYANERRGG AVSV---EVA ----------
NCLTo2      AFVTMP-AKE AETACNKVNG MELDGRT-VR VNEAQPKVSS SGG---GGNP PTPLI-FNLH
NCLTw       GHIEFADTES TDKAVE-MAG TEIMGRA-VR VDYANERX-- ---------- ----------
NCLAg       GHIEFCATES TDAAVK-MAG TEIMGRA-VR VDYANDRRNN AG-----GGG G---------
NCLAf       AFVTMP-SKD AEEACNKLNG YEVDGRA-MR VNEAQPKGGD SGGG---GGR GG--------
NCLGo       GHVEFATTES TDIAVTKMQG VEVMGRA-LR VDYAKDRKSF GGGG---GRF G---------
NCLScom     GHVEFYDTEA TDKAVE-LAG TDIMGRA-VR VDFANERRQS FG------GG G---------
NCLSr       GHVEFVESDS CDEAMK-MIG TDIAGRA-VR VDYAAD-RKP PGGG--FGGG G---------
NCLTan      GHIEFTETAA TDAAMQ-LAG TDICGRQ-VR VDYAADKRKS FGGA--GGGG G---------
NCLA1       GHIEFAETEA TDKAVG-MAG TDIMGRP-VR VDFANDRRKS GGFG---GGG ----------
NCLA2       GGIRVVRHSA TGHSKGFAYV EFVQQDDAVA LVQHCQQKNS SSGGLVVHGR ACR-------
NCLAp       GHIEFAETDA TDKAVA-MAG TDVMGRA-VR VDFANDRR-- GGGG---GGG ----------
NCLAr1      GHIEFYETEA TDKAVE-MAG TEILGRQ-VR VDYAVDRRRS LGGG---PGR ----------
NCLAr2      GHVQFSSVED ATSVMENVSS ISIAGRS-VR LDFAPERDNP P--------- ----------
NCLE        GHIEFSESSA TDKAVL-MAG TDICGRS-VR VDFANDRRKS GGFG---GGG ----------
NCLFc1      AFVTMP-AAD AEQACNKLNG YEMDGRA-LR VNEAQPKG-S SGGG---GGR DG--------
NCLFc2      GFVVMD-AED AKVALAELDG IELDGRV-IR VTEADGRKGS RNTDSNSGGA GR--------
NCLFk       GHIEFGETEA TDEALK-LIG TEILGRA-VR VDYANDRRQS FGGE---GGG RGRGGGRGGG
NCLFs1      GHIEFTDTEA TDKAIA-MAG TDILGRQ-VR VDYANDKKAG GGGG---GFG ----------
NCLFs2      CFVTMP-SAA AEDACNKLTG YELDGRA-LR VNEAQPK--G SERGGPGGGR GGG-------
NCLFs3      CFVTMP-AAA AEEACNKLSG YELDGRA-LR VNEAQPKGMG SERGPGGGGR GGG-------
NCLPa       GHIEFAETEA TDAAVK-LAG TDIIGRA-VR VDFANDRRQS FGGG---GG- ----------
NCLPau      GHIEFTETDA TDLAVK-MAG TDILGRA-VR VDFANDRRQS SGG----FGG G---------
NCLPd       GHIEFAETEA TDAAVK-IAG TDICGRP-VR VDFANDRRQS FGSG---GGG RGGFG----G
NCLPf       GHIEFAETEA TDAAVK-LAG TDILGRA-VR VDFANDADNP LAVA---EAG VVAT-----E
```

```
NCLPh         GHIEFAESHS TDAAIK-LAG TDILGRA-VR VDFANDRRQS FGGG---RQS FGG------G
NCLPp         GHIEFAETEA TDAAVK-MAG TDILGRA-VR VDFAQDKRQS FGGG---FGG GRG-------
NCLNpa        CFVTMP-ASD AEVACTKMNG VEVDGRP-LR VNEAQPRGGG GGGGG--GGG GG--------
NCLNp         GHIEFAESSA TDAAVK-LAG TEILGRP-VR VDYANDRKPS GGGG---FG- ----------
NCLNi1        CFVTMP-SAD AEKACNGVNG REIDGRA-LR VNEAQPRGGG GGGGGGGG-R GG--------
NCLNi2        CFVTMP-SAD AEKACNGVNG REIDGRA-LR VNEAQPRGGG GGGGGGGGGR GG--------
NCLPsp        GHIEFSATEA TDEAVK-MAG TMVAGRA-IR VDYAAVKQKR EFGAGGGGGR G---------
NCLTv         GHVEFEETEA TDAAVA-LAG QDVCGRP-IR VDFAAVKERK SFGDRSPQGG R---------
NCLTs         GHVEFEETEA TDAAVA-LAG QDVCGRP-IR VDFAAVKERK SFGDRSPQGG R---------
NCLTc         GHVEFADTMS TDKAVA-MAG TDVVGRP-IR VDFAAVKQG- -WSPR--QGG S---------
NCLNsa        GHVDFADESA PEAAVQ-MAG TPVMGRE-IR VDYAQPRPPR EGGF---GGG G---------
NCLNg         GHVDFADETA PEAAVQ-MAG TPVMGRE-IR VDYAQPRPPR EGGF---GGG G---------
NCLTmi        GHVEFVDGEG VDAAID-MAG TEIAGRP-VR VDYAAPRAPR ESFG---GGG G---------
NCLCr         GHAEFADVET ARAAKNALFG VEIAGRK-VR LDFAPPRNNS PGGGRGGRGG GRGGGFGGRG
NCLBs         AHVEFESTDS ADAAIK-LNG KELEGRE-LK IDFAGEKRSG G--------- ----------
NCLMc         AHCEFSTPEE AKASL-AAHG ERVEGRA-LR IDLSAPRANR GGDRGGNRGG FGGDRNGGNR
NCLSl         AHVEFE-EAE STQQAVALNG KDLDGRD-VK VDISEKLKNK RAEGGSWEDR RG--------
NCLCsu        AFIEFDSVES ATKAVE-YNN TEVDGRT-VW IDFALPRNSA GGRGG--GRG G---------
NCLEv         GFISYDTFEA SDAALAGMNG QFLCNRP-IS VSYAYKKETK GERHGSAAER LI--------
NCLPg         GFISYDSFEG SDAALAGMNG QFLCNRP-IS ASYAYKKETK GERHGSAAER LI--------
NCLEt         AFLEFDSVDS ATKALE-YNN TELDGRT-IW VAYALPRGAG GDRGGRGGRG G---------
NCLEn         AFLEFDSVES ATKALE-YNN TELDGRT-IW VAYALPRGSG GDRGGRGGRG G---------
NCLEm         AFLEFDSVDS ATKALE-FNN TELDGRT-IW VAYALPRGAG GDRGGRGGRG G---------
NCLEb         AFLEFDSVDS ATKALE-FNN TELDGRT-IW VAYALPRGAV GDRGGRGGRG G---------
NCLEmi        AFLEFDSVDS ATKALE-FNN TELDGRT-IW VAYALPRGAG GDRGGRGGRG G---------
NCLEa         AFLEFDSVDS ATKALK-FNN TELDGRT-IW VAYALPRGAG GDRGGRGGRG G---------
NCLEp         AFLEFDSVDS ATKALE-FNN TELDGRT-IW VAYALPRGAG GDRGGRGGRG G---------
NCLPo1        AYIDFDSEDS AEKATK-LSG TDLEGRQ-IR VNYNQPRESS GKGGKGKGKG ----------
NCLPo2        AYIDFDSEDS AEKATK-LSG TDLEGRQ-IR VNYNQPRESS GKGGKGKGKG ----------
NCLPm         AYIDFDTEDS AEKATK-LSG TDLEGRQ-IR VNYNQPRESS GKGGKGKGKG K---------
NCLCc1        CFIDFETEEG AKKALE-YNE TDIDGRT-VW IDYVRPREDK PSNGFRGGRG G---------
NCLCc2        CFIDFESEEG AKKALE-YNE TDIDGRT-VW IDYVRPREDK PSNGFRGGRG G---------
NCLSmi        GHVQFTSGES TDAAVA-MYG TELNGRA-IR VDYAPPRNRD ----SLGGG- ----------
NCLPbr        GYVEFTSTDA VDAAMK-KAG TDLAGRQ-VR VDFAAARGEG AGDR------ ----------
NCLGt         GHVQFAESSA TDLAVA-KGG EFVAGRA-IR VDFAEDRKPP GSAGSAGGG- ----------
NCLG          GHVQFSEEEA TEKGIA-KAG EFVAGRA-IR LDYAEDKKTQ GGAGGAGGR- ----------
NCLDlu        AHVEFASHAD AQKGVA-LNG LDLLGRS-IR IDPANPSAGG GGRG------ ----------
NCLP          GYVLFEERVS VERAIG-LHD SEFAKRK-LR VFRCLKSGEQ PGDKGGRGGS RG--------

              ....|....| ....|....| ....|....| ....|....| ....|....| ....|....|
                  1085       1095       1105       1115       1125       1135

NCLHs         TTEETLKESF DGSVRARIVT D--------- -RETGSSKGF GFVDFNSEED AKAAKEAMED
NCLAt1        -RFGSGGGRG RD-------- ---------- ----GGRG-R FGSGGG---- ----------
NCLAt2        -RGAPRGRHS DR-------- ---------- ----APRGGR FSDRAP---- ----------
Nsr1pSc       ---GGNRG-- ---------- ---------- ---FGGRG-- GARGG----- ----------
NCLAs         -RDGGRGGGR GG---GRGGG ---------- ------GGFG RGRGDG---- ----------
NCLAsu        -RGGDRF--- ---------- ---------- ----GGDRGG GRGGRG---- ----------
NCLCd         -RGGGRG-RG GGRGRSDGGF ---------- ---GRGRGGG RGGGGG---- ----------
NCLCp         ---------- ---------- ---------- ---------- ---------- ----------
NCLCh         --GRG----- ---------- ---------- ---------- ---------- ----------
NCLCw         -RGGGR---- ---------- ---------- ----GGGRGG GRFGGR---- ----------
NCLCm         -RGGGRGR-- ---------- ---------- --------GY MGGGGG---- ----------
NCLCcr        -GGGGYGGGR GGGGGYGGYN D--------- -GYAGGRGGY GGGG-GGYGG GG----YGA-
```

```
NCLEa        -RGGGRGGGG GGFGRGRGGD ---------- ------RGGG RGGGGG---- ----------
NCLEs        -RGGGRGGRG YMS------- ---------- ----GGGGGY GRGGGG---- ----------
NCLO         --GGGTDRPF AVGEKPEGCT TVFVGNLAFS VDEGSVIDAF KDCGEI---- ----------
NCLOs        --GGGGGRKF EVSPKPEGCT TVFVANLPFS CDDDALRGAL DSCGTI---- ----------
NCLMp        -RGGGRGGGG GRGYMGGGGG GGG------- ---FGGRGGG RGFGGR---- ----------
NCLPi        -RGGGRGGG- ---------- ---------- ------RGGG RGGGRG---- ----------
NCLDf        -RGRGRG-FG GGRNGG---- ---------- -----GFGGG RNGGG----- ----------
NCLDc        -RGGGRGGG- ---------- ---------- ----GGRGGY MGGGGG---- ----------
NCLPj        -FQGGRGQGR GGGGRFGGR- ---------- --GDQNQGGG GGGGSSGTSR G-------GG
NCLLda       --GGGRGGR- ---------- ---------- ---GGGRGYM RDGGRG---- ----------
NCLCa        ---RGGG-RG GGRG------ ---------- ------YGGG RGGGSRGF-- ----------
NCLCc        -RGRGGG-FG GGRGR----- ---------- ------GGGG RGYG------ ----------
NCLChd       -GGGRGGGRG GG-------- ---------- -------RGG GRGG------ ----------
NCLCmu       -GGGGYVGGY GGG------- ---------- -RGGYDRGGY GGGRGGGYDA G-----YDRG
NCLCn        ---RGGGGRG GGRGG----- ---------- ------YGGG RGGG------ ----------
NCLC         -GRGFGGGRG GGE------- ---------- -------RGG RGGG------ ----------
NCLCt1       -GGGGFGGRG GGGGRG---- ---------- -----YMRGG GGGG------ ----------
NCLCt2       --YNDRGG-- GYGGGYGGG- ---------- --YGGNDYGY GGGGGRGGYG G------GRG
NCLDbGSO10   -PGRTPGRSF TPSEKEEGCK QVFIGNLSFN VDEDTIRGAF KDCGTI---- ----------
NCLDbGSO10   YIGGLSSSIT QEDVRDLFKD CG----EIEE VILSMDNETV RCKGFGHIRF KETES-----
NCLSco1      -RGGGRGGR- ---------- ---------- ----GGRGGY MGGGGG---- ----------
NCLSco2      RGGGGYGYGG GGG----GYG ---------- -GRGGGRDNY RGGGGGGYGD NS----RGGG
NCLSd        -RGGGRGGR- ---------- ---------- ----GGRGGY MGGGGG---- ----------
NCLSj        -RGGGX---- ---------- ---------- ---------- ---------- ----------
NCLSma       -RGGGRGGR- ---------- ---------- ----GGRGGY MGGGGG---- ----------
NCLSme       -RGGGRGGR- ---------- ---------- ----GGRGGY MGGGGG---- ----------
NCLTa        -GGRGGGRG- ---------- ---------- ----GGRGGY MGGGGGRD-- ----------
NCLTg        -RG------- ---------- ---------- ---------- ---------- ----------
NCLTm        -GGRGRGGG- ---------- ---------- ----RGRGGY MGGGGG---- ----------
NCLTn        -RGGGRGGRS PGGFG-GGGR ---------- ---GRGRGGG RDGGR----- ----------
NCLTp        -RGGFGGR-- ---------- ---------- ---------- ---------- ----------
NCLTo1       -LEGAEVAV- ---------- ---------- ----VADEAV VDEAGAGT-- ----------
NCLTo2       RGGGGYGGGY GGG--YEPTD D--------- -GRGGG-GGW GGGGRGGSGG GG----YGG-
NCLTw        ---------- ---------- ---------- ---------- ---------- ----------
NCLAg        -RGGGGGGRG GRGYMGGGGG ---------- ------RGGG RG-GGR---- ----------
NCLAf        -GGGGRGGYD GGGGGYSGG- ---------- --GGGYSGGY DDRGGGG--- --------YG
NCLGo        -GGGGRGGX- ---------- ---------- ---------- ---------- ----------
NCLScom      -RGGGRGGRG GR-----GGG ---------- ------RGGG RG--GR---- ----------
NCLSr        -RGGGRGG-- ---------- ---------- ---GRGRGG- -YGGAS---- ----------
NCLTan       -RGGGRGGRS PGGRSPGRGY ---------- ---GRGRGGD RGGGGG---- ----------
NCLA1        -GFGGGVVDS AVVVDSAVEV REAVVVEAEA TWAEAEAEVV GDEAAAEEEV AVDVV-VAVV
NCLA2        -IDYDHGRVR ---------- ---------- -------GSF RTADRT---- ----------
NCLAp        -GFGGGGRG- ---------- ---------- ----GGGGGF GGGGRG---- ----------
NCLAr1       -SGDGRGDG- ---------- ---------- ----RGGGRF GGRGRGAG-- ----------
NCLAr2       ---------- ---------- ---------- ---------- ---------- ----------
NCLE         -GGGGRGGG- ---------- ---------- ----GSFGGR GGGGRGGD-- ----------
NCLFc1       -GGGGYGG-- GGGGGYGGG- ---------- --GGGGRGGY DDRGSGGWKP GTWSRTSCYG
NCLFc2       --LAGGGGGR GGG------- ---------- --GGGRLGAA GGRAVGGFGG GP--------
NCLFk        GRGGFGGGGR F--------- ---------- ---GSGDRGR GGGGGGRSYG S---------
NCLFs1       ---GGGGRS- ---------- ---------- ----NDRGGR GGGRGR---- ----------
NCLFs2       --GYDRGGYD SYGGGYGGG- ---------- --GGG---GY GGGYDNDRGG G------RGG
NCLFs3       --GYDRGGYD SYGGGYSGG- ---------- --GGG---GY GGGYDNDRGG G------RGG
NCLPa        -RGGGRGGSR F--------- ---------- ---GSGDRGR GGGRGGFGGR ----------
NCLPau       GRG---GGGR F--------- ---------- ---GSGDRGR GGG-GRFG-- ----------
```

```
NCLPd        GRGGGRGGSR F--------- ---------- ---GSGDRGR GGGRGGFG-- ----------
NCLPf        VAGVAAVEAE V--------- ---------- ---ASEAATE AAAVADEVAT ----------
NCLPh        GRGGGRGGGR F--------- ---------- ---GSADRGR GGGRGGYG-- ----------
NCLPp        GRGRGRGGGR F--------- ---------- ---GSGDRGR GGG-GR---- ----------
NCLNpa       -RGGGYSES- YSGGGYSGG- ---------- --GGGGYGGS SG-GGGGYGG G-----RGYG
NCLNp        -RGGGRGGGR G--------- ---------- ---GRGGGGR GRGGGR---- ----------
NCLNi1       -GGGGYGDG- YGGGGYGGGS ---------- --GSNSYGGY NDSGYGGGGG G-----SGYR
NCLNi2       -GGGGYGDG- YGGG-YGG-- ---------- --GTNSYGGY NDSGYGGGGG G-----GGYR
NCLPsp       -GGGGRGGG- ---------- ---------- ------RGGG -GGGFG---- ----------
NCLTv        -GGGGRGGF- ---------- ---------- ------GGGG RGGGGR---- ----------
NCLTs        -GGGGRGGF- ---------- ---------- ------GGGG RGGGGR---- ----------
NCLTc        -PKGGRGDY- ---------- ---------- ------GGKG DYGGKG---- ----------
NCLNsa       -RGGGRGGG- ---------- ---------- ------RGGG RSFGGGEGAG ----------
NCLNg        -RGGGRGGG- ---------- ---------- ------RGGG RSFGGG---- ----------
NCLTmi       -GGGGRGGG- ---------- ---------- ------RGGG R--------- ----------
NCLCr        GGRGGFGGR- ---------- ---------- ---GGGRGGF GGRGGGFGGR ----------
NCLBs        ---------- ---------- ---------- -----SSRGG RGRG------ ----------
NCLMc        GGFGGDRGG- ---------- ---------- ---FGGRGGF GGRGGF---- ----------
NCLSl        -GRGGRGGFR GGD------- ---------- --RGGFRGGF RGGDRGGFRG GS--------
NCLCsu       -RGAFGGR-- ---------- ---------- ---------- --GGRG---- ----------
NCLEv        --AANRPKEG PT-PAAAPGR ---------- -------SGA PAMRPPMMPP G---------
NCLPg        --AANRPSDL PKGPGAAPAK G--------- -GGKGGPPGM PSMRPPMLPP G---------
NCLEt        -RGGAGGGGR ---------- ---------- -------GGG RGGGRGGF-- ----------
NCLEn        -RGGAGGGGR ---------- ---------- -------GGG RGGGRGGF-- ----------
NCLEm        -RGGAAGGRG ---------- ---------- -------GG- RGGGRGGF-- ----------
NCLEb        -RGGAAGGRG ---------- ---------- -------GG- RGGGRGGF-- ----------
NCLEmi       -RGGAAGGRG ---------- ---------- -------GG- RGGGRGGF-- ----------
NCLEa        -RGGAGGGRG ---------- ---------- -------GG- RGGGRGGF-- ----------
NCLEp        -RGGASGGRG ---------- ---------- -------GGG RGGGRGGF-- ----------
NCLPo1       ---------- ---------- ---------- -------GKG KGGKGKGK-- ----------
NCLPo2       ---------- ---------- ---------- -------GKG KGGKGKGK-- ----------
NCLPm        ---------- ---------- ---------- -------GKG KG-KGKGK-- ----------
NCLCc1       -RGGFGGRGG GFG------- ---------- ------GGGF GGRGGGGG-- ----------
NCLCc2       -RGGFGGRGG ---------- ---------- ------GGGF GGRGGGGG-- ----------
NCLSmi       -GRGGRGGG- ---------- ---------- ------LGGG RGRGDM---- ----------
NCLPbr       ---------- ---------- ---------- -----APRGG RGGGRG---- ----------
NCLGt        -GGGGRGGGR G--------- ---------- -----GFGGG RGGGG----- ----------
NCLG         -GSFGSAGG- ---------- ---------- -----GRGGG RGGGAA---- ----------
NCLDlu       --GGGRDGG- ---------- ---------- -----RGGGG RGGTPG---- ----------
NCLP         -GGRGGAGGR GGG------- ---------- --RGGSRDGG GGMGGGGGRG G---------

             ....|....| ....|....| ....|....| ....|....| ....|....| ....|....|
                 1145       1155       1165       1175       1185       1195

NCLHs        GEIDGNKVTL DWAKPKGEGG FGGRGGGRGG FGGRGGG--- -RGGRGGFGG RGRGGFGGRG
NCLAt1       ---------- ---------- -RGRDGGRGR FGSGGGR--- ------GSDR GRG--RPSFT
NCLAt2       ---------- ---------- -RGRHSDRG- --APRGR--- ------FSTR GRGPSKPSVM
Nsr1pSc      ---------- ---------- ---------- ---RGGFR-- ---PSGSGAN TAPLGRSRNT
NCLAs        ---------- ---------- GFGRGRGGGG G--------- ----GRGAPS PFAAKKNGAI
NCLAsu       ---------- ---------- GGGRGGG--- -RGGGRGG-- ---GRGDNSR TFSMKKTGGI
NCLCd        ---------- ---------F GRGRGG---- GRD------- ----SGGGGN Y-NPKKTGGI
NCLCp        ---------- ---------- ---------- ---------- ---------- ----------
NCLCh        ---------- ---------- ---------- ---------- ---------- ----------
NCLCw        ---------- ---------- GGGFGGG--- -RGGGRGG-- ---GRG---- -------GGX
```

| | | | | | | |
|---|---|---|---|---|---|---|
| NCLCm | ---------- | ------RGGR | GGR--GGRGG | GRGGGR---- | ----GRGGFD | PAKAKRSGGI |
| NCLCcr | GYDRGGYSDR | GG--YGGHG- | --GGGGYDRG | GYDRGHG--- | GGYSGGGGGY | DDGYGAGGAG |
| NCLEa | ---------- | ---------- | GFGRGGGRGG | GRGGDFG--- | ---RGRGGAS | SSGSKQHGSI |
| NCLEs | ---------- | ---------- | GYGRGGGGGY | GRGGGGRG-- | ---GGGRGKS | PGFAKRTGGI |
| NCLO | ---------- | -------AAT | RFITDRETGD | FKGMGFVE-- | ---FADTESA | DKAVKMGQEG |
| NCLOs | ---------- | -------VDV | RLMNDRETGR | FKGYAHVE-- | ---FEGTECT | DKAVELN--G |
| NCLMp | ---------- | ---------- | GGGRGGGRGG | GRG------- | ----G-GGGG | GFNAKRTGGI |
| NCLPi | ---------- | ---------- | GGGRSGGGGG | YGGGGRS--- | ---GGRGGAD | PARSRNNGSI |
| NCLDf | ---------- | ---------F | GRGRGG---- | GRG------- | -----GGRGN | MTSAKKSGGI |
| NCLDc | ---------- | --GGGFGGGR | GGGRGGGRGG | GRGGGR---- | ----GGGPTS | TFSAKKSGGI |
| NCLPj | GSSSGSNSAS | SGSASNTGGR | ESASSGTDGA | ASRYRSS--- | ------GFAI | VDGQSEG--- |
| NCLLda | ---------- | -------RGG | R---GRGRGR | GRGNG----- | ---------- | ---------- |
| NCLCa | ---------- | ---------G | GGGRGGSRGY | GGG------- | ----GGGRDS | YSSAKKKGGI |
| NCLCc | ---------- | ---------- | GRGRGG---G | GRG------- | ----RGGGGG | FASAKKNGGI |
| NCLChd | ---------- | ---------- | GRGRGGF--G | GGG------- | ----RGGGGG | YEKKSFG--- |
| NCLCmu | GYDTG--YDR | GGYDAGVIG- | --GYGG-DRG | GYGAP----- | ----RGGQGY | ND---GGYGG |
| NCLCn | ---------- | ---------Y | GRGRGG--GY | GRG------- | ----RGGGRG | EGSAR-TGGI |
| NCLC | ---------- | ---------- | GRGGGGR--G | GG-------- | ----RGRGQG | FAKKSGG--I |
| NCLCt1 | ---------- | ---------- | GRG-GGR--G | GGG------- | ----RGGG-- | FSKPSFAKKS |
| NCLCt2 | GYDRGGYGGG | NDGGYNDRGY | GGGGGGGGYG | GGNRGGGYNS | YNDRSGGGGY | GGGQGGG--- |
| NCLDbGSO10 | ---------- | -------TSI | RFGEDRETGE | FKGFGHIE-- | ---FEETEAT | DKAVAMAG-- |
| NCLDbGSO10 | ---------T | DKAVEIRGTF | FMGRFIKVDY | SASKKKKGKN | SSSAKGGGKS | KSKTDKGAEA |
| NCLSco1 | ---------- | --GGRGRGGF | GGGRGGGRGG | GRGGGR---- | ----DFG--- | AGGNKRSGGI |
| NCLSco2 | GYG----EDR | GGYGGGGGG- | --GGGSGGGG | GYDQG----- | ----GYGGGR | GDYDRSGGGG |
| NCLSd | ---------- | --GGRGRGGF | GGGRGGGRGG | GRGGGR---- | ----DFG--- | AGGNKRSGGI |
| NCLSj | ---------- | ---------- | ---------- | ---------- | ---------- | ---------- |
| NCLSma | ---------- | --GGRGRGGF | GGGRGGGRGG | GRGGGR---- | ----DFG--- | AGGNKRSGGI |
| NCLSme | ---------- | --G-RGRGGR | GGGRGGGFGG | GRGGGR---- | ----GGN--- | SFGAKKSGSI |
| NCLTa | ---------- | --GGRGRGGR | GGGRGGGRGG | -RDGGR---- | ----GGG-RG | MMGAKR--GI |
| NCLTg | ---------- | ---------- | ---------- | ---------- | ---------- | ---------- |
| NCLTm | ---------- | --GR---GGR | GGR--GGRGG | GRGGGR---- | ----GGG--- | -FGAKKTGGI |
| NCLTn | ---------- | ---------- | --GRGGGRDG | GRG------- | ------RGGN | SFGAKKHGSI |
| NCLTp | ---------- | ---------- | ---------- | ---------- | ---------- | ---------- |
| NCLTo1 | ---------- | --CQVAVDAV | AE-------- | ---------- | ---------- | ---------- |
| NCLTo2 | GRG-GGYDDR | GGGGYGDRG- | --GGGGGGGG | YNDRG----- | ----GGGGGY | NDRGGGARPG |
| NCLTw | ---------- | ---------- | ---------- | ---------- | ---------- | ---------- |
| NCLAg | ---------- | ---------- | GGGRGGGRGG | GRG------- | ----G-GGGG | -FGAKKHGSI |
| NCLAf | G--SGGYGGR | GG---GGGGY | GGSRGG-GGG | YDDR-GS--- | SGGYGGGGGG | --GYSGGGGR |
| NCLGo | ---------- | ---------- | ---------- | ---------- | ---------- | ---------- |
| NCLScom | ---------- | ---------- | GRGRGGSSGG | FGS------- | ----G-GSGG | GFKAKRTGAI |
| NCLSr | ---------- | ---------- | PGGRGGGRGG | GRG------- | ------GGAS | AFGAKKHGSI |
| NCLTan | ---------- | ---------- | YGGRSGGRSG | GRG------- | ----GGRGTD | AFAAKKHGSI |
| NCLA1 | VTPPSPGTRV | QFLPLREPRL | HLTKNWKRGN | PRQDRA---- | ----GLGLKN | DAPAVQTVRM |
| NCLA2 | ---------- | ---------- | ---------- | ---------- | ---------- | LWHKEYNNNN |
| NCLAp | ------GGGR | GYMGGGGGGR | GGGRGGGRGG | GRGGGR---- | ----GGGFSS | PSKARNSGAI |
| NCLAr1 | ------GGRX | -----GGGRF | GGGRFGGRGG | -RGGGR---- | ----GGRGDS | TSGSKX---- |
| NCLAr2 | ---------- | ---------- | ---------- | ---------- | ---------- | -QAAVAAVVV |
| NCLE | ------GGGR | GYMG-GGGGR | GGGRGGGRGG | GRGGGR---- | ----GGGFS- | ATKSKNSGAI |
| NCLFc1 | GGRSGGYDDR | GGGYSGGGGY | GGGGGGYSGG | YDDR-GG--- | ----GRGGGT | --GYGGG--- |
| NCLFc2 | ---------- | ---------- | -FGGGGLGAG | G--------- | ---------- | ---------- |
| NCLFk | ---------- | ------SGDR | GGRGGRGGGG | GGRGGN---- | ----SLH--- | ----KASGGI |
| NCLFs1 | ---------- | --------GR | GGDSGGRGGR | GRGGGN---- | ----SFN--- | ---AKKTGGI |
| NCLFs2 | GGYGGGGYGG | GGGGYDRGGY | GGGGGGYDRG | GYDRGG---- | YDDRRGGGG- | GGGYGGA--- |
| NCLFs3 | G-YGGGYGGG | GGGGYDRGGY | GGGGGGYDRG | GYDRGG---- | YDDRRGGGG- | GGGYGGG--- |

```
NCLPa      ---------- ---------- -GGFGGRGGG RDGGGS---- ----TLH--- ----KKSGGI
NCLPau     ---------- ---------- --GRGGGRGG GRGGGS---- ----TLH--- ----KASGGI
NCLPd      ---------- ---------- -SGDRGRGGG R-GGGS---- ----SLH--- ----KKSGGI
NCLPf      ---------- ---------- -EADEEAAAA VVAAPF---- ----TRN--- ----RAASQI
NCLPh      ---------- ---------- --GGRGRGGG RGGDSN---- ----SLH--- ----KKSGGI
NCLPp      ---------- ---------- --GRG----- GRGGGS---- ----TLH--- ----KKSGGI
NCLNpa     GGYGGGYSDR GG-YDDRGGY GGSSGGGGGG GYDR-GY--- ERGGGSGGGG --GYSGSHRG
NCLNp      ---------- ---------- --------GF SPANNT---- ----LKA--- ----KKSGGI
NCLNi1     G-GGG-YEDR SG--YGGGGS GGGYRGGGGG YDDRSGG--- YGDRGGGGGG --GYRGG---
NCLNi2     G-GGGGYEDR SG--YGGGG- --GYRGGGGG YDDRSGG--- YGDRGSGGGG YRGYVGG---
NCLPsp     ---------- ---------- GAGRGGGRG- ----ASP--- ---TGRGGAS NPRSKNAGAI
NCLTv      ---------- ---------- GGGGGFGAR- ---------- ----DRSTSP KPVNKNKGSI
NCLTs      ---------- ---------- GGGGGFGAR- ---------- ----DRSTSP KPVNKNKGSI
NCLTc      ---------- ---------- GGFKNGGDR- ---------- ----GRSTSP KPVNKNKGAI
NCLNsa     ---------- -------SEE GAGSGGGRGG GRGGGRGG-- ---GRGGGET GRSTRTEGAF
NCLNg      ---------- ---------G GGGFGGGRGG GRGGGRGG-- ---GRGGGRD GAINKNRGSI
NCLTmi     ---------- ---------- GGGRGGGRGG RDGGGRGG-- ---GRGGGR- ---CVCAGGV
NCLCr      ---------- ---------- GGGRGGFGGR GGGRGGGR-- ---GGGRGGS FTAAVSKGAI
NCLBs      ---------- ---------- ---------- --GRGRP--- ---------- ---RRY----
NCLMc      ---------- ---------- ---------- -GSRGGSG-- ---RGGRGGF SKPPAGKGNI
NCLSl      ---------- ------RGGF RGGDRGGSRG GFRGGDR--- ---GGSRGGF RGGDRGG---
NCLCsu     ---------- ---------- -GGRGGA--- ---------- --------QS AAKAAKDGTV
NCLEv      ---------- -----MPP-- -M-------- ---------- ---------- ----------
NCLPg      ---------- -----MPPPP GMGMGGPPPG MPRY------ ---------- ----------
NCLEt      ---------- ---------- RGGRGGG--- ---------- --------AS AAKAANAGIV
NCLEn      ---------- ---------- RGGRGGG--- ---------- --------AS AVKAANAGNV
NCLEm      ---------- ---------- RGGRGGG--- ---------- --------AS SAKAANAGNV
NCLEb      ---------- ---------- RGGRGGG--- ---------- --------SS SAKAANAGNV
NCLEmi     ---------- ---------- RGGRGGG--- ---------- --------AS SAKAANAGNV
NCLEa      ---------- ---------- RGGRGGG--- ---------- --------AS SAKAANAGTV
NCLEp      ---------- ---------- RGGRGGG--- ---------- --------AS SAKAANAGNV
NCLPo1     ---------- ---------- GKGKGKG--- ---------- ---------S GIAAAARGTI
NCLPo2     ---------- ---------- GKGKGKG--- ---------- ---------S GIAAAR-GTI
NCLPm      ---------- ---------- GKGKGKG--- ---------- ---------S GIAAAARGTI
NCLCc1     ---------- ---------- FGGRGGG-RG RGGYGGRG-- -------GPN RAQAQSRGSI
NCLCc2     ---------- ---------- FGGRGGGGRG GSGFGGRGRG RGGGFGGGAS RAQAQNKGSI
NCLSmi     ---------- ---------- GRGRGGGRGR GGTPT----- ---------- -SGSKNKGSI
NCLPbr     ---------- ---------- -GGRGGGRGA PRGRGHP--- ---------- ---RGRGSGV
NCLGt      ---------- ---------- RGGGRG-GGR GGFGG----- ---------- -TPNKNKGSI
NCLG       ---------- ---------- RGGAFGSGGR GGSAST---- ---------- -TPNRNRGSI
NCLDlu     ---------- ---------R TPGGRGGTPG GRGGSRGG-- ---GRSGGRG SDRAKGAGSI
NCLP       ---------- ---------- --RGRGDAAD GQKRPRS--- ---------- ----------

           ....|....| ....|....| ....|....| ....|....| ....|....| ....|....|
              1205       1215       1225       1235       1245       1255

NCLHs      GFRGGRGGGG DHKPQGKKTK FE-------- ---------- ---------- ----------
NCLAt1     P--QGKKTTF GDE------- ---------- ---------- ---------- ----------
NCLAt2     ESSKGTKTVF NDEE------ ---------- ---------- ---------- ----------
Nsr1pSc    ASFAGSKKTF D--------- ---------- ---------- ---------- ----------
NCLAs      AGFAGKKITF D--------- ---------- ---------- ---------- ----------
NCLAsu     ADFKGNKITF ---------- ---------- ---------- ---------- ----------
NCLCd      SEFKGNKITF D--------- ---------- ---------- ---------- ----------
NCLCp      ---------- ---------- ---------- ---------- ---------- ----------
```

```
NCLCh        ---------- ---------- ---------- ---------- ---------- ----------
NCLCw        ---------- ---------- ---------- ---------- ---------- ----------
NCLCm        AAFSGTKMTF D--------- ---------- ---------- ---------- ----------
NCLCcr       V-TKGYGGGG Y---GGGGY- ---------- ---------- ---------- ----------
NCLEa        AEFKGNKITF D--------- ---------- ---------- ---------- ----------
NCLEs        AEFSGNKITF D--------- ---------- ---------- ---------- ----------
NCLO         AELMGRAMRI DYAKSGGVSG GRDGGGGRGG GGGRGGGG-- ---------- ----------
NCLOs        VECLGRPMRI DFGKPRPPP- ---------- ---------- ---------- ----------
NCLMp        QEFSGNKITF D--------- ---------- ---------- ---------- ----------
NCLPi        SGFAGKKITF D--------- ---------- ---------- ---------- ----------
NCLDf        AEFAGNKITF D--------- ---------- ---------- ---------- ----------
NCLDc        AAFAGNKVTF D--------- ---------- ---------- ---------- ----------
NCLPj        --GVVSGGGG GISGGGGGG- ---------- ---------- ---------- ----------
NCLLda       ---------- ---------- ---------- ---------- ---------- ----------
NCLCa        TEFAGNKITF D--------- ---------- ---------- ---------- ----------
NCLCc        AAFTGNKITF D--------- ---------- ---------- ---------- ----------
NCLChd       AEFSGNRTTF D--------- ---------- ---------- ---------- ----------
NCLCmu       G-YGGGYDDR S---GGGG-- ---------- ---------- ---------- ----------
NCLCn        AAFAGNKITF D--------- ---------- ---------- ---------- ----------
NCLC         QEFSGNKIKF DD-------- ---------- ---------- ---------- ----------
NCLCt1       TEFAGTKITF D--------- ---------- ---------- ---------- ----------
NCLCt2       --YEDRGDNG G---GGGG-- ---------- ---------- ---------- ----------
NCLDbGSO10   TDIMGRAVRV DFAGGRRGDG GGRGGGRGGG RGGGRGGGRG GGFGGGRGGG RGRGGGRGGS
NCLDbGSO10   EQRKKRKKNK KVKEETNDAK ---------- ---------- ---------- ----------
NCLSco1      AAFSGKKMTF D--------- ---------- ---------- ---------- ----------
NCLSco2      G-YSGGGGDR Y---QGGG-- ---------- ---------- ---------- ----------
NCLSd        AAFSGKKMTF D--------- ---------- ---------- ---------- ----------
NCLSj        ---------- ---------- ---------- ---------- ---------- ----------
NCLSma       AAFSGKKMTF D--------- ---------- ---------- ---------- ----------
NCLSme       AAFSGNKITF D--------- ---------- ---------- ---------- ----------
NCLTa        AEFAGTKMTF D--------- ---------- ---------- ---------- ----------
NCLTg        ---------- ---------- ---------- ---------- ---------- ----------
NCLTm        AAFSGNKITF D--------- ---------- ---------- ---------- ----------
NCLTn        SEFKGNKITF D--------- ---------- ---------- ---------- ----------
NCLTp        ---------- ---------- ---------- ---------- ---------- ----------
NCLTo1       ---------- ---------- ---------- ---------- ---------- ----------
NCLTo2       G-YGGGGGGG Y---GGG--- ---------- ---------- ---------- ----------
NCLTw        ---------- ---------- ---------- ---------- ---------- ----------
NCLAg        AAFQGNKITF D--------- ---------- ---------- ---------- ----------
NCLAf        SSYDDRGGGG G-YSGGGGGY DRG------- ---------- ---------- ----------
NCLGo        ---------- ---------- ---------- ---------- ---------- ----------
NCLScom      AEFSGNKITF D--------- ---------- ---------- ---------- ----------
NCLSr        SEFKGTKVTF D--------- ---------- ---------- ---------- ----------
NCLTan       SEFKGNKITF D--------- ---------- ---------- ---------- ----------
NCLA1        LPVSGQRLPT HDDPSTSX-- ---------- ---------- ---------- ----------
NCLA2        NNHTGNPRQQ QT-------- ---------- ---------- ---------- ----------
NCLAp        SAFSGTKITF D--------- ---------- ---------- ---------- ----------
NCLAr1       ---------- ---------- ---------- ---------- ---------- ----------
NCLAr2       AVAASVAVAX ---------- ---------- ---------- ---------- ----------
NCLE         AGFTGT---- ---------- ---------- ---------- ---------- ----------
NCLFc1       --YDDRGGGG GGYSGGGGG- ---------- ---------- ---------- ----------
NCLFc2       ---------- ---------- ---------- ---------- ---------- ----------
NCLFk        ADFAGKKITF ---------- ---------- ---------- ---------- ----------
NCLFs1       TAFAGKKISF D--------- ---------- ---------- ---------- ----------
```

```
NCLFs2       --ADNNTDGG G---GGG--- ---------- ---------- ---------- ----------
NCLFs3       --YGEIKKGG G---GGGG-- ---------- ---------- ---------- ----------
NCLPa        ASFAGKKITX XXXSGGIASF AGKKX----- ---------- ---------- ----------
NCLPau       ASFAGKKITF D--------- ---------- ---------- ---------- ----------
NCLPd        ADFAGKKITF D--------- ---------- ---------- ---------- ----------
NCLPf        S--PGRK--- ---------- ---------- ---------- ---------- ----------
NCLPh        ADFAGKKITF D--------- ---------- ---------- ---------- ----------
NCLPp        ASFAGKKITF D--------- ---------- ---------- ---------- ----------
NCLNpa       GGYDDRGGGG Y---GWSGR- ---------- ---------- ---------- ----------
NCLNp        AGFAGKKITF D--------- ---------- ---------- ---------- ----------
NCLNi1       --YEDRGGGG G---GGGGG- ---------- ---------- ---------- ----------
NCLNi2       --YEDRGGGG G---GGGGG- ---------- ---------- ---------- ----------
NCLPsp       SEFKGNKITF D--------- ---------- ---------- ---------- ----------
NCLTv        VAGTGKKITF ---------- ---------- ---------- ---------- ----------
NCLTs        VAGTGKKITF ---------- ---------- ---------- ---------- ----------
NCLTc        MAGSGKKITF D--------- ---------- ---------- ---------- ----------
NCLNsa       RHLRETRRHS TTEQMAVGDV VGADRCGGKG GEREGKREGE GAGRTWRKGA RAERAHAAGV
NCLNg        QAFAGNKKTF DD-------- ---------- ---------- ---------- ----------
NCLTmi       ---------- ---------- ---------- ---------- ---------- ----------
NCLCr        QEFAGNKMTF D--------- ---------- ---------- ---------- ----------
NCLBs        ---------- ---------- ---------- ---------- ---------- ----------
NCLMc        AEYQGNKMKF ---------- ---------- ---------- ---------- ----------
NCLSl        --FRGRGGFR G---NRD--- ---------- ---------- ---------- ----------
NCLCsu       QEFKGTSKTF DSDSE----- ---------- ---------- ---------- ----------
NCLEv        ---------- ---------- ---------- ---------- ---------- ----------
NCLPg        ---------- ---------- ---------- ---------- ---------- ----------
NCLEt        QEFQGKKTTF D-DSD----- ---------- ---------- ---------- ----------
NCLEn        QEFQGKKMTF D-DSD----- ---------- ---------- ---------- ----------
NCLEm        QEFQGKKTTF E-DSD----- ---------- ---------- ---------- ----------
NCLEb        QEFQGKKTTF E-DSD----- ---------- ---------- ---------- ----------
NCLEmi       QEFQGKKTTF E-DSD----- ---------- ---------- ---------- ----------
NCLEa        QEFQGKKTTF D-DSD----- ---------- ---------- ---------- ----------
NCLEp        QEFQGKKTTF D-DSD----- ---------- ---------- ---------- ----------
NCLPo1       QEFAGHKTTF ADDDDSDSD- ---------- ---------- ---------- ----------
NCLPo2       QEFAGHKTTF ADDDDSDSD- ---------- ---------- ---------- ----------
NCLPm        QEFAGHKTTF ADDDDSDSD- ---------- ---------- ---------- ----------
NCLCc1       QQFQGKSMTF DD-------- ---------- ---------- ---------- ----------
NCLCc2       QQFQGKSMTF DD-------- ---------- ---------- ---------- ----------
NCLSmi       AVSTGKKMTF D--------- ---------- ---------- ---------- ----------
NCLPbr       GAFSGQKTKF ---------- ---------- ---------- ---------- ----------
NCLGt        QESTGKKITF DD-------- ---------- ---------- ---------- ----------
NCLG         QESQGKKISF DDDE------ ---------- ---------- ---------- ----------
NCLDlu       QEFSGKKMSF DD-------- ---------- ---------- ---------- ----------
NCLP         ---EGREEPK GKRQHAAY-- ---------- ---------- ---------- ----------

             ....|....| ....|....| .
                1265       1275
NCLHs        ---------- ---------- -
NCLAt1       ---------- ---------- -
NCLAt2       ---------- ---------- -
Nsr1pSc      ---------- ---------- -
NCLAs        ---------- ---------- -
NCLAsu       ---------- ---------- -
NCLCd        ---------- ---------- -
```

```
NCLCp        ---------- ---------- -
NCLCh        ---------- ---------- -
NCLCw        ---------- ---------- -
NCLCm        ---------- ---------- -
NCLCcr       ---------- ---------- -
NCLEa        ---------- ---------- -
NCLEs        ---------- ---------- -
NCLO         ---------- ---------- -
NCLOs        ---------- ---------- -
NCLMp        ---------- ---------- -
NCLPi        ---------- ---------- -
NCLDf        ---------- ---------- -
NCLDc        ---------- ---------- -
NCLPj        ---------- ---------- -
NCLLda       ---------- ---------- -
NCLCa        ---------- ---------- -
NCLCc        ---------- ---------- -
NCLChd       ---------- ---------- -
NCLCmu       ---------- ---------- -
NCLCn        ---------- ---------- -
NCLC         ---------- ---------- -
NCLCt1       ---------- ---------- -
NCLCt2       ---------- ---------- -
NCLDbGSO10   SFGAKKSGSI AAFQGNKITF D
NCLDbGSO10   ---------- ---------- -
NCLSco1      ---------- ---------- -
NCLSco2      ---------- ---------- -
NCLSd        ---------- ---------- -
NCLSj        ---------- ---------- -
NCLSma       ---------- ---------- -
NCLSme       ---------- ---------- -
NCLTa        ---------- ---------- -
NCLTg        ---------- ---------- -
NCLTm        ---------- ---------- -
NCLTn        ---------- ---------- -
NCLTp        ---------- ---------- -
NCLTo1       ---------- ---------- -
NCLTo2       ---------- ---------- -
NCLTw        ---------- ---------- -
NCLAg        ---------- ---------- -
NCLAf        ---------- ---------- -
NCLGo        ---------- ---------- -
NCLScom      ---------- ---------- -
NCLSr        ---------- ---------- -
NCLTan       ---------- ---------- -
NCLA1        ---------- ---------- -
NCLA2        ---------- ---------- -
NCLAp        ---------- ---------- -
NCLAr1       ---------- ---------- -
NCLAr2       ---------- ---------- -
NCLE         ---------- ---------- -
NCLFc1       ---------- ---------- -
NCLFc2       ---------- ---------- -
NCLFk        ---------- ---------- -
```

```
NCLFs1      ---------- ---------- -
NCLFs2      ---------- ---------- -
NCLFs3      ---------- ---------- -
NCLPa       ---------- ---------- -
NCLPau      ---------- ---------- -
NCLPd       ---------- ---------- -
NCLPf       ---------- ---------- -
NCLPh       ---------- ---------- -
NCLPp       ---------- ---------- -
NCLNpa      ---------- ---------- -
NCLNp       ---------- ---------- -
NCLNi1      ---------- ---------- -
NCLNi2      ---------- ---------- -
NCLPsp      ---------- ---------- -
NCLTv       ---------- ---------- -
NCLTs       ---------- ---------- -
NCLTc       ---------- ---------- -
NCLNsa      PRGGWGEEEG RVQAAVA--- -
NCLNg       ---------- ---------- -
NCLTmi      ---------- ---------- -
NCLCr       ---------- ---------- -
NCLBs       ---------- ---------- -
NCLMc       ---------- ---------- -
NCLSl       ---------- ---------- -
NCLCsu      ---------- ---------- -
NCLEv       ---------- ---------- -
NCLPg       ---------- ---------- -
NCLEt       ---------- ---------- -
NCLEn       ---------- ---------- -
NCLEm       ---------- ---------- -
NCLEb       ---------- ---------- -
NCLEmi      ---------- ---------- -
NCLEa       ---------- ---------- -
NCLEp       ---------- ---------- -
NCLPo1      ---------- ---------- -
NCLPo2      ---------- ---------- -
NCLPm       ---------- ---------- -
NCLCc1      ---------- ---------- -
NCLCc2      ---------- ---------- -
NCLSmi      ---------- ---------- -
NCLPbr      ---------- ---------- -
NCLGt       ---------- ---------- -
NCLG        ---------- ---------- -
NCLDlu      ---------- ---------- -
NCLP        ---------- ---------- -
```
