## Supplementary material for "Structural features of the amino acid sequences of Chromista nucleolin-like proteins": alignment of sequences used for phylogenetic reconstruction

Supplementary data 2. Sequences using for phylogenetic analysis. Color boxes point the special regions of N-terminal domain according to Figure 3. As group 4 is variable, we pointed only acidic regions of separate sequences by red letters and NLS.

```
                ....|....| ....|....| ....|....| ....|....| ....|....| ....|....|
                     5          15         25         35         45         55
1
NCLSco1         ---------M A--SSSSSSS SS-------- ---------- ---------- --DSSSSSS-
NCLSd           ---------M A--SSSSSSS SS-------- ---------- ---------- --DSSSSSS-
NCLSj           ---------M A--SSSSSSS SS-------- ---------- ---------- --DSSSSSSS
NCLSme          ---------M A--SSSSSSS SSS------- ---------- ---------- --DSSSSSSS
NCLDc           ---------M A--SSSSSSS SS-------- ---------- ---------- ---SDSSS-E
NCLTm           ---------M A--SDSSSSS SS-------- ---------- ---------- ---SDSSSSE
NCLCm           ---------M S--SSSSSSS SS-------- ---------- ---------- ------DSSS
NCLTo1          ---------M S--SSSSSSS SS-------- ---------- ---------- ---SSSSSSS
NCLTw           ---------M T--SSSSSSS SS-------- ---------- ---------- ---SSESSSS
NCLTa           ---------M A--SSSSSSS DS-------- ---------- ---------- ------SSSE
2
NCLFs1          -------MNL V--KQVTSTM GADKKEILR- ---------- ---------- QKTEAAEAAA
NCLEs           ---------M A--KKVLENE ---------- ---------- ---------- KAVEEESSSS
NCLNp           ---------M A--KISKEKL ---------- ---------- ---------A KELKAKQKAV
NCLFk           ---------M A--KSSKKEK KVKDSKPDKK AAKKAKAKEE KAAKKASKKA AELEAQQKAA
NCLPf           ---------M A--KASKD-- ---------- ---------- ---------- KDLKAKQKAA
NCLPh           ---------M A--KANKE-- ---------- ---------- ---------- KVLKAKQKAA
NCLPa           ---------- M--ASSKE-- ---------- ---------- ---------- KDLKAKQKAA
NCLPd           ---------- M--AGNKE-- ---------- ---------- ---------- KELKAKLKAA
3
NCLPsp          ---------M G--KTK---T DKKAKLAQE- ---------- ---------- -LAEQKKKMA
NCLPi           ---------M G--SDK---D KK-------- ---------- ---------- -LKKAAKKEK
NCLEa           ---------M G--SKVRKEK KEKK------ ---------- ---------- -SKKSPKKED
NCLTn           ---------M G--SKV-KND KK-------- ---------- ---------- -AKKQAKKEA
NCLTan          ---------M G--TKD-KSD KKKA------ ---------- ---------- -AKKQAKKEA
NCLSr           ---------M G--TKTDKSE KK-------- ---------- ---------- -AKKQAKKEL
NCLAg           ---------M G--SKS-KED KK-------- ---------- ---------- -AKKAAKKAA
NCLGo           ---------- S--SSSSSDS SS-------- ---------- ---------- ----SSEDEK
NCLChd          ---------M G--SKT---D KK-------- ---------- ---------- -AKKAAKKAL
NCLCt1          ---------M GSKSKA---D KK-------- ---------- ---------- -AKKAAKKAE
NCLCd           ---------M G--SKQ---D KK-------- ---------- ---------- -EKRAKKKAV
NCLDf           ---------M G--SKP---D KK-------- ---------- ---------- -SKKSKESKK
NCLCa           ---------M G--SKS---D KK-------- ---------- ---------- -AKKAQKKAL
NCLCn           ---------M G--SKT---D KK-------- ---------- ---------- -LKKAAKKAL
NCLAsu          ---------- ---------- ---------- -------MGS KVKPD----- KKAAK-----
NCLDbGSO10      ---------- ---MG----- -SKTPKSEKK S-----KKAK KAESKESKLK EKKTD-----
4
NCLPbr          ---------M GSTKAATVVK KPTR------ ---------- ---------- --SKDKYNKN
NCLP            ---------- ---------- ---------- ---------- ---------- ----------
NCLAt1          -------MGK S--KSATKVV AEIK------ ---------- ---------- -ATKPLKKGK
NCLAt2          -------MGK SSKKSVTEVE TPAS------ ---------- ---------- -MTKPLKKGK
Nsr1pSc         ---------- --MAKTTKVK G--------- ---------- ---------- --NKKEVKAS
NCLMc           ---------- --MPKTKKSK KEAK------ ---------- ---------- -LSKKELKAK
NCLSl           ---------- ---------- ---------- ---------- ---------- ----------
NCLBs           ---------- ---------- ---------- ---------- ---------- ----------
NCLCsu          MKQKSAPVAV SSSASSSDSD SSEEEPQAKV QKATLKRVAE GGAKQKAQRK NRGKAPPTDS
NCLHs           ------MVKL AKAGKNQGDP KKMAPPPKEV EEDSEDEEMS EDEEDDSSGE EVVIPQKKGK
NCLCr           ---------M ASTDVELHQL VHQFLKAEG- ---------- ---------- -LSKSAKAFA
NCLDlu          -------MAA TADDHTILSL VHEFLTSKG- ---------- ---------- -LTRTANALL
5
NCLNsa          ---------M GKSSKSKADK TLG------- ---------- --------AD VVVQSKGGSA
NCLNg           ---------M GKSSKSKADK TLG------- ---------- --------AD VVVQSKGGSA
NCLTmi          ---------- ---------- ---------- ---------- ---------- -MVRSPPPSK
NCLGt           ---------- ---------- ---------- ---------- ---------- ----------
NCLG            ---------M TKASAGIKRK AGDEPT---- ---------- ---------- -KKKASKESE
6
NCLCh           ----MGKDKK S--KGKDKSA KK-------- ---------- ---------- ---ASKSKIK
NCLO            ----MGSKSK S--KDSKKDA KR-------- ---------- ---------- ---AEKKVMK
NCLTc           -------MAS A--TVDKKAA KK-------- ---------- ---------- ---LAKKQKA
NCLTv           ---------M G--KSATKDA KK-------- ---------- ---------- ---IAKKAKA
NCLTs           ---------M G--KSATKDA KK-------- ---------- ---------- ---IAKKAKA
```

```
              ....|....| ....|....| ....|....| ....|....| ....|....| ....|....|
                  65         75         85         95        105        115
1
NCLSco1       EDEKK----- ---------- ---------- ---------- ---------VV D--TPKIAKK
NCLSd         EDEKK----- ---------- ---------- ---------- ---------VV D--TPKIAKK
NCLSj         EEEKK----- ---------- ---------- ---------- ---------VV D--TPKIAHK
NCLSme        EDEKK----- ---------- ---------- ---------- ---------VV D--TPKIAHK
NCLDc         EDKKK----- ---------- ---------- ---------- ---------VI E--KKPIATK
NCLTm         EETKK----- ---------- ---------- ---------- ---------VV ET-KKPIAKK
NCLCm         EDEKK----- ---------- ---------- ---------- ---------VV H---KPIATK
NCLTo1        EDEKP----- ---------- ---------- ---------- ---------KV E--EKKVIAR
NCLTw         EDEKK----- ---------- ---------- ---------- ---------IV N--EKPIAVK
NCLTa         EETKK----- ---------- ---------- ---------- ---------VV TTPVAKKTKA
2
NCLFs1        KEAQE----- ---------- ---------- ---------- ---------AT ARAERLAKEV
NCLEs         DSDSS----- ---------- ---------- ---------- ---------SS SDSSDSEDEA
NCLNp         EEAKQ----- ---------- ---------- ---------- ---------IA QEALKKAERL
NCLFk         EEAKK----- ---------- ---------- ---------- ---------AA DDAIKAAEKL
NCLPf         KEAKK----- ---------- ---------- ---------- ---------AA DDAIKKAEKL
NCLPh         KEAKK----- ---------- ---------- ---------- ---------AA DDAIKKAEKL
NCLPa         KEAKK----- ---------- ---------- ---------- ---------AA DDAIKKAEKL
NCLPd         KEAKK----- ---------- ---------- ---------- ---------AA DEAVKKAEKL
3
NCLPsp        DLEAK----- ---------- ---------- ---------- --------LAA EAVVESSSDS
NCLPi         AAKAK----- ---------- ---------- ---------- --------AEA EAAAKKAADL
NCLEa         KERVE----- ---------- ---------- ---------- --------KLA AEKAAAEKAA
NCLTn         AEKARKAA-- ---------- ---------- ---------- -------EQAA EEAA---RKA
NCLTan        AEKAR----- ---------- ---------- ---------- --------KEA EEAA---KKA
NCLSr         LEKAK----- ---------- ---------- ---------- --------KAA EEAA---AKA
NCLAg         AEKAA----- ---------- ---------- ---------- --------KEA EEAA---RKA
NCLGo         PTTKK----- ---------- ---------- ---------- ---------TM T--VTKKKTK
NCLChd        VEKLA----- ---------- ---------- ---------- --------KEA AAAA---AAA
NCLCt1        AERLA----- ---------- ---------- ---------- --------KEA EEAA---KAA
NCLCd         AEKLA----- ---------- ---------- ---------- --------KEA KELA---AAA
NCLDf         AAKLAA---- ---------- ---------- ---------- -------KKEA EEAARKAKEA
NCLCa         AEKLA----- ---------- ---------- ---------- --------REA EEAA---KKA
NCLCn         AEKLA----- ---------- ---------- ---------- --------KEA EEAA---KAA
NCLAsu        ------KAEA AR-------- ---------- --LKADAEEL ARKAEELKKK ATKKDETSSD
NCLDbGSO10    -----KEAKK AAKKLKKE-- ---------- EEKAKKAAAE AAAKKAAEEA AAKKAAEEAA
4
NCLPbr        KDKNKG---- ---------- ---------- ---------- ---------AV KDAPVKNAAK
NCLP          ---------- ---------- ---------- ---------- ---------- ----------
NCLAt1        REPED----- ---------- ---------- ---------- ---------DI DTKVSLKKQK
NCLAt2        RDAEE----- ---------- ---------- ---------- ---------DL DMQVT-KKQK
Nsr1pSc       KQAKE----- ---------- ---------- ---------- ---------EK AKAVSSSSS-
NCLMc         AKAQE----- ---------- ---------- ---------- ---------SE ESSDSDSDSS
NCLSl         ---------- ---------- ---------- ---------- ---------- ----------
NCLBs         ---------- ---------- ---------- ---------- ---------- ----------
NCLCsu        SEDTSSDDES TG-------- ---------- ---------- ---TVRKSLV ESSAAAVQRA
NCLHs         KAAATS---- ---------- ---------- ---------- -------AKKV VVSPTKKVAV
NCLCr         KEAAK----- ---------- ---------- ---------- -------VGAV TEEPGQSVTL
NCLDlu        SEVKTPVQPL KPGTPPLAQL VRLRASAPAA PPAETSEESS DDSDESSKAA AKESSDDSDD
5
NCLNsa        EGTKAV---- ---------- ---------- ---------- -------GKKG KKAAKPVEPS
NCLNg         EGTKAV---- ---------- ---------- ---------- -------GKKG KKAAKPVEPS
NCLTmi        AVAAAN---- ---------- ---------- ---------- -------GKKA Q--------E
NCLGt         ---------- ---------- ---------- ---------- ---------- ----------
NCLG          SDSSSDSE-- ---------- ---------- ---------- -------SDEE EVKAKKPVKA
6
NCLCh         AELKA----- ---------- ---------- ---------- ---------KA DREAAEALAA
NCLO          KAAAE----- ---------- ---------- ---------- ---------AA KKAAEAAAEA
NCLTc         EALKL----- ---------- ---------- ---------- ---------KA KEAAEAAAKA
NCLTv         AAKQK----- ---------- ---------- ---------- ---------EL ELAMKKAQEE
NCLTs         AAKQK----- ---------- ---------- ---------- ---------EL ELAMKKAQEE
```

```
             ....|....| ....|....| ....|....| ....|....| ....|....| ....|....|
                125        135        145        155        165        175
1
NCLSco1      KGSSSSSSS- ---------- ---------- -SSS--DSS- ---------- ----------
NCLSd        KGSSSSSSS- ---------- ---------- -SSSSSDSS- ---------- ----------
NCLSj        KGSSSSSSS- ---------- ---------- -SSD---SS- ---------- ----------
NCLSme       KGSSSSSSS- ---------- ---------- -SSSSSDSS- ---------- ----------
NCLDc        KS-------- ---------- ---------- ---------- ---------- ----------
NCLTm        KGSSDNSSS- ---------- ---------- -SGSDSSDS- ---------- ----------
NCLCm        KKDSSSSSS- ---------- ---------- --SSSSDSS- ---------- ----------
NCLTo1       KKSSSSSSS- ---------- ---------- --SS-SSSS- ---------- ----------
NCLTw        KGSSSSASS- ---------- ---------- --SSGSDSS- ---------- ----------
NCLTa        KASSSSKKSK KEV------- ---------- VASSSSDSS- ---------- ----------
2
NCLFs1       KELAEQLKSS GKK------- ---------- TKATKSVVS- ---------- ----------
NCLEs        PPKKVEDKVV AKKKEK---- ---------- --ESNSSSES ---------- ----------
NCLNp        AEEVAKMESE IEGDGKP--- -------SKP EAKAVENDD- ---------- ----------
NCLFk        AEELLKKE-D SGSDSD---- ---------- ---SSSSS-- ---------- ----------
NCLPf        AEEVAKLESE MEAEAK---- ---------- --KAAKKAES ---------- ----------
NCLPh        AEEVAKLESE IKKEAK---- ---------- --KSDSSSS- ---------- ----------
NCLPa        AEEVAKMESE IAAAAK---- ---------- --KAAESS-- ---------- ----------
NCLPd        AEEVAKLESD INAEAK---- ---------- --KAEESS-- ---------- ----------
3
NCLPsp       DSDSDSSSSE DEKK------ ---------E VTKKVTDES- ---------- ----------
NCLPi        AAAAAKAAAA AAAK------ ---------T ADSSDSSDS- ---------- ----------
NCLEa        AEKAAAEKAA AEKA------ ---------A ESDSDSSSS- ---------- ----------
NCLTn        AELKKIAEAE E--------- ---------- SSDSDSSDS- ---------- ----------
NCLTan       AELAKKAAAE EKS------- ---------S SDDSDSSDD- ---------- ----------
NCLSr        AELAKAAESS D--------- ---------- ---SSSSDS- ---------- ----------
NCLAg        AELAKKAAEA E--------- ---------- SSDSSSSDS- ---------- ----------
NCLGo        KEESSDSSS- ---------- ---------- --SSDSDSS- ---------- ----------
NCLChd       AEAAKKAAEA S-DS------ ---------D SSDSDSSDS- ---------- ----------
NCLCt1       AEAAKKAAEE AADD------ ---------D SSSSSSSSS- ---------- ----------
NCLCd        AAAAAAAAAN DSD------- ---------S DSDSDSSSS- ---------- ----------
NCLDf        AEAAKRAAEE AAAK------ ---------A KEEASSDSS- ---------- ----------
NCLCa        AEAAKKAAEE AAAA------ ---------A EAEGSDSDS- ---------- ----------
NCLCn        AEAAKKAAIE AAAE------ ---------S DSDSSSSSS- ---------- ----------

NCLAsu       EDSDASS--- ---------- ---------- -------SSD SESE------ ----------
NCLDbGSO104  AKKAAEE--- ---------- ---------- -------AAK KAAEEA---- ----------
4
NCLPbr       DVEAKKAAAA AKKVAA---- ---------- AAPAQKPAN- ---------- ----------
NCLP         ---------- ---------- ---------- ---------- ---------- ----------
NCLAt1       KDVIAAVQKE KAVKKV---- ---------- PKKVESSDDS DSESEEEEK- ------AKKV
NCLAt2       KELIDVVQKE KAEKTV---- ---------- PKKVESSSSD ASDSDEEEK- ------TKET
Nsr1pSc      ---------- ---------- ---------- --ESSSSSS- ---------- ----------
NCLMc        VEEPVKTKKT KKAPVV---- ---------- EKKNDSSDSD EAPAAGKKRK PETVKATTKK
NCLSl        ---------- ---------- ---------- ---------- ---------- ----------
NCLBs        ---------- ---------- ---------- ---------- ---------- ----------
NCLCsu       KQAVRATALS KQKAPAKPSA GSDGSSSDST EDETPQTKHA SKAKSVSKSA VADPSSSDDS
NCLHs        ATPAKKAAVT PGKKAAATP- ---------- AKKTVTPAKA VT-------- ----------
NCLCr        SEVFKAFSAQ RASKRP---- ---------- REEPKAETS- ---------- ----------
5
NCLDlu       SDESSKAAAE ESSDDS---- ---------- DESSKAAAE- ---------- ----------
NCLNsa       SSSSSDSSSD EEVTPTTNGT AGKKGGKVVA KPVAASSDS- ---------- ----------
NCLNg        SSSSSDSSSD EETAPTTNGT AGKKGGKVVA KPVAASSDS- ---------- ----------
NCLTmi       SSEESESDSD E--------- ---------- ---------- ---------- ----------
NCLGt        ---------- ---------- ---------- ---------- ---------- ----------
NCLG         AAKAAPAKAD KKKAAKK--- ---------E ESSSEEESSD ---------- ----------
6
NCLCh        AKAAAEEAER LAK--AAE-- ---EAAKKEK PSSDSSSSS- ---------- ----------
NCLO         AKKAAEEDAR KAAEDSSD-- ---SGSDSDS SSSSSSSSSX DEAPAPPKKK EGKKDKAKAK
NCLTc        AKLAAEEAAK AAK------- ---------- -MAESDSDS- ---------- ----------
NCLTv        IKRLQEAND- ---------- ---------- --SSSSSDS- ---------- ----------
NCLTs        IKRLQEAND- ---------- ---------- --SSSSSDS- ---------- ----------
```

```
                    ....|....| ....|....| ....|....| ....|....| ....|....| ....|....|
                          185        195        205        215        225        235
1
NCLSco1        ------DSEA EKKA------ ---------- ----AVVVAK KKESKKEAKK KA--------
NCLSd          ------DSEA EKKA------ ---------- ----AVVVAK KKESKKEAKK KA--------
NCLSj          ------DSEA QKK------- ---------- -----VVAKK KKESKKEAKK KA--------
NCLSme         ------DSEA EKK------- ---------- ------VVSK KKESKKEAKK KA--------
NCLDc          ---------- ---------- ---------- ---------- KKESKKEASK KK--------
NCLTm          ------DSDA KAKAKAKASS KKGEKGKKGK KKEEAPVAKK KKDSKKEAAV KKK-------
NCLCm          ------DSDA KAKT------ ---------- ----KKAVAK KKEPKKEVKK EET-------
NCLTo1         ------ESEA EAEAK----- ---------- ----PAPPPK KAES-KKEKK VKA-------
NCLTw          ------DSEA KKKKK----- ---------- ----KTKAVT KKETPKKETP KKV-------
NCLTa          ------DSES DSKK------ ---------- ----KAKAKK AKKSKKESKK EVA-------
2
NCLFs1         ------DDEQ KKAEESDS-- ---------- ---DSSSSSR DSDSDSDDPK KTAV------
NCLEs          ------SSSS DSSDD----- ---------- -------DSS DDEEDEKAEA KPA-------
NCLNp          ------SSDS DSSSS----- ---------- -------SSS DSDSDDDDDE K---------
NCLFk          ------SSSS SSSSS----- ---------- -------SDS DSDSDDDTK- ----------
NCLPf          ------SSDS DSSSS----- ---------- -------SSS DSDSDDDE-- ----------
NCLPh          ------DSDS DSSSS----- ---------- -------SSS SSDSDSGN-- ----------
NCLPa          ------DSDS DSSSS----- ---------- -------SSS DSDSDD---- ----------
NCLPd          ------SSDS DSSSS----- ---------- -------SSS DSDSDDDA-- ----------
3
NCLPsp         ------DSDS DSDSS----- ---------- ----SSDSDS DSDAAPPAKK AAV-------
NCLPi          ------DSDS DSDSD----- ---------- ----SSSSSS SSEDKKKKTK S---------
NCLEa          ------SDSS SSSSS----- ---------- ----SSSSSE DEKPEKKKSE KVV-------
NCLTn          ------DSSD EETPA----- ---------- ----PKASNK KKKAGAEVKK EES-------
NCLTan         ------DSSS DDGSS----- ---------- ----SSSEEK PKKKAPPAKK KES-------
NCLSr          ------DSSS DE-EE----- ---------- ----KAAPVK KGKDSKKKKK EES-------
NCLAg          ------DSSD SE-------- ---------- -----SEDEK PTKTKTKATK KEE-------
NCLGo          ------SDEE EVKAK----- ---------- ----VVVSKK KKEKKVAVAT KKVE------
NCLChd         ------DSDS SSCSD----- ---------- ----SSVEAP AKKVAVKKTK KAK-------
NCLCt1         ------SSSS SSSSS----- ---------- ----SDSDAE MKDADKKEEK KEE-------
NCLCd          ------SSSS SSSSE----- ---------- ----AP--KE TAKIATKKTK APV-------
NCLDf          ------DSDS DSSDD----- ---------- ----SSSSED ESIVTKNKTK KSE-------
NCLCa          ------SSSS SSSSD----- ---------- ----SSDSEN EAAAPEKKAS DAK-------
NCLCn          ------SDSS DSEAE----- ---------- ----APKSKK ETKVVKEKTK VVK-------

NCLAsu         ---------- ---------A EVKVETKTKG ---------- ---------- ------KKND
NCLDbGSO104    ---------- ------AKKA AEEAAAKKDS SDS------- ---------- ---SSSDSSS
4
NCLPbr         ------DAVP VKQAK----- ---------- -----KEKKK GKKTTAAPEP APAP------
NCLP           ---------- ---------- ---------- ---------- ---------- ----------
NCLAt1         PAKKAASSSD ESSDDS---- -----SSDDE PAP------- -KKAVAATNG ----------
NCLAt2         PSKLKDESSS EEEDDS---- -----SSDEE IAPAKKRPEP IKKAKVESSS SDDD------
Nsr1pSc        -------SSS ESESE----- ---------- --------SE SESESSSSS- ----------
NCLMc          PKVEENLSSD DSDSDSVDVI AKPAKKSKKK SAPAKKESSD SDSDSSSEEE KKVA------
NCLSl          ---------- ---------- ---------- -------MPK NQKAVVATKK EET-------
NCLBs          ---------- ---------- ---------- ---------- ---------- ----------
NCLCsu         SGEDASPENF KSKPVKDTQR KTGAAPAKGT QKAVTPTAES SSDDSDEEEP PRKTVSAPPR
NCLHs          ------TPGK KGATPG---- ---------- --KALVATPG KKGAAIPAKG AKN-------
NCLCr          ------SSSS SSS------- ---------- ---------- ---------- ----------
5
NCLDlu         ------ESSD DSDEGSKAAA KESSDDSDGD SDDSSDDESS DGPAKAPVVK PPVASGSLID
NCLNsa         ------DDDS SSSS------ ---------- ----EDEAPA PKKATPAAAK KEKL------
NCLNg          ------DDDS SSSSS----- ---------- ----EDEAPA PKKATPAAGK KEKL------
NCLTmi         ---------- -SSSE----- ---------- ----EDERPA TKVLKAAAKK ----------
NCLGt          ---------- ---------- ---------- ---------- ---------- ----------
NCLG           ------DEDA KKPAAKKADA KKAAPAKKAA KKEESSSEES SSDEDEKPAA KKAT------
6
NCLCh          ------DSSS DSSSD----- ---------- -----SDSDS SVEAKKPAAK A---------
NCLO           EAAKKKDSTS SSSSS----- ---------- -----SSSSS SEEEKKKADK SKAKASKFK-
NCLTc          ------DSDS DSDSD----- ---------- -----SDSDS DNEKKVVKAE P---------
NCLTv          ------DSDS DSDSD----- ---------- -----DEAAT PVAAKAPVVA K---------
NCLTs          ------DSDS DSDSD----- ---------- -----DEAAA PVAAKAPVVA K---------
```

```
             ....|....| ....|....| ....|....| ....|....| ....|....| ....|....|
                  245        255        265        275        285        295
1
NCLSco1      ---------- ---EAKKKAE SSD------- -SSSSSSSSS S-EDEAPAKK ETKKVAETKK
NCLSd        ---------- ---EAKKKAE SSD------- -SSSSSSSSS SSEDEAPAKK ETKKVAETKK
NCLSj        ---------- ---EAEAKKK AEE------- -SSSSSSSSS S-EDEAPAK- ---KVVETKK
NCLSme       ---------- ---EAEAKKK AEE------- -SSSSSSSSS SDEEEAPAK- ---KVAETKK
NCLDc        ---------- ----AAKKAE SSD------- -SSSSSSSSS SEEEAPKKK- AKSKK-EAKK
NCLTm        ---------- -KKEVKKKEE SSD------- -SSSSSSSSS EEEETKKKAP AKTKETKAKK
NCLCm        ---------- -AVKSKKKE- ---------- ---------- ---------- ---VKKEVKK
NCLTo1       ---------- -KKEAEKKEE SSS------- -SSSSSSSSS SSSEEEAPPP PKKKESKQKA
NCLTw        ---------- -EKKADKKKD TPAKKTKKKE ESSDSSSSSS DSESEDEKPA KKSKKEEPKK
NCLTa        ---------- -VKKKKSKKE VVK------- -AESSDSSDS DSEDEAPPP- ---KKKESKK
2
NCLFs1       ---------- KPKVAAKKEE SDD------- -SSSDDSTSD DEDDKAADKM EVEKTEKSAK
NCLEs        ---------E KPVAVAAAKK DDS------- ----SSDSGS S--------- ----SSDSDS
NCLNp        ---------K KPDTSSGKEK DTA------- ---VKEKSDS S--------- ---DSDSSDS
NCLFk        ---------K KPTIATIAKK DSSGS----- -DSSSSDSDS DDDNKKDTKM KEVKKEVKKE
NCLPf        ---------K KPTPAA-TKK EDS------- ---DSSDSDS D--------- ----------
NCLPh        ---------A TKKAAT-KTK DEE------- ---SDSDSDG EEEK----VE SKKKSDSSSS
NCLPa        ---------E KPAAAS-KKE DSD------- ---SSSSSDS D--------- ---SSD----
NCLPd        ---------K KPTTAV-KKE DSD------- ---SDSDSDS DDEEEKVAVK KEESSDSSSV
3
NCLPsp       ---------- -VVVATKKEA PKK------- -EAPKKEAPK SKAKA----- ----KKEESL
NCLPi        ---------- -------KSK AKK------- -EEEKPAAPK KKVEK----- ----KKAESS
NCLEa        ---------- -KKIETKEVK KSD------- ---SSDDSSS DDSDD----- --------SDS
NCLTn        ---------- -DSSS---EE EVK------- -AKKTKTTVK KKE------- -----KTESS
NCLTan       ---------- -QKNNKKKEE PVKKKKKEEP- -AKKKKSEGK KKEEPVKKKK PAEKKKAPPA
NCLSr        ---------- -SSDSSDSDD SSD------- ---EEEEKPK KKAEK----- -----KTEKK
NCLAg        ---------- -SSDSDSSDS DSD------- -SDSDSDEPK KKEEP----- ----KKVAKK
NCLGo        ---------- SKKKSKKKQE SSSSS----- -SSSSSSSDS SDSEEEGEKA PPKKKTAKKK
NCLChd       ---------- -K-VESSSDS SSS------- -SDSSDDEKE TKEKP----- -----KAKAK
NCLCt1       ---------- -K-AESTDSS SDS------- -SDSESEDEK PAEKK----- -----AAPAK
NCLCd        ---------- -KKSDSSSDS SSD------- -SDEEEKPKP KKAKV----- -----ESSSS
NCLDf        ---------- -EKTNDSSDS DSD------- -SESDSDKNE NKKTA----- -----AVKKK
NCLCa        ---------- -KKDDDSSSS DSD------- -SDSDEEKEK SDVKK----- -----EETAK
NCLCn        ---------- -K-DDSSSDS DSE------- -SDSDSD--- SDSDE----- -----EEKKS

NCLAsu       EG-------- -----SSSES SSSSESEKVV AKKSV----- ---------- ----------
NCLDbGSO104  SS-------- -----DSSDS DSDSSSD--- --DEEE---- ---------- ----------
4
NCLPbr       ---------- VPEYSSSSAS DSD------- -SAEDEPMPP AKEEPKKVAA ESSSDESSSS
NCLP         ---------- ---------- ---------- ---------- ---------- ----------
NCLAt1       ---------- ---------- -----TVAKK SKDDSSSSDD DSSDEE--VA VTKKPAAAAK
NCLAt2       ---------- -STSDEETAP VKKQPAVLEK AKVESSSSDD DSSSDEETVP VKKQPAVLEK
Nsr1pSc      ---------- ---------- ---------- ---SSSDSES SSSSSSDSES EAETK-----
NCLMc        ---------- -KKTTKKAKK TTKKAKKPAP VKESSSDSDS SSSEEEKPKK AAKTKKAAKA
NCLSl        ---------- --KDNSRRQS MDE------- -QKNRRQQKE QKN------- ---VKEPVTK
NCLBs        ---------- ---------- ---------- ---------- ---------- ----------
NCLCsu       TAAPPQPKTG SSHKQTPKSQ STKRLVEPEA ESESASEHSS EEENVPSKSV HVTVTKAKAA
NCLHs        ---------- --GKNAKKED SDE------- -EEDDDSEED EEDDEDEDED EDEIEPAAMK
NCLCr        ---------- --SSSSSDSG SSD------- -SDSDSDSDS DSEKKVEKKA SPSAKRAKPA
NCLDlu       GAAQTPGGAA AESSDESDGD SDD------- -SSDDSSDDD SSADENEQNA ANGAQPAKAA
5
NCLNsa       ---------- TKKKAAPESS DSD------- DSGSESDDSS EEEEAAVKPA TPARGKTTAP
NCLNg        ---------- TKKKAAPESS DSD------- DSGSESDDSS DEEEAAVKPA TPARGKTTAP
NCLTmi       ---------- ---------- ---------- ---------- ---------- ----------
NCLGt        ---------- ---------- ---------- ---------- ---------- ----------
NCLG         ---------- -PAKADKKAT PAKKAAKKEE SSSDESSSDE EDAKPAAKKA TPAKAAPAKA
6
NCLCh        ---------A PAKK--PDSD DSS------- DSDSSSDDEA APAKTSKPAA KKSAKKXAKK
NCLO         ---------A EAKKKESDSD SSS------- DSDSSSDSDS DDEKKEKPKP KAKAKAVVKK
NCLTc        ---------- --EKVSEDSD SSS------- DSDSDSDSD- SDDEKAAPAA KAAPAAAAKK
NCLTv        ---------- --AKVSDSSD SS-------- DSDDSSDSE- DEAPAAKPEV KKA-AAVAKK
NCLTs        ---------- --AKVSDSSD SS-------- DSDDSSDSE- DEAPAAKPEV KKA-AAVAKK
```

```
              ....|....| ....|....| ....|....| ....|....| ....|....| ....|....|
                 305        315        325        335        345        355
1
NCLSco1       ED-------- ---------- ---------- ---SSSSSSS --DSSDSEDE ----------
NCLSd         ED-------- ---------- ---------- ---SSSSSSS SSDSSDSEDE ----------
NCLSj         KD-------- ---------- ---------- ---SSSSSSD S-SDSDSDDE ----------
NCLSme        ED-------- ---------- ---------- ---SSSSSSD S-SDSESEDE ----------
NCLDc         EP-------- -------EKK KKE---AKKV AKKEESSSSS SSDSSDSEAE ----------
NCLTm         ESSPSSSDSD SSSDSDDEKK KKAPPAKKEV AKKDESSSSS SDSSSDEEEE ----------
NCLCm         EE-------- ---------- ---------- ---SSSDSSS SSDSSDSEDE ----------
NCLTo1        EKVV------ ---------- -----AAKKK EESSASSSSS SSDSDSSDDE ----------
NCLTw         EKKK------ ---------- -----AAKEK -------SSS SSDSSGSEDE ----------
NCLTa         SP-------- ---------- -----AKKVV AKKAESSSSD SSDSSDSDDE ----------
2
NCLFs1        KE-------- ---------- -------ESS DSSSSDSDSS SSESEDEEEE ----------
NCLEs         DS-------- ---------- ---------- ---SDDDSDD ED-KAETAKN ----------
NCLNp         DS-------- ---------- ---------- ---DDDEDEK EDSKAKAKKV ----------
NCLFk         ES-------- ---------- ---------- ---SSDSSSD DDSSSDEENE ----------
NCLPf         ---------- ---------- ---------- ---SDSDSDD DEKKQTPAAV ----------
NCLPh         SD-------- ---------- ---------- ---DDSSSDD EAEDTKAAEP ----------
NCLPa         ---------- ---------- ---------- ----DDSSDD EDEKAKAAP- ----------
NCLPd         SD-------- ---------- ---------- ---SDSDSDD EEEKAKAAP- ----------
3
NCLPsp        SS-------- ---------- ---------- ----SDSSSD SSSDSSSDS- ----------
NCLPi         SS-------- ---------- ---------- ----SSDSDS SSSDSDSD-- ----------
NCLEa         DE-------- ---------- ---------- ----EPTKKE VAKKAVAKK- ----------
NCLTn         SSDD------ ---------- -----SSDSS DD--DSSEDE DQKNSSSKEK NKAKVE----
NCLTan        KEDS------ ---------- -----SSSSS DDSSDSSSDE EETKKKAPEK TTKKTP----
NCLSr         TEKK------ ---------- -----TEKKT EK--KTEKKA DSKKKK---- ----------
NCLAg         EESS------ ---------- -----DSDSS DS--DSDSDD DKPKSKVEPK ----------
NCLGo         ED-------- ---------- ---------- ---DSSSDSD SSSDSDTSTK ----------
NCLChd        KE-------- ---------- ---------- ----ESSSSS S-DSSD---- ----------
NCLCt1        KE-------- ---------- ---------- ----DSSSSS SSDSSDSES- ----------
NCLCd         SSSS------ ---------- -----SSDSD SDDEPKKGDK AADKTTAKKT ----------
NCLDf         VESS------ ---------- -----ESSSD SDSSSDESED EKEVSKPEK- ----------
NCLCa         KV-------- ---------- ---------- MEEAKKEVKK ESDGSSSSS- ----------
NCLCn         SK-------- ---------- ---------- TEEKKVDVKK DSD-SSSDS- ----------

NCLAsu        ---------- -------AKT VAKKDE---- ---------- ---------- ------SSSS
NCLDbGSO104   ---------- ----KAKAKP KESSSSSD-- ---------- ---------- ----------
4
NCLPbr        SDDE------ ---------- ---------- KEADVSSSDD ESSSSDDEPE ----------
NCLP          ---------- ---------- ---------- ----MGLADD LLSGVAPQDR ----------
NCLAt1        NGS------- ---------- --------VK AKKESSSEDD SSSEDE---P AKKP------
NCLAt2        AKIESSSSDD DSSSDEETVP MKKQTAVLEK AKAESSSSDD GSSSDEEPTP AKKEPIVVKK
Nsr1pSc       ---------- ---------- ---------- ---KEESKDS ---------- ----------
NCLMc         KKAVKTK--- ---------- --------KA APVKEESSDS DSSDEEMEAP KKTE------
NCLSl         KNVP------ ---------- ---------- ---PPKVVDD SDDESGNESE ----------
NCLBs         ---------- ---------- ---------- ---------- ---------- ----------
NCLCsu        EAPSKQPQGK SQHR------ ---------- --YAASSDSS DGDSSDDETC VKSKKNVTSA
NCLHs         AAAA------ ---------- ---------- ---APASEDE DDEDDEDDED ----------
NCLCr         ES-------- ---------- ---------- ----SSSESS SSDSSD---- ----------
NCLDlu        VT-------- ---------- ---------- ----HDASSS SSSSSDDDDD DDDDDDD---
5
NCLNsa        SKKA------ ---------- ---------- --EPESSSSE SSDGSDEEEE ----------
NCLNg         SKKA------ ---------- ---------- --EPESSSSE SSDSSDEEEE ----------
NCLTmi        ---------- ---------- ---------- ----ESASSE ---------- ----------
NCLGt         ---------- ---------- ---------- ---------- ---------- ----------
NCLG          DKKKA----- ---------- ---------- AKKEESSSEE ESSDEDEKPA AKKATPA---
6
NCLCh         PESD------ ---------- ---------- ---SSDSDSS SD-------- ----------
NCLO          DDS------- ---------- ---------- ---ASDSDSS SDSDSDDDDE KMEDTEE---
NCLTc         EESD------ ---------- ---------- ---SDSSSSD SDSDSDSDDE KAAPAA----
NCLTv         EETD------ ---------- ---------- ---SDSSS-D SDSDSDSEDE KPA-------
NCLTs         EETD------ ---------- ---------- ---SDSSS-D SDSDSDSEDE KPA-------
```

```
                ....|....| ....|....| ....|....| ....|....| ....|....| ....|....|
                    365        375        385        395        405        415
1
NCLSco1         ---------- ---------- TXKEKKTXDA TPAETKSASK S--------- ----------
NCLSd           ---------- ---------- TPKEKKTVDA TPAETKSASK S--------- ----------
NCLSj           ---------- ---------- KPKEKKTVDV TPKETKTASK S--------- ----------
NCLSme          ---------- ---------- KPKEK-TVDA TPAETKTTSK S--------- ----------
NCLDc           ---------- ---------- APAKKAVAKK EAAKKEESSS S--------- ----------
NCLTm           ---------- ---------- KKATTSAKK- ---KKKDSSS S--------- ----------
NCLCm           ---------- ---------- APVKKESKKE TKKEEVTKEE VK-------- ----------
NCLTo1          ---------- ---------- PTEKKEEPKE QVAAKKDESS G--------- ----------
NCLTw           ---------- ---------- APKKKEETKK ---AKEDSSS GE-------- ----------
NCLTa           ---------- ---------- KAPPASAKKE AAKKSDSSSD S--------- ----------
NCLFs1          ---------- ---------- APKVASKKAE IKKKPDDSSS SS-------- ----------
2
NCLEs           ---------- -------AKV ADAKKTEEAA GDSDSDSSSS SS-------- ----------
NCLNp           ---------- -------SED KAEKEDKSDS DSSSSSSSSD SD-------- ----------
NCLFk           ---------- -------KKE VVPATKVTAK KDESSDDSSG SD-------- ----------
NCLPf           ---------- -------KKE ESDSSDSDSD --SDSDDDEK KQ-------- ----------
NCLPh           ---------- -------TKS KAVVKKDDSS SSSDSSDSSD SD-------- ----------
NCLPa           ---------- -------AKA KAAVKKDDSS D-SDSSDSSD SD-------- ----------
NCLPd           ---------- -------AKA AAAVKKEDSS --SDSSDSSD SD-------- ----------
3
NCLPsp          ---------- ---------- -------SSD SSSDSEATAP KK-------- ----------
NCLPi           ---------- ---------- ---------- -----DEKAK TK-------- ----------
NCLEa           ---------- ---------- ------ESSD SSSSSSSDSS DD-------- ----------
NCLTn           ---------- ---------- KVSKDKDSDD SSDDDDSTKS SE-------- --------AP
NCLTan          ---------- ---------- EKTKKADSSS DSDSDSSSSS SEEEKAPVSK KEKGSKRKAM
NCLSr           ---------- ---------- EVAKKESSSD SSDSDDSSSD SD-------- ----------
NCLAg           ---------- ---------- KVAKKEESSD SSSDSSDSDS SD-------- ----------
NCLGo           ---------- ---------- SSDPPPKKKE DDSSSDSDSS SD-------- ----------
NCLChd          ---------- ---------- --------DD EPK----AKT TK-------- ----------
NCLCt1          ---------- ---------- -------EDE KPAEKKPAEK AA-------- ----------
NCLCd           ---------- ---------- ---VKKESSD SSSDSSSDSE SE-------- ----------
NCLDf           ---------- ---------- ----AAESTD SSSDSDSDSD DK-------- ----------
NCLCa           ---------- ---------- -------SSD SEMEDAEXKK TE-------- ----------
NCLCn           ---------- ---------- -------SSD SSSDSDSDSD DD-------- ----------

NCLAsu          SESESDSDDD DAEPV----- ---------- ------PKTL ESKSEATSDS DEDDSEKGRK
NCLDbGSO104     SDSS-DSDSS DDEEEE---- ---------- ------KKSK AVAAKKESSD DSSSSSDS--
4
NCLPbr          ---------- ---------- PPKKVAKAAP APAKEVKASS SE-------- ----------
NCLP            ---------- ---------- ---MLSLFAP PPAEPSAPVE EE-------- ----------
NCLAt1          ---------- ---------AA KIAKPAAKDS SSSDDDSDED SEDEKPATKK AA------PAA
NCLAt2          DSSDESSSDE ETPVVKKKPT TVVKDAKAES SSSEEESSSD DEPTPAKKPT VV-----KNA
Nsr1pSc         ---------- ---------- ---------- SSSSSDSSSD EE-EEEEKEE TK-----KEE
NCLMc           ---------- ---------K KAKKPAPVKE SSSDSDSSSE EE-EKPKKTA AK-----KKA
NCLSl           ---------- ---------- DSVDLEQLIQ AKQKKTTAAL GK-------- ----------
NCLBs           ---------- ---------- ---------- ---------- ---------- ----------
NCLCsu          KVVKSAIAVK GQVSDDSDDS DDSSDEESPP RVAHKKSEGT AQQKGAILKK ACPATKPKAP
NCLHs           ---------- ---------- DDDDEEDDSE EEAMETTPAK GK-------- ----------
NCLCr           ---------- ---------- ---------- ---------- ---------- ----------
NCLDlu          ---------- ----DDDDDD DDDDDDDSES SPSSDKKAKE EE-------- ---------K
5
NCLNsa          ---------- ---------- ---EEEAAAK PKAAEESSDE SS-------- ----------
NCLNg           ---------- ---------- ---E-VVAAK PKAAEESSDE SS-------- ----------
NCLTmi          ---------- ---------- ---------- -----ESSDE ---------- ----------
NCLGt           ---------- ---------- ---------- ---------- ---------- ----------
NCLG            ---------- ----APAPAK ADKKKAAKKE ESSSEEESSD DEKPAAKKQA VG-----AKK
6
NCLCh           ---------- --------EE APPAKAAVAK KKADSDSDSD SD-------- ----------
NCLO            ---------- ----KKKEED KPKKKAAAKK ESSDSDSSSK SD-------- ----------
NCLTc           ---------- ----KAAPAA KAAPAAAAKK EESDSDSSS- SD-------- ----------
NCLTv           ---------- ------AKPEV KKAPAAAVAK KD-DSDSSS- SD-------- ----------
NCLTs           ---------- ------AKPEV KKAPAAAVAK KD-DSDSSS- SD-------- ----------
```

```
                ....|....| ....|....| ....|....| ....|....| ....|....| ....|....|
                      425        435        445        455        465        475
1
NCLSco1         ------VSS- ---------- ---------- ----GSSSSS S----DSDSD ----------
NCLSd           ------VSS- ---------- ---------- ----GSSSSS SSSSSDSDSD ----------
NCLSj           ------VSS- ---------- ---------- ----GSSSSS SSS--DSDSD ----------
NCLSme          ------VSSS ---------- ---------- ----GSSSSS SSSSSDSDSD S---------
NCLDc           ------SSDS SDSEAEAP-V KKAKAEE--- KKKD--SDSG SSSSSDSSSD S---------
NCLTm           ------SSDS SDSEEEAPAA KKTKVES--- KKDDDGSSSS SSSDSDSSSD S---------
NCLCm           ------KEVS ---------- ---------- -----DSSSD SSDSDSSDED E---------
NCLTo1          ------SSSS SDSSESEDEK PEAKKAK--- VEDVKKDESG SSSDSDSDSD SDSSSGSDSS
NCLTw           ------DSSG SDSSDSEDEK EDDKKAK--- EATKKKSESS SSSDS-SDSE ----------
NCLTa           ------DSSS DEEEKP---V AKKEEKA--- VAKKEDSSSS SSSSSGSNSD S---------
NCLFs1          ------SSDS DSDEEEEKVA PKAKAAA--- KKEESDDSSS SSSDSDSDND D---------
2
NCLEs           ------SSSS DSDSD----- -----SD--- ----DDDDST KSSNPPDDMD E---------
NCLNp           ------SDDD DDDKDAGEKD TEMKVADE-- ---KKEEDSD SSSDSDSSSD SD--------
NCLFk           ------SSDS DDDDDDDNK- VEKMDVD--- ----KKDDDD SSSDSDSDSD S---------
NCLPf           ------TPAA VKKEESDEEE ADKMEVD--- ---KKSSDSD GSS-SDSDSD S---------
NCLPh           ------SSDD EDDEKEADKK ADKMDVD--- ---KKSSDSD SSSDSDSDSD S---------
NCLPa           ------SSDD EDDKKEDKMD VEKKAAD--- ---SDSSSSD SDSDSDSDSD SD--------
NCLPd           ------SSDD EDDKKTEKMD VEKKASD--- ---SDSSGSD SDSDSDSDSD S---------
3
NCLPsp          ------AAPA PVSTKN-KKD DSDSSS---- ---DSSSDSS SDSSSDSDAD S---------
NCLPi           ------AAPA KTAAKTEKKD DSDSDS---- ---DSSSSSS SSSSSDSDDD D---------
NCLEa           ---------- -----SDSSD SDSDD----- ---SDSSDST KKSEPPANDD N---------
NCLTn           EEAAEEKPSK EGKTKAVSKD EDSSSDSD-- ---SSSDDSS SDDS--TASS E---------
NCLTan          DSDSEEKPTK KVDKKKQAAK EDSSDDDS-- ---SSSDDSS SSDSDSTKSS E---------
NCLSr           ---SDEKPKK AEAKKADTKK KDDS-DSD-- ---SDSDDSS DDDT---KSS D---------
NCLAg           ----DEEPAK K---KVEAKK DDSSSSSD-- ---S---DSD SDDS--TKSS E---------
NCLGo           ------SDTS ---------- ---------- -----TKSSD PPATMDVDDD K---------
NCLChd          ------AETK K----ASDSS SDSSS----- ---SSDSDSS SDSDSDEDTK E---------
NCLCt1          ------AEVK KDDGSDSDSS SSSSS----- ---SSDSDSS SDSDSDSDSE D---------
NCLCd           ------EEKS TPKAKEIKKE ESDS------ ---SSSSDSS SDS--DSDSD S---------
NCLDf           ------EGEK K---MEVEKE DSSSS----- ---SSDSDSS SDSDDDDDTK S---------
NCLCa           ------AAKP T-EAKAVEKE DSSS------ ---DSSSDSS SDSDSDSDSD S---------
NCLCn           ------DAKS N-DKKEIVKE AKATAAKE-- ---DSSSDSS SDS-SDSDSD S---------

NCLAsu          EDD------- ---------- ---------- ---------- ---------- ----------
NCLDbGSO104     ---SDSDSDD EV-------- ---------- ---------- ---------- --------KE
4
NCLPbr          -----DSSSS EEDDDEEPTK NGATEAKAAS SDEDDSSDES DEEESDDEES GDD-------
NCLP            ---------- ---------- ---------- --DEDEDDES EDGDADADGD G---------
NCLAt1          AKAASSSDSS DEDSDEE-SE DEKPAQKKAD T--KASKK-S SSDESS-ESE EDE-------
NCLAt2          KPAAKDSSSS EEDSDEEESD DEKPPTKKAK VSSKTSKQES SSDESSDESD KEE-------
Nsr1pSc         S---KESSSS DSSS------ ---------- ----SSSSDS ESE------- ----------
NCLMc           APAKEESSES DSSSEEESEP KKAKKTKKAA PVKESSSSES DSSDEAPKPS KAKK------
NCLSl           ------RSRG ASDVSATKKV AKKPVQAAES DDEESDGSDD EKKENDTENS PE--------
NCLBs           -----MGKYA KGSDESSGSE QEYKSSSKKY EHDSDSDDPR YKGSSSKKNK GY--------
NCLCsu          VGESTSSEES DDSDENEAAQ MQNKVVNGKT IRSARKEEQD EEAESGADET GK--------
NCLHs           ------KAAK VVPVKAKNVA EDEDEEEDDE DEDDDDDEDD EDDDDEDDEE EEEEEEEEPV
NCLCr           ------SSDG DSSDDEKAAP APAAAAAAS- ---SDSSSSS DSDSSDSDSS D---------
NCLDlu          EGGGNHASDS DSDDDAPSHK AAAKAVTAD- ---ESDGSSS SDDSSDDDSE DEP----DSS
5
NCLNsa          ------SDSE EEEEEEEK-- KAATKAE--- ----SSESGG ESSSSEEDED E---------
NCLNg           ------SDSE EEEEEEEEEE KAATKAE--- ----SSESGS ESSSSEEDED ED--------
NCLTmi          ---------- EEEEEEEE-- --APKAD--- ----TAADG- ---------- ----------
NCLGt           ---------- ---------- ---------- --MAEASKKR KEPEKAKKED G---------
NCLG            PEAAVVTKKA STKKDESSSD ESSSDEEDAK PVKKAAAKKA TPAKKAKKEE S---------
6
NCLCh           ------SSSS DTKSSEEPPK AAPAAATKPP KKAASSSDSS SSSDSDSXSD S---------
NCLO            ------SSS- DSDDDEE--K EKPKAKKEEV KKGSSSSDSS SSSDSDSDSD DD--------
NCLTc           ------SDS- DSDSDDEKAA PAAKAAPAAA AKKEESDSDS SSSDSDSDSD S---------
NCLTv           ------SDS- DSDSEDEKPA SKPEEKKAAV AKKDDSDSDS SSSDSDSDSD ----------
NCLTs           ------SDS- DSDSEDEKPA SKPEEKKAAV AKKDDSDSDS SSSDSDSDSD ----------
```

```
                 ....|....| ....|....| ....|....| ....|....| ....|....| ....|....|
                     485        495        505        515        525        535
1
NCLSco1          --DDDTKESD VP-ADDGVPS KKRKAETVLS NDN---GDKK QHTGEEG--- ----------
NCLSd            --DDDTKESD VP-ADDGVPS KKRKAETVLS NDN---GDKK QHTGEEG--- ----------
NCLSj            --DEDTKESE VPAADEGVPS KKRKAETVLS NDN---GDKK QHTGEEG--- ----------
NCLSme           --DDDTEESA VP-ADDGVPS KKRKAETVLS NDN---GDKK QHTGEEG--- ----------
NCLDc            --DDEKE-AE VP---DGVPS KKRKAEAVLE STE----DKK VAVGDD---- ----------
NCLTm            --EDETEKSD VP---VGVPT KKRKAEAVTP DAADEHADKK MHVAED---- ----------
NCLCm            -KSNDTKESE IP----ATNS KKRKSEAEDT TTD----AKR QAVSDEYVV- ----------
NCLTo1           DSEDEKEDSE EAEKTSNGDS KKRKADSVLE S-G---EDKK QSLDPE---- ----------
NCLTw            --EDETKESE APATTAATPN KKRKADTQEE TNG---DYKR QNVDSD---- ----------
NCLTa            --SSDSD-AD VP---SGVPS KKRKAEAVVA DATEEQPEKI AKVDEE---- ----------
NCLFs1           --DDDDEAEK AP----EKSS GKRKAEDIEA AAP---TPKK AAVSDEAT-- ----------
2
NCLEs            ---------- ----DEKESK KKRKAEADT- EDGFKDHSKK AAITPAE-D- ----------
NCLNp            -SDSEQTPAS IPPKETESTS KKRKEESS-- NESSEPPMKK QAFSPKE-S- ----------
NCLFk            ---------- --DED-EKAS KKRKEANDDD EQTSEPPMKK QKDS----T- ----------
NCLPf            ----DSTPAS VPPVD-DSSS KKRKAD---- DEESEPPVKK QAVT------ ----------
NCLPh            ----DATPAS VPPVE-DSAS KKRKAD---- DEPSEPATKK QAVA------ ----------
NCLPa            -SDSDSTPAS VPPVD-DSSS KKRKAD---- DEPVEPAVKK QAVG------ ----------
NCLPd            ----DSTPAS VPPVD-DSAS KKRKAD---- EETVEPAFKK QAVG------ ----------
3
NCLPsp           ------DAEK AKPP-----S KKRKAED--A PAEEPKAKVE ASSSSVNG-- ----------
NCLPi            ------TKDS DPPA-----S KKRKADD--T QDEVVHKTPK KEPEAAGG-- ----------
NCLEa            ------EAEN SGSS------ KKRKADD--A DESEENKRPA VSDDGAK--- ----------
NCLTn            ------APED VDMK--DTES KKRKGSD--- AMEPPTKKVA IDGDAN---- ----------
NCLTan           ------APVK SKES--ATPS KKRKNDDNED SEEPPTKKVA VEESGKS--- ----------
NCLSr            ------VPED DKME---EAT KKRKAGD--D DDAPPAKKAA GAEEETE--- ----------
NCLAg            ------APGD TEMENVDTTS KKRKAED--- SGEEPSKKVA VE--GEN--- ----------
NCLGo            --DFKRKTSS AT---EEPPV KKIKTTTATD GDD------- -DADNSN--- ----------
NCLChd           ------SDT- PAASK----K RKFDGVG--E EGEAVSKKAA AVADDDGA-- ----------
NCLCt1           ------TKD- SAVPE----S KKRKAED--A ESEPSAKKTA VADESE---- ----------
NCLCd            ------EDV- IDKK------ RKSGAVE--D EPAT--KKAA VVSDEG---- ----------
NCLDf            ------SDPP IEKKE----S KKRKKED--E AETEPAKKPA VEDDAV---- ----------
NCLCa            ------TKS- SDAPN----S KKRKADN--E SGNQ--SKKV AVEDGV---- ----------
NCLCn            ------TQS- SEAP-----V KKRKATD--D GGNE--SKKA AVEDGE---- ----------

NCLAsu           ---------- ---ASSSSEG SSSDEEEDVE P--------- -AKNNKRKN- --EDVSATAV
NCLDbGSO104      EKMPGAKKSS NDTTESSSVA SSSDSSSSDE DD-------- --------DE KEEEKANPKK
4
NCLPbr           EEESESESEK EDEPMPDATA KRKKPDTKAS SGGEPAKKSK VDESAKS--- ----------
NCLP             ---------- -------APD TKPKADAVLR KPRVDDAEKL SR-------- ----------
NCLAt1           ---------- -SEDEEETPK KKSSDVEMVD AE-KSSAKQP KTPSTPAAG- ----------
NCLAt2           ---------- -SKDEKVTPK KKDSDVEMVD AEQKSNAKQP KTPTNQTQG- ----------
Nsr1pSc          ---------- ----KEESND KKRKSE--DA EEEEDEESSN KKQKNEETE- ----------
NCLMc            ----SAEIES SSESEEESEE KPQKEEHQDA EMEEEKQEAP AQEEAEETE- ----------
NCLSl            ---------- ---------- ---------- -----IFVG- ---------- ----------
NCLBs            ------GSPG SSGSDSHPQK RQREPASYED KG-------- ---------- ----------
NCLCsu           ------GKTA RKRKTPSPDD GEEEPAAKKQ KESFSQPKPR VQVKGGGVV- ----------
NCLHs            KEAPGKRKKE MAKQKAAPEA KKQKVEGTEP TTAFNLFVGN LNFNKSAPEL KTGISDVFAK
NCLCr            SEDEKEEEEE KPAAK----- ----EEEAEE AVETPKPAAA TPSKGGN--- ----------
NCLDlu           SDDDDDDASD APAAQPPAAG SKRVRDEYDA SAGTPAPKRA APASSGN--- ----------
5
NCLNsa           ---------- EEEETAVAAS KKRKGGEEGG QEKKKSKGAA SPDEDR---- ----------
NCLNg            -------EDE EEEETAVAAS KKRKGGEEGG QEKKKSKGAA SPDEDR---- ----------
NCLTmi           ---------- -------ASA KKAPKAEEG- ----VSTGAG T--------- ----------
NCLGt            ----AAGD-- ---------- ---------- ---------- ---------- ----------
NCLG             ----SSDESS SSDEEDEKPA KKKAAKDSDD EEESPKKKVK VEEKEENAEE K---------
6
NCLCh            --------SSD EEDDKEKPVA KKRKVEKND- -EPTPIKKPA ETPAEEH--- ----------
NCLO             ---DSTKSSE PPAADDAPKS KKRKAEDTSP VEATPVKKAA AS--DDN--- ----------
NCLTc            --------DSD SDDEKAATSN KRKAEEEEAP VAAKASKAEA DSSPADAK-- ----------
NCLTv            ---------- --DEKE-TKA PKRKASDAAL EEAPAKKAAV SDDAGNK--- ----------
NCLTs            ---------- --DEKEETKA PKRKASDAAL EEAPAKKAAV SDDAGNK--- ----------
```

```
              ....|....| ....|....| ....|....| ....|....| ....|....| ....|....|
                   545        555        565        575        585        595
1
NCLSco1       ---------- ---------- ---------- ---------- ---------- ------GENT
NCLSd         ---------- ---------- ---------- ---------- ---------- ------GENT
NCLSj         ---------- ---------- ---------- ---------- ---------- ------GENT
NCLSme        ---------- ---------- ---------- ---------- ---------- ------GENT
NCLDc         ---------- ---------- ---------- ---------- ---------- -------ENS
NCLTm         ---------- ---------- ---------- ---------- ---------- -------EQT
NCLCm         ---------- ---------- ---------- ---------- ---------- ---LDDSENP
NCLTo1        ---------- ---------- ---------- ---------- ---------- -------ANC
NCLTw         ---------- ---------- ---------- ---------- ---------- -------DNC
NCLTa         ---------- ---------- ---------- ---------- ---------- -------DNT
NCLFs1        ---------- ---------- ---------- ---------- ---------- -------EGN
2
NCLEs         ---------- ---------- ---------- ---------- ---------- ---GSDGINT
NCLNp         ---------- ---------- ---------- ---------- ---------- ---ESGDSNT
NCLFk         ---------- ---------- ---------- ---------- ---------- ---ESEGPPP
NCLPf         ---------- ---------- ---------- ---------- ---------- ---DSDENTT
NCLPh         ---------- ---------- ---------- ---------- ---------- ---DGDEATT
NCLPa         ---------- ---------- ---------- ---------- ---------- ---GDEEGTT
NCLPd         ---------- ---------- ---------- ---------- ---------- ---GDDEG-T
3
NCLPsp        ---------- ---------- ---------- ---------- ---------- ---------H
NCLPi         ---------- ---------- ---------- ---------- ---------- --------NT
NCLEa         ---------- ---------- ---------- ---------- ---------- ---------S
NCLTn         ---------- ---------- ---------- ---------- ---------- ---------T
NCLTan        ---------- ---------- ---------- ---------- ---------- ---------V
NCLSr         ---------- ---------- ---------- ---------- ---------- ---------T
NCLAg         ---------- ---------- ---------- ---------- ---------- ---------T
NCLGo         ---------- ---------- ---------- ---------- ---------- -------DAC
NCLChd        ---------- ---------- ---------- ---------- ---------- --------GK
NCLCt1        ---------- ---------- ---------- ---------- ---------- --------NT
NCLCd         ---------- ---------- ---------- ---------- ---------- --------KL
NCLDf         ---------- ---------- ---------- ---------- ---------- --------NL
NCLCa         ---------- ---------- ---------- ---------- ---------- --------NR
NCLCn         ---------- ---------- ---------- ---------- ---------- --------NL

NCLAsu        TKVAKVANIA ATNG------ ---------E VSDDAN---- ---------- ----------
NCLDbGSO104   RQRTDDAAAE SS-------F STPAKKKSAV SNGKTK---- ---------- ----------
4
NCLPbr        ---------- ---------- ---------- ---------- ---------- ----------
NCLP          ---------- ---------- ---------- ---------- ---------- ----------
NCLAt1        ---------- ---------- ---------- ---------- ---------- -------GSK
NCLAt2        ---------- ---------- ---------- ---------- ---------- -------GSK
Nsr1pSc       ---------- ---------- ---------- ---------- ---------- -------EPA
NCLMc         ---------- ---------- ---------- ---------- ---------- -------GPT
NCLSl         ---------- ---------- ---------- ---------- ---------- ----------
NCLBs         ---------- ---------- ---------- ---------- ---------- ------DEQS
NCLCsu        ---------- ---------- ---------- ---------- ---------- -----EKMRT
NCLHs         NDLAVVDVRI GMTRKFGYVD FESAEDLEKA LELTGLKVFG NEIKLEKPKG KDSKKERDAR
NCLCr         ---------- ---------- ---------- ---------- ---------- ---GRG--EF
5
NCLDlu        ---------- ---------- ---------- ---------- ---------- ---GGGSDKP
NCLNsa        ---------- ---------- ---------- ---------- ---------- ----------
NCLNg         ---------- ---------- ---------- ---------- ---------- ----------
NCLTmi        ---------- ---------- ---------- ---------- ---------- ----------
NCLGt         ---------- ---------- ---------- ---------- ---------- --------KR
NCLG          ---------- ---------- ---------- ---------- ---------- ---AVGVENR
6
NCLCh         ---------- ---------- ---------- ---------- ---------- ----------
NCLO          ---------- ---------- ---------- ---------- ---------- ----------
NCLTc         ---------- ---------- ---------- ---------- ---------- ---------L
NCLTv         ---------- ---------- ---------- ---------- ---------- ----------
NCLTs         ---------- ---------- ---------- ---------- ---------- ----------
```

```
                          ....|....| ....|....| ....|....| ....|....| ....|....| ....|....|
                                605        615        625        635        645        655
1
NCLSco1       KIYIRGLPWR AGEDEVGTFF TSCG------ ---------- ---------- ----------
NCLSd         KIYIRGLPWR AGEDEVRDFF TSCG------ ---------- ---------- ----------
NCLSj         KIYIRGLPWR ATEEEVRDFF TSCG------ ---------- ---------- ----------
NCLSme        KVYVRGLPWR ATEEEVRDFF TSCG------ ---------- ---------- ----------
NCLDc         KIYVRGLPWR ATEDEVREFF AACG------ ---------- ---------- ----------
NCLTm         KIYIRGLPWR ATEEEVRDFF ASCG------ ---------- ---------- ----------
NCLCm         KVYVRGLPWK ATYDEIKEFF GGCG------ ---------- ---------- ----------
NCLTo1        KVYIRGLPWR ATEDEVREFF AECG------ ---------- ---------- ----------
NCLTw         KIYVRGLPWK AGEDDVKEFF KACG------ ---------- ---------- ----------
NCLTa         KIYVRGLPWR TSDEEVREYF AACG------ ---------- ---------- ----------
NCLFs1        KVYVRGLPWK AEEHEVQEFF AACG------ ---------- ---------- ----------
2
NCLEs         KIFVRGLPWK TTEEEVTDFF KDCG------ ---------- ---------- ----------
NCLNp         KIYVRGLPWR ATKDQVEVFF QSCG------ ---------- ---------- ----------
NCLFk         KLYIRGLPWR ADEEQVRDFF KSCG------ ---------- ---------- ----------
NCLPf         KLYIRGLPWR TTEEAVKEYF QTCG------ ---------- ---------- ----------
NCLPh         KLYIRGLPWR ATEEAVKEFF QSCG------ ---------- ---------- ----------
NCLPa         KLYIRGLPWR ATEDAVRDYF KACG------ ---------- ---------- ----------
NCLPd         KLYIRGLPWR ATEDGVRDFF KACG------ ---------- ---------- ----------
3
NCLPsp        KVFVRGFPWA TTENEINDFF QSCG------ ---------- ---------- ----------
NCLPi         KIFIRGLPWR ATEEEVMDFF KDCGDGPTSI EMPLQDDGRS SGTAIMDFAN ATGFDAGLAL
NCLEa         KIFIRGLPWR ASEDEIREHF GGCGE----- ---------- ---------- ----------
NCLTn         KIYIRGLPWR ATEDELRDFF QTCGT----- ---------- ---------- ----------
NCLTan        KVFVRGLPWR ASEDEVWEFF GECGK----- ---------- ---------- ----------
NCLSr         KVYVRGLPWR ASEEEIREWF VSCGT----- ---------- ---------- ----------
NCLAg         KIYIRGLPWR ATEDEVREFF QGCGKG---- ---------- ---------- ----------
NCLGo         KLYVKGLPWK ATYDDVHGYF SQSSTKA--- ---------- ---------- ----------
NCLChd        KMYIRGLPWR ATDDELREYF ASCGE----- ---------- ---------- ----------
NCLCt1        TMYIRGLPWR ATEEEVRAFF DSCGE----- ---------- ---------- ----------
NCLCd         KMFVRGLPWR ASEDDVRNYF TECGE----- ---------- ---------- ----------
NCLDf         KMFVRGLPWR ATEEEVREYF ATCGE----- ---------- ---------- ----------
NCLCa         KMYVRGLPWK ATEDEVRDYF ASCGE----- ---------- ---------- ----------
NCLCn         TLFIRGLPWR ATEDDLREYF KSCGE----- ---------- ---------- ----------
NCLAsu        ---------- -CKIFVRGLP WAATEDEVED FFKGCG---- ---------- ----------
4
NCLDbGSO104   ---------- ---VGVWGLP YSVTEEEIKD FFKDCG---- ---------- ----------
NCLPbr        SLFIGGLSYE ATEDDLRDLF AECG------ ---------- ---------- ----------
NCLP          TVFIGNVPVS TTQKVLKRLF TECGQIDSIR IRSAAFAN-- ---------- ----------
NCLAt1        TLFAANLSFN IERADVENFF KEAG------ ---------- ---------- ----------
NCLAt2        TLFAGNLSYQ IARSDIENFF KEAG------ ---------- ---------- ----------
Nsr1pSc       TIFVGRLSWS IDDEWLKKEF EHIG------ ---------- ---------- ----------
NCLMc         ELFVGNLSWN VDDAYLKETF EYYG------ ---------- ---------- ----------
NCLSl         -----NIPFT MENDALKELF EEFG------ ---------- ---------- ----------
NCLBs         EVFVGGLSWE ADDSDLQKFF KN-CG----- ---------- ---------- ----------
NCLCsu        EIFCGGLPYA TTESELKELF EADCG----- ---------- ---------- ----------
NCLHs         TLLAKNLPYK VTQDELKEVF EDAA------ ---------- ---------- ----------
NCLCr         AVTVRGLPYE MDEDSILELF ATSG------ ---------- ---------- ----------
NCLDlu        TAMLTGLPHE VTDAAIRAFL APAG------ ---------- ---------- ----------
5
NCLNsa        KVFVQGLPWS ATEEEVRAFF EKCG------ ---------- ---------- ----------
NCLNg         KVFVQGLPWS ATEEEVRAFF QKCG------ ---------- ---------- ----------
NCLTmi        KIYVRGLPWS ATEEEVRDFF KECG------ ---------- ---------- ----------
NCLGt         RVFLGGLPFK ATEKDIKKMF ESCG------ ---------- ---------- ----------
NCLG          RVFLGGLPFR ATEDEIKDMF KSCG------ ---------- ---------- ----------
6
NCLCh         KIIAQGIPWA CTEEDVQQFF ADCG------ ---------- ---------- ----------
NCLO          VVIAKGLPWK ATPEEVTDFF KDCG------ ---------- ---------- ----------
NCLTc         KLYVRGLTWT AGEADVRDFF KDCG------ ---------- ---------- ----------
NCLTv         KLYVRGLPWK ATEDEVRDFF KECG------ ---------- ---------- ----------
NCLTs         KLYVRGLPWK ATEDEVRDFF KECG------ ---------- ---------- ----------
```

```
               ....|....| ....|....| ....|....| ....|....| ....|....| ....|....|
                     665        675        685        695        705        715
1
NCLSco1        ---------- ---------- ---------- ---------- ---------- ----------
NCLSd          ---------- ---------- ---------- ---------- ---------- ----------
NCLSj          ---------- ---------- ---------- ---------- ---------- ----------
NCLSme         ---------- ---------- ---------- ---------- ---------- ----------
NCLDc          ---------- ---------- ---------- ---------- ---------- ----------
NCLTm          ---------- ---------- ---------- ---------- ---------- ----------
NCLCm          ---------- ---------- ---------- ---------- ---------- ----------
NCLTo1         ---------- ---------- ---------- ---------- ---------- ----------
NCLTw          ---------- ---------- ---------- ---------- ---------- ----------
NCLTa          ---------- ---------- ---------- ---------- ---------- ----------
NCLFs1         ---------- ---------- ---------- ---------- ---------- ----------
2
NCLEs          ---------- ---------- ---------- ---------- ---------- ----------
NCLNp          ---------- ---------- ---------- ---------- ---------- ----------
NCLFk          ---------- ---------- ---------- ---------- ---------- ----------
NCLPf          ---------- ---------- ---------- ---------- ---------- ----------
NCLPh          ---------- ---------- ---------- ---------- ---------- ----------
NCLPa          ---------- ---------- ---------- ---------- ---------- ----------
NCLPd          ---------- ---------- ---------- ---------- ---------- ----------
3
NCLPsp         ---------- ---------- ---------- ---------- ---------- ----------
NCLPi          NGETFGERWL SIKAERVSFL VILNLSCCFD FYAFRETEVL PSFFXANFKH FCLTYMIIYV
NCLEa          ---------- ---------- ---------- ---------- ---------- ----------
NCLTn          ---------- ---------- ---------- ---------- ---------- ----------
NCLTan         ---------- ---------- ---------- ---------- ---------- ----------
NCLSr          ---------- ---------- ---------- ---------- ---------- ----------
NCLAg          ---------- ---------- ---------- ---------- ---------- ----------
NCLGo          ---------- ---------- ---------- ---------- ---------- ----------
NCLChd         ---------- ---------- ---------- ---------- ---------- ----------
NCLCt1         ---------- ---------- ---------- ---------- ---------- ----------
NCLCd          ---------- ---------- ---------- ---------- ---------- ----------
NCLDf          ---------- ---------- ---------- ---------- ---------- ----------
NCLCa          ---------- ---------- ---------- ---------- ---------- ----------
NCLCn          ---------- ---------- ---------- ---------- ---------- ----------
NCLAsu         ---------- ---------- ---------- ---------- ---------- ----------
NCLDbGSO104    ---------- ---------- ---------- ---------- ---------- ----------
4
NCLPbr         ---------- ---------- ---------- ---------- ---------- ----------
NCLP           ---------- ---------- ---------- ---------- ---------- ----------
NCLAt1         ---------- ---------- ---------- ---------- ---------- ----------
NCLAt2         ---------- ---------- ---------- ---------- ---------- ----------
Nsr1pSc        ---------- ---------- ---------- ---------- ---------- ----------
NCLMc          ---------- ---------- ---------- ---------- ---------- ----------
NCLSl          ---------- ---------- ---------- ---------- ---------- ----------
NCLBs          ---------- ---------- ---------- ---------- ---------- ----------
NCLCsu         ---------- ---------- ---------- ---------- ---------- ----------
NCLHs          ---------- ---------- ---------- ---------- ---------- ----------
NCLCr          ---------- ---------- ---------- ---------- ---------- ----------
NCLDlu         ---------- ---------- ---------- ---------- ---------- ----------
5
NCLNsa         ---------- ---------- ---------- ---------- ---------- ----------
NCLNg          ---------- ---------- ---------- ---------- ---------- ----------
NCLTmi         ---------- ---------- ---------- ---------- ---------- ----------
NCLGt          ---------- ---------- ---------- ---------- ---------- ----------
NCLG           ---------- ---------- ---------- ---------- ---------- ----------
6
NCLCh          ---------- ---------- ---------- ---------- ---------- ----------
NCLO           ---------- ---------- ---------- ---------- ---------- ----------
NCLTc          ---------- ---------- ---------- ---------- ---------- ----------
NCLTv          ---------- ---------- ---------- ---------- ---------- ----------
NCLTs          ---------- ---------- ---------- ---------- ---------- ----------
```

```
               ....|....| ....|....| ....|....| ....|....| ....|....| ....|....|
                     725        735        745        755        765        775
1
NCLSco1        ---------- ---------- ---------- ---------- ---------- ----------
NCLSd          ---------- ---------- ---------- ---------- ---------- ----------
NCLSj          ---------- ---------- ---------- ---------- ---------- ----------
NCLSme         ---------- ---------- ---------- ---------- ---------- ----------
NCLDc          ---------- ---------- ---------- ---------- ---------- ----------
NCLTm          ---------- ---------- ---------- ---------- ---------- ----------
NCLCm          ---------- ---------- ---------- ---------- ---------- ----------
NCLTo1         ---------- ---------- ---------- ---------- ---------- ----------
NCLTw          ---------- ---------- ---------- ---------- ---------- ----------
NCLTa          ---------- ---------- ---------- ---------- ---------- ----------
NCLFs1         ---------- ---------- ---------- ---------- ---------- ----------
2
NCLEs          ---------- ---------- ---------- ---------- ---------- ----------
NCLNp          ---------- ---------- ---------- ---------- ---------- ----------
NCLFk          ---------- ---------- ---------- ---------- ---------- ----------
NCLPf          ---------- ---------- ---------- ---------- ---------- ----------
NCLPh          ---------- ---------- ---------- ---------- ---------- ----------
NCLPa          ---------- ---------- ---------- ---------- ---------- ----------
NCLPd          ---------- ---------- ---------- ---------- ---------- ----------
3
NCLPsp         ---------- EIES------ ---------- ---------- ---------- ----------
NCLPi          AHNSQNNFTP ALPNXTFSLK PFLFHKTLTM DKTINIPISA XNHFFNCFQP LWILLMPLDL
NCLEa          ---------- ---------- ---------- ---------- ---------- ----------
NCLTn          ---------- ---------- ---------- ---------- ---------- ----------
NCLTan         ---------- ---------- ---------- ---------- ---------- ----------
NCLSr          ---------- ---------- ---------- ---------- ---------- ----------
NCLAg          ---------- ---------- ---------- ---------- ---------- ----------
NCLGo          ---------- ---------- ---------- ---------- ---------- ----------
NCLChd         ---------- ---------- ---------- ---------- ---------- ----------
NCLCt1         ---------- ---------- ---------- ---------- ---------- ----------
NCLCd          ---------- ---------- ---------- ---------- ---------- ----------
NCLDf          ---------- ---------- ---------- ---------- ---------- ----------
NCLCa          ---------- ---------- ---------- ---------- ---------- ----------
NCLCn          ---------- ---------- ---------- ---------- ---------- ----------
NCLAsu         ---------- ---------- --EIVK---- ---------- ---------- ----------
NCLDbGSO104    ---------- ---------- --TIQS---- ---------- ---------- ----------
4
NCLPbr         ---------- ---------- ---------- ---------- ---------- ----------
NCLP           ---------- ---------- ---------- ---------- ---------- ----------
NCLAt1         ---------- ---------- ---------- ---------- ---------- ----------
NCLAt2         ---------- ---------- ---------- ---------- ---------- ----------
Nsr1pSc        ---------- ---------- ---------- ---------- ---------- ----------
NCLMc          ---------- ---------- ---------- ---------- ---------- ----------
NCLSl          ---------- ---------- ---------- ---------- ---------- ----------
NCLBs          ---------- ---------- ---------- ---------- ---------- ----------
NCLCsu         ---------- ---------- ---------- ---------- ---------- ----------
NCLHs          ---------- ---------- ---------- ---------- ---------- ----------
NCLCr          ---------- ---------- ---------- ---------- ---------- ----------
NCLDlu         ---------- ---------- ---------- ---------- ---------- ----------
5
NCLNsa         ---------- ---------- ---------- ---------- ---------- ----------
NCLNg          ---------- ---------- ---------- ---------- ---------- ----------
NCLTmi         ---------- ---------- ---------- ---------- ---------- ----------
NCLGt          ---------- ---------- ---------- ---------- ---------- ----------
NCLG           ---------- ---------- ---------- ---------- ---------- ----------
6
NCLCh          ---------- ---------- ---------- ---------- ---------- ----------
NCLO           ---------- ---------- ---------- ---------- ---------- ----------
NCLTc          ---------- ---------- ---------- ---------- ---------- ----------
NCLTv          ---------- ---------- ---------- ---------- ---------- ----------
NCLTs          ---------- ---------- ---------- ---------- ---------- ----------
```

```
                 ....|....| ....|....| ....|....| ....|....| ....|....| ....|....|
                       785        795        805        815        825        835
1
NCLSco1          ---------- -------EMT S-VELPLQDD GRX-SGTAII QFSDAAGAAA AMEH-NGADF
NCLSd            ---------- -------EMT S-VELPLQDD GRS-SGTAII QFSDAAGAAA AMEH-NGADF
NCLSj            ---------- -------EMT S-VELPLQDD GRS-SGTAII AFSNAASAAA AMEH-NGADF
NCLSme           ---------- -------EMA S-VEMPLQDD GRS-SGTAII AFSDAAGAAA AMEH-NGADF
NCLDc            ---------- -------EIT T-LELPMQDD GRS-SGTAII DFKESAGAAA AMEQ-NGADF
NCLTm            ---------- -------EIE S-CELPLMDD GRS-SGTAII AFKEATAAAA ALEH-NGADF
NCLCm            ---------- -------KIK S-VDLPLLAD GRS-SGTAVI EFESPAGSAA AMEQ-NGADF
NCLTo1           ---------- -------EIK S-VDMPLQDD GRS-SGTAII EFSDPSGSAS ALEH-NGADF
NCLTw            ---------- -------DIV N-IELPLMND GRS-SGTAII EFKDSTGAAS ALEH-NGADF
NCLTa            ---------- -------EVT S-VELPLQDD GRS-SGTAII DFKDNAGSAA AFEL-NGADY
NCLFs1           ---------- -------PIK S-CEMPLMDD GRS-SGTAII EFETNEGAAA AIEL-NGGDF
2
NCLEs            ---------- -------NGP KVVELPLQDD GRS-SGTAII DFGSTEGAEA AIAL-NGATY
NCLNp            ---------- -------SGP RSVDLPLQDD GRS-SGTAII EFEDAESAAA AIEL-NGADF
NCLFk            ---------- -------SGP TTIELPLQDD GRS-SGTAVV EFKDTASAEA AIEL-NGADF
NCLPf            ---------- -------SGP TFVELPLQPD GRS-SGTAII DFADAASAAA AMEL-NGADF
NCLPh            ---------- -------SGP KSVELPLQDD GRS-SGTAIV DFHDSSSAAA ALEL-NGADF
NCLPa            ---------- -------SGP KSVELPLQED GRS-SGTAII DFHDAASAAA ALEL-NGADF
NCLPd            ---------- -------SGP KSVELPLQED GRS-SGTAIL DFHDGASAAA AMEL-NGADF
3
NCLPsp           ---------- ---------- --IEQPLGDD GRA-SGTAII AFKTQEGANK SYEL-DRATF
NCLPi            MRRATEEEVM DFFKDCGDGP TSIEMPLQDD GRS-SGTAIM DFANATGFDA GLAL-NGETF
NCLEa            ---------- ---------I KEVEQPMNSD GRS-SGTALI VFGSASSAES AVEM-NGGDF
NCLTn            ---------- ---------I VSCELPLQDD GRS-SGTAVL EFESAESAAK GIEL-NGEDF
NCLTan           ---------- ---------I VTCELPLQDD GRS-SGTALV EFGSDAEAAK AIEM-NGADF
NCLSr            ---------- ---------I TNCELPLQDD GRS-SGTAIL DFDSADAAVK AIEL-NGEDF
NCLAg            ---------- ---------P TNLELPLQDD GRS-SGTAII DFGSAEDAAA AMEL-NGADF
NCLGo            ---------- -------KVV S-CELPVQDD GRS-SGTAIL EFESPKEASA VMEECQGADF
NCLChd           ---------- ---------I VSCELPLQSD GRS-SGTAEI EFSTKDACEK CMAL-NGEDF
NCLCt1           ---------- ---------M VSCELPLQDD GRS-SGTAII KFSDTAGAEA CLAL-NGEDF
NCLCd            ---------- ---------M KTCELPLNHD GRS-SGTAII VFATKDACEA CIAL-NGADF
NCLDf            ---------- ---------I ESLELPLQDD GRS-SGTAII VFKTKEACEA CLAC-NGEDF
NCLCa            ---------- ---------I TLLELPLMDD GRS-SGTAVI EYATKDACEA CLKL-NGEDF
NCLCn            ---------- ---------I TSLELPLQDD GRS-SGTAVM KFSSKKECEA CLQL-DGEDF
NCLAsu           ---------- ---------- ---------- ----IDLPLS EDGRASGTAF VEFKTTSGSA
NCLDbGSO104      ---------- ---------- ---------- ----IEF-QE KLGRFSGRAI VEFDSSDACE
4
NCLPbr           ---------- --------TI SSIRIPVFED SGKPRGIAFI EFEEGDAAKK GLAK-DGAEI
NCLP             ---------- ----KKISRK AAVISKAFDK DRRDTYNAYI VFVAPEGAQA ALGK-NGTVV
NCLAt1           ---------- --------EV VDVRFSTNRD DGSFRGFGHV EFASSEEAQK ALEFHG-RPL
NCLAt2           ---------- --------EV VDVRLSS-FD DGSFKGYGHI EFASPEEAQK ALEMNG-KLL
Nsr1pSc          ---------- --------GV IGARVIYERG TDRSRGYGYV DFENKSYAEK AIQEMQGKEI
NCLMc            ---------- --------EI KRANVLIRDG --RSQGIGFV EFGNHAQAKA ALEQANEYEL
NCLSl            ---------- ---------- TIENVSVPMS GRKKKGYAFI RFSSHEEAAA AVKGMNNKEI
NCLBs            ---------- ---------S ITNIKILKND QGKSKGSAFI KFSSPEEAQE AIRL-NGSEH
NCLCsu           ---------- ---------P TTRIKMLE-- ---GKGIAFI TFQTEEGAQK AVEY-NNTQY
NCLHs            ---------- ---------- EIRLVSK--- DGKSKGIAYI EFKTEADAEK TFEEKQGTEI
NCLCr            ---------- -------EVV RVEVSRFEDT GRP-KGIANV VFATEEGKDN AIAR-DGETC
NCLDlu           ---------- -------CGP VEIFMPGEST GQTGRGVAFV SVSSTDALAK LIAH-SGKQL
5
NCLNsa           ---------- --------PM TSVELPLKD- GRS-SGTAYI VFGEEDGVGK AVAM-NEEIF
NCLNg            ---------- --------PM TSVELPLKD- GRS-SGTAYI VFGEEDGVGK AVAM-NEEIF
NCLTmi           ---------- --------EI ESCDLPTDHN GRA-SGTAFV VFADSDAANK S---------
NCLGt            ---------- --------AI ENIELPMNAD SRP-AGFGFL TFKDADSVAK AVAM-DGQEL
NCLG             ---------- --------QV ETLELPMNNE GRP-SGFGFL TFKSAAAVAK AVAM-DGQEL
6
NCLCh            ---------- -------EIL S-CKIPMQDD GRS-SGRAFV VFSTDEALQA AIAM-DGQTM
NCLO             ---------- -------SGP SDVDLPLDYS GRS-SGTAIL KFDSSDGVNA ALAL-NGETF
NCLTc            ---------- -------EMV A-CELPLSDD GRS-SGTAFI TMKDQAGVDA ALAL-DHADF
NCLTv            ---------- -------EIE S-CELPLDDT GRS-SGTCFL VFKDTAGAEQ GLAL-DQATF
NCLTs            ---------- -------EIE S-CELPLDDT GRS-SGTCFL VFKDTAGAEQ GLAL-DQATF
```

```
             ....|....| ....|....| ....|....| ....|....| ....|....| ....|....|
                    845        855        865        875        885        895
1
NCLSco1      NG--RWLXIK ---------- ---------- ---------- ---------- ---------Y
NCLSd        NG--RWLNIK ---------- ---------- ---------- ---------- ---------Y
NCLSj        NG--RWLNIK ---------- ---------- ---------- ---------- ---------Y
NCLSme       GG--RWLNIK ---------- ---------- ---------- ---------- ---------Y
NCLDc        GG--RWLSIK ---------- ---------- ---------- ---------- ---------Y
NCLTm        GG--RWLNIK ---------- ---------- ---------- ---------- ---------Y
NCLCm        GG--RWLNIK ---------- ---------- ---------- ---------- ---------Y
NCLTo1       GG--RWLNIK ---------- ---------- ---------- ---------- ---------Y
NCLTw        QG--RWLNIK ---------- ---------- ---------- ---------- ---------Y
NCLTa        GG--RWLSIK ---------- ---------- ---------- ---------- ---------Y
NCLFs1       QG--RWLSIK ---------- ---------- ---------- ---------- ---------Y
2
NCLEs        GDSGRWLNIK ---------- ---------- ---------- ---------- ---------Y
NCLNp        --DGRWLSIK ---------- ---------- ---------- ---------- ---------Y
NCLFk        --EGRWLSIK ---------- ---------- ---------- ---------- ---------Y
NCLPf        --EGRWLSIK ---------- ---------- ---------- ---------- ---------Y
NCLPh        --EGRWLSIK ---------- ---------- ---------- ---------- ---------Y
NCLPa        --EGRWLSIK ---------- ---------- ---------- ---------- ---------Y
NCLPd        --EGRWLSIK ---------- ---------- ---------- ---------- ---------Y
3
NCLPsp       GE--RWLSVK EN-------- ---------- ---------- ---------- ----------
NCLPi        GE--RWLSIK AN-------- ---------- ---------- ---------- ----------
NCLEa        GG--RWLDIK LA-------- ---------- ---------- ---------- ----------
NCLTn        QG--RWLSIK YS-------- ---------- ---------- ---------- ----------
NCLTan       QG--RWLDIK LDG------- ---------- ---------- ---------- ----------
NCLSr        QG--RWLSIK YS-------- ---------- ---------- ---------- ----------
NCLAg        NG--RWLSIK YS-------- ---------- ---------- ---------- ----------
NCLGo        DG--RWLNIS ---------- ---------- ---------- ---------- ---------L
NCLChd       GG--RWLSIK YA-------- ---------- ---------- ---------- ----------
NCLCt1       NG--RWLNIK YS-------- ---------- ---------- ---------- ----------
NCLCd        EG--RWLNII YS-------- ---------- ---------- ---------- ----------
NCLDf        NG--RWLNIK YS-------- ---------- ---------- ---------- ----------
NCLCa        NG--RWLSIK YS-------- ---------- ---------- ---------- ----------
NCLCn        DG--RWLSIK YS-------- ---------- ---------- ---------- ----------
NCLAsu       AAIEL-NGNT FG--ERWLSI KYDTPKQQGG NF-------- ---------- ----------
NCLDbGSO104  AAIKL-NKMT FFDSGRWVGV EDADAPAQTR TPT------- ---------- ----------
4
NCLPbr       RG--RWIKVQ EAS------- ---------- ---------- ---------- ----------
NCLP         DEKTLRVDKA TAPRVG---- ---------- ---------- ---------- ---------Q
NCLAt1       LGREIRLDIA QERGERGERP AF-------- ---------- ---------- -------TPQ
NCLAt2       LGRDVRLDLA NERG------ ---------- ---------- ---------- -------TPR
Nsr1pSc      DGRPINCDMS TSKPA----- ---------- ---------- ---------- ----------
NCLMc        DGRPMRVSFS ADKP------ ---------- ---------- ---------- ----------
NCLSl        EGRELKCNFS SGKV------ ---------- ---------- ---------- ----------
NCLBs        MGRTLRINLS GDKPNK---- ---------- ---------- ---------- --------HR
NCLCsu       NGRTLRINLT ADKQQNQQTR AGDR------ ---------- ---------- -------GER
NCLHs        DGRSISLYYT GEKG------ ---------- ---------- ---------- ----------
NCLCr        GA--RWLSIG EK-------- ---------- ---------- ---------- ----------
NCLDlu       GG--RAVGIR EA-------- ---------- ---------- ---------- ----------
5
NCLNsa       GDSGRWVRVR KYEP------ ---------- ---------- ---------- ---------V
NCLNg        GDSGRWVRVR KYEP------ ---------- ---------- ---------- ---------T
NCLTmi       --RERWLKIV LATE------ ---------- ---------- ---------- ---------R
NCLGt        MG--RWVKVK EAD------- ---------- ---------- ---------- ----------
NCLG         QG--RWIKVK EAD------- ---------- ---------- ---------- ----------
6
NCLCh        QE--RWIGVR ---------- ---------- ---------- ---------- --------KW
NCLO         GDSGRWLKIE KYDCKPQAQA VSPKPEGCTR IFVGNLSFDI DEDTIKNFFA QCGTVTDVKW
NCLTc        QG--RWLSIK ---------- ---------- ---------- ---------- ---------L
NCLTv        GE--RWLSVK ---------- ---------- ---------- ---------- ---------Y
NCLTs        GE--RWLSVK ---------- ---------- ---------- ---------- ---------Y
```

```
                ....|....| ....|....| ....|....| ....|....| ....|....| ....|....|
                      905        915        925        935        945        955
1
NCLSco1         SXXKPVT--- ---------- ---------- ---------- ---------- -------AXR
NCLSd           STSKPVT--- ---------- ---------- ---------- ---------- -------AAR
NCLSj           STSKPVT--- ---------- ---------- ---------- ---------- -------AAR
NCLSme          SSSKPIT--- ---------- ---------- ---------- ---------- -------AAR
NCLDc           SSSKPIG--- ---------- ---------- ---------- ---------- -------EVR
NCLTm           SSSKPIS--- ---------- ---------- ---------- ---------- -------APR
NCLCm           SSNKPVT--- ---------- ---------- ---------- ---------- -------APR
NCLTo1          STSRPIT--- ---------- ---------- ---------- ---------- -------EAR
NCLTw           STSKSIT--- ---------- ---------- ---------- ---------- -------APR
NCLTa           SSNKPIN--- ---------- ---------- ---------- ---------- -------EAR
NCLFs1          STPKPIL--- ---------- ---------- ---------- ---------- -------SAR
2
NCLEs           STSKPIT--- ---------- ---------- ---------- ---------- -------APR
NCLNp           STPKPIL--- ---------- ---------- ---------- ---------- -------GAR
NCLFk           STPKSII--- ---------- ---------- ---------- ---------- -------APR
NCLPf           STPKPIL--- ---------- ---------- ---------- ---------- -------AAR
NCLPh           SSPKPIL--- ---------- ---------- ---------- ---------- -------AAR
NCLPa           STPKPIL--- ---------- ---------- ---------- ---------- -------AAR
NCLPd           STPKPIL--- ---------- ---------- ---------- ---------- -------AAR
3
NCLPsp          -SEKPK---- ---------- ---------- ---------- ---------- -------FER
NCLPi           -GDKPVH--- ---------- ---------- ---------- ---------- -------TPR
NCLEa           -DDKPIQ--- ---------- ---------- ---------- ---------- -------TER
NCLTn           -SAKPIT--- ---------- ---------- ---------- ---------- -------APR
NCLTan          -MQKPAA--- ---------- ---------- ---------- ---------- -------GAR
NCLSr           -SPKPIT--- ---------- ---------- ---------- ---------- -------SGR
NCLAg           -TPKPIL--- ---------- ---------- ---------- ---------- -------AAR
NCLGo           YTQRKPSRM- ---------- ---------- ---------- ---------- -------GAY
NCLChd          -SDKPAF--- ---------- ---------- ---------- ---------- -------AAR
NCLCt1          -NSKPIH--- ---------- ---------- ---------- ---------- -------SPR
NCLCd           -NDKPIT--- ---------- ---------- ---------- ---------- -------APR
NCLDf           -TPKPIF--- ---------- ---------- ---------- ---------- -------AKR
NCLCa           -SPKPIN--- ---------- ---------- ---------- ---------- -------TPR
NCLCn           -TPKPIN--- ---------- ---------- ---------- ---------- -------APR
NCLAsu          ---------- ---------- ---------- ---------- ---------- ---SPRPVSK
NCLDbGSO104     ---------- ---------- ---------- ---------- ---------- ---------A
4
NCLPbr          AKREPQK--- ---------- ---------- ---------- ---------- --------QR
NCLP            GPGKAVPGGP E--------- ---------- ---------- ---------- -------GEQ
NCLAt1          SGNFRSG--- ---------- ---------- ---------- ---------- ---------G
NCLAt2          NSNPGRK--- ---------- ---------- ---------- ---------- ---------G
Nsr1pSc         GNNDRAK--- ---------- ---------- ---------- ---------- ---------K
NCLMc           --NKHEK--- ---------- ---------- ---------- ---------- ---------R
NCLSl           VEKKPRG--- ---------- ---------- ---------- ---------- ------EGQE
NCLBs           ESGGSGRG-- ---------- ---------- ---------- ---------- ----------
NCLCsu          GGGRGGRGGG K--------- ---------- ---------- ---------- ----NTGERD
NCLHs           QNQDYRG--- ---------- ---------- ---------- ---------- ------GKN-
NCLCr           -RLREEA--- ---------- ---------- ---------- ---------- -------EAS
NCLDlu          -VSADGG--- ---------- ---------- ---------- ---------- -------AVT
5
NCLNsa          AAGKDGGKGM G--------- ---------- ---------- ---------- -------FAR
NCLNg           AAGKDGGKGM G--------- ---------- ---------- ---------- -------FAR
NCLTmi          AARPSFG--- ---------- ---------- ---------- ---------- --------AP
NCLGt           GTEGSAGKK- ---------- ---------- ---------- ---------- ----PFTPNR
NCLG            GEEKNKAPG- ---------- ---------- ---------- ---------- ----RFG---
6
NCLCh           EPPRAS---- ---------- ---------- ---------- ---------- ----------
NCLO            AEDRDTGRFR GFGHLEFEES DATDKAVAMA GSDLLGRPIK IDYAKAQARK SFGGGTDRPF
NCLTc           STEKPKTFS- ---------- ---------- ---------- ---------- ----------
NCLTv           ATDKPQGFNK ---------- ---------- ---------- ---------- ----------
NCLTs           ATDKPQGFNK ---------- ---------- ---------- ---------- ----------
```

```
                 ....|....| ....|....| ....|....| ....|....| ....|....| ....|....|
                    965        975        985        995        1005       1015
1
NCLSco1         EPSQKXEGCM TVFVGXLSFH XDEETVRETF KD-XGEITAV RFAEDK-ETG QFKGFGHVEF
NCLSd           EPSQKEEGCM TVFVGNLSFH IDEETVRETF KD-CGEITAV RFAEDK-ETG QFKGFGHVEF
NCLSj           EPSQKEEGCM TVFVGNLSFH IDEETVRDTF KD-CGEITAV RFAEDK-ETG QFKGFGHVEF
NCLSme          EASQKEEGCM TVFVGNLSFH IDEDTIRDTF KD-CGEITAV RFAEDK-ETG QFKGFGHVEF
NCLDc           APSQKDEGCM TVFVGNLSFH IDEDTVRETF KD-CGEIASI RFAEDR-ETG QFKGFGHVEF
NCLTm           APSQKEDGCV TVFVGNLSFN IDEETLRDAF KD-CGEIASV RFAEDR-ETG QFKGFGHVEF
NCLCm           EPSHKEEGCT TVFVGNLSFA IDEDTLREAF AS-CGEIASV RFAEDR-ETG AFKGFGHIEF
NCLTo1          APSQKEEGCV TVFVGNMSFH IDEDTVRETF KD-CGEIASI RFAEDR-ETG AFKGYGHVEF
NCLTw           EPSKKDPGCV TVFVGNLSFD IDEDALRDAF KD-CGEICSI RFAEDR-ETG AFKGFGHIEF
NCLTa           PVSQKEEGCM TVFVGNLSFN IEEDALREAF QE-CGEITNI RFATDR-ETG DFKGFGHVEF
NCLFs1          EPTQKQDGCV TVFVGNLAWD VDEDTLRQAF GE-CGEITSV RFATDR-ETG EFKGFGHIEF
2
NCLEs           GPSEKQEGCV TVFIGNLDWN IDEETIRATF GD-CGEITQV RFAEDR-ESG QFKGFGHIEF
NCLNp           EPTMKEEGCV TVFVGNLSWD IDEDSLRDAF KD-CGTITQV RFSTDR-ETG DFKGYGHIEF
NCLFk           AASEKQEGCM TVFVGNLSWD VDEDTLRDAF KD-CGTITQV RFSTDR-ETG DFKGYGHIEF
NCLPf           EASQKQEGCC TVFVGNLSWD IDEDSLRAAF AD-CGTINQI RFSTDR-ETG DFKGYGHIEF
NCLPh           QVSEKQEGCC TVFVGNLSWD IDEDSLRAAF AD-CGTITQV RFSTDR-ETG DFKGYGHIEF
NCLPa           EVTQKQEGCC TVFVGNLSWD IDEETLRAAF AD-CGTITQV RFSTDR-ETG DFKGYGHIEF
NCLPd           EVSQKQEGCC TVFVGNLSWD IDEETLRAAF AD-CGTITQI RFSTDR-ETG DFKGYGHIEF
3
NCLPsp          EVSPKMDGCM TVFMGNLSWD VDEDSIRETF GA-CGEITEV RFAMDR-ESG EFKGFGHIEF
NCLPi           EASEKQEGCT TVFVGNLSWD IDEETLRQTF GE-CGTICSV RFATDR-ETG EFRGYGHVEF
NCLEa           APSEKPEGCL TIFIGNLSWQ IDEDSVREAF QE-CGEITTV RFAEDK-ETG DFRGFGHVEF
NCLTn           QPSEKPPGCL TVFVGNLSWE IDEESLRAAF AE-CGEITQV RFATDR-ETQ EFKGFGHIEF
NCLTan          EPTEKPEGCL SVFVGNLSWS IDEESLRAAF AE-CGEISQV RFATDR-ETQ EFKGYGHIEF
NCLSr           EPTPKPPGCT TVFVGNLSWS IDEDSLREAF AH-CGEIASV RFAMDR-ETQ EFKGFGHVEF
NCLAg           EPSQKEEGCI TVFIGNLSWD VDEDAIRQAF GD-CGEINQV RFATDR-ETG EFKGFGHIEF
NCLGo           VPSEKTEGTT TIFVGNMSFD IDEETLRGAL EETCGEITSV RFAMDR-DTG NFKGFGHVEF
NCLChd          ETSEKPAGCK TVFCGNLSWN VDEDTLRETF AE-CGEISSI RFATDR-ETG EFKGFGHIEF
NCLCt1          GPSEKPEGCN TVFCGNLSWN VDEDSLRAVF QD-CGEITQI RFAEDK-ETG EFKGFGHIEF
NCLCd           PTSEKPPGCT TVFVGNLSFS IDEPTLREAF AE-CGEISEV RFAEDR-ETG EFKGFGHIEF
NCLDf           EPSEKQEGTL TVFVGNLSFN IDEETLKEAF AD-CGEITQV RFAEDK-ETG QFKGFGHIEF
NCLCa           QPSEKPEGCT TVFVGNLSWQ IDEETLRAAF AD-CGEIAQV RFAEDR-ETG DFKGFGHVEF
NCLCn           QPSEKPEGCT TVFVGNLSFN IDEDTLREAF AE-CGEISQI RFATDR-ETG DFRGYGHMEF
NCLAsu          KEPGCTT--L FMGNLS---- ---WNVDEDS VREFFKG-CG EIAGVRFSED RE-TGEFKGF
NCLDbGSO104     KTPGSVT--V FVANLS---- ---FSIDEET VRDTFKD-CG TITAIRFGED RD-TGAFKGY
4
NCLPbr          GQQPKPDGCT TVFIGNLSFQ ADEYAISQVF ED-CGEIVSV RICTDR-DTG KSRGFGYVEF
NCLP            AAAPEYDRKR SVFVGNMPFD AQEQSLREHF AS-CGNVESV RIVRDP-YEN MGKGFGYVLF
NCLAt1          DGGDEKKIFV KGFDASLSED DIKNTLREHF SS-CGEIKNV SVPIDR-DTG NSKGIAYLEF
NCLAt2          EGSQSRTIYV RGFSSSLGED EIKKELRSHF SK-CGEVTRV HVPTDR-ETG ASRGFAYIDL
Nsr1pSc         FGDTPSEPSD TLFLGNLSFN ADRDAIFELF AK-HGEVVSV RIPTHP-ETE QPKGFGYVQF
NCLMc           PVNPDAEGSC CIFIGNVGYN TTKEALWDFF AN-YGNITDL RV-AHD-MDG NPRGFAHCEF
NCLSl           GGDKPEQKST TVFVGNLSYT TTEKSLEKFF SG-CGAIKAV RIAKME--DG KLKGFAHVEF
NCLBs           -------STS TVFVGNLPYN ADENNLTDFF SD-CGSIKQV RIAHDQ-D-G NKRGFAHVEF
NCLCsu          GSRPKAAPNK QIVVRNLSFN ATEESIRGLF EE-CGDIQEV RMPVFE-DSG KFKGQAFIEF
NCLHs           --STWSGESK TLVLSNLSYS ATEETLQEVF EK-ATFIKVP ---QNQ--NG KSKGYAFIEF
NCLCr           ---------A TLFMGGLSYT ATEDDVYNAF AG-VAEPVRV RIATDR-DSG EPKGFGHAEF
NCLDlu          YSMTVPPGVT KLFVGNLSFK VTEAALRDAF AP-CGHVTAV RLGMDKEDPT KFAGWAHVEF
5
NCLNsa          PPSEKPEGCM SVFIGNLSWN IDEETIRSAF AS-CGEVMSV RFATDR-ETG EFRGFGHVDF
NCLNg           PPSEKPEGCM SVFIGNLSWN IDEETIRSAF AS-CGEVTSV RFATDR-ETG EFRGFGHVDF
NCLTmi          DKKDKPEGCT TVFIGNLSWD VDEDAVRAAF GE-CGEIASV RFATDR-ETG DFKGFGHVEF
NCLGt           EPKPKPDGCT TIFMGNLSWD VDEDTIRSFF AD-CGEVVNV RFATDR-ETG DFKGFGHVQF
NCLG            EQRPKPAGCV SLFMGNLSWE VTEEEVRGLF AD-CGEVTQV RFATDR-ETG EFKGFGHVQF
6
NCLCh           ---EKPEGCD TVFVGNLSYD VDEPTVRDFF KD-CGAIQTI RFAEDR-ETG EFRGFGHVQF
NCLO            AVGEKPEGCT TVFVGNLAFS VDEGSVIDAF KD-CGEIAAT RFITDR-ETG DFKGMGFVEF
NCLTc           GPSEKQEGCM TVFIGNLSWD VDEDAIRQTF ES-CGEITQI RFAEDR-ETG RFKGFGHVEF
NCLTv           GPSEKMAGCM TVFIGNLSWD VDEETIRSIF EG-CGTISQI RFAEDR-ETG RFKGFGHVEF
NCLTs           GPSEKMAGCM TVFIGNLSWD VDEETIRSIF EG-CGTISQI RFAEDR-ETG RFKGFGHVEF
```

```
                  ....|....| ....|....| ....|....| ....|....| ....|....| ....|....|
                      1025       1035       1045       1055       1065       1075
1
NCLSco1       AETESTDLAV A--LAGTYVM DRPLRVDFAN ERKDRGFGGG GG-------- ---RGGGRGG
NCLSd         ADTESTDAAV A--LAGTYVM DRPLRVDFAN ERKDRGFGGG G--------- ---RGGGRGG
NCLSj         AETESTDAAV A--LAGTYVM DRPLRVDFAN ERRDRGFGGG G--------- ---RGGGX--
NCLSme        ADTESTDKAV A--LAGTYVM DRPLRVDFAN ERRDRGFGGG G--------- ---RGGGRGG
NCLDc         VETESTDKAV A--MAGTYVM DRALRVDFAN ERKSFGGGGG GG-------- ---RGGGRGG
NCLTm         VESESTDKAV E--LAGTYVM DRPLRVDYAN DRKKFGGGGG GGGFGG---- ---GGRGRGG
NCLCm         VDTESTDLAV K--MAGEDVM GRAIRVDYAN SRNHGEGRGG GG-------- ---RGGGRGR
NCLTo1        VETEATDKAV A--LAGTYVM DRPIRVDYAN ERRGGAVSVE VA---L---- ---EGAEVAV
NCLTw         ADTESTDKAV E--MAGTEIM GRAVRVDYAN ERX------- ---------- ----------
NCLTa         AETESTDKAV L--MAGTYVM DRALRVDYAN DRRGGAGGGG GG-------- ---GGRGGGR
NCLFs1        TDTEATDKAI A--MAGTDIL GRQVRVDYAN DKKAGGGGGG FGG------- ---GGGRSND
2
NCLEs         AETEATDKAV A--LAGTDML GRAVRVDYAN DRRGGGGGGG GFGGGRGGG- ---RGGGRG-
NCLNp         AESSATDAAV K--LAGTEIL GRPVRVDYAN DRKPSGGGGF G--------- ---RGGGRG-
NCLFk         GETEATDEAL K--LIGTEIL GRAVRVDYAN DRRQSFGGEG GGRGRGGGR- ---GGGGRGG
NCLPf         AETEATDAAV K--LAGTDIL GRAVRVDFAN DADNPLAVAE AGVVAT-EV- ---AGVAAV-
NCLPh         AESHSTDAAI K--LAGTDIL GRAVRVDFAN DRRQSFGGGR QSFGG--GG- ---RGGGRG-
NCLPa         AETEATDAAV K--LAGTDII GRAVRVDFAN DRRQSFGGGG G--------- ---RGGGRG-
NCLPd         AETEATDAAV K--IAGTDIC GRPVRVDFAN DRRQSFGSGG GGRGGFGGG- ---RGGGRG-
3
NCLPsp        SATEATDEAV K--MAGTMVA GRAIRVDYAA VKQKREFGAG GGGG------ ---RGGG--G
NCLPi         EETEATDKAV G--MAGQDVM GRAIRVDFAA VRQN-SFGGG GGRG------ ---GGRG--G
NCLEa         ANTDSTTQAV A--MAGTDIM GRAVRVDYAQ DRYS--GGGG GGGG------ ---RGGG-RG
NCLTn         AETEATDKAI A--MAGTDIV GRQVRVDFAA DRRQSFG--- GGGG------ ---RGGGRGG
NCLTan        TETAATDAAM Q--LAGTDIC GRQVRVDYAA DKRKSFGGAG GGGG------ ---RGGGRGG
NCLSr         VESDSCDEAM K--MIGTDIA GRAVRVDYAA DRKPPGGG-F GGGG------ ---RGGG---
NCLAg         CATESTDAAV K--MAGTEIM GRAVRVDYAN DRRNNAG--- GGGG------ ---RGGGGGG
NCLGo         ATTESTDIAV TK-MQGVEVM GRALRVDYAK DRKSFGGGGG RFG------- ---GGGGRGG
NCLChd        FSSDSTDAAI A--MAGTDIM GRAVRVDYAA DKR--AGGG- -FGG------ ---GRG---G
NCLCt1        AETEATDAAV A--LAGTDIM GRAVRVDYAG DKRKQAGGGR GFGG------ ---RGG---G
NCLCd         VDSASTDEAL K--LVGTEIM GRGVRIDYAA DKRG-EGGGG GRGR------ ---GGGRGRG
NCLDf         AATESTDAAV K--LAGTDIM GRAVRVDYAN DRRS-NGGGR GFG------- ---GGG--RG
NCLCa         VNSESTEAAV K--MAGTDIM GRPCRVDYAA DKRK-QNGGF GGGR------ ---GGG--RG
NCLCn         VNSESTDLAV Q--MAGTDIM GRACRVDFAA DKR--AGGGF GGGR------ ---GGGG-RG
NCLAsu        GHVDFVNTES CDKGVL-LSG QELMGRA-IR LDFANPRPSG GGGAGGRGGG RG--------
NCLDbGSO104   AFMDFEETEA TDKAVA-MAG TEIMGRA-VR VDHTTGASNT PGRT------ ----------
4
NCLPbr        TSTDAVDAAM K--KAGTDLA GRQVRVDFAA ARGEGAGDRA PRGG------ ---RGGGRGG
NCLP          EERVSVERAI G--LHDSEFA KRKLRVFRCL KSGEQPGDKG GRGGS----- ---RGGGRGG
NCLAt1        S--EGKEKAL E-LNGSDMGG GFYLVVDEPR PRGDSSGGGG FGRGNG---- ------RFGS
NCLAt2        T--SGFDEAL Q-LSGSEIGG GN-IHVEESR PR-DSDEGRS SNRAPA---- ------RGAP
Nsr1pSc       SNMEDAKKAL D-ALQGEYID NRPVRLDFSS PRPNNDGGRG GSRG------ -------FGG
NCLMc         STPEEAKASL --AAHGERVE GRALRIDLSA PRANRGGDRG GNRGG----- -------FGG
NCLSl         EEAESTQQAV A-LNGK-DLD GRDVKVDISE KLKNKRAEGG S--------W EDRRGGRGGR
NCLBs         ESTDSADAAI K--LNGKELE GRELKIDFAG EKRSGGSSRG GRG------- -------RGG
NCLCsu        DSVESATKAV E--YNNTEVD GRTVWIDFAL PRNSAGGRGG GRGGRG---- ---AFGGRGG
NCLHs         ASFEDAKEAL N-SCNKREIE GRAIRLELQG PRGSPNARSQ PSKTLFVKGL SEDTTEETLK
NCLCr         AD-VETARAA KNALFGVEIA GRKVRLDFAP PRNNSPGGGR GGRGGGRGGG FGGRGGGRGG
NCLDlu        AS-HADAQKG -VALNGLDLL GRSIRIDPAN PSAGGGGRGG GGRDGGRGG- -----GGRGG
5
NCLNsa        ADESAPEAAV Q--MAGTPVM GREIRVDYAQ PRPPREGGFG GGG------- ---RGGGRGG
NCLNg         ADETAPEAAV Q--MAGTPVM GREIRVDYAQ PRPPREGGFG GGG------- ---RGGGRGG
NCLTmi        VDGEGVDAAI D--MAGTEIA GRPVRVDYAA PRAPRESFGG GGG------- ---GGGGRGG
NCLGt         AESSATDLAV AK--GGEFVA GRAIRVDFAE DRKPPGSAG- ---------- ---SAGGGGG
NCLG          SEEEATEKGI AK--AGEFVA GRAIRLDYAE DKKTQGGAG- ---------- ---GAGGRGS
6
NCLCh         CDTMSTDEAV KLN--GQAMC GRNARIDYAP D--------- ----KR---- ---GGGGRG-
NCLO          ADTESADKAV KMGQEGAELM GRAMRIDYAK SGGV------ -SGGRD---- ---GGGGRGG
NCLTc         ADTMSTDKAV AMA--GTDVV GRPIRVDFAA VKQG--WSPR --QGGS---- ---PKGGRGD
NCLTv         EETEATDAAV ALA--GQDVC GRPIRVDFAA VKERKSFGDR SPQGGR---- ---GGGGRGG
NCLTs         EETEATDAAV ALA--GQDVC GRPIRVDFAA VKERKSFGDR SPQGGR---- ---GGGGRGG
```

```
                ....|....| ....|....| ....|....| ....|....| ....|....| ....|....|
                    1085       1095       1105       1115       1125       1135
1
NCLSco1         RGGRGGYMGG GGG------- ---------- ---------- ---------- -GGRGRGGFG
NCLSd           RGGRGGYMGG GGG------- ---------- ---------- ---------- -GGRGRGGFG
NCLSj           ---------- ---------- ---------- ---------- ---------- ----------
NCLSme          RGGRGGYMGG GGG------- ---------- ---------- ---------- -G-RGRGGRG
NCLDc           GGGRGGYMGG GGG------- ---------- ---------- ---------- -GGGFGGGRG
NCLTm           GRGRGGYMGG GGG------- ---------- ---------- ---------- -GR---GGRG
NCLCm           -----GYMGG GGG------- ---------- ---------- ---------- -----RGGRG
NCLTo1          VADEAVVDEA GAG------- ---------- ---------- ---------- ----------
NCLTw           ---------- ---------- ---------- ---------- ---------- ----------
NCLTa           GGGRGGYMGG GGGRD----- ---------- ---------- ---------- -GGRGRGGRG
NCLFs1          RGGRGGGRGR GRG------- ---------- ---------- ---------- ----------
2
NCLEs           ---GRGYMSG GGGGY----- ---------- ---------- ---------- -GRGGGGGYG
NCLNp           ---GGRGGRG GGG------- ---------- ---------- ---------- -RGRG-----
NCLFk           FGGGGRFGSG DRG------- ---------- ---------- ---------- -RGGGGGGRS
NCLPf           ---EAEVASE AAT------- ---------- ---------- ---------- -EAAAVADEV
NCLPh           ---GGRFGSA DRG------- ---------- ---------- ---------- -RGGGRGGYG
NCLPa           ---GSRFGSG DRG------- ---------- ---------- ---------- -RGGGRGGFG
NCLPd           ---GSRFGSG DRG------- ---------- ---------- ---------- -RGGGRGGFG
3
NCLPsp          GRGGGRGGG- GGGFG----- ---------- ---------- ---------- ----GAGRGG
NCLPi           GRGGGRGGGR GGGRG----- ---------- ---------- ---------- ----GGGRSG
NCLEa           GGRGGGGGGF GRGRG----- ---------- ---------- ---------- -GDRGGGRGG
NCLTn           RSPGGFG-GG GRGR------ ---------- ---------- ---------- --GRGGGRDG
NCLTan          RSPGGRSPGR GYGR------ ---------- ---------- ---------- --GRGGDRGG
NCLSr           ---------R GGGR------ ---------- ---------- ---------- --GRGG--YG
NCLAg           R--GGRGYMG GGG------- ---------- ---------- ---------- --GRGGGRGG
NCLGo           X--------- ---------- ---------- ---------- ---------- ----------
NCLChd          GRGGGRGGG- ---------- ---------- ---------- ---------- ---RGGGRGG
NCLCt1          GGFGGRGGGG GRGY------ ---------- ---------- ---------- --MRGGGGGG
NCLCd           GGRGRSDGGF GRG------- ---------- ---------- ---------- ---RGGGRGG
NCLDf           RGRGFGGGRN GGG------- ---------- ---------- ---------- ---FGGGRNG
NCLCa           GGRG-YGGGR GGGSR----- ---------- ---------- ---------- -GFGGGGRGG
NCLCn           GGRGGYGGGR GGG------- ---------- ---------- ---------- ---YGRGRGG
NCLAsu          -RGGDRF--- ---------- ---------- ----GGDRGG GRGGRG---- ----------
NCLDbGSO104     -PGRTPGRSF TPSEKEEGCK QVFIGNLSFN VDEDTIRGAF KDCGTI---- ----------
4
NCLPbr          GRGGGRGAPR GRG------- ---------- ---------- ---------- ----------
NCLP            AGGRGGGRGG SRD------- ---------- ---------- ---------- ------GGGG
NCLAt1          GGGRGRDGGR G-RFG----- ---------- ---------- ---------- -SGGGRGRDG
NCLAt2          RGRHSDRAPR GGRFS----- ---------- ---------- ---------- -DRAPRGRHS
Nsr1pSc         -RGGGR---- GGNRG----- ---------- ---------- ---------- --FGGRG--G
NCLMc           DRNGGNRGGF GGDRG----- ---------- ---------- ---------- -GFGGRGGFG
NCLSl           GGFRGGDRGG FRGGFRGGDR GGF------- ---------- ---------- ----RGGSRG
NCLBs           RGRPRRY--- ---------- ---------- ---------- ---------- ----------
NCLCsu          RGGGRGGAQS AAK------- ---------- ---------- ---------- ----------
NCLHs           ESFDGSVRAR IVTDRETGSS KGFGFVDFNS EEDAKAAKEA MEDGEIDGNK VTLDWAKPKG
NCLCr           FGGRGGGRGG FGGRG----- ---------- ---------- ---------- -GGFGGRGGG
NCLDlu          TPGRTPG--- --GRG----- ---------- ---------- ---------- -GTPGGRGGS
5
NCLNsa          GRGGGRSFGG GEGAG----- ---------- ---------- ---------- -SEEGAGSGG
NCLNg           GRGGGRSFGG G--------- ---------- ---------- ---------- ---GGGGFGG
NCLTmi          GRGGGR---- ---------- ---------- ---------- ---------- ----GGGRGG
NCLGt           GGRGGGRGGF GGG------- ---------- ---------- ---------- -------RGG
NCLG            FGSAGG--GR GGG------- ---------- ---------- ---------- -------RGG
6
NCLCh           ---------- ---------- ---------- ---------- ---------- ----------
NCLO            GGGRGGGG-- ---------- ---------- ---------- ---------- ----------
NCLTc           YGGKGDYGGK GGG------- ---------- ---------- ---------- ----------
NCLTv           FGGGGRGGGG RGG------- ---------- ---------- ---------- ----------
NCLTs           FGGGGRGGGG RGG------- ---------- ---------- ---------- ----------
```

```
                   ....|....| ....|....| ....|....| ....|....| ....|....| ....|....|
                       1145       1155       1165       1175       1185       1195
1
NCLSco1            GGRGGGRGGG RGGGRDFG-- -------AGG NKRSGGIAAF SGKKMTFD-- ----------
NCLSd              GGRGGGRGGG RGGGRDFG-- -------AGG NKRSGGIAAF SGKKMTFD-- ----------
NCLSj              ---------- ---------- ---------- ---------- ---------- ----------
NCLSme             GGRGGGFGGG RGGGRGGN-- -------SFG AKKSGSIAAF SGNKITFD-- ----------
NCLDc              GGRGGGRGGG RGGGRGGGP- -----TSTFS AKKSGGIAAF AGNKVTFD-- ----------
NCLTm              GR--GGRGGG RGGGRGGG-- --------FG AKKTGGIAAF SGNKITFD-- ----------
NCLCm              GR--GGRGGG RGGGRGRGG- -----FDPAK AKRSGGIAAF SGTKMTFD-- ----------
NCLTo1             ---------- ---------- --------TC QVAVDAVAE- ---------- ----------
NCLTw              ---------- ---------- ---------- ---------- ---------- ----------
NCLTa              GGRGGGRGG- RDGGRGGG-- -----RGMMG AKR--GIAEF AGTKMTFD-- ----------
NCLFs1             ----GDSGGR GGRGRGGG-- ------NSFN AKKTGGITAF AGKKISFD-- ----------
2
NCLEs              RGGGGGYGRG GGGRGGGGRG -----KSPGF AKRTGGIAEF SGNKITFD-- ----------
NCLNp              -------GGR GFSPANNTL- ---------K AKKSGGIAGF AGKKITFD-- ----------
NCLFk              YGSSGDRGGR GGRGGGGGGR -----GGNSL HKASGGIADF AGKKITF--- ----------
NCLPf              A-TEADEEAA AAVVAAPFT- ---------R NRAASQIS-- PGRK------ ----------
NCLPh              ----GGRGRG GGRGGDSNS- ---------L HKKSGGIADF AGKKITFD-- ----------
NCLPa              G-RGGFGGRG GGRDGGGST- ---------L HKKSGGIASF AGKKITXXXX SGGIASFAGK
NCLPd              ---SGDRGRG GGR-GGGSS- ---------L HKKSGGIADF AGKKITFD-- ----------
3
NCLPsp             GRG-----AS PTGRGG---- -----ASNPR SKNAGAISEF KGNKITFD-- ----------
NCLPi              GGGGYGGGGR SGGRGG---- -----ADPAR SRNNGSISGF AGKKITFD-- ----------
NCLEa              GGGGFGRGGG RGGGRGGDFG RGRGGASSSG SKQHGSIAEF KGNKITFD-- ----------
NCLTn              GR---GRGGG RDGGRG---- ---RGGNSFG AKKHGSISEF KGNKITFD-- ----------
NCLTan             GGGYGGRSGG RSGGRGG--- --GRGTDAFA AKKHGSISEF KGNKITFD-- ----------
NCLSr              GASPGGRGGG RGGGRG---- ---GGASAFG AKKHGSISEF KGTKVTFD-- ----------
NCLAg              GRG-GGRGGG RGGGRG---- ---GGGGGFG AKKHGSIAAF QGNKITFD-- ----------
NCLGo              ---------- ---------- ---------- ---------- ---------- ----------
NCLChd             GRGRGGFGGG GRGGGG---- -----G-YEK KSFG---AEF SGNRTTFD-- ----------
NCLCt1             GRG-GGRGGG GRGGG----- -------FSK PSFAKKSTEF AGTKITFD-- ----------
NCLCd              GGGFGRGRGG GRDSGG---- -----GGNYN PKKTGGISEF KGNKITFD-- ----------
NCLDf              G-GFGRGRGG GRGGGR---- -----GNMTS AKKSGGIAEF AGNKITFD-- ----------
NCLCa              SRGYGGGGGG RDSYS----- ---------S AKKKGGITEF AGNKITFD-- ----------
NCLCn              --GYGRGRGG GRGEG----- ---------S AR-TGGIAAF AGNKITFD-- ----------
NCLAsu             ---------- ---------- GGGRGGG--- -RGGGRGG-- ---GRGDNSR TFSMKKTGGI
NCLDbGSO104        ---------- -------TSI RFGEDRETGE FKGFGHIE-- ---FEETEAT DKAVAMAG--
4
NCLPbr             ---------- ---------- --------HP RGRGSGVGAF SGQKTKF--- ----------
NCLP               MGGGGGRGGR GRGDAADGQ- ---------K RPRSEGREEP KGKRQHAAY- ----------
NCLAt1             GRGRFGSGGG RGSDRGRG-- --------RP SFTP------ QGKKTTFGDE ----------
NCLAt2             DRG---APRG RFSTRGRGP- -------SKP SVMES----S KGTKTVFNDE E---------
Nsr1pSc            ARGG---RGG FRPSGSGAN- -------TAP LGRSRNTASF AGSKKTFD-- ----------
NCLMc              GRGGFGSRGG SGRGGRGGF- -------SKP PAGKGNIAEY QGNKMKF--- ----------
NCLSl              GFRGGDRGGS RGGFRG-GDR GG--SRGGFR GGDRGGFRGR GGFRGNRD-- ----------
NCLBs              ---------- ---------- ---------- ---------- ---------- ----------
NCLCsu             ---------- ---------- ---------- AAKDGTVQEF KGTSKTFDSD SE--------
NCLHs              EGGFGGRGGG RGGFGGRGGG RG--GRGGFG GRGRGGFGGR GGFRGGRGGG GDHKPQGKKT
NCLCr              RGGFGGRGGG RGGGRGGGRG -----GSFTA AVSKGAIQEF AGNKMTFD-- ----------
NCLDlu             RG--GGRSGG RGSDRAKG-- ---------- ---AGSIQEF SGKKMSFDD- ----------
5
NCLNsa             GRGGGRGGGR GGGRGGGET- -------GRS TRTEGAFRHL RETRRHSTTE QMAVGDVVGA
NCLNg              GRGGGRGGGR GGGRGGGRD- -------GAI NKNRGSIQAF AGNKKTFDD- ----------
NCLTmi             GRGGRDGGGR GGGRGGGR-- ---------- CVCAGGV--- ---------- ----------
NCLGt              GG-RGGGRG- GGRGGFGG-- --------TP NKNKGSIQES TGKKITFDD- ----------
NCLG               GAARGGAFGS GGRGGSAST- --------TP NRNRGSIQES QGKKISFDDD E---------
6
NCLCh              ---------- ---------- ---------- ---------- ---------- ----------
NCLO               ---------- ---------- ---------- ---------- ---------- ----------
NCLTc              --------FK NGGDRGRST- -----SPKPV NKNKGAIMAG SGKKITFD-- ----------
NCLTv              --------GG GFGARDRST- -----SPKPV NKNKGSIVAG TGKKITF--- ----------
NCLTs              --------GG GFGARDRST- -----SPKPV NKNKGSIVAG TGKKITF--- ----------
```

```
               ....|....| ....|....| ....|....| ....|....| ....|....| ....|....|
                   1205       1215       1225       1235       1245       1255
1
NCLSco1        ---------- ---------- ---------- ---------- ---------- ----......
NCLSd          ---------- ---------- ---------- ---------- ---------- ----......
NCLSj          ---------- ---------- ---------- ---------- ---------- ----......
NCLSme         ---------- ---------- ---------- ---------- ---------- ----......
NCLDc          ---------- ---------- ---------- ---------- ---------- ----......
NCLTm          ---------- ---------- ---------- ---------- ---------- ----......
NCLCm          ---------- ---------- ---------- ---------- ---------- ----......
NCLTo1         ---------- ---------- ---------- ---------- ---------- ----......
NCLTw          ---------- ---------- ---------- ---------- ---------- ----......
NCLTa          ---------- ---------- ---------- ---------- ---------- ----......
NCLFs1         ---------- ---------- ---------- ---------- ---------- ----......
2
NCLEs          ---------- ---------- ---------- ---------- ---------- ----......
NCLNp          ---------- ---------- ---------- ---------- ---------- ----......
NCLFk          ---------- ---------- ---------- ---------- ---------- ----......
NCLPf          ---------- ---------- ---------- ---------- ---------- ----......
NCLPh          ---------- ---------- ---------- ---------- ---------- ----......
NCLPa          KX-------- ---------- ---------- ---------- ---------- ----......
NCLPd          ---------- ---------- ---------- ---------- ---------- ----......
3
NCLPsp         ---------- ---------- ---------- ---------- ---------- ----......
NCLPi          ---------- ---------- ---------- ---------- ---------- ----......
NCLEa          ---------- ---------- ---------- ---------- ---------- ----......
NCLTn          ---------- ---------- ---------- ---------- ---------- ----......
NCLTan         ---------- ---------- ---------- ---------- ---------- ----......
NCLSr          ---------- ---------- ---------- ---------- ---------- ----......
NCLAg          ---------- ---------- ---------- ---------- ---------- ----......
NCLGo          ---------- ---------- ---------- ---------- ---------- ----......
NCLChd         ---------- ---------- ---------- ---------- ---------- ----......
NCLCt1         ---------- ---------- ---------- ---------- ---------- ----......
NCLCd          ---------- ---------- ---------- ---------- ---------- ----......
NCLDf          ---------- ---------- ---------- ---------- ---------- ----......
NCLCa          ---------- ---------- ---------- ---------- ---------- ----......
NCLCn          ---------- ---------- ---------- ---------- ---------- ----......
NCLAsu         ADFKGNKITF ---------- ---------- ---------- ---------- ----------
NCLDbGSO104    TDIMGRAVRV DFAGGRRGDG GGRGGGRGGG RGGGRGGGRG GGFGGGRGGG RGRGGGRGGS
4
NCLPbr         ---------- ---------- ---------- ---------- ---------- ----......
NCLP           ---------- ---------- ---------- ---------- ---------- ----......
NCLAt1         ---------- ---------- ---------- ---------- ---------- ----......
NCLAt2         ---------- ---------- ---------- ---------- ---------- ----......
Nsr1pSc        ---------- ---------- ---------- ---------- ---------- ----......
NCLMc          ---------- ---------- ---------- ---------- ---------- ----......
NCLSl          ---------- ---------- ---------- ---------- ---------- ----......
NCLBs          ---------- ---------- ---------- ---------- ---------- ----......
NCLCsu         ---------- ---------- ---------- ---------- ---------- ----......
NCLHs          KFE------- ---------- ---------- ---------- ---------- ----......
NCLCr          ---------- ---------- ---------- ---------- ---------- ----......
NCLDlu         ---------- ---------- ---------- ---------- ---------- ----......
5
NCLNsa         DRCGGKGGER EGKREGEGAG RTWRKGARAE RAHAAGVPRG GWGEEEGRVQ AAVA......
NCLNg          ---------- ---------- ---------- ---------- ---------- ----......
NCLTmi         ---------- ---------- ---------- ---------- ---------- ----......
NCLGt          ---------- ---------- ---------- ---------- ---------- ----......
NCLG           ---------- ---------- ---------- ---------- ---------- ----......
6
NCLCh          ---------- ---------- ---------- ---------- ---------- ----......
NCLO           ---------- ---------- ---------- ---------- ---------- ----......
NCLTc          ---------- ---------- ---------- ---------- ---------- ----......
NCLTv          ---------- ---------- ---------- ---------- ---------- ----......
NCLTs          ---------- ---------- ---------- ---------- ---------- ----......
```

```
             ....|....| ....|....| .
                1265       1275
1
NCLSco1      .......... .......... .
NCLSd        .......... .......... .
NCLSj        .......... .......... .
NCLSme       .......... .......... .
NCLDc        .......... .......... .
NCLTm        .......... .......... .
NCLCm        .......... .......... .
NCLTo1       .......... .......... .
NCLTw        .......... .......... .
NCLTa        .......... .......... .
NCLFs1       .......... .......... .
2
NCLEs        .......... .......... .
NCLNp        .......... .......... .
NCLFk        .......... .......... .
NCLPf        .......... .......... .
NCLPh        .......... .......... .
NCLPa        .......... .......... .
NCLPd        .......... .......... .
3
NCLPsp       .......... .......... .
NCLPi        .......... .......... .
NCLEa        .......... .......... .
NCLTn        .......... .......... .
NCLTan       .......... .......... .
NCLSr        .......... .......... .
NCLAg        .......... .......... .
NCLGo        .......... .......... .
NCLChd       .......... .......... .
NCLCt1       .......... .......... .
NCLCd        .......... .......... .
NCLDf        .......... .......... .
NCLCa        .......... .......... .
NCLCn        .......... .......... .
NCLAsu       ---------- ---------- -
NCLDbGSO104  SFGAKKSGSI AAFQGNKITF D
4
NCLPbr       .......... .......... .
NCLP         .......... .......... .
NCLAt1       .......... .......... .
NCLAt2       .......... .......... .
Nsr1pSc      .......... .......... .
NCLMc        .......... .......... .
NCLSl        .......... .......... .
NCLBs        .......... .......... .
NCLCsu       .......... .......... .
NCLHs        .......... .......... .
NCLCr        .......... .......... .
NCLDlu       .......... .......... .
5
NCLNsa       .......... .......... .
NCLNg        .......... .......... .
NCLTmi       .......... .......... .
NCLGt        .......... .......... .
NCLG         .......... .......... .
6
NCLCh        .......... .......... .
NCLO         .......... .......... .
NCLTc        .......... .......... .
NCLTv        .......... .......... .
NCLTs        .......... .......... .
```
